# Genomic insights into evolutionary journey of the porcine endogenous retroviruses

**DOI:** 10.1101/431858

**Authors:** Yicong Chen, Mingyue Chen, Xiaoyan Duan, Jie Cui

**Affiliations:** CAS Key Laboratory of Special Pathogens and Biosafety, Center for Emerging Infectious Diseases, Wuhan Institute of Virology, Chinese Academy of Sciences, Wuhan 430071, China.; University of Chinese Academy of Sciences, Beijing 100049, China.

**Keywords:** porcine endogenous retrovirus, genomic rearrangement, cross-species transmission, evolution, origin, recombination

## Abstract

**Background:** Xenotransplantation may overcome significant shortage of human allotransplant. Porcine organs are considered favorable for xenotransplantation duo to similar size and function to human organ. However, porcine endogenous retroviruses (PERVs) are potential infectious agents during xenotransplantation as they are able to infect and horizontally transfer among human cells. Furthermore, PERVs can be endogenized in pig genomes and are transmitted genetically in a Mendelian fashion. Here, we depict a complex evolutionary history of modern PERVs.

**Results:** We *in silico* mined 142 mammalian genomes and 14 pig genomes. This led to the documentation of 185 PERVs and a new viral cluster. Large-scale genomic alterations were found in most PERVs including many insertion-deletion events and which are suggestive of ancient origins, and pig genomes have been shaped by PERV-mediated genomic rearrangement during evolution. Notably, we found that lesser Egyptian jerboa and rock hyrax harbor ancestral PERV-related elements indicative of ancient cross-species transmission events from none-porcine species to pigs. A comprehensive analysis of these viral “fossils” suggested that recombination among none-porcine endogenous retroviruses led to the origination of PERVs.

**Conclusion:** For the first time, using large scale genomic mining, we decipher a complex evolutionary history for the PERVs. These new findings help us to understand the past of PERVs which pose the potential risk in clinical trials of xenotransplantation and provide novel insights into the origin and evolution of a human-infecting pathogen.

## Background

Xenotransplantation, the transplantation of tissues and organs from one species to another, may alleviate shortages of human donor organs(1, 2). Porcine organs are suitable for xenotransplantation due to their similar size and function to human organs, and the fact that pigs can be bred in large numbers(3). However, porcine endogenous retroviruses (PERVs) are able to infect and horizontally transfer among human cells, raising major concerns about the safety of xenotransplantation(3, 4), particularly given that retroviruses are often associated with severe infectious disease or oncogenic disease(3, 5-7).

PERVs are endogenous gammaretroviruses, and exist in the genomes of all pig strains(3, 8). The envelope (*env*) genes of three PERV classes (PERV-A, -B and -C) differ with respect to the receptor-binding domain (RBD)(9). Although there is no evidence of PERV transmission in patients receiving encapsulated pig islets(10-12), PERV-A and -B have been observed to infect both human cells and pig cells while PERV-C infects only pig cells(13). PERVs may also integrate into the human genome *in vitro*(14, 15). In pig cells, PERV-C can recombine with the *env* of PERV-A to produce A/C recombinants, which can infect human cells more efficiently(13). This increases the inherent risk in xenotransplantation and xenogeneic cell therapies.

While several studies have examined the evolutionary relationships between PERVs and other viruses, the origin and evolution of PERVs remains uncertain, particularly why they are unable to infect humans(16, 17). At least two species that belong to the same order as pigs, *Tayassu pecari* (of Eocene origin) and *Babyrousa babyrussa* (of Miocene origin) lack PERVs(18). However, the common warthog (*Phacochoerus africanus*) carries PERVs, suggesting that an ancestral porcine species carried PERVs(18). PERVs have two different types of long terminal repeat (LTRs), one with a 39-bp repeat structure in the U3 region, and the other without this repeat structure(19, 20). The 39-bp repeats carried by PERV-A and -B confer strong promoter activity and thus increase transcription(19, 20). However, the 39-bp repeat structure is absent in some PERV-A and all PERV-C. These PERVs thus have low transcriptional activity(19, 20). BLAST search analysis confirmed that the R and U5 regions of the PERV LTRs are highly conserved in the pig and mouse genomes (74–87% identity)(21). Indeed, the LTRs of PERV-A, -B and LTR-IS (a LTR family found solely in the mouse genome) have similar structures(21). The conserved LTR sequences found in pigs and mice might have originated from a common exogenous viral element, but evolved independently(21). Using genome mining with all available mammalian genomes, we clearly illustrate the evolutionary journey of the modern PERVs and reveal that the genesis of PERVs is much more complex than previously thought.

## Results

### *In silico* characterization of putative PERVs

Using previously reported PERV sequences as queries, we mined 14 pig genomes (Additional files 2: Table S1) available in GenBank, and showed a detailed genome-wide distribution of full-length PERVs (i.e. containing two LTRs). We initially compiled a PERV data set that included 185 putative PERVs (containing at least one LTR) (Additional files 2: Table S2). A total of 84 classified (30 PERV-A, 39 PERV-B and 15 PERV-C), 18 unclassified full-length PERVs (i.e., lacking of env gene) were retrieved from pig genomes. We identified 2–10 full-length PERVs in each of 12 other pig breeds, including Meishan, Goettingen, and Large White. We removed 19 previously classified PERV sequences that were low quality fragments (> 200 “N” bases). Hence, the final data set comprised 65 high quality classified PERVs (27 PERV-A, 29 PERV-B, and 9 PERV-C). Their viral genomic structures are summarized in Figure 1.

**Fig. 1.**
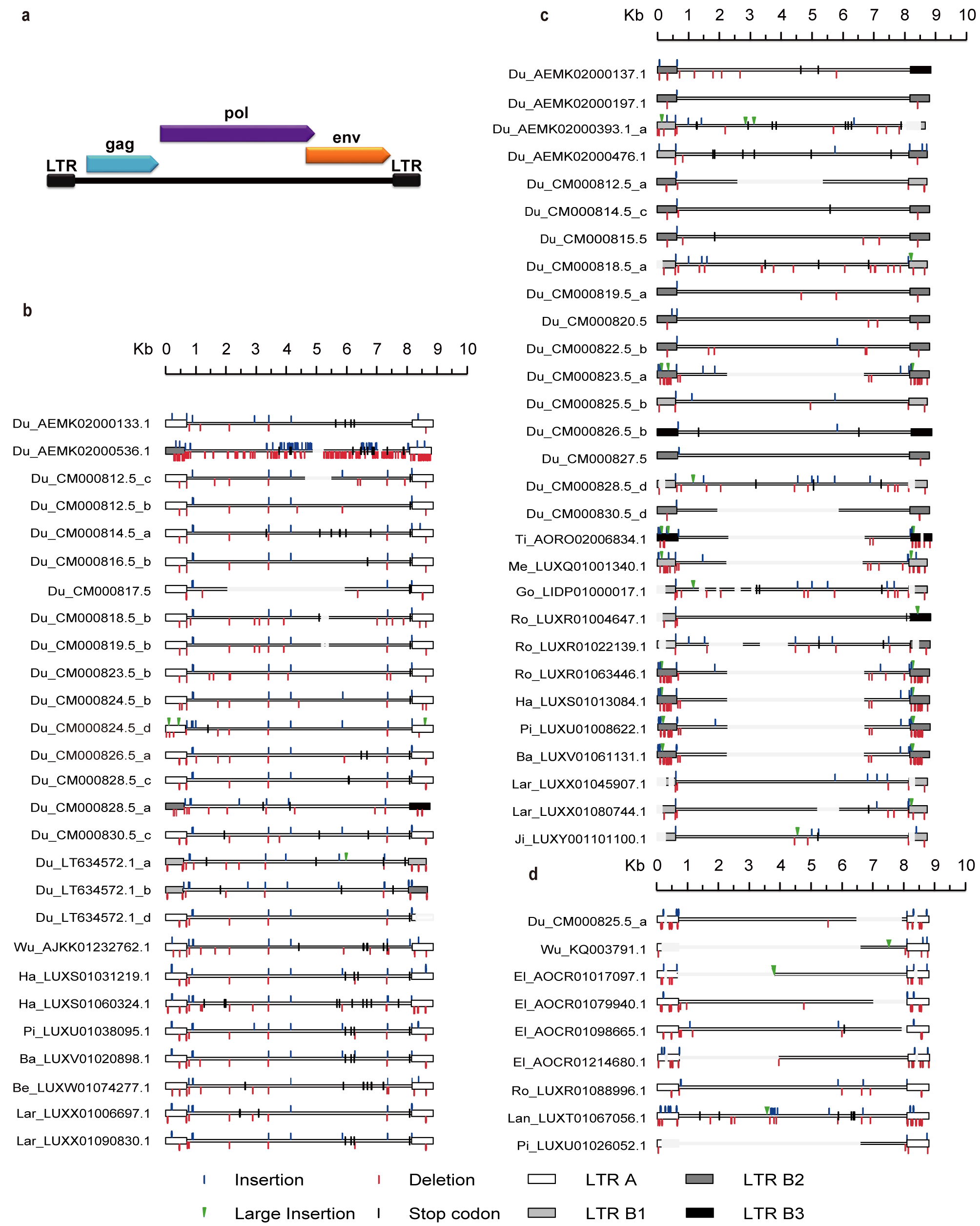
PERV proviruses in porcine genomes. (**A**) Genomic structure of PERVs, including *gag, pol* and *env* genes and LTRs. Proviruses of PERV-A and PERV-A like (**B**), PERV-B and PERV-B like (**C**) and PERV-C and PERV-C like (**D**) groups depicted based on reference PERV-A (accession number: AF435967.1), PERV-B (accession number: EU523109.1) and PERV-C (accession number: HQ536015.1). LTRs of PERVs were classified by the presence (type B) or absence (type A) of the 18 bp and 21 bp repeat structure. Type B LTRs were divided into three subtypes (LTR B1, LTR B2 and LTR B3). LTR A, B1, B2 and LTR B3 are presented in white, light gray, dark gray and black, respectively. Insertions and deletions (< 50 bp) are depicted with blue and red flags, respectively. Larger insertions (>50 bp) are labeled with green arrow. Large deletions (>50 bp) are shown without lines. Stop codons are showed with a black flag. (Du, Duroc pig; Wu, Wuzhishan pig; El, Ellegaard pig; Ti, Tibetan pig; Go, Goettingen pig; Me, Meishan pig; Ro, Rongchang pig; Ha, Hampshire pig; Lan, Landrace pig; Pi, Pietrain pig; Ba, Bamei pig; Be, Bekshire pig; Lar, LargeWhite pig; Ji, Jinhua pig).

Most of the PERVs exhibited large-scale genetic alterations induced by indels and stop codons (Fig. 1), suggestive of a relatively long evolutionary history. PERV LTRs were classified by the presence (LTR B) or absence (LTR A) of the 18 bp and 21 bp repeat structure reported previously(9, 19, 22). Three different type B LTRs in the PERV were identified, distinguished by the number of 18 bp and 21 bp repeat sequences: LTR B1 (two 18-bp and one 21-bp repeats), LTR B2 (three 18-bp and two 21-bp repeats), and LTR B3 (four 18 bp and three 21 bp repeats). Of the 65 high-quality PERVs we analyzed, we assigned 57, of which 32 (>55%) carried LTR A, 10 carried LTR B1, 13 carried LTR B2, and 2 carried LTR B3. LTR A was identified in PERV-A and -C, and LTR B1 was identified in PERV-A and -B. LTR B2 and LTR B3 were only identified in PERV-B. The remaining eight PERVs contained different types of 5’- and 3’- LTR which may reflect recombination between PERVs with different LTR types (Fig. 1). For example, we discovered one PERV (AEMK02000536.1) with LTR B2 at the 5’ end and LTR A at the 3’ end (Fig. 1).

### Potential genomic rearrangement via PERVs

Retrovirus integration creates a short duplication called target site duplication (TSD) flanking the LTR(23, 24). If chromosomal rearrangement through homologous recombination between distant proviruses occurred, the flanked TSDs should be different, as mentioned in a previous study of primate genomic rearrangement via ERVs (25). To identify pig genomic rearrangement via PERVs, we firstly constructed a maximum likelihood (ML) tree representing the 5’- and 3’- LTR sequences of full-length PERVs (Additional files 1: Fig. S1). The phylogenetic tree was divided into three large clusters (Additional files 1: Fig. S1), suggesting that three major integration events had occurred. We collected PERVs with 5’- and 3’- LTR sequences did not clustered together in the phylogenetic. Remarkably, 11 PERVs did not share the same TSD (4bp in length) (Table 1, Additional files 2: Table S3), thus these PERVs could reflect porcine genomic rearrangement during evolution.

**Table 1.** PERVs with different target site duplications (TSDs).

| Name | Accession number | Divergent<br>LTRs based<br>on structure<sup>*</sup> | Divergent<br>LTR based<br>on tree | Flanking TSD<sup>**</sup> |  |
| --- | --- | --- | --- | --- | --- |
|  |  |  |  | 5' | 3' |
| AEMK02000536.1 | AEMK02000536.1 | yes | yes | AGCC | CTTT |
| CM000818.5_a | CM000818.5 | no | yes | GTTC | CTTC |
| CM000826.5_c | CM000826.5 | no | no | ACCA | AATC |
| CM000828.5_d | CM000828.5 | no | yes | CCAC | CACC |
| KQ001967.1 | KQ001967.1 | no | yes | CCAC | CACC |
| LIDP01000017.1 | LIDP01000017.1 | no | yes | CCAC | CACC |
| LUXR01004647.1 | LUXR01004647.1 | yes | yes | GTTC | CTTC |
| LUXR01022139.1 | LUXR01022139.1 | yes | yes | CCAC | CACC |
| LUXX01045907.1 | LUXX01045907.1 | no | yes | CCAC | CACC |
| LUXX01080744.1 | LUXX01080744.1 | no | yes | GTTC | CTTC |
| LUXY01101100.1 | LUXY01101100.1 | no | yes | CCAC | CACC |

\*LTRs of PERVs comprise four types (LTR A, B1, B2, and B3). If two different types of LTRs are flanking the PERV, the LTRs are considered "divergent".
\*\*Only TSDs flanking the intact 5' and 3' LTRs sequences were analyzed

### Detection of PERV-related sequences in mammalian genomes

After screening 142 mammalian genomes (Additional files 2: Table S4) available on GenBank using tBLASTn and choosing three major proteins (Gag, Pol and Env) of PERVs as queries, a significant sequence (accession number: NW_004504334.1) in the genome of lesser Egyptian jerboa (*Jaculus jaculus*) that exhibited strong sequence similarity (for *gag* and *pol*: >75% nucleotide identity over 95% region; for *env*: >75% nucleotide identity over 55% region) to PERVs. Using this PERV-related sequence as query, three other possible PERV related sequences were identified in *J. jaculus* (accession number: NW_004504375.1, NW_004504378.1, and NW_004504445.1) with >85% nucleotide identity over 80% of the query sequence. The four PERV-related sequences identified in *J. jaculus* were designated as eJJRVs. These four significant hits located in large scaffolds > 5 Mb in length (Additional files 2: Table S5), with several host genes identified, indicated that the eJJRV sequences were relatively reliable.

We were only able to identify one pair of eJJRV LTRs. This full-length eJJRV (containing 2 LTRs) (accession number: NW_004504334.1) is annotated in Additional files 1: Fig. S2. The length of 3’- LTR of this eJJRV is 674 bp while 5’- LTR is 932 bp with a 258 bp insertion. We aligned eJJRV LTRs with PERV LTRs. The start of the U3 region and the end of the U5 region were distinct and not included in the alignment (Additional files 1: Fig. S3). The eJJRV LTRs included a repeat structure (three 18 bp and two 21 bp repeat sequences) in the U3 region, identical to that of the PERV LTR B2. Furthermore, 3’ LTR of eJJRV had high identity with LTR B2 of PERV (∼73%). Alignment analysis revealed a closer relationship between LTRs of the eJJRV and LTR B2 of PERVs (Additional files 1: Fig. S3). Notably, the alignment of the conserved R region supported a close evolutionary relationship between the eJJRV and PERVs (Fig. 2a). To highlight the similarity between PERVs and eJJRVs we generated pairwise alignments of eJJRV and PERV nucleotides using the full-length ERVs, and performed a sliding window analysis of these pairwise alignments (Fig. 2b)(26, 27). For comparison, we determined the similarity of HIV-1 provirus sequence to that of its closest relative (chimpanzee SIVcpz) (28, 29). Interestingly, *gag* and *pol* were more similar between eJJRV and PERV-A, -B and -C than HIV-1 and SIVcpz (Fig. 2b). Therefore, our results indicate that eJJRVs and PERVs are homologous. However, the RBD and the proline rich-region (PRR) of the surface subunit (SU) of *env* were dissimilar between eJJRV and PERV-A, -B and -C, as also seen in HIV and SIVcpz. As RBD determined host range(30-33), this observation suggests that eJJRVs and PERVs have distinct host ranges.

**Fig. 2.**
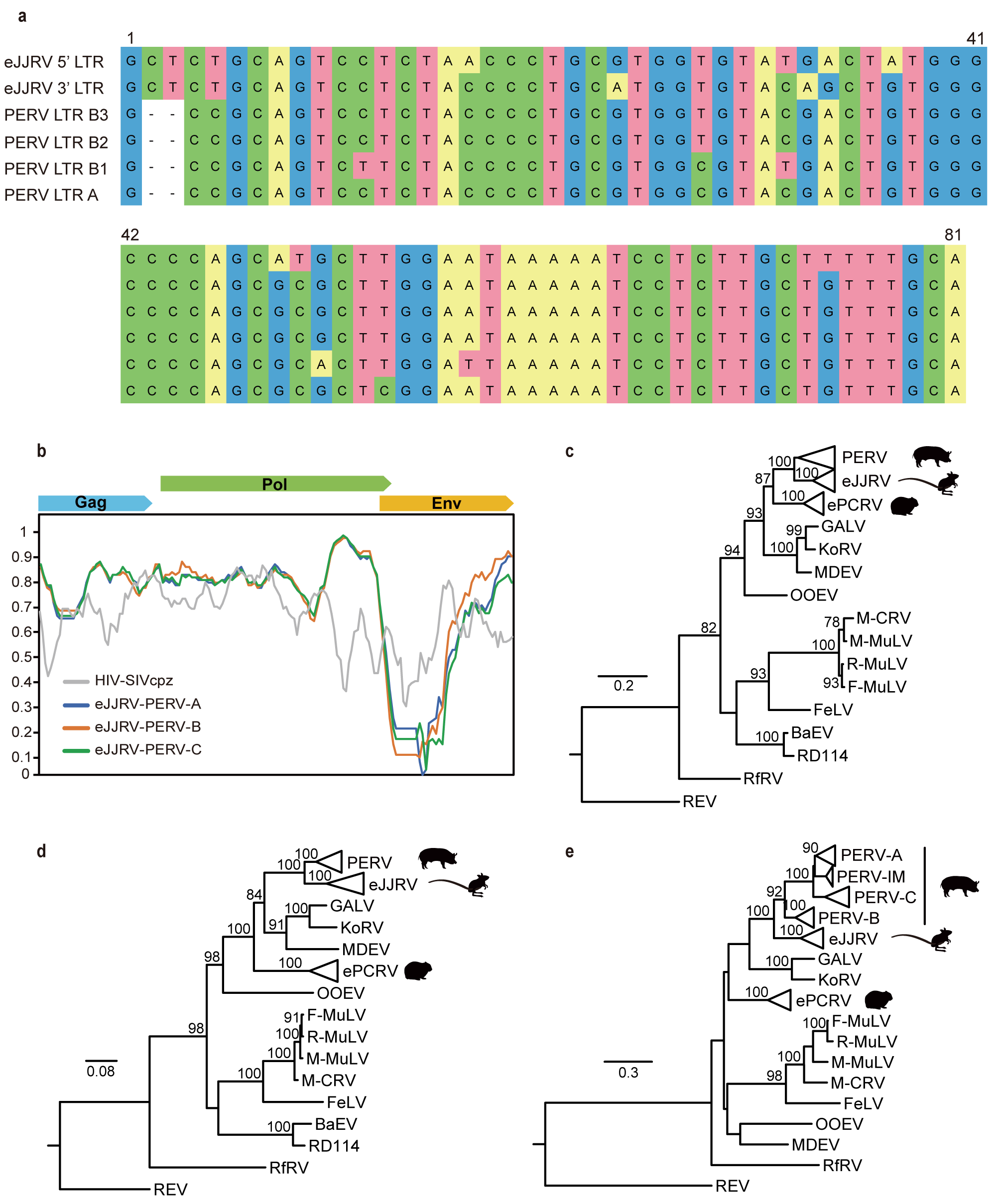
Comparison of PERVs, eJJRVs and ePCRVs. (**A**) Alignment of R region of LTR in eJJRV and PERVs. (**B**) Sliding window analysis of percent sequence identity along pairwise alignments of proviruses without LTRs. Phylogenetic trees of Gag (**C**), Pol (**D**) and Env (**E**) inferred using the amino acid sequences of PERVs, eJJRVs, ePCRVs and other representative gammaretroviruses (Table S7). Bootstrap values < 70% are not shown in phylogenetic trees. Trees were rooted to Reticuloendotheliosis virus (REV). The complete phylogenetic trees of Gag (**C**), Pol (**D**) and Env (**E**) are presented in additional files 1: Fig. S4-S6, respectively. All abbreviations can be found in Table S7.

We used the RBD amino acid sequences from PERV-A, -B and -C as queries to screen for homologous viral elements. The eight significant hits (>60% amino acid identity over 80% region) were obtained in rock hyrax (*Procavia canpensis*) of *Procaviidae*, and all eight hits flanking with genes located in large scaffolds >0.3 Mb in length (accession number: KN678690.1, KN676491.1, KN678005.1, KN677924.1, KN676905.1, KN676182.1, KN680906.1, and KN676638.1) (Additional files 2: Table S5). We examined the sequences flanking the eight hits (especially *pol*), and found that ERVs including these hits were endogenous gamma-retroviruses. These hits were therefore designated ePCRVs. We aligned the RBDs of PERVs and ePCRVs, and found that ePCRVs were highly similar to PERVs (Fig. 3). Pairwise comparisons revealed that ePCRV_1 and ePCRV_2 had a high identity to PERV-B (63%) but a low identity with PERV-A, -C and PERV-IM (40–43%). Therefore, the RBD of ePCRVs and PERV-B are homologous.

**Fig. 3.**
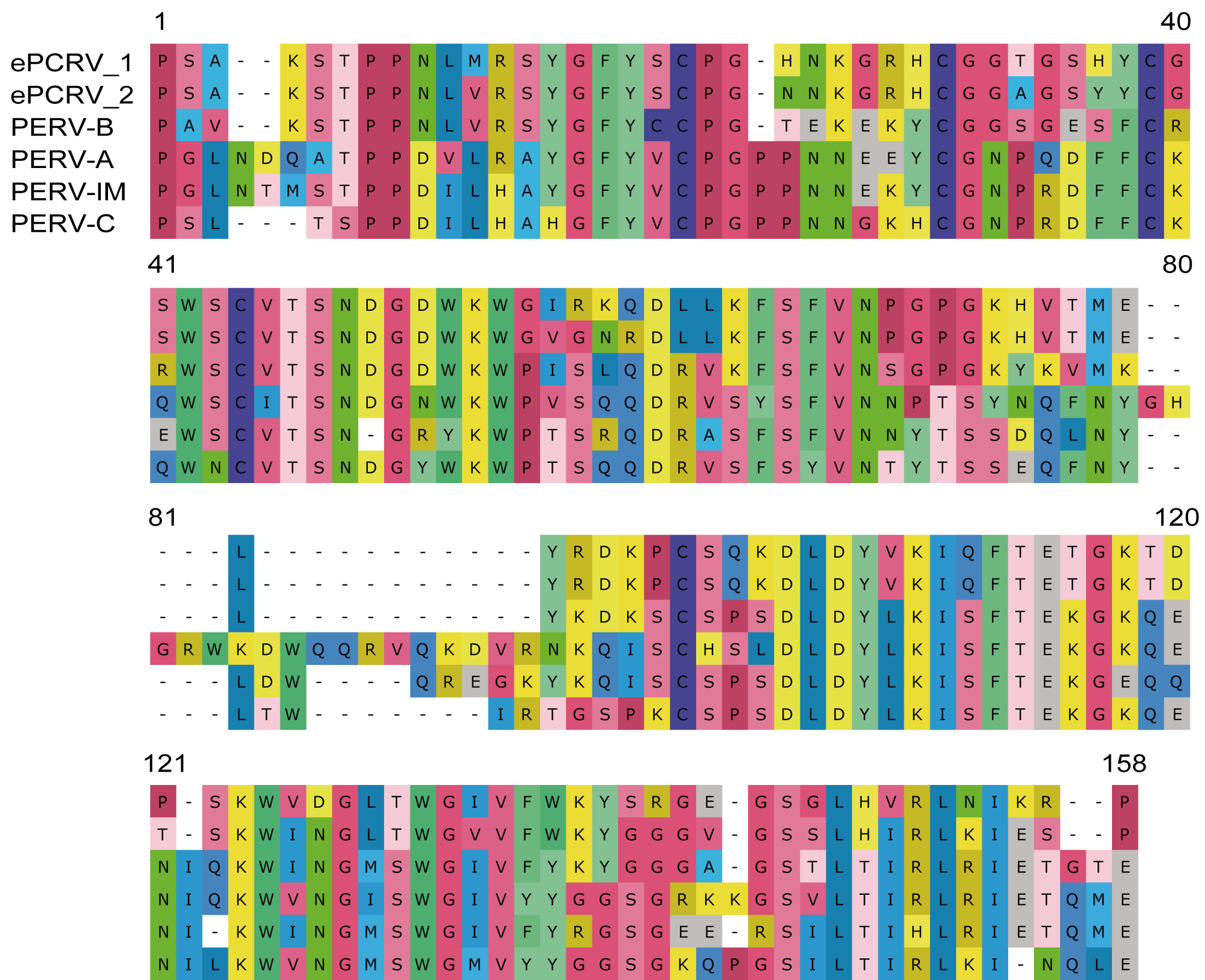
Amino acid sequence comparison of the RBD of PERVs and ePCRVs. Two ePCRVs (ePCRV_1(accession number: KN678005.1) and ePCRV_2 (accession number: KN678690.1)) were aligned to PERV-A, -B, -C, and the newly discovered PERV-IM.

To characterize the relationship within eJJRVs, ePCRVs and PERVs, we inferred phylogenetic trees of Gag, Pol and Env, first removing the variable RBD (Fig. 2c-e). Our maximum likelihood (ML) phylogenetic tree revealed that eJJRVs and PERVs clustered together with high bootstrap supports in three phylogenies (Fig. 2c-e), suggesting that they shared the most recent common ancestry. However, the Gag of ePCRVs clustered with eJJRVs and PERVs while Pol and Env (without RBD) of ePCRVs were distant related to eJJRVs and PERVs which might indicate recombinant events among ancient retroviruses (Fig. 2c-e)

Remarkably, a new lineage close to PERV-A and PERV-C was observed, named PERV-IM (as designation of PERV-intermediate type), and presented in all 14 pig genomes (Fig. 2e, Additional files 2: Table S2). The Env proteins of PERV-IMs showed relatively low similarity to PERV-A, -B, and -C, and they were clearly distinct in the RBD region (Fig. 3).

## Discussion

Using systematic large-scale genome mining, we analyzed the origin and evolution of the modern PERVs. We found homologous LTRs of PERVs (∼73% identity) in eight *Muroidea* species (*Mus caroli, M. pahari, M. musculus, M. spretus, Apodemus speciosus, A. sylvaticus, Rattus norvegicus*, and *Phodopus sungorus*). The coding genes (*gag, pol*, and *env*) near these homologous LTRs were identified (Additional files 2: Table S6). Also, ERVs in two *Muroidea* species (*M. musculus* and *R. norvegicus*) found in previous study were also presented(34). However, phylogenetic analysis of the Gag (>220 amino acids), Pol (>560 amino acids) and Env (>330 amino acids) sequences of these ERVs suggested that they were distantly related to the modern PERVs (Additional files 1: Fig. S7). We provided robust phylogenetic results that eJJRV shared the most recent ancestry with the PERVs (Fig. 2), and ePCRV also contributed the origin of Gag (Additional files 1: Fig. S7). Failure to detect any other eJJRV and PERV-related elements in the remaining rodent genome screening suggests that the virus was not vertically transmitted, suggesting ancient horizontal transmission occurred during evolution.

PERVs, eJJRVs, and ePCRVs integrated into *Suidae, Dipodidae*, and *Procaviidae*, respectively. Miocene (23 - 5.33 MY) *Suidae* fossils have been found in East Africa, Europe and Asia (http://fossilworks.org/?a=taxonInfo&taxon_no=42381). Miocene *Dipodidae* fossils have been found in North Africa, Europe and Asia (http://fossilworks.org/?a=taxonInfo&taxon_no=41695); Pliocene (5.3 - 2.59 MY) *Dipodidae* fossils have been identified in East Africa, thus suggesting that the *Dipodidae* may have spread to East Africa during the Miocene. Miocene *Procaviidae* fossils have been found in South of Africa and East Africa (http://fossilworks.org/?a=taxonInfo&taxon_no=43293). According to the current fossil records, the only shared region for Miocene *Dipodidae, Procaviidae* and *Suidae* fossils is East Africa. It is likely that ancestral PERV might be derived from two ancient retroviruses carried by *Dipodidae* and *Procaviidae*, denoted as JJRV and PCRV in the illustration (Fig. 4), in East Africa during the Miocene time. The ancestral PERV then split into different classes (PERV-A, -B, -IM and PERV-C), in which the PERV-B diverged earlier than other classes of PERVs (Fig. 4).

**Fig. 4.**
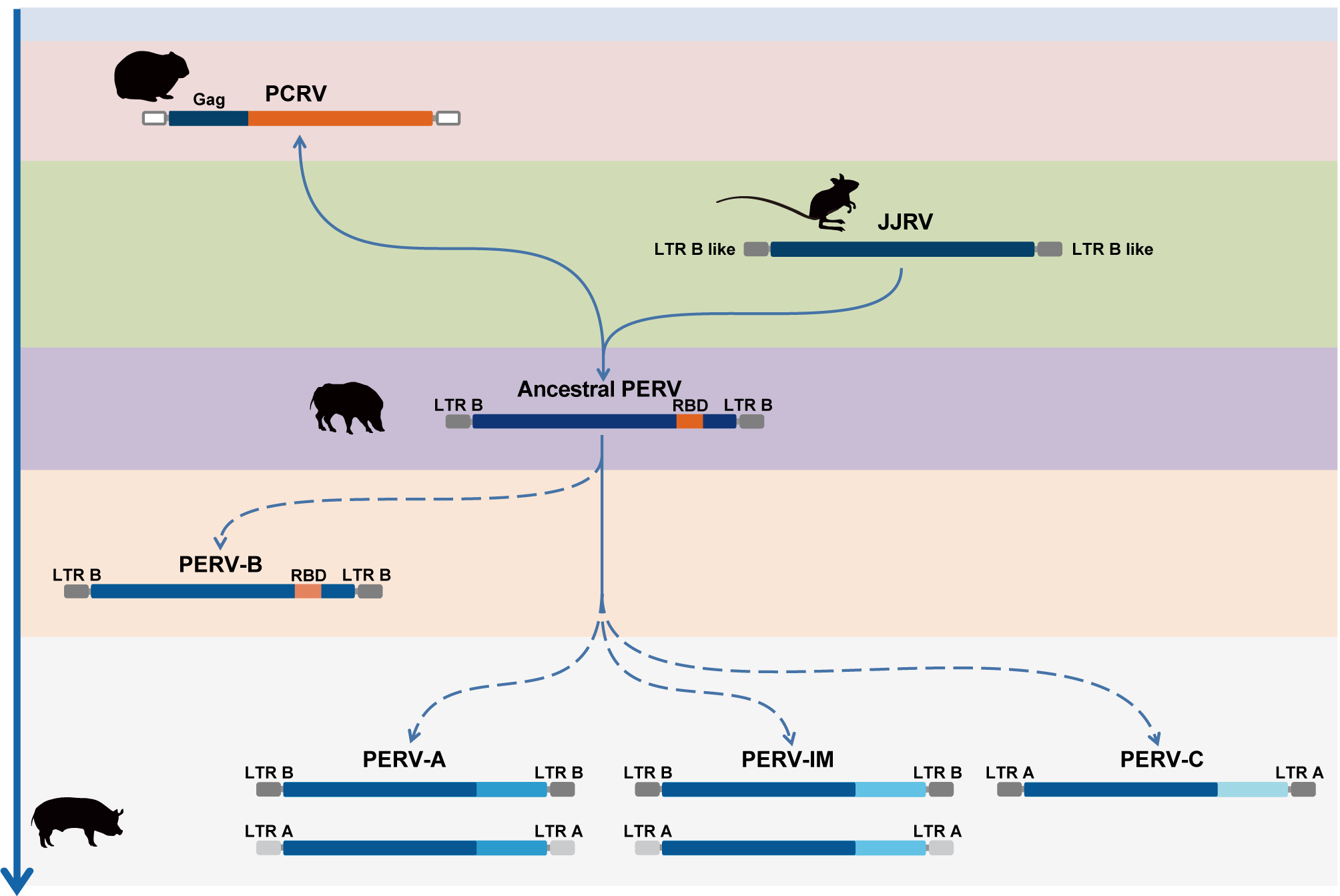
Scenario of genesis of PERVs. A schematic representation of PERV evolutionary history is shown, summarizing our hypothesis regarding the origin and evolution of PERVs. Arrow on the left indicates putative timeline of each type of ERVs. Different background colors illustrate putative evolution periods of modern PERVs. JJRV, PCRV, ancestral PERV represent the exogenous forms of the retroviruses from their hosts. Solid blue arrows show origination and dotted blue arrows show speciation. Abbreviation: RBD, receptor binding domain; LTR, long terminal repeat; Gag, group antigens gene.

We also discovered a new class of PERV named PERV-IM and they are presented in all pig genomes we scanned (Additional files 2: Table S1). It remains unknown whether PERV-IMs are able to infect human cells. However, it is possible that non-human trophic PERV was able to participate in recombination, leading to human trophic retroviruses(35, 36). The identification of the new PERVs is crucial for future assessment of pathogen transmission during xenotransplantation.

## Conclusions

For the first time, we decipher a complex evolutionary history for the PERVs. The ancestral PERV might be derived from two ancient retroviruses carried by non-porcine species. We also suggest that pig genomes have been shaped by PERVs, as specifically reflected by PERV-associated genomic rearrangement that have occurred during porcine evolution. In a word, modern PERVs have a complex evolutionary history than previously thought prior to their appearance in pigs.

## Materials and methods

### *In silico* identification of PERV and PERV-related proviruses

To identify PERV-related elements in *Sus scrofa*, tBLASTn (37) was used and the amino acid sequences of Gag, Pol and Env of 20 representative PERV proviruses (accession numbers: HQ536016.1, HQ536015.1, HQ536013.1, KC116220.1, AY570980.1, HQ540592.1, HQ536007.1, AX546209.1, AF435967.1, AY953542.1, HQ540591.1, AY099323.1, AJ133817.1, EU523109.1, EF133960.1, AY056035.1, AY099324.1, A66553.1, HQ536011.1, and HQ536009.1) were chosen as queries to screen the 14 pig genomes available in GenBank that were released before November 2017. A 50% identity over 50% of the match region was used to filter significant hits. It has been shown that PERVs harbor two LTR structures, one with and one without a repeat structure in the U3 region(9, 19). Using two typical LTRs as queries we extended flanking sequences of coding domains of PERVs to identify LTRs with BLASTn, and TSDs were used to define PERV boundaries. LTR lengths were defined as 100–1,000 bp. PERVs with at least one LTR and one coding gene were used in the evolutionary analysis.

To identify PERV-related proviruses in mammals, tBLASTn was used with the queries described above in 20 representative PERV proviruses to search the 142 mammal genomes available in GenBank as of November 2017. A 50% identity over 80% region was used to filter significant hits. LTRs were identified using LTR finder(38), LTRharvest (39) and BLASTn. LTR lengths were also defined as 100–1,000 bp.

### Detection of potential genomic rearrangement via PERVs

To search for proviruses involved in recombination and genomic rearrangement, we constructed a maximum likelihood (ML) tree of the 5’- and 3’- LTRs of full-length PERVs using PhyML 3.1(40) with GTR+I+Γ nucleotide substitution model. LTRs less than 250 bp were not considered. Sequence alignment was performed with MAFFT 7.222(41).

### Phylogenetic analyses

To determine the evolutionary relationship among PERVs, eJJRVs, ePCRVs and representative gammaretroviruses (Additional files 2: Table S7), phylogenetic trees were inferred using the amino acid sequences of full-length PERVs and PERVs with one LTR and at least one coding gene. All Gag, Pol and Env protein sequences (Dataset S1) were aligned in MAFFT 7.222 and confirmed manually in MEGA7(42). The phylogenetic history of these gammaretroviruses was then determined using the maximum likelihood (ML) method available in PhyML 3.1(40), incorporating 100 bootstrap replicates to assess node robustness. The best-fit JTT+Γ amino acid substitution model was selected for Gag, Pol and JTT+Γ+I for Env using the ProtTest 3.4.2(43).

## Supporting information

## Additional Files

**Additional files 1. Fig. S1.** Maximum likelihood (ML) tree of the 5’ and 3’ LTRs of all full-length PERVs. **Fig. S2.** Detailed descriptions of eJJRV genome. **Fig. S3.** The alignment of LTRs of PERVs and eJJRV. **Fig. S4.** The complete phylogenetic tree of Gag. Fig. **S5.** The complete phylogenetic tree of Pol. **Fig. S6.** The complete phylogenetic tree of Env. **Fig. S7.** The phylogenetic trees of Gag, Pol and Env of Muroidea ERVs, eJJRVs, ePCRVs, PERVs and representative retroviruses.

**Additional files 2. Table S1.** The information of pig, rodent, and rock hyrax genomes used for data mining. **Table S2.** The information of full-length and near full-length PERVs. **Table S3.** The recombination-related information of full-length PERVs shown in Fig. S1. **Table S4.** The information of 142 mammals used for PERV-like sequences mining. **Table S5.** The information of genes flanking the eJJRVs and ePCRVS. **Table S6.** The information of full-length and near full-length Muroidea ERVs. **Table S7**. The information of representative retroviruses used for phylogenetic analysis.

**Additional files 3. Data set S1.** The alignments used to build the phylogenetic trees of Gag, Pol and Env represented in Fig. S4, S5, and S6, respectively. **Data set S2.** The alignments used to build the phylogenetic trees of Gag, Pol and Env represented in Fig. S7.

## Declarations

### Ethics approval and consent to participate

Not applicable.

## Acknowledgement

Not applicable.

## Funding

J.C. is supported by National Natural Science Foundation of China under grant no. 31671324 and CAS Pioneer Hundred Talents Program.

## Availability of data and materials

The dataset supporting the conclusions of this article is described in the main text (and additional files)

## Author contributions

J.C. conceived and designed the research. Y.C. and M.C. conducted the analyses. X.D. participated in the analysis of full-length PERVs. J.C. supervised the whole project. All authors participated in the project discussion and manuscript preparation.

## Competing interests

The authors declare that they have no competing interests

## References

1. Ekser B, Cooper DKC, Tector AJ. The need for xenotransplantation as a source of organs and cells for clinical transplantation. International journal of surgery (London, England). 2015;23(Pt B):199–204.

2. Ekser B, Ezzelarab M, Hara H, van der Windt DJ, Wijkstrom M, Bottino R, et al. Clinical xenotransplantation: the next medical revolution? Lancet (London, England). 2012;379(9816):672–83.

3. Niu D, Wei HJ, Lin L, George H, Wang T, Lee IH, et al. Inactivation of porcine endogenous retrovirus in pigs using CRISPR-Cas9. Science (New York, NY). 2017;357(6357):1303–7.

4. Denner J. Paving the Path toward Porcine Organs for Transplantation. The New England journal of medicine. 2017;377(19):1891–3.

5. Wegman-Points LJ, Teoh-Fitzgerald ML, Mao G, Zhu Y, Fath MA, Spitz DR, et al. Retroviral-infection increases tumorigenic potential of MDA-MB-231 breast carcinoma cells by expanding an aldehyde dehydrogenase (ALDH1) positive stem-cell like population. Redox biology. 2014;2:847–54.

6. Denner J, Young PR. Koala retroviruses: characterization and impact on the life of koalas. Retrovirology. 2013;10:108.

7. Aiewsakun P, Katzourakis A. Marine origin of retroviruses in the early Palaeozoic Era. Nature communications. 2017;8:13954.

8. Denner J, Tonjes RR. Infection barriers to successful xenotransplantation focusing on porcine endogenous retroviruses. Clinical microbiology reviews. 2012;25(2):318–43.

9. Tonjes RR, Niebert M. Relative age of proviral porcine endogenous retrovirus sequences in Sus scrofa based on the molecular clock hypothesis. Journal of virology. 2003;77(22):12363–8.

10. Morozov VA, Wynyard S, Matsumoto S, Abalovich A, Denner J, Elliott R. No PERV transmission during a clinical trial of pig islet cell transplantation. Virus research. 2017;227:34–40.

11. Crossan C, Mourad NI, Smith K, Gianello P, Scobie L. Assessment of porcine endogenous retrovirus transmission across an alginate barrier used for the encapsulation of porcine islets. Xenotransplantation. 2018:e12409.

12. Wynyard S, Nathu D, Garkavenko O, Denner J, Elliott R. Microbiological safety of the first clinical pig islet xenotransplantation trial in New Zealand. Xenotransplantation. 2014;21(4):309–23.

13. Moalic Y, Blanchard Y, Félix H, Jestin A. Porcine Endogenous Retrovirus Integration Sites in the Human Genome: Features in Common with Those of Murine Leukemia Virus. Journal of virology. 2006;80(22):10980–8.

14. Czauderna F, Fischer N, Boller K, Kurth R, Tonjes RR. Establishment and characterization of molecular clones of porcine endogenous retroviruses replicating on human cells. Journal of virology. 2000;74(9):4028–38.

15. Patience C, Takeuchi Y, Weiss RA. Infection of human cells by an endogenous retrovirus of pigs. Nature medicine. 1997;3(3):282–6.

16. Li Z, Ping Y, Shengfu L, Hong B, Youping L, Yangzhi Z, et al. Phylogenetic relationship of porcine endogenous retrovirus (PERV) in Chinese pigs with some type C retroviruses. Virus research. 2004;105(2):167–73.

17. Cui J, Tachedjian G, Tachedjian M, Holmes EC, Zhang S, Wang LF. Identification of diverse groups of endogenous gammaretroviruses in mega-and microbats. The Journal of general virology. 2012;93(Pt 9):2037–45.

18. Niebert M, Tonjes RR. Evolutionary spread and recombination of porcine endogenous retroviruses in the suiformes. Journal of virology. 2005;79(1):649–54.

19. Scheef G, Fischer N, Krach U, Tonjes RR. The number of a U3 repeat box acting as an enhancer in long terminal repeats of polytropic replication-competent porcine endogenous retroviruses dynamically fluctuates during serial virus passages in human cells. Journal of virology. 2001;75(15):6933–40.

20. Wilson CA, Laeeq S, Ritzhaupt A, Colon-Moran W, Yoshimura FK. Sequence analysis of porcine endogenous retrovirus long terminal repeats and identification of transcriptional regulatory regions. Journal of virology. 2003;77(1):142–9.

21. Huh JW, Cho BW, Kim DS, Ha HS, Noh YN, Yi JM, et al. Long terminal repeats of porcine endogenous retroviruses in Sus scrofa. Archives of virology. 2007;152(12):2271– 6.

22. Niebert M, Kurth R, Tonjes RR. Retroviral safety: analyses of phylogeny, prevalence and polymorphisms of porcine endogenous retroviruses. Annals of transplantation. 2003;8(3):56–64.

23. Mayer J, Blomberg J, Seal RL. A revised nomenclature for transcribed human endogenous retroviral loci. Mobile DNA. 2011;2(1):7.

24. Johnson WE, Coffin JM. Constructing primate phylogenies from ancient retrovirus sequences. Proceedings of the National Academy of Sciences of the United States of America. 1999;96(18):10254–60.

25. Hughes JF, Coffin JM. Evidence for genomic rearrangements mediated by human endogenous retroviruses during primate evolution. Nature genetics. 2001;29(4):487–9.

26. Lole KS, Bollinger RC, Paranjape RS, Gadkari D, Kulkarni SS, Novak NG, et al. Full-length human immunodeficiency virus type 1 genomes from subtype C-infected seroconverters in India, with evidence of intersubtype recombination. Journal of virology. 1999;73(1):152–60.

27. Zhuo X, Feschotte C. Cross-Species Transmission and Differential Fate of an Endogenous Retrovirus in Three Mammal Lineages. PLoS pathogens. 2015;11(11):e1005279.

28. Martoglio B, Graf R, Dobberstein B. Signal peptide fragments of preprolactin and HIV-1 p-gp160 interact with calmodulin. The EMBO journal. 1997;16(22):6636–45.

29. Corbet S, Muller-Trutwin MC, Versmisse P, Delarue S, Ayouba A, Lewis J, et al. env sequences of simian immunodeficiency viruses from chimpanzees in Cameroon are strongly related to those of human immunodeficiency virus group N from the same geographic area. Journal of virology. 2000;74(1):529–34.

30. Argaw T, Wilson CA. Detailed mapping of determinants within the porcine endogenous retrovirus envelope surface unit identifies critical residues for human cell infection within the proline-rich region. Journal of virology. 2012;86(17):9096–104.

31. Watanabe R, Miyazawa T, Matsuura Y. Cell-binding properties of the envelope proteins of porcine endogenous retroviruses. Microbes and infection. 2005;7(4):658–65.

32. Denner J. Recombinant porcine endogenous retroviruses (PERV-A/C): a new risk for xenotransplantation? Archives of virology. 2008;153(8):1421–6.

33. Ericsson TA, Takeuchi Y, Templin C, Quinn G, Farhadian SF, Wood JC, et al. Identification of receptors for pig endogenous retrovirus. Proceedings of the National Academy of Sciences of the United States of America. 2003;100(11):6759–64.

34. Hayward A, Cornwallis CK, Jern P. Pan-vertebrate comparative genomics unmasks retrovirus macroevolution. Proceedings of the National Academy of Sciences of the United States of America. 2015;112(2):464–9.

35. Harrison I, Takeuchi Y, Bartosch B, Stoye JP. Determinants of high titer in recombinant porcine endogenous retroviruses. Journal of virology. 2004;78(24):13871–9.

36. Kaulitz D, Mihica D, Adlhoch C, Semaan M, Denner J. Improved pig donor screening including newly identified variants of porcine endogenous retrovirus-C (PERV-C). Archives of virology. 2013;158(2):341–8.

37. Altschul SF, Gish W, Miller W, Myers EW, Lipman DJ. Basic local alignment search tool. Journal of molecular biology. 1990;215(3):403–10.

38. Xu Z, Wang H. LTR_FINDER: an efficient tool for the prediction of full-length LTR retrotransposons. Nucleic acids research. 2007;35(Web Server issue):W265-8.

39. Ellinghaus D, Kurtz S, Willhoeft U. LTRharvest, an efficient and flexible software for de novo detection of LTR retrotransposons. BMC Bioinformatics. 2008.

40. Guindon S, Dufayard JF, Lefort V, Anisimova M, Hordijk W, Gascuel O. New algorithms and methods to estimate maximum-likelihood phylogenies: assessing the performance of PhyML 3.0. Systematic biology. 2010;59(3):307–21.

41. Katoh K, Standley DM. MAFFT multiple sequence alignment software version 7: improvements in performance and usability. Molecular biology and evolution. 2013;30(4):772–80.

42. Kumar S, Stecher G, Tamura K. MEGA7: Molecular Evolutionary Genetics Analysis Version 7.0 for Bigger Datasets. Molecular biology and evolution. 2016;33(7):1870–4.

43. Abascal F, Zardoya R, Posada D. ProtTest: selection of best-fit models of protein evolution. Bioinformatics (Oxford, England). 2005;21(9):2104–5.

