## Supplementary material for "Genomic insights into evolutionary journey of the porcine endogenous retroviruses"

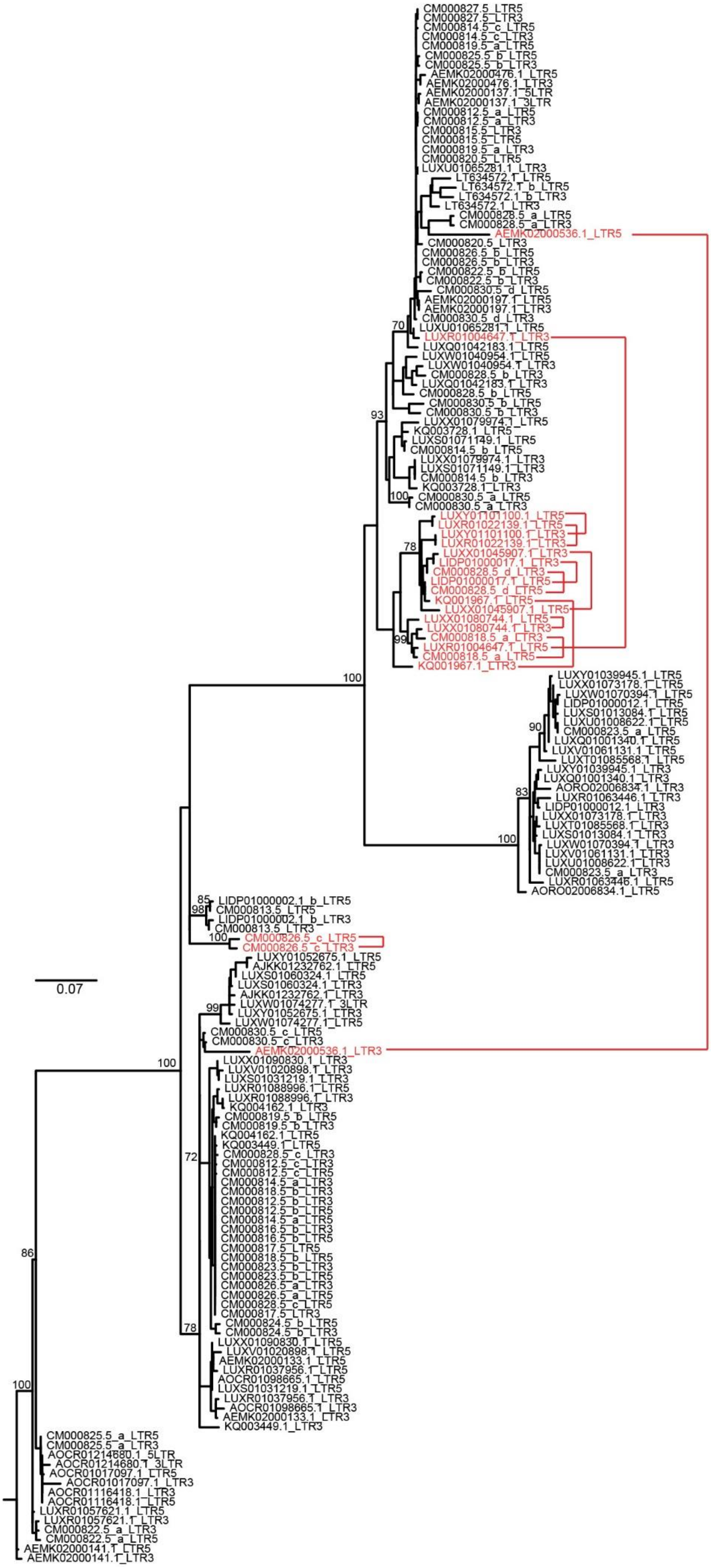
CM000827.5_LTR5
CM000827.5_LTR3
CM000814.5_c_LTR5
CM000814.5_c_LTR3
CM000819.5_a_LTR5
CM000825.5_b_LTR5
CM000825.5_b_LTR3
AEMK02000476.1_LTR5
AEMK02000476.1_LTR3
AEMK02000137.1_5LTR
AEMK02000137.1_3LTR
CM000812.5_a_LTR5
CM000812.5_a_LTR3
CM000815.5_LTR3
CM000815.5_LTR5
CM000819.5_a_LTR3
CM000820.5_LTR5
LUXU01065281.1_LTR3
LT634572.1_LTR5
LT634572.1_b_LTR5
LT634572.1_b_LTR3
LT634572.1_LTR3
CM000828.5_a_LTR5
CM000828.5_a_LTR3
AEMK02000536.1_LTR5
CM000820.5_LTR3
CM000826.5_b_LTR5
CM000826.5_b_LTR3
CM000822.5_b_LTR5
CM000822.5_b_LTR3
CM000830.5_d_LTR5
AEMK02000197.1_LTR5
AEMK02000197.1_LTR3
CM000830.5_d_LTR3
70
LUXU01065281.1_LTR5
LUXR01004647.1_LTR3
LUXQ01042183.1_LTR5
LUXW01040954.1_LTR5
LUXW01040954.1_LTR3
CM000828.5_b_LTR3
LUXQ01042183.1_LTR3
CM000828.5_b_LTR5
CM000830.5_b_LTR5
CM000830.5_b_LTR3
93
LUXX01079974.1_LTR5
KQ003728.1_LTR5
LUXS01071149.1_LTR5
CM000814.5_b_LTR5
LUXX01079974.1_LTR3
LUXS01071149.1_LTR3
CM000814.5_b_LTR3
KQ003728.1_LTR3
100
CM000830.5_a_LTR5
CM000830.5_a_LTR3
LUXY01101100.1_LTR5
LUXR01022139.1_LTR5
LUXY01101100.1_LTR3
LUXR01022139.1_LTR3
78
LUXX01045907.1_LTR3
LIDP01000017.1_LTR3
CM000828.5_d_LTR3
LIDP01000017.1_LTR5
CM000828.5_d_LTR5
KQ001967.1_LTR5
LUXX01045907.1_LTR5
LUXX01080744.1_LTR5
LUXX01080744.1_LTR3
99
CM000818.5_a_LTR3
LUXR01004647.1_LTR5
CM000818.5_a_LTR5
KQ001967.1_LTR3
100
LUXY01039945.1_LTR5
LUXX01073178.1_LTR5
LUXW01070394.1_LTR5
LIDP01000012.1_LTR5
LUXS01013084.1_LTR5
LUXU01008622.1_LTR5
90
CM000823.5_a_LTR5
LUXQ01001340.1_LTR5
LUXV01061131.1_LTR5
LUXT01085568.1_LTR5
LUXY01039945.1_LTR3
LUXQ01001340.1_LTR3
83
AORO02006834.1_LTR3
LUXR01063446.1_LTR3
LIDP01000012.1_LTR3
LUXX01073178.1_LTR3
LUXT01085568.1_LTR3
LUXS01013084.1_LTR3
100
LUXW01070394.1_LTR3
LUXV01061131.1_LTR3
LUXU01008622.1_LTR3
CM000823.5_a_LTR3
LUXR01063446.1_LTR5
AORO02006834.1_LTR5
85
LIDP01000002.1_b_LTR5
98
CM000813.5_LTR5
LIDP01000002.1_b_LTR3
CM000813.5_LTR3
100
CM000826.5_c_LTR5
CM000826.5_c_LTR3
LUXY01052675.1_LTR5
AJKK01232762.1_LTR5
LUXS01060324.1_LTR5
LUXS01060324.1_LTR3
AJKK01232762.1_LTR3
99
LUXW01074277.1_3LTR
LUXY01052675.1_LTR3
LUXW01074277.1_LTR5
0.07
CM000830.5_c_LTR5
CM000830.5_c_LTR3
AEMK02000536.1_LTR3
100
LUXX01090830.1_LTR3
LUXV01020898.1_LTR3
LUXS01031219.1_LTR3
LUXR01088996.1_LTR5
LUXR01088996.1_LTR3
KQ004162.1_LTR3
CM000819.5_b_LTR5
CM000819.5_b_LTR3
KQ004162.1_LTR5
KQ003449.1_LTR5
CM000828.5_c_LTR3
72
CM000812.5_c_LTR3
CM000812.5_c_LTR5
CM000814.5_a_LTR3
CM000818.5_b_LTR3
CM000812.5_b_LTR3
CM000812.5_b_LTR5
CM000814.5_a_LTR5
CM000816.5_b_LTR3
CM000816.5_b_LTR5
CM000817.5_LTR5
86
CM000818.5_b_LTR5
CM000823.5_b_LTR3
CM000823.5_b_LTR5
CM000826.5_a_LTR3
CM000826.5_a_LTR5
CM000828.5_c_LTR5
CM000817.5_LTR3
CM000824.5_b_LTR5
78
CM000824.5_b_LTR3
LUXX01090830.1_LTR5
LUXV01020898.1_LTR5
AEMK02000133.1_LTR5
LUXR01037956.1_LTR5
100
AOCR01098665.1_LTR5
LUXS01031219.1_LTR5
LUXR01037956.1_LTR3
AOCR01098665.1_LTR3
AEMK02000133.1_LTR3
KQ003449.1_LTR3
CM000825.5_a_LTR5
CM000825.5_a_LTR3
AOCR01214680.1_5LTR
AOCR01214680.1_3LTR
AOCR01017097.1_LTR5
AOCR01017097.1_LTR3
AOCR01116418.1_LTR3
AOCR01116418.1_LTR5
LUXR01057621.1_LTR5
LUXR01057621.1_LTR3
CM000822.5_a_LTR3
CM000822.5_a_LTR5
AEMK02000141.1_LTR5
AEMK02000141.1_LTR3

**Fig. S1 Maximum likelihood (ML) tree of the 5' and 3' LTRs of all full-length PERVs.** Bootstrap values lower than 70% are not shown in the phylogenetic tree. ML tree was constructed using PhyML with the GTR+I+Γ nucleotide substitution model. LTRs of PERVs with different target site duplication (TSD) flanking are labelled in red. Red line connect the pairwise LTRs with different TSD.

**eJJRV**

**TSD | 5′ LTR start**

GTAC**TG**AAAGGCCCAAGAAACCAGAAGCCTAGGAATACTATCATGCTCCAAAAGGAGCTTTAGCAGGGCCCTGCATACCT
CAAATTCCAGGCCCTACTCTCTGCTATAAAGTAAGTAAAAAGTCACTCTTCTTATTGTATCAGAGCCCGCCCTGGATATA
AAACTAGGGAAAAGGGCGTAAGACAGGCCTTTGAGGATGGGAAGTGAGTAATGGATTAAACCGCTTATGTGGCTTCTGTA
AACTGCTCGCACCATAAGCCCTAATAGCCTGTCTTAACCACACTTTGTCACCCCCACCCAAGAGCCCGCGCAAATGCTGA

**Large insertion**

CTGCCAGGGTTATAGGAAACTGATTGGTCCACAGAGTGTGGGCTTGAATAATTTAAAATGATTGGCTTAAAACTATGGAG
TGATAGGCTGGAGAGATGGCTTAGCAGTTAAGCACTTGCCTGTGAAGCCTAAGGACCCCGGTTCGAGGCTCAATTCCTCA
GGACCCACGTTAGCCAGATGCATAAGGGGGTGCACACATCTGGAGTTTGTTTGCAGTGGCTGGAGGCCCTGGCACGCCCA
TTCTCGCTCTCTCTATCTCTATCTGCCTGTTTCTCTCTCTGTCGCTCTCAAATAAATAAATAAATAAAATGAACAAAAAT
TTTTTTAAAAACTATGTAGTGGCATACACCGACTAGCCCACATTGCGCGGGCTCCTGATACTAAGAAATGATTGGTTTTA
ATTCACAGGCTTTGTTAGAAACCATATAAAAGCTGTCCCGTTCCTGCATTCAGGGCTCTGCAGTCCTCTAACCCTGCGTG
GTGTATGACTATGGGCCCCAGCATGCTTGGAATAAAAATCCTCTTGCTTTTTGCATCAAGACCATTTCTCGTGAGTGATT

**5′ LTR end |**

TGGGGTGTTGCCATTTCCTGGCAGAGTGTGGGGTCCTCGTTCTGGAGGTCTTA**CA**TTTGGGGGCTCATCCGGGATTTACG
ACCACCCCCTCATGCCCGAGAACTGACTTGGAGGTAAGGGGGACCCCCCCTGCAGTGAATGTGTGCTGGCCAGTGTTTCT
GCTCTGAGTATGTGTTTTCTGGGACACGCGCTTTCTGTTTTCAGTTTGTAGCCATCCACTCGGGCCGTAAAGACCAGAGG
ACTGTGATCAGCAGATGTGCTAGGAGGATCACAGGCTGCTGCCCTGGGGGATGCCCTGGGAGGTGAGAAGAGCCAGGGAC
ACCTGATGATCTTCTATTGTCAGATTGTCAGTCAGAGGACCAAGTTCTGTTGTTAGAGTGGAAGCTTCCCCTTCTGCGGA
TGTCTGATCCTTTCGCCTGCTTATGGAGGACATGGATGGGCTGCTGGTGTCTGGATCTGTTAGTCTCAGTTGTTGTGT

**| Gag Start**

GCATTGTTGTCCTTTGTGTCTTTGTCTATAGTTCTGTTATGGACAGATGGTGATGACTCCCCTTAGCTTGACTCTTGAC
CATTGGACTGAAGTTAAATCTAGGGCTCATAATTTGTCAGTTGAAGTTTAAAAGAGACTATGGCAGACCTTTTGTGCCTC
TGAATGGCCAACTTACAAGGTCGGGTGGCCATCAGAAGGAACCTTTAATTCTGAGACTATTCTGGCTGTTAAAGATATTA
TTTTTCAGAATGGACCAGGCTCTCATCCCGATAAAAGCCTTACATCCTCACATGGAAAAATTTGGTGGACAGCCCCCCAC
CATGGGTTAAGCCATGGTTGAACAGACCAAGGAAGCCAAGCTCCTGAGTCCTGGCCCTTGAAAATGAAAGAAAACCCTCT
CCGGAGAAAGTCAGACCTCCTCCTCCTCCTCGCATCTATCCCAAGATCGAGGAGCTCCTGGCTTGGCCAGAAACTCAGCC
TGCCCCTCCTCCTCCTTACCCAGCTCCGGGTGCCAGGGCAGGGCCCTCCAACCTTCCCAAGCCTCCGCCCTCCACCTTTC
CCATGCCTCCGCCTCTTGCCCCTCCCATGCCTCCGCCCCTTGCCCCTCCCGAAGCTCCGAGCTCCGGCGTGACTGCTCCG
CCCCTAGCCCCTCCCGAAGTTCCGCCCTCCACCTTTCCCAAGCCTCCGCCCTCTGCCCTTCCTGAGCCTCCACAGCCCTC
CGCCCCTACCAAACCTCTAGCAATGGGAGGACCAGCTGCAAGAATGTGGAGCTGGAGATATGCCACCCTGGAGGGAGCAG
ATGGGATTGTAACATTGCCGCTATGCACATCAGGTCCTCTCGTGCTGGGGGCCAACTACAGCCTCTCCAATATTGGCCTT
TCTCCTCTGCAGATCTTTATAACTGGAAAGCTAAGTACCCTCCTTTTTCAGAAGAACCCCGGCGCCTCACAGTGTTGATA
GAGTCTCTTATGTTTTCTCATCAGCCTACTTGGGATGATTGACAACAGCTGTTGCAGACACTCTTCACAACAGAGGAGCG
AGAGGGAATTCTAATAGAGGCCAGAAAGAATGTTCCTGGGGCCTAAGGGCAGCCCTCACAGCTGCAGCATGAGATAGACA
TGGGGTTCCCTCTGACTCACCCTGCCTGGGAATACAATACAGCTGAAGGTAGGCAGAGCTTAAAAATCTATTGCCAGGCT
CTGGTGGCGGATCTCCAAGATGCCTCCAGAAGGCCCCCCCGTTTGGCCAGGGTGAGAGAGATGATGCAGGGACCAACTGA
ATCCCCCTCAATGTTCCTTGAGAGGCTCATGGAGGCCTTTTGGAGGTTCACTCCCTTTGACCCCACCTCTGAAATCCAGA
AAGGCTCAGTAGCATTAGCCTTCATAGGATAGTCAGCTCCTGATATCAGAAAAAAAACTTCAGAGATTGGAAGGATTACA
GGAAGCTGAGTTGCATGATTTAGTAAAAGAGGCAGAAAAGGTATATTACAAAAGAGAGACAGAAGAGGAAAAGGAGCAAA

GGAAAGAGAGAGGGAAAGAAGAACGCAAGGAGAGACCAGAGAAAGAGAGAGAGGAAAGAAAAGATAGGCGCAATAGGCGA

CAAGAGAAAAATCTAACTAGGATCTTGGCCACAGTGGTGTAGAAGAAAGGAGAAAGGGAGAGAGATATTTAAAAATTTAG

GACAGGTCCTAGGCAGATAGGGAATTTGGGCAACATGACCCAACTTTATAGAGATCAATGTGCCTGTTGCAAGGAGAAAG

**Gag end |**

GGCACTAGGCAAGGGATTGCCCTAAGAAGAACAGCAAAGGACCTAGGGTTCTAGCTCTTGAAGAGGATGAAGATTAGGGA

AGATTGGGCTCGGAGCCCCTCCCTGAGCCCAGGGTAATTCTAAAGGTGGAGGGGAGGCCAGTCAAGTTCCTGGTTGATAC

**| Pol start**

CAGAGCAGAACATTCAGTACTCTTACAACCACTGGGAGGACTAAAAGATAAAAAATCCTGGGTGTTGGGCGCTACTGGAC

AATGGCAATATTCATGGACTACCCAAAGAACTGTTGACCTGGGAGTGGGACGGGTAACCCACTCATTCTTGGTCATCCCT

GAGTGCCCAGCGCCCCTCCTTCTTAGGGACCTGTTAACCAAGATGGGTGTTCAAATTTCCCTCAAATCAGAAATACTGGA

AGTGTCTGCCAGAGGTCAACCCATCACTGTGTTGACTCTCCAACTAGATGATGAGTATCGATTATACTCTTCCCCAGTAG

CCTAGTCAAGATCTAAAGCCCTGGTTAGAGCAGTTTCCCCAAGCCTGGGCAGAAACCATGGGCATGGGCTTGGCGAAACA

AGTACCCCCACAGGTCATTCAGCTAAAAGCCAGTGCTATGCCAGTATCAGTCAAACAATATCCCATGAGCTGAGAGGCTC

AAGAAGGAATTTGGCCCCATATTCAAAGACTGATTCAGCAAGGTATCCTGGTTCCAGTTCAGTCCCCCTGGAACACTCCC

TTACTGCCAGTTAGAAAGCCTGGGACTAATGATTACCACCCAGTACAGGACTTGAAAAAAATTAATAAAAGAGTACAGGA

CATACATCCAACTGTCCCAAACCCTTACAACCTCCTTAGTGCACTCCCACCCAAGCGGAGTTGATACACAGTGTTGGATC

TAAAAGATGCCTTCTTCTGCCTAAGATTACATTTGGACAGCCAATCTCTTTTTGCTTTTGAATGGAGAGACCCAGACACA

GGGAGAACTGGGTAGGTCAACTGGACTCACTTACCCTAAGGATTCAAGAACTCCCCAACCATCTTTGATAAAGCCCTACA

CAGGGATCTAGCCAACTTTAGGGTTCAGCACCCCCACCTAACCCTGCTGCAATACGTTGATGACTTACTCCTGGCAGGAG

CTACCCAGCAGAACTGTCTAAAAGGTATGAAGGCACAACTACTGGAGCTGACTGACCTTGGTTACTGAGCCTCAGCTAAA

AAGGCTCAAATATGCAAAAGAGAGGTAACGTACTTGGGGTACTCCTTGTGGGATGGGAAAAGATGGCTGGCAGAGGCACA

AAAGAGAACTGTAGTACAAATACTGGTCCCAACTATGGCCAGACAAGTCAGAGAGTTCCTGGGGACAGCTGGGTTCTGCA

GGCTATGGATCCTGGGGTTTGCAACCTTAGCAGCCCCCCTCTATTCACTAACTAAAGAGAAAGAGGGATTCACCTGGACC

TCCAAACACCAGAGGGTGTTTGATACCATCAAGAAAGCACTGCTAAGTGCACCTGCTCTAGCCCTCTCCGATGTGACTAA

ACCATTCACCCTTTATGTAGATGAGCATAAGGGTATAGCCCAGGGAGTCCTGACCCAATCTTTAGGACCATGGAGAAGAC

CTGTCGCCTACCTGTCAAAGAAACTTGACTCCGTAGCTAGTGGATGGCCTGTCTGTCTAAAATCCATAGTGGCAATGCCC

ACACTGATCAAAGATGCTGATAAACTAACTCTAGGGCAGAAGATAACTGTCATTGCCCCCCCATGCTCTGTAGAGCACTG

TCCAGCAGCCCCCAGACCAATGGATGACCAATGCCCAAATGGCACACTACCAGAGCCTGCTCCTCACTGACAGGGTAATG

TTCACTCCACCTGCCATCCTTAATCCCGCTACTCTTCTGCCTGAAGAAAATGATGAGCCAGTGACTCATGACTTTCATCA

ACTGCTGGTTGAGGAAACTGGGATCCGTAAGGACCTTACAGATGTCCCACTGACCAGAGGAACATTAACCTGGTTCACAG

ATGGGAGCAGTTATATGGTAGAAGGTAAGAGGATGGCTGGGGCAGCGATAGTAGATGGAACGGTCTGGGCCAGCAGTCTG

CCAGAAGGAACTGCAGCACAAAAGGCTGAGCTCGTGGCCCTCACACAGGCCTTATGATTAGCTGAAGGAAAGTCTGTGAA

TATCTACATGGACAGTAGGTATGCTTTTGCTACTGCACATGTGCATGGGGCCATCTATCAACAGAGAGGGCTGCTTACTT

CAGCAGGGAAAGAAGTTAAAAACAAAGAAGAAATACTGTGTTTGCTAGAAGCTTTACATCTACCCAAAAGACTGGCCATT

ATGCATTGTCCTGGCCACCAAAAAAGCCAAAGATTTTGTCTCCAAAGGGAACCAAATGGCTGACCAAATAGCCAAACAGG

CAGCCCAGGGGCTCAGCCTTTTGCCCATAATAGAAAAGCCAGAAAACCAAGAAAGTGGGCAATGGTATACCCCAGAGGAT

TGGCAAGAAGTCAGAAAATTAGCCCAGTTCTCTGAAACCCCGTAGGGGGCCTGTTTCACCTTAGAGGGAAAAGAGATTCT

ACCCCAAGTGAGGGGACTAGAATACGTCCAAGAAATACACTGCCTAACCCACCCAGGGGCCAAACATCTACAAAAACTAC

TACAAACGTCTTTTTACCATATTCTGAGGTTGCCAGAGATAGCTGACTCAGTATTCAAGCATTGTGTACCCTGCCAGATG

GTCAATGCTAATCCTTCAAAAATGCCTCCAGGTAAGAGACTGAGGGGAAGCAGCCCAGGCACTCACTGGGAAATGGACTT

CACTGAGGTAAAGCCAGCTAAGTGTGGTAACAAATACTTACTAGTTTTGGTAGACACTTTTTCAGGCTGGGTAGAAGCTT

ATACTACCAAGAAAGAGACCTCAACCGTGGTGGCTAAGAAAATACTGGAGGAAATTTTCCCAAGATTCGGAATACCTAAG

GTAATAGGGTCAGACAATGGTCCAGCTTTTGTTGCCCAAGTAGTCAGGGACTGGCCAAAATATTGGGGATTGATTGGAAA

TTACATTGTGCATACAGACCCCAAAGTTCAGGACAGGTAGAGAGGATGAATAGAACCATTAATGAGACCCTCACCAAACT

GACTGCAGAGACTGGCGCTAATGATGCGATAGCTCTCCTACCCCTTGTGCTCTTTAGGGTCAGAAACACCCCCGGACAGT

TTGGACTGGCCCCCTATGAATTGCTATACGGGGAGCCCCCTCCACTGATAGAAATAGCCTCTGTACATAGTGCTGATGTG

CCATTTTCCCAGCCTTTGTTCTCTAGGCTCAAGGTGCTCGAGTGGGTGAGATGATGAACGTGGAAGCGGCTCTGGGAGGC

CTACTCAGAAGAAGGAGACTTACAGGTCCCACCTCGCTTCCAAGTGGGAGATTCGGTCTATGTTAGACGCCACCACGCAG

GAAACCTTGATACTTGGTGGAAGGGCCCCTACCTTGTCCTACTGACCACACCGACAGCTGTGAAAGTTGAAGGAATACCC

**| Env start**

ACCTGGATCCATGCATCTCACGTCAAGCCTGCTCCGCCACATGGTCCTGAATGGCAAGCCAAGAGGACGGACAATCCCCT

**Pol end |**

TAAGCTTAGGCTTCGCCGTGTGGCTTCTCCTTCTCCAGCTAGGTGATTCCTCCAATCTCCATGCCCCCCAAAAGTTGACT

TGGCATGTACTTTCCCAGATGGGGGATATGATATGGACTGCAACTATGGAAGCACCCCCCTGGACTTGGTGGCCTGTCCT

GACCCTGGACATATGTGCCCTGGTAGCAGGGCTATAATCTTGGGATATCCCAGAACTCACTTCCGAGGAGGTCCCTAGAG

ATGTCCACTCTCATGTAGCTAAATGGGCACTCATACAAGAGGTCGCCTCCCAAGGCCAATAAATGGCAGATGACCCAGGG

TGCAGCTCTAGGCTAGCCCAACGTAGGATGCAGCATACTCCATTTTATGTCTGCCCCCGAACTTCTAAAAGAAGTTGTGG

AGGGGTAAAACAGTTCTACTGTAAATCCTGGGGTTGTGAGGCCACAGGCAATACTTACTGGCACCCAGCTTCCTCCTGGG

ACCTGATCACCATTCAGGATGGATTAGACGGGAAGAGAGCAGTTCCTCTCAACATTTCTTTCACCCCCAAGGTCGAGGGT

CCTGAGACTGGATAAAAGGAAAAACTTGGGGACTCAGGTTCTATATGACAGGATGGGCTAAGGGCCTAACCTTCACTATA

TGACTCAAGATAGAGCCTCCCACCCCAAGTTCAATAGGACCTAATAATGTTCTAGCAGACCAGGGACTCCCTAATGCGGC

ATTGCCTATGTCCCTACCCACAGTGGCAGCTGGGACCTGCCGCGTAACGACCACTAGAATCACTCCGACTAGCCCATTCT

TGGTAGCCCCACACAAGACAGGACAAAGGCTTTTAAACCTTATCCAGGGGGCCTTCCTCACCATAAATGCCACCAACCCC

AATGCTACCTCTGCCTGCTGGCTTTGTTTATCCTCAGGACCACCCAACTATGAGGGGGTGGCAAACGTGGGAGAGTTTAA

TGTTACTAAAGAACATAATAGACAGTGTTCATGGGGAATTAAAAACAAGCTGACTTTCCCAGACATCTCAGGAAGTGGGA

CATGTATAGGCTCCCCCGTCCCACCAACATCTCTGTAATAATACTCTGATCTATAATCAGACCTCAGACAACCAATATCT

AATGCCAGGATATAACAGGTGGTGGGCATGTGATACTGGGCTAACTCCCTGTGTTTCCCCCTCAGTTTTTAAGCAATCCA

AAAATTTGTATATAATGGTCCGGCTTGTCCCCCAAGTTTACTACCATCCTGAGGAAGTGGTCGTCGATGAGTATGACTAC

CTGATCCAAAAGGGAGCCTGTGACCCTTACCCTAGCAGTCATGCTTGGGTTAGGGATAGCCACCGGTGTAGGGACAGGGA

CCACAGCCCTGATCACAGGACCACAACAATTAGAAAAAGGACTTGGTGAATTACATGCTGTCATAACTGAAGATCTCCAA

GCCTTACAGAAATCCATTAGCAATCTCGAAGAAACCCTGACCTCCTTGTCTGAAGTGGTCTTACAAAACTGGAGGGAATT

AGACCTACTGTTTCTAAAAGAAGGCGGGTTGTGTGCAGCCTTAAAGGAAGAATGTTGCTTTTATGTAGATCACTCAGGAG

CTATCAGAGACTCCATGAGCAAGCTCAGAGAAAGGCTAGAGAAATGTCAAAGGGAGAGGGAAGCTGACCAAGGATGGTTT

GAGTGATGGTTCAACAAGTCCCCCTAGATGACCACTCTGCTTTCTACTCTAATGGGACCCTTAATGGTTTTGCTCCTGAT

GCTTACAGTTGGGCCTTGCATAATTAATAAGTTTATTGCTTTTGTTAGAGAACAAGTGAGTGCAGTCCAGAACATGGTAC

**Env end |**

ACAGACAACAGTACAAGGGCCTTCCAAGCCAAGAGGAAACTGGCATTTAGTCCTCCCAGTTCTAGGATTAGAGCTATTAA

**|3′ LTR start**

CGAGAGAAAAAGTGGGACA**TG**TAAGGCCCAAGAAGTCAGGAGCCTAGAAATACTATCACGCTCCAAAAGGAGCTTTAGCA

GGGCCCTGCATACCTCAAATTCCAGGCCCTACTCTCTGTTAGAAAGTAAGTAAAACGTCACTCTTCTTATTGTGTCAGAG

CCCGCTCTCGAAATAAAACTAGGTAAAAGGGTGTAAAACAGGCCCTTGAGGATGGGAAGTGAGTAACGGATTAAACCGCT

TATGTGGCTTCTGTAAATTGCTCGCACCATAAGCCCTAATAGCCTGTCTTAACCACACTTTGTCACCCCCACCCGAGAGC

CCGCGCAAATGCTGACTGCCAGGGTTATAGGAAACTGATTGGTCCATGTAGCGTGGGCTTGAGTAATTTAAAATGATTGG

CTTAAAAACTATGGAGTGGCATAAACTGACTGGCCCGCACTGCTCAGGCTTCTGATACTAAGAAATGATTGATTTTAATG

CACAGGCTTTGTTAGAAACCATATAAAAGCTGTCCCATTCCTGCATTCGGGGCTCTGCAGTCCTCTACCCCTGCATGGTG

TACAGCTGTGGGCCCCAGCGCGCTTGGAATAAAAATCCTCTTGCTGTTTGCATCAAGACCATTTCTCATGAGTGATTTGG

**3′ LTR end |TSD**

GGTGTCACCATTTCCAGGCAGAGTGTGGGGTCCTCATTCTGGAGGTCTTA**CA**GTAC

**Fig. S2 Detailed descriptions of eJJRV genomes.** One full-length eJJRV annotated in detail. Protein open reading frame (ORF) locations were determined by ORFfinder (https://www.ncbi.nlm.nih.gov/orffinder/) and BLASTn. LTR, *gag*, *pol* and *env* sequences are highlighted. The start and the end of each LTR and ORF are labeled. The end of *pol* and *env* is uncertain.

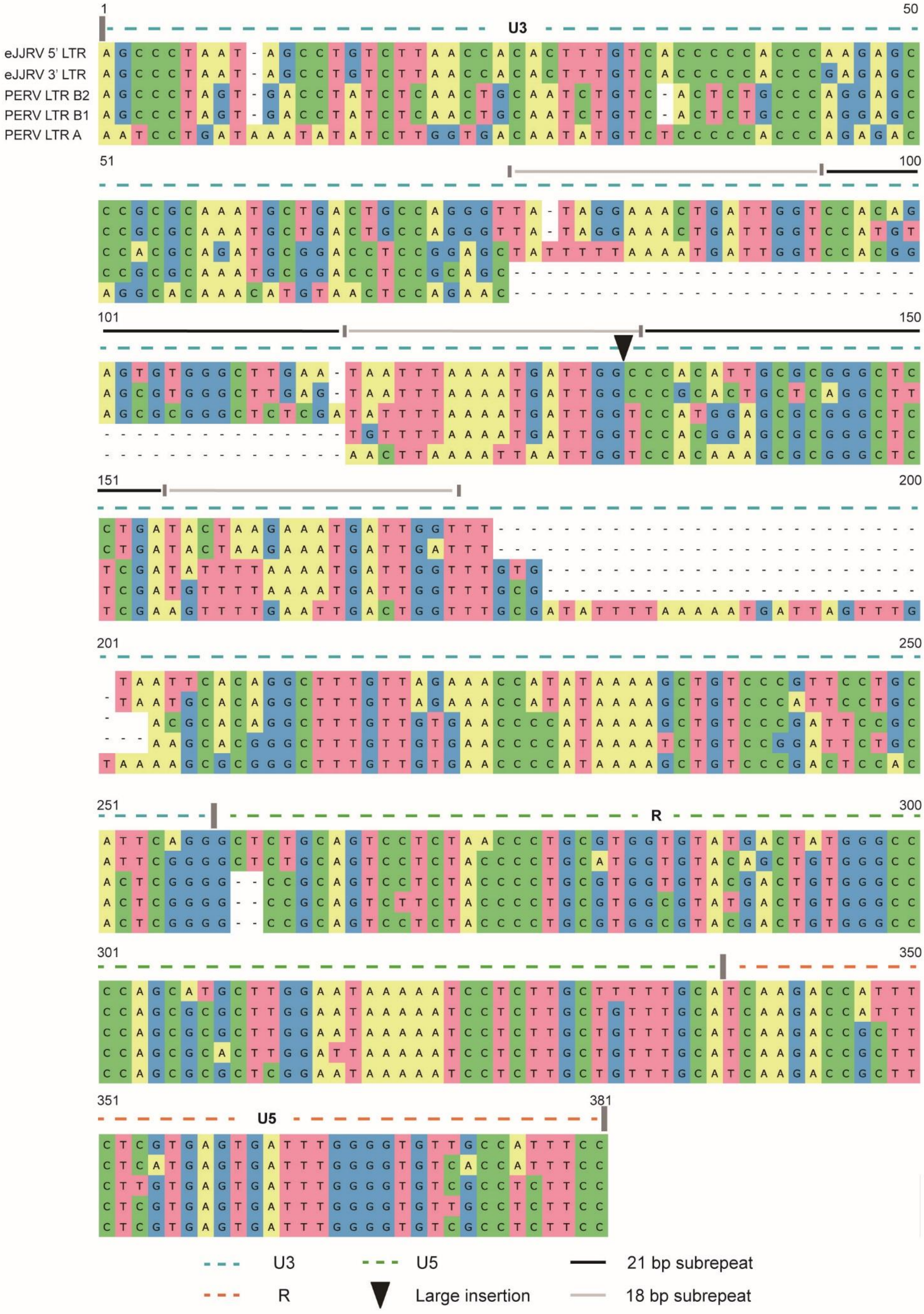


**Fig. S3 The alignment of LTRs of PERVs and eJJRV.** The start of U3 region and the end of U5 region in PERVs and eJJRV are distinct, and these regions are not showed in the alignment. Three types of LTR (LTR A, B1, and B2) of PERVs and one pairwise LTRs of full-length eJJRV (accession number: NW_004504334.1) are selected. Part of U3 region including repeat box, R region and part of U5 region are highly similar between eJJRV and PERVs.

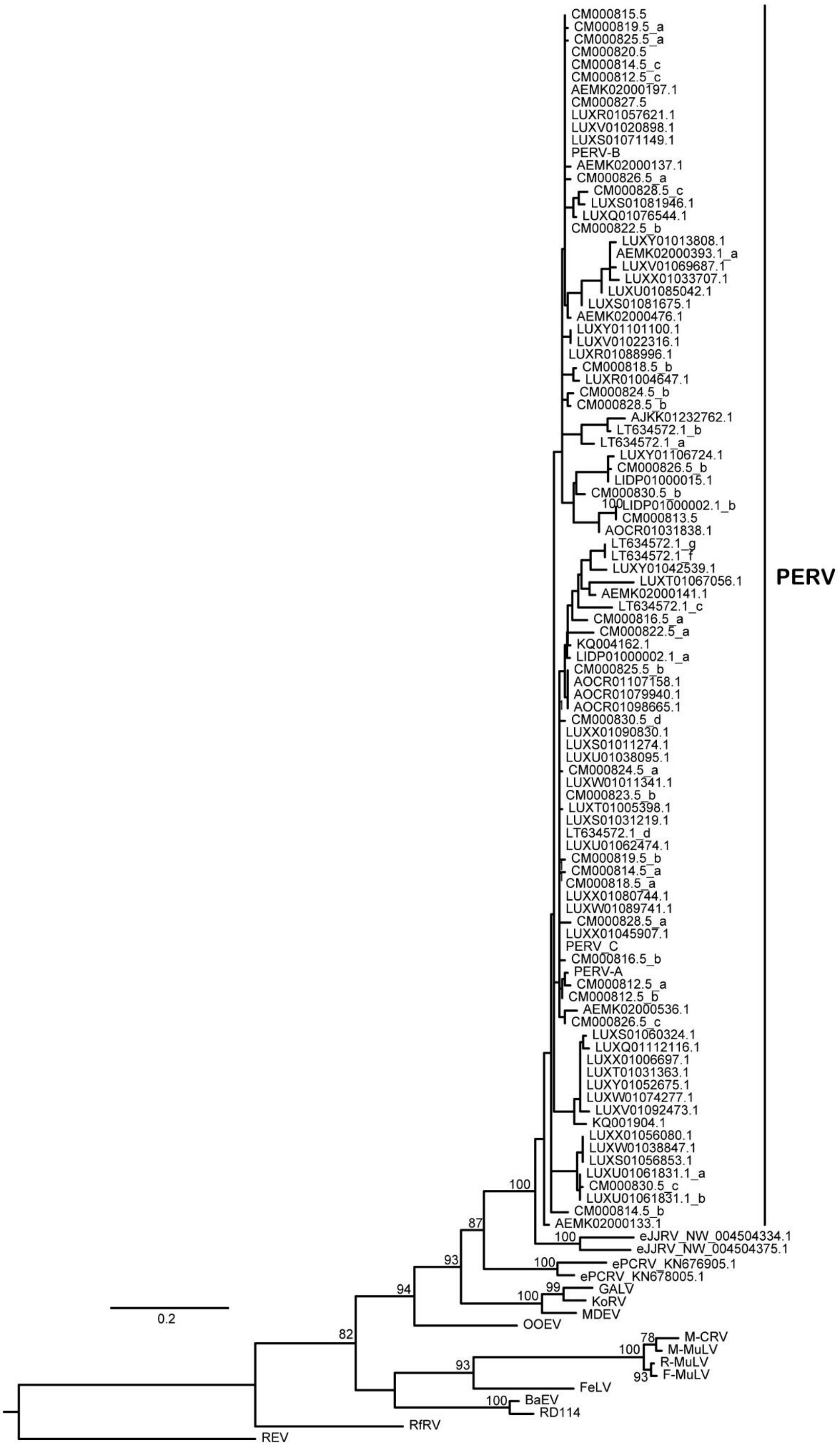
PERV
CM000815.5
CM000819.5_a
CM000825.5_a
CM000820.5
CM000814.5_c
CM000812.5_c
AEMK02000197.1
CM000827.5
LUXR01057621.1
LUXV01020898.1
LUXS01071149.1
PERV-B
AEMK02000137.1
CM000826.5_a
CM000828.5_c
LUXS01081946.1
LUXQ01076544.1
CM000822.5_b
LUXY01013808.1
AEMK02000393.1_a
LUXV01069687.1
LUXX01033707.1
LUXU01085042.1
LUXS01081675.1
AEMK02000476.1
LUXY01101100.1
LUXV01022316.1
LUXR01088996.1
CM000818.5_b
LUXR01004647.1
CM000824.5_b
CM000828.5_b
AJKK01232762.1
LT634572.1_b
LT634572.1_a
LUXY01106724.1
CM000826.5_b
LIDP01000015.1
CM000830.5_b
100
LIDP01000002.1_b
CM000813.5
AOCR01031838.1
LT634572.1_g
LT634572.1_f
LUXY01042539.1
LUXT01067056.1
AEMK02000141.1
LT634572.1_c
CM000816.5_a
CM000822.5_a
KQ004162.1
LIDP01000002.1_a
CM000825.5_b
AOCR01107158.1
AOCR01079940.1
AOCR01098665.1
CM000830.5_d
LUXX01090830.1
LUXS01011274.1
LUXU01038095.1
CM000824.5_a
LUXW01011341.1
CM000823.5_b
LUXT01005398.1
LUXS01031219.1
LT634572.1_d
LUXU01062474.1
CM000819.5_b
CM000814.5_a
CM000818.5_a
LUXX01080744.1
LUXW01089741.1
CM000828.5_a
LUXX01045907.1
PERV_C
CM000816.5_b
PERV-A
CM000812.5_a
CM000812.5_b
AEMK02000536.1
CM000826.5_c
LUXS01060324.1
LUXQ01112116.1
LUXX01006697.1
LUXT01031363.1
LUXY01052675.1
LUXW01074277.1
LUXV01092473.1
KQ001904.1
LUXX01056080.1
LUXW01038847.1
LUXS01056853.1
LUXU01061831.1_a
CM000830.5_c
LUXU01061831.1_b
CM000814.5_b
AEMK02000133.1
100
eJJRV_NW_004504334.1
eJJRV_NW_004504375.1
100
87
ePCRV_KN676905.1
ePCRV_KN678005.1
93
GALV
99
100
KoRV
MDEV
94
OOEV
0.2
82
M-CRV
78
M-MuLV
100
R-MuLV
93
F-MuLV
93
FeLV
100
BaEV
RD114
RfRV
REV

**Fig. S4 The complete phylogenetic tree of Gag.** The phylogenetic tree constructed using the amino acid sequences of Gag of PERVs, eJJRVs and other representative gammaretroviruses. Bootstrap values lower than 70% are not shown in the phylogenetic tree. The alignment used to build the phylogenetic tree is represented in Dataset S1.

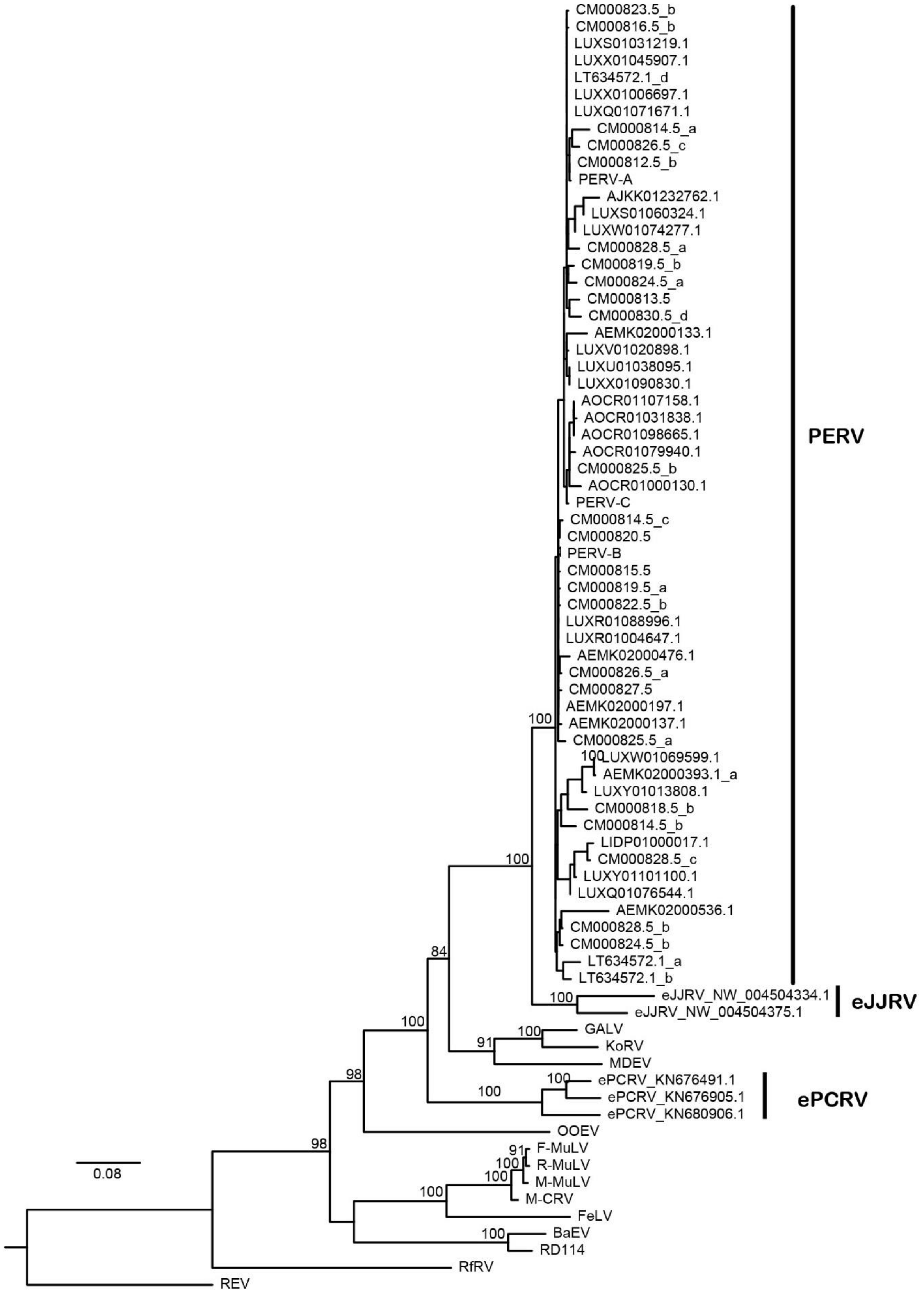


**Fig. S5 The complete phylogenetic tree of Pol.** The phylogenetic tree constructed using the amino acid sequences of Pol of PERVs, eJJRVs and other representative gammaretroviruses. Bootstrap values lower than 70% are not shown in the phylogenetic tree. The alignment used to build the phylogenetic tree is represented in Dataset S1.

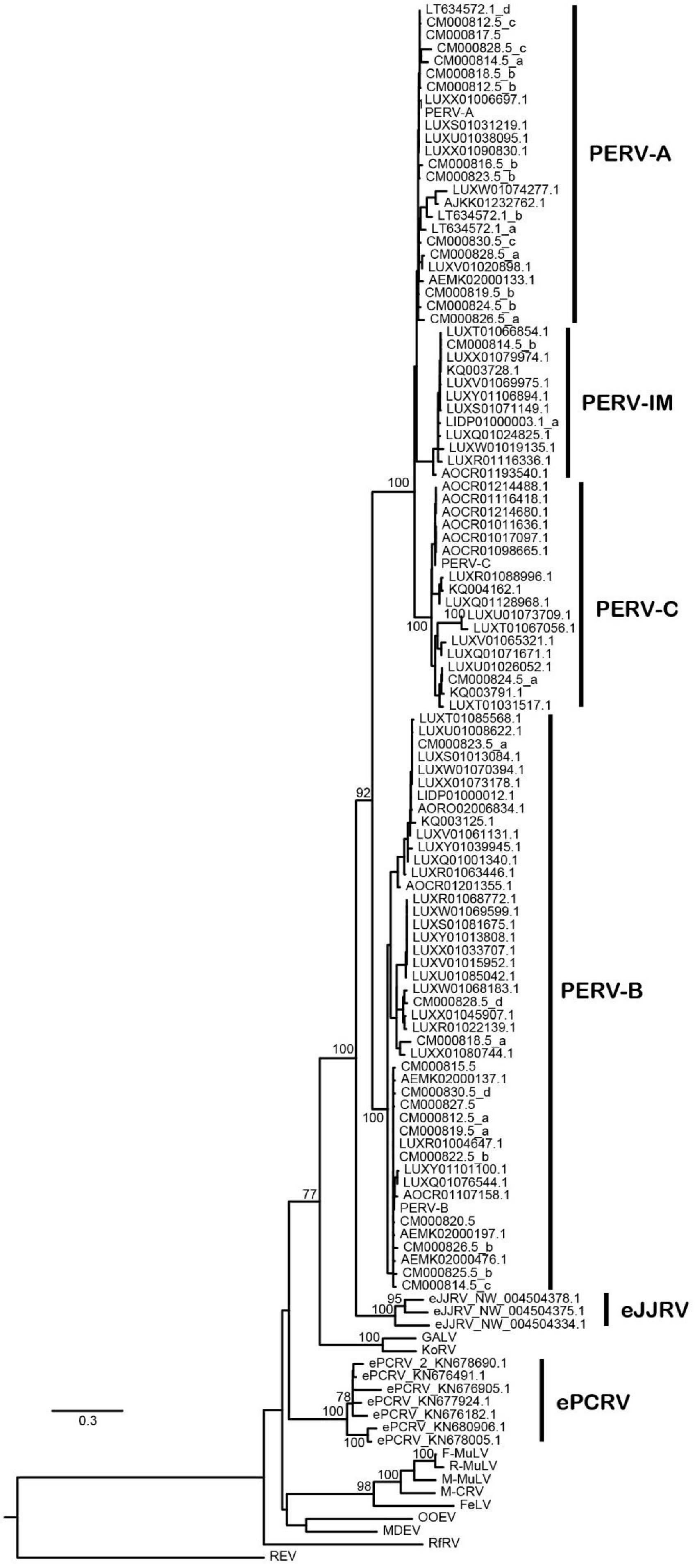
LT634572.1_d
CM000812.5_c
CM000817.5
CM000828.5_c
CM000814.5_a
CM000818.5_b
CM000812.5_b
LUXX01006697.1
PERV-A
LUXS01031219.1
LUXU01038095.1
LUXX01090830.1
CM000816.5_b
CM000823.5_b
LUXW01074277.1
AJKK01232762.1
LT634572.1_b
LT634572.1_a
CM000830.5_c
CM000828.5_a
LUXV01020898.1
AEMK02000133.1
CM000819.5_b
CM000824.5_b
CM000826.5_a
PERV-A
LUXT01066854.1
CM000814.5_b
LUXX01079974.1
KQ003728.1
LUXV01069975.1
LUXY01106894.1
LUXS01071149.1
LIDP01000003.1_a
LUXQ01024825.1
LUXW01019135.1
LUXR01116336.1
AOCR01193540.1
PERV-IM
100
AOCR01214488.1
AOCR01116418.1
AOCR01214680.1
AOCR01011636.1
AOCR01017097.1
AOCR01098665.1
PERV-C
LUXR01088996.1
KQ004162.1
LUXQ01128968.1
100
LUXU01073709.1
100
LUXT01067056.1
LUXV01065321.1
LUXQ01071671.1
LUXU01026052.1
CM000824.5_a
KQ003791.1
LUXT01031517.1
PERV-C
LUXT01085568.1
LUXU01008622.1
CM000823.5_a
LUXS01013084.1
LUXW01070394.1
LUXX01073178.1
LIDP01000012.1
AORO02006834.1
KQ003125.1
LUXV01061131.1
LUXY01039945.1
LUXQ01001340.1
LUXR01063446.1
AOCR01201355.1
92
LUXR01068772.1
LUXW01069599.1
LUXS01081675.1
LUXY01013808.1
LUXX01033707.1
LUXV01015952.1
LUXU01085042.1
LUXW01068183.1
CM000828.5_d
LUXX01045907.1
LUXR01022139.1
CM000818.5_a
LUXX01080744.1
PERV-B
100
CM000815.5
AEMK02000137.1
CM000830.5_d
CM000827.5
CM000812.5_a
CM000819.5_a
LUXR01004647.1
CM000822.5_b
LUXY01101100.1
LUXQ01076544.1
AOCR01107158.1
PERV-B
CM000820.5
AEMK02000197.1
CM000826.5_b
AEMK02000476.1
CM000825.5_b
CM000814.5_c
100
77
95
eJJRV_NW_004504378.1
100
eJJRV_NW_004504375.1
eJJRV_NW_004504334.1
eJJRV
100
GALV
KoRV
ePCRV_2_KN678690.1
ePCRV_KN676491.1
ePCRV_KN676905.1
78
ePCRV_KN677924.1
100
ePCRV_KN676182.1
ePCRV_KN680906.1
100
ePCRV_KN678005.1
ePCRV
0.3
100
F-MuLV
R-MuLV
100
M-MuLV
98
M-CRV
FeLV
OOEV
MDEV
RfRV
REV

**Fig. S6 The complete phylogenetic tree of Env.** The phylogenetic tree constructed using the amino acid sequences of Env of PERVs, eJJRVs and other representative gammaretroviruses. Bootstrap values lower than 70% are not shown in the phylogenetic tree. The alignment used to build the phylogenetic tree is represented in Dataset S1.

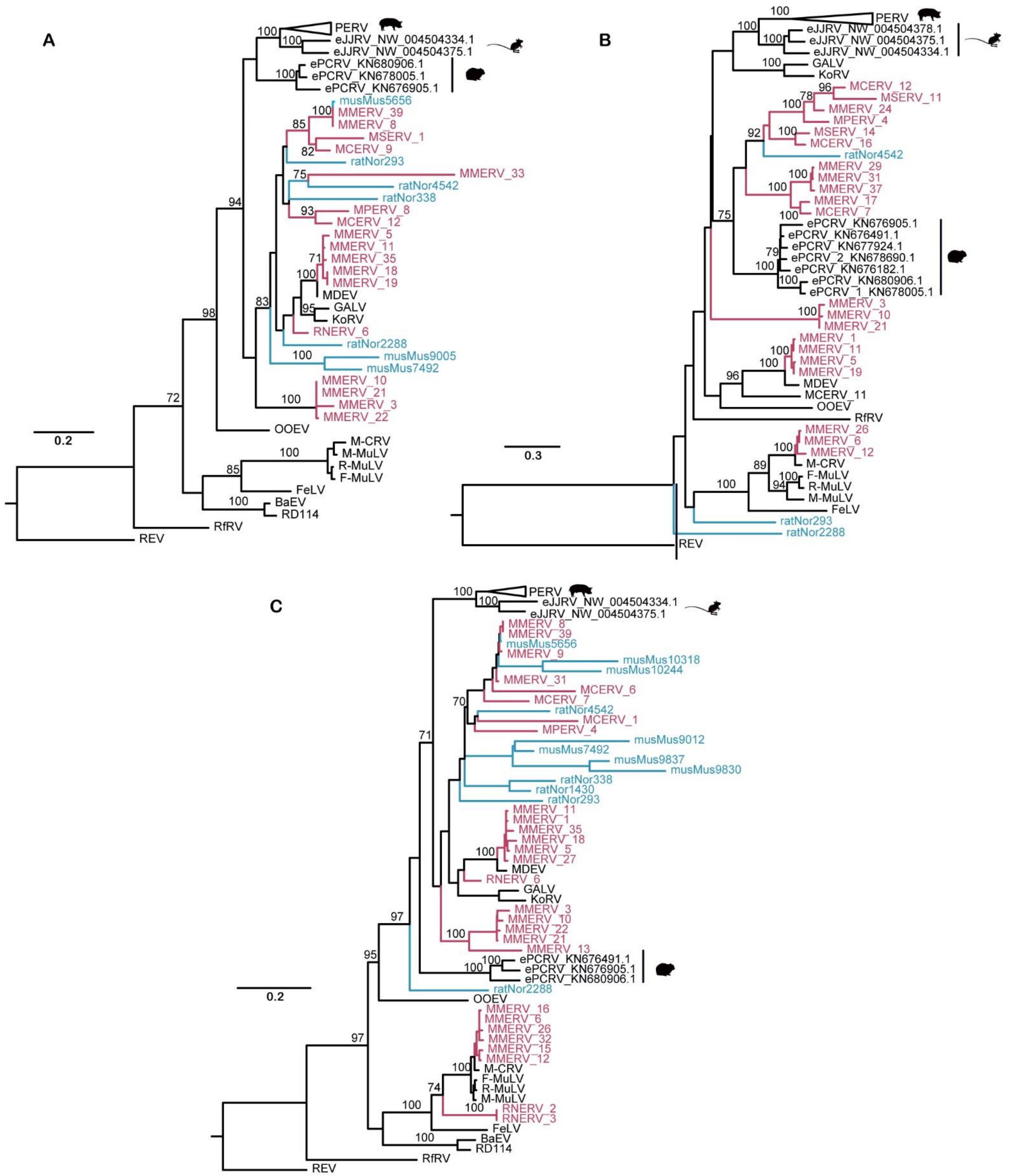


**Fig. S7. The complete phylogenetic tree of Gag, Pol and Env of *Muroidea* ERVs, eJJRVs, ePCRVs, PERVs and representative retroviruses**. Phylogenetic trees of Gag (A), Pol (B) and Env (C) constructed using amino acid (aa) sequences of PERVs, eJJRV,

*Muroidea* ERVs (Table S6) and other representative gammaretroviruses (Table S7). The lengths of amino acid are 225aa (Gag), 567aa (Pol) and 334aa (Env), respectively. he best-fit JTT+Γ amino acid substitution model was selected for Gag, Pol and JTT+Γ+I for Env. Bootstrap values <70% are not presented in the phylogenetic trees. Trees were rooted using Reticuloendotheliosis virus (REV). The information of *Muroidea* genomes presents in Table S1. Previous found ERVs are labeled in blue. *Muroidea* ERVs are labeled in red. The alignment used to build the phylogenetic tree is represented in Dataset S2.
