## Supplementary material for "Genomic insights into evolutionary journey of the porcine endogenous retroviruses"

**Table S1.** The information of pig, rodent, and rock hyrax genomes used for data mining.

| Species | Breed | Geographic location | GenBank assembly accession | Copy number of ERVs |
|---|---|---|---|---|
| *Sus scrofa* (pig) | Duroc (TJ Tabasco) | Europe | GCA_000003025.6 | 47 |
| | Wuzhishan | China | GCA_000325925.2 | 10 |
| | Ellegaard Gottingen minipig | Germany, Europe | GCA_000331475.1 | 5 |
| | Tibetan | China | GCA_000472085.2 | 1 |
| | Goettingen | Denmark, Europe | GCA_001292865.1 | 7 |
| | Meishan | China | GCA_001700195.1 | 3 |
| | Rongchang | China | GCA_001700155.1 | 7 |
| | Hampshire | Europe | GCA_001700165.1 | 6 |
| | Landrace | Denmark, Europe | GCA_001700215.1 | 2 |
| | Pietrain | Europe | GCA_001700255.1 | 8 |
| | Bamei | China | GCA_001700235.1 | 2 |
| | Berkshire | Europe | GCA_001700575.1 | 4 |
| | LargeWhite | Europe | GCA_001700135.1 | 9 |
| | Jinhua | China | GCA_001700295.1 | 5 |
| *Jaculus jaculus* (lesser Egyptian jerboa) | - | Northern Africa, the Middle East and Central Asia | GCA_000280705.1 | 4 |
| *Procavia capensis* (Cape rock hyrax) | - | Africa and Central Asia | GCA_000152225.2 | 8 |
| *Mus caroli* (Ryukyu mouse) | - | - | GCF_900094665.1 | 20 |
| *Mus pahari* (shrew mouse) | - | - | GCF_900095145.1 | 15 |
| *Mus musculus* (house mouse) | - | - | GCF_000001635.26 | 39 |
| *Mus spretus* (western wild mouse) | - | - | GCA_001624865.1 | 14 |

| | | | | |
|---|---|---|---|---|
| *Apodemus speciosus* (large Japanese field mouse) | - | - | GCA_002335545.1 | 4 |
| *Apodemus sylvaticus* (European woodmouse) | - | - | GCA_001305905.1 | 0 |
| *Rattus norvegicus* (Norway rat) | - | - | GCF_000001895.5 | 6 |
| *Phodopus sungorus* | - | - | GCA_001707965.1 | 0 |

- The corresponding information is unknown

**Table S2. The information of full-length and near full-length PERVs.**

| Breed | Name | Class of PERV§ | Locus | Orientation (+/-) | Start | End |
|---|---|---|---|---|---|---|
| Duroc | AEMK02000133.1 * | A | AEMK02000133.1 | - | 563793 | 572725 |
| Duroc | AEMK02000137.1 * | B | AEMK02000137.1 | + | 1949 | 10743 |
| Duroc | AEMK02000141.1 * | UN | AEMK02000141.1 | + | 19304 | 23938 |
| Duroc | AEMK02000197.1 * | B | AEMK02000197.1 | + | 13836 | 22597 |
| Duroc | AEMK02000319.1 | UN | AEMK02000319.1 | + | 3229 | 6461 |
| Duroc | AEMK02000393.1_a * | B | AEMK02000393.1 | + | 820148 | 829191 |
| Duroc | AEMK02000393.1_b | UN | AEMK02000393.1 | + | 909902 | 914028 |
| Duroc | AEMK02000476.1 * | B | AEMK02000476.1 | - | 42158 | 50876 |
| Duroc | AEMK02000536.1 * | A | AEMK02000536.1 | + | 14633 | 22827 |
| Duroc | AEMK02000643.1 | UN | AEMK02000643.1 | + | 4001 | 7228 |
| Duroc | CM000812.5_a * | B | CM000812.5 | + | 38459013 | 38465006 |
| Duroc | CM000812.5_b * | A | CM000812.5 | + | 262166347 | 262175261 |
| Duroc | CM000812.5_c * | A | CM000812.5 | + | 132020281 | 132028360 |
| Duroc | CM000813.5 * | UN | CM000813.5 | - | 76648002 | 76655788 |
| Duroc | CM000814.5_a * | A | CM000814.5 | - | 10660619 | 10669533 |
| Duroc | CM000814.5_b * | IM | CM000814.5 | + | 17778858 | 17788086 |
| Duroc | CM000814.5_c * | B | CM000814.5 | - | 51108601 | 51117360 |
| Duroc | CM000815.5 * | B | CM000815.5 | - | 45599494 | 45608252 |
| Duroc | CM000816.5_a | UN | CM000816.5 | + | 29356398 | 29361616 |
| Duroc | CM000816.5_b * | A | CM000816.5 | - | 92185133 | 92194050 |
| Duroc | CM000817.5 * | A | CM000817.5 | + | 10402614 | 10407657 |
| Duroc | CM000818.5_a * | B | CM000818.5 | + | 21236160 | 21244976 |
| Duroc | CM000818.5_b * | A | CM000818.5 | + | 105710884 | 105719193 |
| Duroc | CM000819.5_a * | B | CM000819.5 | + | 15319428 | 15328187 |
| Duroc | CM000819.5_b * | A | CM000819.5 | + | 51570546 | 51579460 |
| Duroc | CM000820.5 * | B | CM000820.5 | - | 138895584 | 138904340 |

| | | | | | | |
|---|---|---|---|---|---|---|
| Duroc | CM000822.5_a * | UN | CM000822.5 | - | 29084884 | 29090328 |
| Duroc | CM000822.5_b * | B | CM000822.5 | + | 38201361 | 38210116 |
| Duroc | CM000823.5_a * | B | CM000823.5 | - | 22986890 | 22991220 |
| Duroc | CM000823.5_b * | A | CM000823.5 | + | 28221374 | 28230286 |
| Duroc | CM000824.5_a * | C | CM000824.5 | + | 21308535 | 21310710 |
| Duroc | CM000824.5_b * | A | CM000824.5 | - | 142030847 | 142039759 |
| Duroc | CM000824.5_d * | A | CM000824.5 | - | 146750531 | 146760098 |
| Duroc | CM000825.5_a * | C | CM000825.5 | - | 62550397 | 62557645 |
| Duroc | CM000825.5_b * | B | CM000825.5 | + | 119667621 | 119676299 |
| Duroc | CM000826.5_a * | A | CM000826.5 | - | 66864153 | 66873044 |
| Duroc | CM000826.5_b * | B | CM000826.5 | - | 110958945 | 110967782 |
| Duroc | CM000826.5_c * | IM | CM000826.5 | + | 116321668 | 116334781 |
| Duroc | CM000827.5 * | B | CM000827.5 | + | 59571885 | 59580646 |
| Duroc | CM000828.5_a * | A | CM000828.5 | - | 3794010 | 3802775 |
| Duroc | CM000828.5_b * | IM | CM000828.5 | - | 9536784 | 9542014 |
| Duroc | CM000828.5_c * | A | CM000828.5 | + | 33062883 | 33071800 |
| Duroc | CM000828.5_d * | B | CM000828.5 | - | 41467516 | 41476101 |
| Duroc | CM000830.5_a * | UN | CM000830.5 | + | 71402222 | 71409911 |
| Duroc | CM000830.5_b * | UN | CM000830.5 | + | 72244018 | 72249170 |
| Duroc | CM000830.5_c * | A | CM000830.5 | - | 73752986 | 73761903 |
| Duroc | CM000830.5_d * | B | CM000830.5 | - | 112842765 | 112847657 |
| Duroc | LT634572.1_a * | A | LT634572.1 | - | 20194782 | 20203572 |
| Duroc | LT634572.1_b * | A | LT634572.1 | - | 21867301 | 21876037 |
| Duroc | LT634572.1_c | UN | LT634572.1 | + | 24407524 | 24411155 |
| Duroc | LT634572.1_d * | A | LT634572.1 | + | 25245606 | 25253936 |
| Duroc | LT634572.1_f * | UN | LT634572.1 | - | 25269797 | 25273827 |
| Duroc | LT634572.1_g | UN | LT634572.1 | - | 25299029 | 25302045 |
| Wuzhishan | AJKK01232762.1 * | A | AJKK01232762.1 | + | 23882 | 32746 |
| Wuzhishan | KQ001810.1 * | UN | KQ001810.1 | + | 268790 | 282360 |
| Wuzhishan | KQ001904.1 | UN | KQ001904.1 | - | 360801 | 363847 |

| | | | | | | |
|---|---|---|---|---|---|---|
| Wuzhishan | KQ001967.1 * | B | KQ001967.1 | + | 701263 | 709367 |
| Wuzhishan | KQ001997.1 * | IM | KQ001997.1 | - | 317985 | 322992 |
| Wuzhishan | KQ003125.1 * | B | KQ003125.1 | + | 442452 | 446737 |
| Wuzhishan | KQ003449.1 * | UN | KQ003449.1 | + | 2431965 | 2439733 |
| Wuzhishan | KQ003477.1 * | IM | KQ003477.1 | - | 4611805 | 4618707 |
| Wuzhishan | KQ003728.1 * | IM | KQ003728.1 | + | 602469 | 610981 |
| Wuzhishan | KQ003791.1 * | C | KQ003791.1 | - | 297736 | 300067 |
| Wuzhishan | KQ004162.1 * | C | KQ004162.1 | + | 1378452 | 1386833 |
| Ellegaard | AOCR01000130.1 | C | AOCR01000130.1 | + | 5 | 5410 |
| Ellegaard | AOCR01011636.1 | C | AOCR01011636.1 | + | 2 | 3124 |
| Ellegaard | AOCR01017097.1 * | C | AOCR01017097.1 | + | 925 | 9470 |
| Ellegaard | AOCR01031838.1 | UN | AOCR01031838.1 | - | 3235 | 9338 |
| Ellegaard | AOCR01035274.1 | C | AOCR01035274.1 | + | 18917 | 20586 |
| Ellegaard | AOCR01079940.1 * | C | AOCR01079940.1 | + | 47557 | 55276 |
| Ellegaard | AOCR01098665.1 * | C | AOCR01098665.1 | - | 83072 | 91753 |
| Ellegaard | AOCR01107158.1 | B | AOCR01107158.1 | - | 3 | 7772 |
| Ellegaard | AOCR01116418.1 * | C | AOCR01116418.1 | - | 30836 | 39272 |
| Ellegaard | AOCR01122800.1 | B | AOCR01122800.1 | - | 9560 | 12138 |
| Ellegaard | AOCR01193540.1 | IM | AOCR01193540.1 | - | 3726 | 6667 |
| Ellegaard | AOCR01201355.1 | B | AOCR01201355.1 | + | 1 | 2403 |
| Ellegaard | AOCR01214488.1 | C | AOCR01214488.1 | - | 2687 | 5809 |
| Ellegaard | AOCR01214680.1 * | C | AOCR01214680.1 | + | 9997 | 18501 |
| Tibetan | AORO02006834.1 * | B | AORO02006834.1 | + | 28813 | 33341 |
| Tibetan | AORO02050002.1 | IM | AORO02050002.1 | + | 1 | 2752 |
| Goettingen | LIDP01000002.1_a | UN | LIDP01000002.1 | - | 743130 | 746983 |
| Goettingen | LIDP01000002.1_b * | C | LIDP01000002.1 | + | 77151802 | 77159575 |
| Goettingen | LIDP01000003.1_a | IM | LIDP01000003.1 | + | 18145309 | 18150139 |
| Goettingen | LIDP01000003.1_b | UN | LIDP01000003.1 | + | 18232716 | 18237157 |
| Goettingen | LIDP01000005.1 | UN | LIDP01000005.1 | + | 32344699 | 32349877 |
| Goettingen | LIDP01000008.1 | UN | LIDP01000008.1 | + | 54505184 | 54507036 |

| | | | | | | |
|---|---|---|---|---|---|---|
| Goettingen | LIDP01000012.1 * | B | LIDP01000012.1 | - | 23435735 | 23440056 |
| Goettingen | LIDP01000013.1_a * | A | LIDP01000013.1 | + | 151258154 | 151267071 |
| Goettingen | LIDP01000013.1_b * | C | LIDP01000013.1 | - | 156298083 | 156311680 |
| Goettingen | LIDP01000015.1 * | IM | LIDP01000015.1 | - | 128579984 | 128592582 |
| Goettingen | LIDP01000017.1 * | B | LIDP01000017.1 | - | 46901312 | 46909894 |
| Goettingen | LIDP01000019.1 * | UN | LIDP01000019.1 | + | 83266097 | 83269702 |
| Meishan | LUXQ01001340.1 * | B | LUXQ01001340.1 | + | 91288 | 95595 |
| Meishan | LUXQ01004839.1 | UN | LUXQ01004839.1 | + | 625458 | 627570 |
| Meishan | LUXQ01024825.1 | IM | LUXQ01024825.1 | + | 3 | 4124 |
| Meishan | LUXQ01042183.1 * | IM | LUXQ01042183.1 | + | 424777 | 429352 |
| Meishan | LUXQ01071671.1 | C | LUXQ01071671.1 | + | 356347 | 362858 |
| Meishan | LUXQ01076544.1 * | B | LUXQ01076544.1 | + | 3044660 | 3052938 |
| Meishan | LUXQ01091036.1 | UN | LUXQ01091036.1 | + | 11388 | 20566 |
| Meishan | LUXQ01112116.1 | UN | LUXQ01112116.1 | - | 1 | 3678 |
| Meishan | LUXQ01128968.1 | C | LUXQ01128968.1 | + | 92 | 7467 |
| Rongchang | LUXR01004647.1 * | B | LUXR01004647.1 | - | 55158 | 64105 |
| Rongchang | LUXR01007605.1 | UN | LUXR01007605.1 | + | 7009 | 17141 |
| Rongchang | LUXR01012890.1 * | IM | LUXR01012890.1 | - | 361433 | 366793 |
| Rongchang | LUXR01015861.1 | UN | LUXR01015861.1 | + | 19049 | 21918 |
| Rongchang | LUXR01022139.1 * | B | LUXR01022139.1 | + | 3865481 | 3874153 |
| Rongchang | LUXR01030059.1 | UN | LUXR01030059.1 | + | 53893 | 56745 |
| Rongchang | LUXR01037956.1 * | A | LUXR01037956.1 | + | 37902 | 46313 |
| Rongchang | LUXR01057621.1 * | UN | LUXR01057621.1 | + | 106155 | 112848 |
| Rongchang | LUXR01063446.1 * | B | LUXR01063446.1 | - | 1595639 | 1599939 |
| Rongchang | LUXR01068772.1 | B | LUXR01068772.1 | - | 10 | 2439 |
| Rongchang | LUXR01088996.1 * | C | LUXR01088996.1 | - | 1998263 | 2007096 |
| Rongchang | LUXR01116336.1 | IM | LUXR01116336.1 | + | 1 | 4618 |
| Hampshire | LUXS01011274.1 * | UN | LUXS01011274.1 | + | 400996 | 406126 |
| Hampshire | LUXS01013084.1 * | B | LUXS01013084.1 | - | 26330 | 30654 |
| Hampshire | LUXS01031219.1 * | A | LUXS01031219.1 | - | 220229 | 229152 |

| | | | | | | |
|---|---|---|---|---|---|---|
| Hampshire | LUXS01037717.1 | B | LUXS01037717.1 | - | 1178453 | 1180422 |
| Hampshire | LUXS01045203.1 | IM | LUXS01045203.1 | + | 35 | 3505 |
| Hampshire | LUXS01050965.1 * | IM | LUXS01050965.1 | + | 2502235 | 2507449 |
| Hampshire | LUXS01056853.1 | UN | LUXS01056853.1 | - | 212954 | 216275 |
| Hampshire | LUXS01060324.1 * | A | LUXS01060324.1 | + | 26936 | 35791 |
| Hampshire | LUXS01071149.1 * | IM | LUXS01071149.1 | - | 1863478 | 1872541 |
| Hampshire | LUXS01081675.1 | B | LUXS01081675.1 | - | 353 | 8766 |
| Hampshire | LUXS01081946.1 | UN | LUXS01081946.1 | - | 3 | 2916 |
| Landrace | LUXT01005398.1 | UN | LUXT01005398.1 | + | 866715 | 870298 |
| Landrace | LUXT01022621.1 | UN | LUXT01022621.1 | + | 256840 | 259949 |
| Landrace | LUXT01031363.1 | UN | LUXT01031363.1 | + | 29086 | 32079 |
| Landrace | LUXT01031517.1 | C | LUXT01031517.1 | + | 1 | 2094 |
| Landrace | LUXT01050601.1 | UN | LUXT01050601.1 | - | 5944 | 16044 |
| Landrace | LUXT01066854.1 | IM | LUXT01066854.1 | - | 1517491 | 1520442 |
| Landrace | LUXT01067056.1 * | C | LUXT01067056.1 | - | 872799 | 881829 |
| Landrace | LUXT01085568.1 * | B | LUXT01085568.1 | - | 1107449 | 1111798 |
| Landrace | LUXT01085818.1 | IM | LUXT01085818.1 | + | 1 | 1871 |
| Pietrain | LUXU01005948.1 | IM | LUXU01005948.1 | + | 3 | 3038 |
| Pietrain | LUXU01008622.1 * | B | LUXU01008622.1 | - | 820208 | 824532 |
| Pietrain | LUXU01015352.1 | UN | LUXU01015352.1 | - | 1761 | 8413 |
| Pietrain | LUXU01026052.1 * | C | LUXU01026052.1 | + | 64738 | 66913 |
| Pietrain | LUXU01029113.1 | UN | LUXU01029113.1 | + | 2547 | 12691 |
| Pietrain | LUXU01036526.1 * | IM | LUXU01036526.1 | + | 2502653 | 2507784 |
| Pietrain | LUXU01038095.1 * | A | LUXU01038095.1 | - | 221923 | 230985 |
| Pietrain | LUXU01061831.1 * | UN | LUXU01061831.1 | + | 256260 | 263999 |
| Pietrain | LUXU01062474.1 * | UN | LUXU01062474.1 | - | 1642044 | 1650290 |
| Pietrain | LUXU01065281.1 * | UN | LUXU01065281.1 | - | 835413 | 839986 |
| Pietrain | LUXU01073709.1 * | C | LUXU01073709.1 | - | 873408 | 881687 |
| Pietrain | LUXU01085042.1 | B | LUXU01085042.1 | - | 356 | 8873 |
| Bamei | LUXV01015952.1 | B | LUXV01015952.1 | + | 52 | 2622 |

| | | | | | | |
|---|---|---|---|---|---|---|
| Bamei | LUXV01020898.1 * | A | LUXV01020898.1 | - | 220830 | 229741 |
| Bamei | LUXV01022316.1 | UN | LUXV01022316.1 | + | 3046222 | 3049009 |
| Bamei | LUXV01061131.1 * | B | LUXV01061131.1 | + | 2274019 | 2278341 |
| Bamei | LUXV01065321.1 | C | LUXV01065321.1 | + | 2 | 3300 |
| Bamei | LUXV01069687.1 | UN | LUXV01069687.1 | - | 1 | 3929 |
| Bamei | LUXV01069975.1 | IM | LUXV01069975.1 | - | 2398090 | 2402284 |
| Bamei | LUXV01092473.1 | UN | LUXV01092473.1 | + | 27905 | 30890 |
| Bamei | LUXV01098735.1 | IM | LUXV01098735.1 | - | 44874 | 46744 |
| Berkshire | LUXW01011341.1 | UN | LUXW01011341.1 | - | 1 | 3644 |
| Berkshire | LUXW01019135.1 | IM | LUXW01019135.1 | + | 9 | 3427 |
| Berkshire | LUXW01038847.1 | UN | LUXW01038847.1 | - | 277987 | 281309 |
| Berkshire | LUXW01040954.1 * | IM | LUXW01040954.1 | + | 2475476 | 2480328 |
| Berkshire | LUXW01049000.1 | UN | LUXW01049000.1 | + | 2 | 2913 |
| Berkshire | LUXW01050896.1 | UN | LUXW01050896.1 | + | 46 | 2237 |
| Berkshire | LUXW01068183.1 | B | LUXW01068183.1 | - | 3943490 | 3946828 |
| Berkshire | LUXW01069599.1 * | B | LUXW01069599.1 | + | 55747 | 64183 |
| Berkshire | LUXW01070394.1 * | B | LUXW01070394.1 | - | 372279 | 376655 |
| Berkshire | LUXW01074277.1 * | A | LUXW01074277.1 | - | 3026 | 11942 |
| Berkshire | LUXW01089741.1 | UN | LUXW01089741.1 | - | 1 | 4941 |
| LargeWhite | LUXX01006697.1 * | A | LUXX01006697.1 | + | 14787 | 23496 |
| LargeWhite | LUXX01027740.1 * | UN | LUXX01027740.1 | - | 2537039 | 2543883 |
| LargeWhite | LUXX01033707.1 * | B | LUXX01033707.1 | + | 223594 | 232510 |
| LargeWhite | LUXX01033772.1 | UN | LUXX01033772.1 | + | 342 | 6101 |
| LargeWhite | LUXX01045907.1 * | B | LUXX01045907.1 | + | 2628683 | 2637041 |
| LargeWhite | LUXX01056080.1 | UN | LUXX01056080.1 | - | 416976 | 420298 |
| LargeWhite | LUXX01071374.1 * | UN | LUXX01071374.1 | + | 2476375 | 2481476 |
| LargeWhite | LUXX01073178.1 * | B | LUXX01073178.1 | + | 451456 | 455777 |
| LargeWhite | LUXX01079974.1 * | IM | LUXX01079974.1 | + | 2109764 | 2119026 |
| LargeWhite | LUXX01080744.1 * | B | LUXX01080744.1 | + | 111478 | 120196 |
| LargeWhite | LUXX01090830.1 * | A | LUXX01090830.1 | - | 33028 | 41951 |

| | | | | | | |
|---|---|---|---|---|---|---|
| Jinhua | LUXY01006678.1 * | A | LUXY01006678.1 | + | 2493985 | 2498941 |
| Jinhua | LUXY01013808.1 | B | LUXY01013808.1 | - | 356 | 8398 |
| Jinhua | LUXY01039945.1 * | B | LUXY01039945.1 | + | 2311187 | 2315339 |
| Jinhua | LUXY01042539.1 * | UN | LUXY01042539.1 | + | 101981 | 105208 |
| Jinhua | LUXY01049233.1 | UN | LUXY01049233.1 | + | 11041 | 21148 |
| Jinhua | LUXY01052675.1 * | UN | LUXY01052675.1 | - | 2657 | 11285 |
| Jinhua | LUXY01101100.1 * | B | LUXY01101100.1 | + | 3039859 | 3048982 |
| Jinhua | LUXY01106724.1 | UN | LUXY01106724.1 | + | 81968 | 84391 |
| Jinhua | LUXY01106894.1 | IM | LUXY01106894.1 | - | 456844 | 461221 |

* The full-length PERVs containing both 5' and 3' LTR. PERVs without * represent those only having one LTR.

[§] UN represents the unclassified PERVs for loss of *env*.

**Table S3.** The recombination-related information of full-length PERVs shown in Fig. S1.

| Name | Accession number | Orientation | Divergent LTRs based on structure* | Divergent LTR based on tree | Flanking TSD** | | Are they the same flanking TSD sequence? |
|---|---|---|---|---|---|---|---|
| | | | | | 5' | 3' | |
| AEMK02000133.1 | AEMK02000133.1 | - | No | Yes | GTTG | GTTG | Yes |
| AEMK02000137.1 | AEMK02000137.1 | + | Yes | No | GAAC | GAAC | Yes |
| AEMK02000141.1 | AEMK02000141.1 | + | No | No | GTAT | GTAT | Yes |
| AEMK02000197.1 | AEMK02000197.1 | + | No | No | GGAG | GGAG | Yes |
| AEMK02000476.1 | AEMK02000476.1 | - | No | No | GAAC | GAAC | Yes |
| AEMK02000536.1 | AEMK02000536.1 | + | Yes | Yes | AGCC | CTTT | No |
| AJKK01232762.1 | AJKK01232762.1 | + | No | Yes | GTCT | GTCT | Yes |
| AOCR01017097.1 | AOCR01017097.1 | + | No | No | CTAA | CTAA | Yes |
| AOCR01098665.1 | AOCR01098665.1 | - | No | Yes | GTTG | GTTG | Yes |
| AOCR01116418.1 | AOCR01116418.1 | - | No | No | CTAG | CTAG | Yes |
| AOCR01214680.1 | AOCR01214680.1 | + | No | Yes | ACAT | ACAT | Yes |
| AORO02006834.1 | AORO02006834.1 | + | No | Yes | CAAG | CAAG | Yes |
| CM000812.5_c | CM000812.5 | + | No | Yes | GACC | GACC | Yes |
| CM000812.5_b | CM000812.5 | + | No | No | CTAC | CTAC | Yes |
| CM000812.5_a | CM000812.5 | + | Yes | No | CCAC | CCAC | Yes |
| CM000813.5 | CM000813.5 | - | No | Yes | GTTC | GTTC | Yes |
| CM000814.5_a | CM000814.5 | - | No | Yes | CAGA | CAGA | Yes |
| CM000814.5_b | CM000814.5 | + | No | Yes | GAAG | GAAG | Yes |
| CM000814.5_c | CM000814.5 | - | No | No | CACC | CACC | Yes |
| CM000815.5 | CM000815.5 | - | No | No | GTTT | GTTT | Yes |
| CM000816.5_b | CM000816.5 | - | No | No | GTTC | GTTC | Yes |
| CM000817.5 | CM000817.5 | + | No | No | GTTC | GTTC | Yes |
| CM000818.5_b | CM000818.5 | + | No | No | GTAG | GTAG | Yes |
| CM000818.5_a | CM000818.5 | + | No | Yes | GTTC | CTTC | No |
| CM000819.5_a | CM000819.5 | + | No | No | CTTT | CTTT | Yes |
| CM000819.5_b | CM000819.5 | + | No | No | AGAG | AGAG | Yes |
| CM000820.5 | CM000820.5 | - | No | No | CCAG | CCAG | Yes |

| | | | | | | | |
|---|---|---|---|---|---|---|---|
| CM000822.5_a | CM000822.5 | - | No | Yes | GTAT | GTAT | Yes |
| CM000822.5_b | CM000822.5 | + | No | No | TGAC | TGAC | Yes |
| CM000823.5_a | CM000823.5 | - | No | Yes | CAAG | CAAG | Yes |
| CM000823.5_b | CM000823.5 | + | No | No | GAAT | GAAT | Yes |
| CM000824.5_b | CM000824.5 | - | No | No | ACAC | ACAC | Yes |
| CM000825.5_b | CM000825.5 | + | No | No | GTAG | GTAG | Yes |
| CM000825.5_a | CM000825.5 | - | No | No | AAAG | AAAG | Yes |
| CM000826.5_b | CM000826.5 | - | No | No | ATCC | ATCC | Yes |
| CM000826.5_c | CM000826.5 | + | No | No | ACCA | AATC | No |
| CM000826.5_a | CM000826.5 | - | No | No | CTTC | CTTC | Yes |
| CM000827.5 | CM000827.5 | + | No | No | CTAT | CTAT | Yes |
| CM000828.5_c | CM000828.5 | + | No | Yes | AATA | AATA | Yes |
| CM000828.5_a | CM000828.5 | - | Yes | No | GGCT | GGCT | Yes |
| CM000828.5_d | CM000828.5 | - | No | Yes | CCAC | CACC | No |
| CM000828.5_b | CM000828.5 | - | No | Yes | ATAC | ATAC | Yes |
| CM000830.5_d | CM000830.5 | - | No | Yes | CAAC | CAAC | Yes |
| CM000830.5_a | CM000830.5 | + | No | No | GTGT | GTGT | Yes |
| CM000830.5_b | CM000830.5 | + | No | No | CCTC | CCTC | Yes |
| CM000830.5_c | CM000830.5 | - | No | Yes | CTTT | CTTT | Yes |
| KQ001967.1 | KQ001967.1 | + | No | Yes | CCAC | CACC | No |
| KQ003449.1 | KQ003449.1 | + | No | Yes | CCTA | CCTA | Yes |
| KQ003728.1 | KQ003728.1 | + | No | Yes | GAAG | GAAG | Yes |
| KQ004162.1 | KQ004162.1 | + | No | Yes | TGAC | TGAC | Yes |
| LIDP01000002.1_b | LIDP01000002.1 | + | No | Yes | GAAC | GAAC | Yes |
| LIDP01000012.1 | LIDP01000012.1 | - | No | Yes | CAAG | CAAG | Yes |
| LIDP01000017.1 | LIDP01000017.1 | - | No | Yes | CCAC | CACC | No |
| LT634572.1_a | LT634572.1 | - | No | Yes | CCTT | CCTT | Yes |
| LT634572.1_b | LT634572.1 | - | Yes | No | CCTT | CCTT | Yes |
| LUXQ01001340.1 | LUXQ01001340.1 | + | No | Yes | CAAG | CAAG | Yes |
| LUXQ01042183.1 | LUXQ01042183.1 | + | Yes | Yes | ATAC | ATAC | Yes |

| | | | | | | | |
|---|---|---|---|---|---|---|---|
| LUXR01004647.1 | LUXR01004647.1 | - | Yes | Yes | GTTC | CTTC | No |
| LUXR01022139.1 | LUXR01022139.1 | + | Yes | Yes | CCAC | CACC | No |
| LUXR01037956.1 | LUXR01037956.1 | + | No | Yes | GTTG | GTTG | Yes |
| LUXR01057621.1 | LUXR01057621.1 | + | No | Yes | GTAT | GTAT | Yes |
| LUXR01063446.1 | LUXR01063446.1 | - | No | Yes | CAAG | CAAG | Yes |
| LUXR01088996.1 | LUXR01088996.1 | - | No | Yes | TGAC | TGAC | Yes |
| LUXS01013084.1 | LUXS01013084.1 | - | No | Yes | CAAG | CAAG | Yes |
| LUXS01031219.1 | LUXS01031219.1 | - | No | Yes | GTTG | GTTG | Yes |
| LUXS01060324.1 | LUXS01060324.1 | + | No | Yes | GTCT | GTCT | Yes |
| LUXS01071149.1 | LUXS01071149.1 | - | No | Yes | GAAG | GAAG | Yes |
| LUXT01085568.1 | LUXT01085568.1 | - | Yes | Yes | CAAG | CAAG | Yes |
| LUXU01008622.1 | LUXU01008622.1 | - | No | Yes | CAAG | CAAG | Yes |
| LUXU01065281.1 | LUXU01065281.1 | - | No | Yes | CAAC | CAAC | Yes |
| LUXV01020898.1 | LUXV01020898.1 | - | No | Yes | GTTG | GTTG | Yes |
| LUXV01061131.1 | LUXV01061131.1 | + | No | Yes | CAAG | CAAG | Yes |
| LUXW01040954.1 | LUXW01040954.1 | + | No | Yes | ATAC | ATAC | Yes |
| LUXW01070394.1 | LUXW01070394.1 | - | No | Yes | CAAG | CAAG | Yes |
| LUXW01074277.1 | LUXW01074277.1 | - | No | Yes | GTCT | GTCT | Yes |
| LUXX01045907.1 | LUXX01045907.1 | + | No | Yes | CCAC | CACC | No |
| LUXX01073178.1 | LUXX01073178.1 | + | No | Yes | CAAG | CAAG | Yes |
| LUXX01079974.1 | LUXX01079974.1 | + | No | Yes | GAAG | GAAG | Yes |
| LUXX01080744.1 | LUXX01080744.1 | + | No | Yes | GTTC | CTTC | No |
| LUXX01090830.1 | LUXX01090830.1 | - | No | Yes | GTTG | GTTG | Yes |
| LUXY01039945.1 | LUXY01039945.1 | + | No | Yes | CAAG | CAAG | Yes |
| LUXY01052675.1 | LUXY01052675.1 | - | No | Yes | GTCT | GTCT | Yes |
| LUXY01101100.1 | LUXY01101100.1 | + | No | Yes | CCAC | CACC | No |

*LTRs of PERVs are divided into 4 types (LTR A, B1, B2 and B3). If two different kinds of LTRs are flanking the PERV, the LTRs are divergent.

**Only TSDs flanking the intact 5’ and 3’ LTRs sequences were analyzed.

**Table S4.** The information of 142 mammal used for PERV-like sequences mining.

| No. | Order | Species name | Common name |
|---|---|---|---|
| 1 | Afrosoricida | *Chrysochloris asiatica* | Cape Golden Mole |
| 2 | Afrosoricida | *Echinops telfairi* | Small Madagascar Hedgehog |
| 3 | Carnivora | *Acinonyx jubatus* | Cheetah |
| 4 | Carnivora | *Ailuropoda melanoleuca* | Giant Panda |
| 5 | Carnivora | *Ailurus fulgens* | Lesser Panda |
| 6 | Carnivora | *Canis lupus familiaris* | Dog |
| 7 | Carnivora | *Felis catus* | Domestic Cat |
| 8 | Carnivora | *Leptonychotes weddellii* | Weddell Seal |
| 9 | Carnivora | *Lycaon pictus* | African Hunting Dog |
| 10 | Carnivora | *Mustela putorius furo* | Domestic Ferret |
| 11 | Carnivora | *Neomonachus schauinslandi* | Hawaiian Monk Seal |
| 12 | Carnivora | *Odobenus rosmarus divergens* | Pacific Walrus |
| 13 | Carnivora | *Panthera pardus* | Leopard |
| 14 | Carnivora | *Panthera tigris* | Tiger |
| 15 | Carnivora | *Ursus arctos* | Brown Bear |
| 16 | Carnivora | *Ursus maritimus* | Polar Bear |
| 17 | Cetartiodactyla | *Ammotragus lervia* | Aoudad |
| 18 | Cetartiodactyla | *Balaenoptera acutorostrata* | Minke Whale |
| 19 | Cetartiodactyla | *Balaenoptera bonaerensis* | Antarctic Minke Whale |
| 20 | Cetartiodactyla | *Bison* | American Bison |
| 21 | Cetartiodactyla | *Bos indicus* | Zebu Cattle |
| 22 | Cetartiodactyla | *Bos mutus* | Wild Yak |
| 23 | Cetartiodactyla | *Bos taurus* | Cattle |
| 24 | Cetartiodactyla | *Bubalus bubalis* | Water Buffalo |
| 25 | Cetartiodactyla | *Camelus bactrianus* | Bactrian Camel |
| 26 | Cetartiodactyla | *Camelus dromedarius* | Arabian Camel |
| 27 | Cetartiodactyla | *Camelus ferus* | Wild Bactrian Camel |
| 28 | Cetartiodactyla | *Capra aegagrus* | Wild Goat |
| 29 | Cetartiodactyla | *Capra hircus* | Goat |
| 30 | Cetartiodactyla | *Capreolus capreolus* | Western Roe Deer |
| 31 | Cetartiodactyla | *Cervus elaphus* | Red Deer |
| 32 | Cetartiodactyla | *Eschrichtius robustus* | Grey Whale |
| 33 | Cetartiodactyla | *Giraffa tippelskirchi* | Giraffa |
| 34 | Cetartiodactyla | *Lipotes vexillifer* | Yangtze River Dolphin |
| 35 | Cetartiodactyla | *Odocoileus virginianus* | White-Tailed Deer |
| 36 | Cetartiodactyla | *Okapia johnstoni* | Okapi |
| 37 | Cetartiodactyla | *Orcinus orca* | Killer Whale |
| 38 | Cetartiodactyla | *Ovis aries* | Sheep |
| 39 | Cetartiodactyla | *Ovis canadensis* | Bighorn Sheep |
| 40 | Cetartiodactyla | *Pantholops hodgsonii* | Chiru |

| | | | |
|---|---|---|---|
| 41 | Cetartiodactyla | *Physeter catodon* | Sperm Whale |
| 42 | Cetartiodactyla | *Tursiops truncatus* | Bottlenose Dolphin |
| 43 | Cetartiodactyla | *Vicugna pacos* | Alpaca |
| 44 | Chiroptera | *Eidolon helvum* | Straw-Colored Fruit Bat |
| 45 | Chiroptera | *Eptesicus fuscus* | Big Brown Bat |
| 46 | Chiroptera | *Hipposideros armiger* | Great Roundleaf Bat |
| 47 | Chiroptera | *Megaderma lyra* | Indian False Vampire |
| 48 | Chiroptera | *Miniopterus natalensis* | Natal Long-Fingered Bat |
| 49 | Chiroptera | *Myotis brandtii* | Brandt'S Bat |
| 50 | Chiroptera | *Myotis davidii* | David'S Myotis |
| 51 | Chiroptera | *Myotis lucifugus* | Little Brown Bat |
| 52 | Chiroptera | *Pteronotus parnellii* | Parnell'S Mustached Bat |
| 53 | Chiroptera | *Pteropus alecto* | Black Flying Fox |
| 54 | Chiroptera | *Pteropus vampyrus* | Large Flying Fox |
| 55 | Chiroptera | *Rhinolophus ferrumequinum* | Greater Horseshoe Bat |
| 56 | Chiroptera | *Rhinolophus sinicus* | Chinese Rufous Horseshoe Bat |
| 57 | Chiroptera | *Rousettus aegyptiacus* | Egyptian Rousette |
| 58 | Dasypodidae | *Dasypus novemcinctus* | Nine-Banded Armadillo |
| 59 | Dasyuromorphia | *Sarcophilus harrisii* | Tasmanian Devil |
| 60 | Dermoptera | *Galeopterus variegatus* | Sunda Flying Lemur |
| 61 | Didelphimorphia | *Monodelphis domestica* | Gray Short-Tailed Opossum |
| 62 | Diprotodontia | *Notamacropus eugenii* | Tammar Wallaby |
| 63 | Diprotodontia | *Phascolarctos cinereus* | Koala |
| 64 | Hyracoidea | *Procavia capensis* | Cape Rock Hyrax |
| 65 | Insectivora | *Condylura cristata* | Star-Nosed Mole |
| 66 | Insectivora | *Erinaceus europaeus* | Western European Hedgehog |
| 67 | Insectivora | *Sorex araneus* | European Shrew |
| 68 | Lagomorpha | *Ochotona princeps* | American Pika |
| 69 | Lagomorpha | *Oryctolagus cuniculus* | Rabbit |
| 70 | Macroscelidea | *Elephantulus edwardii* | Cape Elephant Shrew |
| 71 | Monotremata | *Ornithorhynchus anatinus* | Platypus |
| 72 | Perissodactyla | *Ceratotherium simum simum* | Southern White Rhinoceros |
| 73 | Perissodactyla | *Equus asinus* | Ass |
| 74 | Perissodactyla | *Equus caballus* | Horse |
| 75 | Perissodactyla | *Equus przewalskii* | Przewalski'S Horse |
| 76 | Pholidota | *Manis javanica* | Malayan Pangolin |
| 77 | Pholidota | *Manis pentadactyla* | Chinese Pangolin |
| 78 | Pilosa | *Choloepus hoffmanni* | Hoffmann'S Two-Fingered Sloth |
| 79 | Primates | *Aotus nancymaae* | Ma'S Night Monkey |
| 80 | Primates | *Callithrix jacchus* | White-Tufted-Ear Marmoset |
| 81 | Primates | *Carlito syrichta* | Philippine Tarsier |
| 82 | Primates | *Cebus capucinus* | White-Faced Sapajou |
| 83 | Primates | *Cercocebus atys* | Sooty Mangabey |
| 84 | Primates | *Chlorocebus sabaeus* | Green Monkey |

| | | | |
|---|---|---|---|
| 85 | Primates | *Colobus angolensis* | Angolan Colobus |
| 86 | Primates | *Daubentonia madagascariensis* | Aye-Aye |
| 87 | Primates | *Eulemur flavifrons* | Sclater'S Lemur |
| 88 | Primates | *Eulemur macaco* | Black Lemur |
| 89 | Primates | *Gorilla gorilla* | Western Gorilla |
| 90 | Primates | *Homo sapiens* | Human |
| 91 | Primates | *Macaca fascicularis* | Crab-Eating Macaque |
| 92 | Primates | *Macaca mulatta* | Rhesus Monkey |
| 93 | Primates | *Macaca nemestrina* | Pig-Tailed Macaque |
| 94 | Primates | *Mandrillus leucophaeus* | Drill |
| 95 | Primates | *Microcebus murinus* | Gray Mouse Lemur |
| 96 | Primates | *Nasalis larvatus* | Proboscis Monkey |
| 97 | Primates | *Nomascus leucogenys* | Northern White-Cheeked Gibbon |
| 98 | Primates | *Otolemur garnettii* | Small-Eared Galago |
| 99 | Primates | *Pan paniscus* | Pygmy Chimpanzee |
| 100 | Primates | *Pan troglodytes* | Chimpanzee |
| 101 | Primates | *Papio anubis* | Olive Baboon |
| 102 | Primates | *Pongo abelii* | Sumatran Orangutan |
| 103 | Primates | *Propithecus coquereli* | Coquerel'S Sifaka |
| 104 | Primates | *Rhinopithecus bieti* | Black Snub-Nosed Monkey |
| 105 | Primates | *Rhinopithecus roxellana* | Golden Snub-Nosed Monkey |
| 106 | Primates | *Saimiri boliviensis boliviensis* | Bolivian Squirrel Monkey |
| 107 | Proboscidea | *Loxodonta africana* | African Savanna Elephant |
| 108 | Rodentia | *Apodemus speciosus* | Large Japanese Field Mouse |
| 109 | Rodentia | *Apodemus sylvaticus* | European Woodmouse |
| 110 | Rodentia | *Castor canadensis* | American Beaver |
| 111 | Rodentia | *Cavia aperea* | Brazilian Guinea Pig |
| 112 | Rodentia | *Cavia porcellus* | Domestic Guinea Pig |
| 113 | Rodentia | *Chinchilla lanigera* | Long-Tailed Chinchilla |
| 114 | Rodentia | *Cricetulus griseus* | Chinese Hamster |
| 115 | Rodentia | *Dipodomys ordii* | Ord'S Kangaroo Rat |
| 116 | Rodentia | *Ellobius lutescens* | Transcaucasian Mole Vole |
| 117 | Rodentia | *Ellobius talpinus* | Northern Mole Vole |
| 118 | Rodentia | *Fukomys damarensis* | Damara Mole-Rat |
| 119 | Rodentia | *Heterocephalus glaber* | Naked Mole-Rat |
| 120 | Rodentia | *Ictidomys tridecemlineatus* | Thirteen-Lined Ground Squirrel |
| 121 | Rodentia | *Jaculus jaculus* | Lesser Egyptian Jerboa |
| 122 | Rodentia | *Marmota marmota* | European Marmot |
| 123 | Rodentia | *Meriones unguiculatus* | Mongolian Gerbil |
| 124 | Rodentia | *Mesocricetus auratus* | Golden Hamster |
| 125 | Rodentia | *Microtus agrestis* | Short-Tailed Field Vole |
| 126 | Rodentia | *Microtus ochrogaster* | Prairie Vole |
| 127 | Rodentia | *Mus caroli* | Ryukyu Mouse |

| | | | |
|---|---|---|---|
| 128 | Rodentia | *Mus musculus* | House Mouse |
| 129 | Rodentia | *Mus pahari* | Shrew Mouse |
| 130 | Rodentia | *Mus spretus* | Western Wild Mouse |
| 131 | Rodentia | *Myodes glareolus* | Bank Vole |
| 132 | Rodentia | *Nannospalax galili* | Upper Galilee Mountains Blind Mole Rat |
| 133 | Rodentia | *Neotoma lepida* | Desert Woodrat |
| 134 | Rodentia | *Octodon degus* | Degu |
| 135 | Rodentia | *Peromyscus maniculatus* | North American Deer Mouse |
| 136 | Rodentia | *Phodopus sungorus* | Djungarian Hamster |
| 137 | Rodentia | *Psammomys obesus* | Fat Sand Rat |
| 138 | Rodentia | *Rattus norvegicus* | Norway Rat |
| 139 | Scandentia | *Tupaia belangeri* | Northern Tree Shrew |
| 140 | Scandentia | *Tupaia chinensis* | Chinese Tree Shrew |
| 141 | Sirenia | *Trichechus manatus latirostris* | Florida Manatee |
| 142 | Tubulidentata | *Orycteropus afer* | Aardvark |

**Table S5.** The information of genes flanking the eJJRVs and ePCRVs.

| Name | Accession numbers | Gene | | |
|---|---|---|---|---|
| | | Name | Product | Accession numbers |
| eJJRV | NW_004504334.1 | Rab10 | ras-related protein Rab-10 | XM_004663738 |
| | | Gareml | GRB2-associated and regulator of MAPK protein-like isoform X2 | XM_004663737 |
| | | Hadha | trifunctional enzyme subunit alpha, mitochondrial | XM_004663732 |
| | NW_004504375.1 | U2surp | U2 snRNP-associated SURP motif-containing protein isoform X2 | XM_004663625 |
| | | Trpc1 | short transient receptor potential channel 1 isoform X3 | XM_004663621 |
| | | Atr | serine/threonine-protein kinase ATR | XM_004663591 |
| | NW_004504378.1 | Adcy3 | adenylate cyclase type 3 isoform X2 | XM_004663749 |
| | | Dnmt3a | DNA (cytosine-5)-methyltransferase 3A | XM_004663744 |
| | | Efr3b | protein EFR3 homolog B | XM_012949933 |
| | NW_004504445.1 | Brwd3 | bromodomain and WD repeat-containing protein 3 | XM_004669595 |
| | | Rps6ka6 | ribosomal protein S6 kinase alpha-6 isoform X1 | XM_004669599 |
| | | Apool | MICOS complex subunit MIC27 | XM_004669601 |
| ePCRV | KN678690.1 | LOC101360595 | potassium voltage-gated channel subfamily C member 3 | XM_023736208 |
| | | LRRC4B | leucine-rich repeat-containing protein 4B | XM_003406838 |
| | KN676491.1 | TMED9 | transmembrane emp24 domain-containing protein 9 | XM_003404689 |
| | KN678005.1 | TIGD1 | tigger transposable element-derived protein 1 | XM_004092051 |
| | KN677924.1 | CTTNBP2NL | CTTNBP2 N-terminal-like protein isoform X2 | XM_008515240 |
| | | CAPZA1 | F-actin-capping protein subunit alpha-1 | XM_007949157 |
| | KN676905.1 | LMX1A | LIM homeobox transcription factor 1-alpha | XM_006876512 |
| | KN676182.1 | CDKN2AIP | CDKN2A-interacting protein | XM_010595654 |
| | | ING2 | inhibitor of growth protein 2 | XM_023554959 |
| | | IRF2 | interferon regulatory factor 2 | XM_007940873 |
| | KN676638.1 | HTR2A | 5-hydroxytryptamine receptor 2A | XM_023551308 |

**Table S6.** The information of full-length and near full-length *Muroidea* ERV.

| Species | Name | Locus | Orientation (+/-) | Start | End |
|---|---|---|---|---|---|
| *Apodemus speciosus* (large Japanese field mouse) | ASERV_1 | BDUI01007340.1 | + | 7091 | 18424 |
| *Apodemus speciosus* (large Japanese field mouse) | ASERV_2 | BDUI01011695.1 | + | 27086 | 39487 |
| *Apodemus speciosus* (large Japanese field mouse) | ASERV_3 | BDUI01014616.1 | + | 4402 | 16562 |
| *Apodemus speciosus* (large Japanese field mouse) | ASERV_4 | BDUI01165896.1 | + | 43 | 2078 |
| *Mus spretus* (western wild mouse) | MSERV_1 * | CM004094.1 | + | 12428920 | 12433423 |
| *Mus spretus* (western wild mouse) | MSERV_2 * | CM004094.1 | + | 13440056 | 13445675 |
| *Mus spretus* (western wild mouse) | MSERV_3 * | CM004094.1 | + | 39404880 | 39411011 |
| *Mus spretus* (western wild mouse) | MSERV_4 | CM004094.1 | + | 53370667 | 53377344 |
| *Mus spretus* (western wild mouse) | MSERV_5 * | CM004095.1 | + | 124164246 | 124168804 |
| *Mus spretus* (western wild mouse) | MSERV_6 * | CM004099.1 | + | 64159005 | 64161997 |
| *Mus spretus* (western wild mouse) | MSERV_7 | CM004100.1 | + | 19124698 | 19134398 |
| *Mus spretus* (western wild mouse) | MSERV_8 * | CM004100.1 | - | 49390701 | 49397604 |
| *Mus spretus* (western wild mouse) | MSERV_9 * | CM004104.1 | - | 147121298 | 147126766 |
| *Mus spretus* (western wild mouse) | MSERV_10 * | CM004104.1 | + | 63559926 | 63563981 |
| *Mus spretus* (western wild mouse) | MSERV_11 | CM004105.1 | + | 21055717 | 21062066 |
| *Mus spretus* (western wild mouse) | MSERV_12 | CM004108.1 | - | 49541470 | 49544422 |
| *Mus spretus* (western wild mouse) | MSERV_13 * | CM004110.1 | + | 18030051 | 18038614 |
| *Mus spretus* (western wild mouse) | MSERV_14 | CM004112.1 | + | 74127053 | 74129822 |
| *Mus musculus* (house mouse) | MMERV_1 | NC_000067.6 | - | 195342993 | 195357606 |
| *Mus musculus* (house mouse) | MMERV_2 | NC_000067.6 | + | 54602252 | 54609807 |
| *Mus musculus* (house mouse) | MMERV_3 * | NC_000067.6 | + | 58633572 | 58652013 |
| *Mus musculus* (house mouse) | MMERV_4 * | NC_000069.6 | - | 4910069 | 4915466 |
| *Mus musculus* (house mouse) | MMERV_5 | NC_000069.6 | - | 96363402 | 96371676 |
| *Mus musculus* (house mouse) | MMERV_6 | NC_000070.6 | - | 108146034 | 108155016 |
| *Mus musculus* (house mouse) | MMERV_7 | NC_000070.6 | + | 122031962 | 122039933 |
| *Mus musculus* (house mouse) | MMERV_8 * | NC_000070.6 | + | 146132189 | 146146516 |
| *Mus musculus* (house mouse) | MMERV_9 * | NC_000070.6 | - | 147653462 | 147667936 |

| | | | | | |
|---|---|---|---|---|---|
| *Mus musculus* (house mouse) | MMERV_10 | NC_000070.6 | + | 39383244 | 39395013 |
| *Mus musculus* (house mouse) | MMERV_11 | NC_000071.6 | + | 143773327 | 143782050 |
| *Mus musculus* (house mouse) | MMERV_12 | NC_000071.6 | + | 25230699 | 25239670 |
| *Mus musculus* (house mouse) | MMERV_13 | NC_000072.6 | - | 43917022 | 43925183 |
| *Mus musculus* (house mouse) | MMERV_14 * | NC_000072.6 | - | 69383698 | 69389523 |
| *Mus musculus* (house mouse) | MMERV_15 | NC_000072.6 | - | 73287906 | 73299314 |
| *Mus musculus* (house mouse) | MMERV_16 | NC_000073.6 | + | 29615207 | 29623905 |
| *Mus musculus* (house mouse) | MMERV_17 | NC_000073.6 | + | 29732067 | 29742864 |
| *Mus musculus* (house mouse) | MMERV_18 | NC_000073.6 | + | 42885280 | 42897233 |
| *Mus musculus* (house mouse) | MMERV_19 | NC_000073.6 | + | 42885280 | 42897233 |
| *Mus musculus* (house mouse) | MMERV_20 | NC_000075.6 | + | 109463789 | 109474910 |
| *Mus musculus* (house mouse) | MMERV_21 | NC_000076.6 | + | 25895611 | 25904025 |
| *Mus musculus* (house mouse) | MMERV_22 | NC_000076.6 | - | 26926972 | 26935378 |
| *Mus musculus* (house mouse) | MMERV_23 | NC_000076.6 | - | 79420549 | 79426320 |
| *Mus musculus* (house mouse) | MMERV_24 | NC_000077.6 | + | 23527118 | 23533328 |
| *Mus musculus* (house mouse) | MMERV_25 | NC_000079.6 | - | 22308443 | 22316117 |
| *Mus musculus* (house mouse) | MMERV_26 | NC_000079.6 | - | 98986275 | 98995255 |
| *Mus musculus* (house mouse) | MMERV_27 | NC_000080.6 | + | 38108618 | 38117415 |
| *Mus musculus* (house mouse) | MMERV_28 * | NC_000081.6 | + | 33063653 | 33073552 |
| *Mus musculus* (house mouse) | MMERV_29 | NC_000082.6 | + | 56040893 | 56053089 |
| *Mus musculus* (house mouse) | MMERV_30 | NC_000083.6 | + | 66173605 | 66181792 |
| *Mus musculus* (house mouse) | MMERV_31 | NC_000083.6 | - | 94663897 | 94672405 |
| *Mus musculus* (house mouse) | MMERV_32 | NC_000085.6 | + | 38375022 | 38382083 |
| *Mus musculus* (house mouse) | MMERV_33 * | NC_000085.6 | + | 9310529 | 9316610 |
| *Mus musculus* (house mouse) | MMERV_34 * | NC_000085.6 | - | 9621419 | 9627503 |
| *Mus musculus* (house mouse) | MMERV_35 | NC_000086.7 | - | 121041808 | 121057724 |
| *Mus musculus* (house mouse) | MMERV_36 * | NC_000086.7 | - | 169136903 | 169141171 |
| *Mus musculus* (house mouse) | MMERV_37 * | NC_000086.7 | - | 8765246 | 8770067 |
| *Mus musculus* (house mouse) | MMERV_38 | NW_012132903.1 | + | 211516 | 215375 |
| *Mus musculus* (house mouse) | MMERV_39 * | NW_019168507.1 | + | 49361 | 63688 |

| | | | | | |
|---|---|---|---|---|---|
| *Rattus norvegicus* (Norway rat) | RNERV_1 | NC_005101.4 | + | 112839250 | 112846717 |
| *Rattus norvegicus* (Norway rat) | RNERV_2 | NC_005101.4 | - | 163930082 | 163941915 |
| *Rattus norvegicus* (Norway rat) | RNERV_3 | NC_005101.4 | - | 164073367 | 164085504 |
| *Rattus norvegicus* (Norway rat) | RNERV_4 | NC_005109.4 | - | 3476351 | 3488162 |
| *Rattus norvegicus* (Norway rat) | RNERV_5 * | NC_005112.4 | - | 89691210 | 89699756 |
| *Rattus norvegicus* (Norway rat) | RNERV_6 * | NC_005116.4 | - | 14138750 | 14147532 |
| *Mus caroli* (Ryukyu mouse) | MCERV_1 * | NC_034574.1 | - | 18620568 | 18628684 |
| *Mus caroli* (Ryukyu mouse) | MCERV_2 * | NC_034570.1 | + | 11917178 | 11923138 |
| *Mus caroli* (Ryukyu mouse) | MCERV_3 | NC_034570.1 | + | 11985084 | 11996915 |
| *Mus caroli* (Ryukyu mouse) | MCERV_4 | NC_034570.1 | + | 39517065 | 39523440 |
| *Mus caroli* (Ryukyu mouse) | MCERV_5 * | NC_034571.1 | + | 116393982 | 116398538 |
| *Mus caroli* (Ryukyu mouse) | MCERV_6 * | NC_034573.1 | - | 44194429 | 44202555 |
| *Mus caroli* (Ryukyu mouse) | MCERV_7 | NC_034576.1 | + | 28969181 | 28976746 |
| *Mus caroli* (Ryukyu mouse) | MCERV_8 | NC_034576.1 | + | 44091152 | 44099831 |
| *Mus caroli* (Ryukyu mouse) | MCERV_9 * | NC_034577.1 | + | 41218138 | 41222010 |
| *Mus caroli* (Ryukyu mouse) | MCERV_10 * | NC_034579.1 | - | 110036279 | 110046371 |
| *Mus caroli* (Ryukyu mouse) | MCERV_11 | NC_034580.1 | + | 116643636 | 116652385 |
| *Mus caroli* (Ryukyu mouse) | MCERV_12 | NC_034580.1 | + | 19360179 | 19372402 |
| *Mus caroli* (Ryukyu mouse) | MCERV_13 | NC_034585.1 | - | 18403459 | 18406398 |
| *Mus caroli* (Ryukyu mouse) | MCERV_14 | NC_034586.1 | - | 83266053 | 83274517 |
| *Mus caroli* (Ryukyu mouse) | MCERV_15 | NC_034587.1 | - | 3193064 | 3201350 |
| *Mus caroli* (Ryukyu mouse) | MCERV_16 | NC_034587.1 | + | 70801494 | 70813308 |
| *Mus caroli* (Ryukyu mouse) | MCERV_17 * | NC_034587.1 | - | 70826334 | 70832363 |
| *Mus caroli* (Ryukyu mouse) | MCERV_18 * | NC_034589.1 | + | 76573015 | 76577046 |
| *Mus caroli* (Ryukyu mouse) | MCERV_19 | NW_018389896.1 | + | 3316 | 8447 |
| *Mus caroli* (Ryukyu mouse) | MCERV_20 | NW_018390126.1 | + | 36367 | 42383 |
| *Mus pahari* (shrew mouse) | MPERV_1 | NC_034605.1 | + | 9841686 | 9845137 |
| *Mus pahari* (shrew mouse) | MPERV_2 * | NC_034592.1 | - | 115548487 | 115554442 |
| *Mus pahari* (shrew mouse) | MPERV_3 | NC_034593.1 | + | 40652727 | 40659930 |
| *Mus pahari* (shrew mouse) | MPERV_4 * | NC_034597.1 | - | 18909417 | 18917267 |

| | | | | | |
|---|---|---|---|---|---|
| *Mus pahari* (shrew mouse) | MPERV_5 * | NC_034597.1 | - | 79286612 | 79292922 |
| *Mus pahari* (shrew mouse) | MPERV_6 * | NC_034601.1 | - | 77665113 | 77669641 |
| *Mus pahari* (shrew mouse) | MPERV_7 | NC_034602.1 | - | 10135886 | 10140830 |
| *Mus pahari* (shrew mouse) | MPERV_8 | NC_034602.1 | + | 26223471 | 26226782 |
| *Mus pahari* (shrew mouse) | MPERV_9 | NC_034602.1 | + | 8076595 | 8087725 |
| *Mus pahari* (shrew mouse) | MPERV_10 * | NC_034604.1 | - | 244802 | 249579 |
| *Mus pahari* (shrew mouse) | MPERV_11 * | NC_034604.1 | + | 64465704 | 64470420 |
| *Mus pahari* (shrew mouse) | MPERV_12 | NW_018391913.1 | - | 158314 | 169263 |
| *Mus pahari* (shrew mouse) | MPERV_13 | NW_018392282.1 | - | 41850 | 46797 |
| *Mus pahari* (shrew mouse) | MPERV_14 * | NW_018392790.1 | + | 17443 | 23944 |
| *Mus pahari* (shrew mouse) | MPERV_15 | NW_018393646.1 | + | 11018 | 15870 |

* The full-length ERVs containing both 5' and 3' LTR. ERVs without * represent those only having one LTR.

**Table S7.** The information of representative retroviruses used for phylogenetic analysis.

| **Virus name** | **Abbreviation** | **Natural host** | **Accession number** |
|---|---|---|---|
| Baboon endogenous virus | BaEV | Baboon | D10032 |
| Feline leukemia virus | FeLV | Cat | NC_001940 |
| Friend murine leukemia virus | F-MuLV | Mouse | NC_001362 |
| Gibbon ape leukemia virus | GALV | Gibbon | NC_001885 |
| Koala retrovirus | KoRV | Koala | NC_021704 |
| Murine type C retrovirus | M-CRV | Mouse | NC_001702 |
| Moloney murine leukemia virus | M-MuLV | Mouse | NC_001501 |
| Mus dunni endogenous retrovirus | MDEV | Mouse | AF053745 |
| Orcinus orca endogenous retrovirus | OOEV | Killer whale | GQ222416 |
| Porcine endogenous retrovirus A | PERV-A | Pig (*Sus scrofa*) | AJ293656 |
| Porcine endogenous retrovirus B | PERV-B | Pig (*Sus scrofa*) | AY099324 |
| Porcine endogenous retrovirus C | PERV-C | Pig (*Sus scrofa*) | HQ536015 |
| RD114 retrovirus | RD114 | Cat | NC_009889 |
| Reticuloendotheliosis virus | REV | Chicken | NC_006934 |
| Rauscher murine leukemia virus | R-MuLV | Mouse | NC_001819 |
| Rhinolophus ferrumequinum retrovirus | RfRV | Greater horseshoe bat | JQ303225 |
