## Supplementary material for "Genomic insights into evolutionary journey of the porcine endogenous retroviruses"

**Data set S1.** The alignments used to build the phylogenetic trees of Gag, Pol and Env represented in Fig. S4, S5, and S6, respectively.

```
Gag
114 225
LUXW01011341.1       NHPPFSEDPQ RLTGLVESLM FSHQPTWDDC QQLLQTLFTT EERERILLEA
LUXS01031219.1       NHPPFSEDPQ RLTGLVESLM FSHQPTWDDC QQLLQTLFTT EERERILLEA
LUXX01090830.1       NHPPFSEDPQ RLTGLVESLM FSHQPTWDDC QQLLQTLFTT EERERILLEA
LT634572.1_d         NHPPFSEDPQ RLTGLVESLM FSHQPTWDDC QQLLQTLFTT EERERILLEA
CM000818.5_a         NHPPFSEDPQ RLTGLVESLM FSHQPTWDDC QQLLQTLFTT EERERILLEA
LUXS01011274.1       NHPPFSEDPQ RLTGLVESLM FSHQPTWDDC QQLLQTLFTT EERERILLEA
LUXU01062474.1       NHPPFSEDPQ RLTGLVESLM FSHQPTWDDC QQLLQTLFTT EERERILLEA
LUXX01045907.1       NHPPFSEDPQ RLTGLVESLM FSHQPTWDDC QQLLQTLFTT EERERILLEA
LUXX01080744.1       NHPPFSEDPQ RLTGLVESLM FSHQPTWDDC QQLLQTLFTT EERERILLEA
LUXU01038095.1       NHPPFSEDPQ RLTGLVESLM FSHQPTWDDC QQLLQTLFTT EERERILLEA
PERV_C               NHPPFSEDPQ RLTGLVESLM FSHQPTWDDC QQLLQTLFTT EERERILLEA
LUXW01089741.1       NHPPFSEDPQ RLTGLVESLM FSHQPTWDDC QQLLQTLFTT EERERILLEA
LUXT01005398.1       NHPPFSEDPQ RLTGLVESLM FSHQPTWDDC QQLLQTLFTT EERERILLEA
CM000826.5_c         NHPPFSEDPQ RLTGLVESLM FSHQPTWDDC QQLLQTLFTT EERERILLEA
CM000812.5_b         NHPPFSEDPQ RLTGLVESLM FSHQPTWDDC QQLLQTLFTT EERERILLEA
CM000814.5_a         NHPPFSEDPQ RLTGLVESLM FSHQPTWDDC QQLLQTLFTT EERERILLEA
CM000824.5_a         NHPPFSEDPQ RLTGLVESLM FSHQPTWDDC QQLLQTLFTT EERERILLEA
CM000812.5_a         NHPPFSEDPQ RLTGLVESLM FSHQPTWDDC QQLLQTLFTT EERERILLEA
PERV-A               NHPPFSEDPQ RLTGLVESLM FSHQPTWDDC QQLLQTLFTT EERKRILLEA
CM000816.5_b         NHPPFSEDPQ RLTGLVESLM FSHQPTWDDC QQLLQTLFTT EERERILLEA
AOCR01107158.1       NHPPFSEDPQ RLTGLVESLM FSHQPTWDDC QQLLQTLFTT EERERILLEA
CM000825.5_b         NHPPFSEDPQ RLTGLVESLM FSHQPTWDDC QQLLQTLFTT EERERILLEA
AOCR01079940.1       NHPPFSEDPQ RLTGLVESLM FSHQPTWDDC QQLLQTLFTT EERERILLEA
AOCR01098665.1       NHPPFSEDPQ RLTGLVESLM FSHQPTWDDC QQLLQTLFTT EERERILLEA
KQ004162.1           NHPPFSEDPQ RLTALVESLM FSHQPTWDDC QQLLQTLFTT EERERILLEA
CM000819.5_b         NHPPFSEDPQ RLTGLVESLM FSHQPTWDDC QQLLQTLFTT EERERILLEA
CM000823.5_b         NHPPFSEDPQ RLTGLVESLM FSHQPTWDDC QQLLQTLFTT EERERILLEA
CM000828.5_a         NHPPFSEDPQ RLTGLVESLM FSHQPTWDNC QQLLQTLFTT EERERILLEA
AEMK02000137.1       NHPPFSEDPQ RLTGLVESLM FSHQPTWDDC QQLLQTLFTT EERERILLEA
CM000820.5           NHPPFSEDPQ RLTGLVESLM FSHQPTWDDC QQLLQTLFTT EERERILLEA
CM000814.5_c         NHPPFSEDPQ RLTGLVESLM FSHQPTWDDC QQLLQTLFTT EERERILLEA
CM000812.5_c         NHPPFSEDPQ RLTGLVESLM FSHQPTWDDC QQLLQTLFTT EERERILLEA
AEMK02000197.1       NHPPFSEDPQ RLTGLVESLM FSHQPTWDDC QQLLQTLFTT EERERILLEA
CM000827.5           NHPPFSEDPQ RLTGLVESLM FSHQPTWDDC QQLLQTLFTT EERERILLEA
LUXR01057621.1       NHPPFSEDPQ RLTGLVESLM FSHQPTWDDC QQLLQTLFTT EERERILLEA
LUXV01020898.1       NHPPFSEDPQ RLTGLVESLM FSHQPTWDDC QQLLQTLFTT EERERILLEA
LUXS01071149.1       NHPPFSEDPQ RLTGLVESLM FSHQPTWDDC QQLLQTLFTT EERERILLEA
PERV-B               NHPPFSEDPQ RLTGLVESLM FSHQPTWDDC QQLLQTLFTT EERERILLEA
```

| | | | | | |
|---|---|---|---|---|---|
| LUXR01088996.1 | NHPPFSEDPQ | RLTGLVESLM | FSHQPTWDDC | QQLLQTLFTT | EERERILLEA |
| CM000825.5_a | NHPPFSEDPQ | RLTGLVESLM | FSHQPTWDDC | QQLLQTLFTT | EERERILLEA |
| CM000819.5_a | NHPPFSEDPQ | RLTGLVESLM | FSHQPTWDDC | QQLLQTLFTT | EERERILLEA |
| CM000826.5_a | NHPPFSEDPQ | RLTGLVESLM | FSHQPTWDDC | QQLLQTLFTT | EERERILLEA |
| CM000815.5 | NHPPFSEDPQ | RLTGLVESLM | FSHQPTWDDC | QQLLQTLFTT | EE-ERILLEA |
| CM000822.5_b | NHPPFSEDPQ | RLTGLVESLM | FSHQPTWDDC | QQLLQTLFTT | E--ERILLEA |
| LUXV01022316.1 | NHPPFSEDPQ | RLTGLVESLM | FSHQPTWDDC | QQLLQTLFTT | EERERILLEA |
| LUXY01101100.1 | NHPPFSEDPQ | RLTGLVESLM | FSHQPTWDDC | QQLLQTLFTT | EERERILLEA |
| CM000828.5_b | NHPPFSEDPQ | CLTGLVESLM | FSHQPTWDDC | QQLLQTLFTT | EERERILLEA |
| CM000824.5_b | NHPPFSEDPQ | RLTGLVESLM | FSHQPTWDDC | QQLLQTLFTT | EERERILLEA |
| LUXR01004647.1 | NHPPFSEDPQ | RLTGLVESLM | FSHQPTWDDC | QQLLQTLFTT | EERERILLEA |
| CM000818.5_b | NHPPFSEDPQ | RLTGLVESLM | FSHQPTWDDC | QQLLQTLFTT | EERERILLEA |
| CM000830.5_d | NHPPFSEDPQ | RLMGLVESLM | FSHQPTWDDC | QQLLQTLFTT | EE-ERILLEA |
| CM000814.5_b | NHPPFSEDPQ | RLTGLVESLM | FSHQPTWDNC | QQLLQTLFTT | EERERILLEA |
| AEMK02000476.1 | NHPPFSEDPQ | RLTGLVESLM | FSHQPTWDDC | QQLLQTLFTT | EE-ERILLEA |
| LUXS01081946.1 | NHPPFSEDPQ | RLTGLVESLM | FSHQPTWDDF | QQLLQTLFTT | EGRERILLEA |
| CM000828.5_c | NHPPFSEDPQ | RLTGLVESLM | FSHQPTWDDC | QQLLQTLFTT | EGRERILLEA |
| LUXQ01076544.1 | NHPPFSEDPQ | RLTGLVESLM | FSHQPTWDDC | QQLLQTLFTT | EGRERILLEA |
| LIDP01000002.1_a | NHPPFSEDPQ | R---LVESLM | FSHQPTWDDC | QQLLQTLFTT | EERERILLEA |
| CM000816.5_a | NHPPFSEDPQ | RLTALVESLM | FSHQPTWNDC | QQLLQTLFTT | EKRERILLEA |
| AEMK02000141.1 | NHPPFSEDPQ | RLTALVESLM | FSHQPTWDDC | QQLLQTLFTT | EERERILLET |
| AOCR01031838.1 | NHPPFSEDPQ | CLTGLVESLM | FSHQPTWDDC | QQLLQTLFTT | EERERILLEA |
| CM000813.5 | NHPPFSEDPQ | CLTGLVESLM | FSHQPTWDDC | QQLLQTLSTT | GERERILLEA |
| LIDP01000002.1_b | NHPPFSEDPQ | CLTGLVESLM | FSHQPTWDDC | QQLLQTLSTT | GERERILLEA |
| AEMK02000133.1 | NHPPFSEDPQ | HLTGLVESLM | FSHQPTWDDC | QQLLQTLFTT | EERERILLEA |
| LUXU01085042.1 | NHPPFSEDPQ | RLTGLVESLM | FSHQPTWDDC | QQLLQTLFTT | EERERILLEA |
| AEMK02000393.1_a | NHPPFSEDPQ | CLVGLVESLM | FSHQPTWDDC | QQLLQTLFTT | EERERILLEA |
| LUXV01069687.1 | NHPPFSEDPQ | CLVGLVESLM | FSHQPTWDDC | QQLLQTLFTT | EERERILLEA |
| LUXY01013808.1 | NHPPFSEDPQ | CLVGLVESLM | FSHQPTWDDC | QQLLQTLFTT | EERERILLEA |
| LUXX01033707.1 | NHPPFSEDPQ | CLVGLVESLM | FSHQPTWDDC | QQLLQTLFTT | EERERILLEA |
| LUXS01081675.1 | NHPPFSEDPQ | RLTGLVESLM | FSHQPTWDDC | QQLLQTLFTT | EERERILLEA |
| CM000830.5_b | NHPPFSEDPQ | CLTGLVESLM | FSHQPTWDDC | QQLLQTLFTT | EE-ERILLEA |
| LT634572.1_f | NHPPFSEDPQ | RLTALVESLM | FSHQPTWDNC | QQLLQTLFTT | EKRERILLEA |
| LT634572.1_g | NHPPFSEDPQ | RLTALVESLM | FSHQPTWDNC | QQLLQTLFTT | EKRERILLEA |
| LUXY01042539.1 | NHPPFSEDPQ | RLTALVESLM | FSHQPT-DNC | QQLLQTLFTT | EEREKILLEA |
| LT634572.1_c | NHPPFSEDPQ | RLTALVESLM | FSHQPTWDDC | QQLLQTHFTT | EERERILLKA |
| CM000822.5_a | NHPPFSEDPQ | CLTALVESLM | FSHQPTWDDC | QQLLQTLFTT | EERERILLEA |
| LT634572.1_a | NHPPSSEDPQ | HLTGLVESLM | FSHQPTWYDC | QQLLQTLFTT | EERERILLEA |
| LT634572.1_b | NHPPSSEDPQ | HLTGLVESLI | FSHQPTWYDC | QQLLQTLFTT | EE-ERILLEA |
| AJKK01232762.1 | NHPPSSEDPQ | HLTGLVESLI | FSHQPTWYDC | QQLLQTLFTT | EERERILLEA |
| LUXW01074277.1 | NHPPFSEDPQ | RLTGLVESLM | FSHQPTWDDC | QQLLQTLFTT | EERERILLEA |
| LUXX01006697.1 | NHPPFSEDPQ | RLTGLVESLM | FSHQPTWDDC | QQLLQTLFTT | EERERILLEA |
| LUXT01031363.1 | NHPPFSEDPQ | RLTGLVESLM | FSHQPTWDDC | QQLLQTLFTT | EERERILLEA |
| LUXY01052675.1 | NHPPFSEDPQ | RLTGLVESLM | FSHQPTWDDC | QQLLQTLFTT | EERERILLEA |

```
LUXQ01112116.1       NHPPFSEDPQ RLTGLVESLM FSHQPTWDDC QQLLQTLFTT EERERILLET
LUXV01092473.1       NHPPFSEDPQ RLTGLVESLM FSHQPTWDDC QQLLQTLFTT EERERILLEA
LUXS01060324.1       NHPPFSEDPQ RLTGLVESLM FSHQPTWDDC QQLLQTLFTT EE-ERILLET
KQ001904.1           NYPPFSEDPQ CLTGLVESLM FSHQPTWDDC QQLLQTLFTT EERERILLEA
CM000830.5_c         NHPPFSENPQ SLMGLVESLM FSHQPTWDDC QQQLQTLFTT EEREKILLEA
LUXU01061831.1_a     NHPPFSEDPQ SLMGLVESLM FSHQPTWDDC QQQLQTLFTT EEREKILLEA
LUXU01061831.1_b     NHPPFSEDPQ SLMGLVESLM FSHQPTWDDC QQQLQTLFTT EEREKILLEA
LUXW01038847.1       NHPPFSENPQ SLMGLVESLM FSHQPTWDDC QQQLQTLFTT EEREKILLEA
LUXX01056080.1       NHPPFSENPQ SLMGLVESLM FSHQPTWDDC QQQLQTLFTT EEREKILLEA
LUXS01056853.1       NHPPFSENPQ SLMGLVESLM FSHQPTWDDC QQQLQTLFTT EEREKILLEA
AEMK02000536.1       NHPPFSEDPQ RLTGLVESLM FSHQPTWDDC QQLLQTLFTT EERERILLEA
LUXT01067056.1       NHPPFSEDPQ RLTALVESLM FSHQPTWDDC QQLLQTLFTT EERERILLET
CM000826.5_b         NHPPFSEDPQ CLTGL-GSLM FSHQPTWDDC QQLLQTLSTT EERERILLEA
LUXY01106724.1       NHPPFLEDPQ CLTGLGESLM FSHQPTWDDC QQLLQTLSTT EERERILLEA
LIDP01000015.1       NHPPFSEDPQ CLTGLGESLM FSHQPTWDDC QQLLQTLSTT EERERILLEA
eJJRV_NW_004504375.1 NHPPFSEEPW HLTGLIESLM FSHQPTWDDC QQLLQTLFTT KEQERILIEA
eJJRV_NW_004504334.1 KYPPFSEEPR RLTVLIESLM FSHQPTWDD- QQLLQTLFTT EEREGILIEA
ePCRV_KN678005.1     NHPSFSDDPQ KLTSLMESLM FSHQPTWDDC QQLLQVLFTT EEHERILLEA
ePCRV_KN676905.1     NHPSFSDDPQ KLTSLMESLM FSHQPTWEDC QQLLQVLFTT EERERILLKA
MDEV                 NHPPFSENPS GLTGLLESLM FSHQPTWDDC QQLLQVLFTT EERERILMEA
KoRV                 NHPSFSENPT GLTGLLESLM FSHQPTWDDC QQLLQVLFTT EERERILLEA
GALV                 NHPSFSENPA GLTGLLESLM FSHQPTWDDC QQLLQILFTT EERERILLEA
OOEV                 HNPTFSENPQ ALTALIESLV FSHQPTWDDC QQLLQTLLTT EERQRVLLEA
RD114                HNPSFSQEPQ ALTSLIESIL LTHQPTWDDC QQLLQVLLTT EERQRVLLEA
BaEV                 HNPSFSQDPQ ALTSLIESIL LTHQPTWDDC QQLLQVLLTT EERQRVLLEA
FeLV                 HNPPFSQDPV ALTNLIESIL VTHQPTWDDC QQLLQALLTG EERQRVLLEA
M-MuLV               NNPSFSEDPG KLTALIESVL ITHQPTWDDC QQLLGTLLTG EEKQRVLLEA
F-MuLV               NNPSFSEDPA KLTALIESVL LTHQPTWDDC QQLLGTLLTG EEKQRVLLEA
R-MuLV               NNPSFSEDPG KLTALIESVL LTHQPTWDDC QQLLGTLLTG EEKQRVLLEA
M-CRV                NNPSFSEDPG KLTALIESVL TTHQPTWDDC QQLLGTLLTG EEKQRVLLEA
RfRV                 QNPPFSEDPK GLTDLFESVM HTHSPTWDDC QQLLKTLFTT EERERILTEA
REV                  QNPSFSQAPD EVISLLESVF YTHQPTWDDC QQLLRTLFTT EERERVRTES

                     RKNVPGADGR PTQLQNEIDM GFPLTRPGWD YNTAEGRESL KIYRQALVAG
                     RKNVPGADGR PTQLQNEIDM GFPLTRPGWD YNTAEGRESL KIYRQALVAG
                     RKNVPGADGR PTQLQNEIDM GFPLTRPGWD YNTAEGRESL KIYRQALVAG
                     RKNVPGADGR PTQLQNEIDM GFPLTRPGWD YNTAEGRESL KIYRQALVAG
                     RKNVPGADGR PTQLQNEIDM GFPLTRPGWD YNTAEGRESL KIYRQALVAG
                     RKNVPGADGR PTQLQNEIDM GFPLTRPGWD YNTAEGRESL KIYRQALVAG
                     RKNVPGADGR PTQLQNEIDM GFPLTRPGWD YNTAEGRESL KIYRQALVAG
                     RKNVPGADGR PTQLQNEIDM GFPLTRPGWD YNTAEGRESL KIYRQALVAG
                     RKNVPGADGR PTQLQNEIDM GFPLTRPGWD YNTAEGRESL KIYRQALVAG
                     RKNVPGADGR PTQLQNEIDM GFPLTRPGWD YNTAEGRESL KIYRQALVAG
                     RKNVPGADGR PTQLQNEIDM GFPLTRPGWD YNTAEGRESL KIYRQALVAG
```

RKNVPGADGR PTQLQNEIDM GFPLTRPGWD YNTAEGRESL KIYRQALVAG
RKNVPGADGR PTQLQNEIDM GFPLTRPGWD YNTAEGRESL KIYRQALVAG
RKNVPGADGR PTQLQNEIDM GFPLTRPGWD YNTAEGRESL KIYRQALVAG
RKNVPGADGR PTQLQNEIDM GFPLTRPGWD YNTAEGRESL KIYRQALVAG
RKNVPGADGR PTQLQNEIDM GFPLTRPGWD YNTAEGRESL KIYRQALVAG
RKNVPGADGR PTQLQNEIDM GFPLTRPGWD YNTAEGRESL KIYRQALVAG
RKNVPGADGR PTQLQNEIDM GFPLTRPGWD YNTAEGRESL KIYRQALVAG
RKNVPGADGR PTQLQNEIDM GFPLTRPGWD YNTAEGRESL KIYRQALVAG
RKNVPGADGR PTQLQNEIDM GFPLTRPGWD YNTAEGRESL NIYRQALVAG
RKNVPGADGR PTRLQNEIDM GFPLTRPGWD YNTAEGRESL KIYRQALVAG
RKNVPGADGR PTRLQNEIDM GFPLTRPGWD YNTAEGRESL KIYRQALVAG
RKNVPGADGR PTRLQNEIDM GFPLTRPGWD YNTAEGRESL KIYRQALVAG
RKNVPGADGR PTRLQNEIDM GFPLTRPGWD YNTAEGRESL KIYRQALVAG
RKNVPGADGR PTQLQNEIDM GFPLTRPGWD YNTAEGRESL KIYRQALVAG
RKNVPGADGR PTQLQNEIDM GFPLTRPGWD YNTAEGRESL KIYRQALVAG
RKNVPGADGR PTQLQNEIDM GFPLTRPGWD YNTAEGRESL KIYRQALVAG
KKNVPGADGR PTQLQNEIDM GFPLTRPGWD YNTAEGRESL KIYRQALVAG
RKNVPGADGR PTQLQNEIDM GFPLTRPGWD YNTAEGRESL KIYCQALVAG
RKNVPGADGR PTQLQNEIDM GFPLTRPGWD YNTAEGRESL KIYRQALVAG
RKNVPGADGR PTQLQNEIDM GFPLTRPGWD YNTAEGRESL KIYRQALVAG
RKNVPGADGR PTQLQNEIDM GFPLTRPGWD YNTAEGRESL KIYRQALVAG
RKNVPGADGR PTQLQNEIDM GFPLTRPGWD YNTAEGRESL KIYRQALVAG
RKNVPGADGR PTQLQNEIDM GFPLTRPGWD YNTAEGRESL KIYRQALVAG
RKNVPGADGR PTQLQNEIDM GFPLTRPGWD YNTAEGRESL KIYRQALVAG
RKNVPGADGR PTQLQNEIDM GFPLTRPGWD YNTAEGRESL KIYRQALVAG
RKNVPGADGR PTQLQNEIDM GFPLTRPGWD YNTAEGRESL KIYRQALVAG
RKNVPGADGR PTQLQNEIDM GFPLTRPGWD YNTAEGRESL KIYRQALVAG
RKNVPGADGR PTQLQNEIDM GFPLTRPGWD YNTAEGRESL KIYRQALVAG
RKNVPGADGR PTQLQNEIDM GFPLTRPGWD YNTAEGRESL KIYRQALVAG
RKNVPGADGR PTQLQNEIDM GFPLTRPGWD YNTAEGRESL KIYRQALVAG
RKNVPGADGR PTQLQNEIDM GFPLTRPGWD YNTAEGRESL KIYRQALVAG
RKNVPGADGR PTQLQNEIDM GFPLTRPGWD YNTAEGRESL KIYRQALVAG
RKNVPGADGR PTQLQNEIDM GFPLTRPGWD YNTAEGRESL KIYRQALVAG
RKNVPGADGR PTQLQNEIDM GFPLTRPGWD YNTAEGRESL KIYRQALVAG
RKNVPGADGR PTQLQNEIDM GFPLTRPGWD YNTAEGRESL KIYRQALVAG
RKNVPGADGR PTQLQNEIDM GFPLTRPSWD YNTAEGRESL KIYRQALVAG
RKNVPGADGR PTQLQNEIDM GFPLTRPSWD YNTAEGRESL KIYRQALVAG
RKNVPGADGR PTQLQNETDM GFPLTPPGWD YNTAEGRESL KIYGQALVAG
RKNVPGADGR PTQLQNETDM GFPLTRPGWD YNTAEGRESL KIYRQALVAG
RKNVPGADGR PTQLQNEIDM GFPLTRPGWD YNTAEGRESL KIYRQALVAG
RKNVPGADGR PTQLQNEIDM GFPLTRPSWD YNTAEGRESL KIYHQALVAG
RKNVPGADG- PTQLQNEIDM GFPLTRPGWD YNTAEGRESL KIYRQALVAG
RKNVPGADGR PTQLQNEIDM GFPLTRPGWD YNTAEGRESL KIYCQALVAG
RKNVPGADGR PTQLQNEIDM GFPLTRPGWD YNTAEGRESL KIYRQALVAG

RKNVPGADGR PTQLQNEIDM GFPLTRPGWD YNTAEGRESL KIYRQALVAG
RKNVPGADGR PTQLQNEIDM RFPLTRPG-D YNTAEGRESL KIYRQALVAG
RKNVPGADGR PTQLQNEIDM GFPLTRPG-D YNTAEGRESL KIYRQALVAG
KKNVPGADGR PTQLQNKIDM GFPLTRPG-D YNTAEGRESL KIYRQALVAG
RKNVPGADGR PTQLQNEIDM GFPLTRPDWD YNTAEGRESL KIYRQALVVG
KKNVPGADGR PTQLQNEIDM GFPLTRP-WD YNTAEGRESL KIYRQALVVG
KKNVPGADGR PTQLQNEIDM GFPLTRP-WD YNTAEGRESL KIYRQALVVG
RKNVPGADGR PTQLQNEIDM GFP-THPGWD YNTAEGRESL KIYRQALVAG
RKNVPGADGR PTQLQNEIDM GFPLTRPSWD YNTAEGRESL KIYHQALVAG
RKNVPGANGR PTQLQNEIDM GFPLTRPSWD YNTAEGRESL KIYHQALVAG
RKNVPGANGR PTQLQNEIDM GFPLTRPSWD YNTAEGRESL KIYRQALVAG
RKNVPGANGR PTQLQNEIDM GFPLTRPSWD YNTAEGRESL KIYRQALVVG
RKNVPGANGR PTQLQNEIDM GFPLTRPSWD YNTAEGRESL KIYHQALVAG
RKNVPGADGR PTQLQNEIDM GFPLTRPGWD YNTAEGRESL KIYRQALVAG
RKNVPGADGQ PTQLQNEIDM GFPLTRPGWD YNTAEGRESL KIYRQALVAG
RKNVPGADGR PTQLQNEIDM GFPLTRPG-D YNTAEGRESL KIYRQALVAD
RKNVPGADGR PTQLQNEIDM GFPLTRPG-D YNTAEGRESL KIYRQALVAD
RKNVPGADGR PTQLQNEIDI GFPLTRPG-D YNTAEGRESL KIYRQALVAG
KKNVPGADGR PTQLQNEIDM RFPLTRPG-N YNTAEGRESL KIYRQALMAG
RKNVPGADGR PTQLQNEIDM GFPLTHPG-D YNTAEGRESL KIYRQALVAG
RKNIPGADGR PTQLQNEIDM GFPLTRHGWD YNTAKGRESL KIYRQALVAG
RKNIPGADGR PTQLQNEIDM GFPLTRPGWD HNTAVGRESL KIYRQALLAG
RKNIPGADGR PTQLQNEIDM GFPLTRPGWD YNTAVGRESL KIYRQALLAG
RKNVPGADGR PTQLQNEIDM GFPLTRPGWD YNTAEGRESL KIYHQALVAG
RKNVPGADGR PTQLQNEIDM GFPLTRPGWD YNTAEGRESL KIYHQALVAG
RKNVPGADGR PTQLQNEIDM GFPLTRPGWD YNTAEGRESL KIYHQALVAG
RKNVPGADG- PTQLQNEIDM GFPLTRPGWD YNTAEGRESL KIYHQALVAG
RKNVPGADGR PTQLQNEIDM GFPLTRPGWD YNTAEGRESL KIYHQALVAG
RKNVPGADGR PTQLQNEIDM GFPLTRPGWD YNTAEGRESL KIYHQALVAG
RRNVPGADG- PTQLQNEIDM GFPLTRPGWD YNTAEGRESL KIYHQALVAG
RKNVPGADGR PTQLQNEIDM GFPLTRPGWD YNTAEGRESL KIYHQALVAG
RKNVPGADGR PTHLQNEIDM GFPLTRPGWD YNTAEGRESV KIYCQSLVAG
RKNVPGADGR PTHLQNEIDM GFPLTRPGWD YNTAEGRESV KIYCQSLVAG
RKNVPGADGR PTHLQNEIDM GFPLTRPGWD YNTAEGRESV KIYCQSLVAG
RKNVPGADGQ PTHLQNEIDM GFPLTRPGWD YNTAEGRESV KIYCQALVAG
RKNVPGADGQ PTHLQNEIDM GFPLTRPGWD YNTAEGRESV KIYCQALVAG
RKNVPGADGQ PTHLQNEIDM GFPLTRPGWD YNTAEGRESV KIYCQALVAG
RKNVPGADGR PTQLQNEIDM GFPLTRPGWD YNTAEGRESL KIYRQALVAG
KKNVPGADGR PTQLQNEIDM GFPLTRPGWD YNTAEGRESL KIYHQALVAG
KKKVPGANGR PTQLQNEIDM EFPLTRPSWD YNTAEGRESL KIYRQALVAG
KKKVPGANGR PTQLQNEIDM EFPLTRPSWD YNTAEGRESL KIYRQALVAG
KKKVPGANGR PTQLQNEIDM EFPLTRPSWD YNTAEGRESL KIYRQALVAG
RKNVPGANRQ PTQLQHEIDM GFPPTCPVWD YNTAEGRESL KICLQALVAG
RKNVPGA-GQ PSQLQHEIDM GFPLTHPAWE YNTAEGRQSL KIYCQALVAD

RKNVPGTDGR PTQLPNEIDQ GFPLTRPLWD FNTPEGRERL TIYRQALVAG
RKNVPGTDGR PTQFTNKIDR GFPLTRHLWD FNTPEGRERQ TTYHQALVAG
RKNVLGEDGT PTALPNLVDE AFPLNRPNWD YNTAEGRGRL LVYRRTLVAG
RKNVLGVNGA PTQLENLINE AFPLNRPQWD HNTAEGRERL LVYRRTLVAG
RKNVLGDNGA PTQLENLINE AFPLNRPHWD YNTAAGRERL LVYRRTLVAG
RKNVLGANGQ PTQLPNEIDV GFPLVRPNWD FNAPEGRERL KMYRQALVVG
RKNVPGPGGF PTQLPNEIDE GFPLTRPDWD YETAPGRESL RIYRQALLAG
RKNVPGPGGL PTQLPNEIDE GFPLTRPDWD YETAPGRESL RIYRQALLAG
RKQVPGEDGR PTQLPNVIDE TFPLTRPNWD FATPAGREHL RLYRQLLLAG
RKAVRGDDGR PTQLPNEVDA AFPLERPDWD YTTQAGRNHL VHYRQLLLAG
RKAVRGEDGR PTQLPNDIND AFPLERPDWD YNTQRGRNHL VHYRQLLLAG
RKAVRGEDGR PTQLPNDIND AFPLERPDWD YNTQRGRNHL VHYRQLLLAG
RKAVRGDDGR PTQLPNEIEA AFPLERPDWD YTTLRGRNHL VLYRQLLLAG
RKNVPGDNGR PTTLPNLIDE RFPLNRPDWD FGNAEGRERL RVYRQTLMAG
RREVRNDQGV QVTDEREIEA QFPATRPDWD PNTGRGNDNL ERYRQILLRG

LRGASRRPTN LAKVREVMQG PNEPPSVFLE RLMEAFRRFT PFDPTSEAQK
LRGASRRPTN LAKVREVMQG PNEPPSVFLE RLMEAFRRFT PFDPTSEAQK
LRGASRRPTN LAKVREVMQG PNEPPSVFLE RLMEAFRRFT PFDPTSEAQK
LRGASRRPTN LAKVREVMQG PNEPPSVFLE RLMEAFRRFT PFDPTSEAQK
LRGASRRPTN LAKVREVMQG PNEPPSVFLE RLMEAFRRFT PFDPTSEAQK
LRGASRRPTN LAKVREVMQG PNEPPSVFLE RLMEAFRRFT PFDPTSEAQK
LRGASRRPTN LAKVREVMQG PNEPPSVFLE RLMEAFRRFT PFDPTSEAQK
LRGASRRPTN LAKVREVMQG PNEPPSVFLE RLMEAFRRFT PFDPTSEAQK
LRGASRRPTN LAKVREVMQG PNEPPSVFLE RLMEAFRRFT PFDPTSEAQK
LRGASRRPTN LAKVREVMQG PNEPPSVFLE RLMEAFRRFT PFDPTSEAQK
LRGASRRPTN LAKVREVMQG PNEPPSVFLE RLMEAFRRFT PFDPTSEAQK
LRGASRRPTN LAKVREVMQG PNEPPSVFLE RLMEAFRRFT PFDPTSEAQK
LRGASRRPTN LAKVREVMQG PNEPPSVFLE RLMEAFRRFT PFDPTSEAQK
LRGASRRPTN LAKVREVMQG PNEPPSVFLE RLMEAFRRFT PFDPTSEAQK
LRGASRRPTN LAKVREVMQG PNEPPSVFLE RLMEAFRRFT PFDPTSEAQK
LRGASRRPTN LAKVREVMQG PNEPPSVFLE RLMEAFRRFT PFDPTSEAQK
LRGASRRPTN LAKVREVMQG LNEPPSVFLE RLMEAFRRFT PFDPTSEAQK
LRGASRRPTN LAKVREVMQG PNEPPSVFLE RLMEAFRRFT PFDPTSEAQK
LRGASRRPTN LAKVREVMQG PNEPPSVFLE RLMEAFRRFT PFDPTSEAQK
LRGASRRPTN LAKVREVMQG PNEPPSVFLE RLMEAFRRFT PFDPTSEAQK
LRGASRRPTN LAKVREVMQG PNEPPSVFLE RLLEAFRRYT PFDPTSEAQK
LRGASRRPTN LAKVREVMQG PNEPPSVFLE RLLEAFRRYT PFDPTSEAQK
LRGASRRPTN LAKVREVMQG PNEPPSVFLE RLLEAFRRYT PFDPTSEAQK
LRGASRRPTN LAKVREVMQG PNEPPSVFLE RLLEAFRRYT PFDPTSEAQK
LRGASRRPTN LAKVREMMQG PNEPPSVFLE RLLEAFRRYT PFDPTSEAQK
LQGASRRPTN LAKVREVMQG PNEPPSVFLE RLMEAFRRFT PFDPTSEAQK
LRGASRR-TN LAKVREVMQG PNEPPSVFLE RLMEAFRRFT PFDPTSEAQK
LRGASRRPTN LAKVREVMQG PNKPPSVFLE RLMEAFRRFT PFDPTSEAQK

LRGASRRPTN LAKVREVMQG PNKPPSVFLE RLMEAFRRFT PFDPTSEAQK
LRGASRRPTN LAKVREVMQG PNEPPSVFLE RLMEAFRRFT PFDPTSEAQK
LRGASRRPTN LAKVREVMQG PNEPPSVFLE RLMEAFRRFT PFDPTSEAQK
LRGASRRPTN LAKVREVMQG PNEPPSVFLE RLMEAFRRFT PFDPTSEAQK
LRGASRRPTN LAKVREVMQG PNEPPSVFLE RLMEAFRRFT PFDPTSEAQK
LRGASRRPTN LAKVREVMQG PNEPPSVFLE RLMEAFRRFT PFDPTSEAQK
LRGASRRPTN LAKVREVMQG PNEPPSVFLE RLMEAFRRFT PFDPTSEAQK
LRGASRRPTN LAKVREVMQG PNEPPSVFLE RLMEAFRRFT PFDPTSEAQK
LRGASRRPTN LAKVREVMQG PNEPPSVFLE RLMEAFRRFT PFDPTSEAQK
LRGASRRPTN LAKVREVMQG PNEPPSVFLE RLMEAFRRFT PFDPTSEAQK
LRGASRRPTN LAKVREVMQG PNEPPSVFLE RLMEAFRRFT PFDPTSEAQK
LRGASRRPTN LAKVREVMQG ANEPPSVFLE RLMEAFRRFT PFDPTSEAQK
LRGASRRPTN LAKVREVMQG PNEPPSVFLE RLMEAFRRFT PFDPTSEAQK
LRGASRRPTN LAKVREVMQG PNEPPSVFLE RLMEAFRRFT PFDPTSEAQK
LRGASRRPTN LAKVREVMQG PNEPPSVFLE RLMEAFRRFT PFDPTSEAQK
LRGASRRPTN LAKVREVMQG PNEPPSVFLE RLMEAFRRFT PFDPTSEAQK
LRGASRRPTN LAKVREVMQG PNEPPSVFLE RLMEAFRRFT PFDPTSEAQK
LRGASRRPTN LAKVREVMQG PNEPPSVFLE RLMEAFRRFT PFDPTSEAQK
LRGASRRPTN LAKVREVMQG PNEPPSVFLE RLMEAFRRYT PFDPTSEAQK
LRGASRRPTN LAKVREMMQG PNEPPSVFLE RLMEAFRRYT PFDPTSEAQK
LRGASKRSTN LAKVREVMQG PKEPPSVFLE RLMEAFRRFT PFDPTSEAQK
LRGASKRSTN LAKVREVMQG PKEPPSVFLE RLMEAFRRFT PFDPTSEAQK
LRGASRRPTN LAKVREVKQG PNEPPSVFLE RLMEAFRRFT PFDPTSEAQK
LRGASRRPTN LAKVREVMQG PNEPPSVFLE RLMEAFRRFT PFDPTSEAQK
LRGASRWPTN LAKVREVMQG PNEPPSVFLE RLMEAFRRFT PFDPTSEAQK
LRGASRRPTN LAKVREMMQG PHEPPSVFLE RLMEAFRRFT PFDPTSEAQK
LRGASRRPTN LAKVREMMQG PHEPP--FLE RLMEAFRRFT PFDPTSEAQK
LRGASRRPTN LAKVREMMQG PHEPP--FLE RLMEAFRRFT PFDPTSEAQK
LRGASRRPTN LAKVREVMQG PNEPPSVFLE RLLEAFRRYT PFDPTSEAQK
LRGASRRPTN LAKVKKVMQG PNEPPSVFLE RLLEAFRRYT PFDPTSEAQK
LRGASKRPTN LAKVK-VMQG PNEPPSVFLE RLLEAFRRYT PFDPTSEAQK
LRGASRRPTN LAKVREVMQG PNEPPSVFLE RIMEAFRWFT PFDPTSEAQK
LWGASRRPTN LAKVREVMQG PNEPPSVFLE RIMEAFRWFT PFDPTSEAQK
LWGASRRPTN LAKVREVMQG PNEPPSVFLE RIMEAFRWFT PFDPTSEAQK
LRGASRRPTN LAKVREVMQG PNEPPSVFLE RLMEAFRRFT PFDPTSEAQK
LWGTSRWPTN LAKVREVMQG LNEPPLVFLE RLMEAFRRFT PFDPTSEAQK
LWGTSRWPTN LAKVREVMQG LNEPPLVFLE RLMEAFRRFT PFDPTSEAQK
LWGASRWPTN LAKVREVMQG LNEPPLVFLE RLMEAFRRFT PFDPTSEAQK
LWGTSRWPTN LAKVREVMQG LNEPPLVFLE RLMEAFRRFT PFDPTSEAQK
LRGTSRWPTN LAKVREVMQG LNEPPLVFLE RLMEAFRRFT PFDPTSEAQK
LRGASRRPTN LAKVREVMQG PNEPPSVFLE RLMEAFRRFT PFDPTSEAQK
LWGASRWPTN LAKVREVMQG PNEPPSVFLE RLMEAFRRFT PFDPTSEAQK
LRGASRRPTN LAKVREVMQG PNEPPSVFLE RLLEAFRRYT PFDPTSEAQK
LRGASRRPTN LAKVREVMQG PNEPPSVFLE RLLEAFRRYT PFDPTSEAQK

LRGASRRPTN LAKVREVMQG PNEPPSVFLE RLLEAFRRYT PFDPTSEAQK
LQGASRRPTN LAKVRKVMQG PNEPPSVFLE RLLEAFRRYT PFDPTSEAQK
LRGASRRPTN LAKVREVMQG PNEPPSVFLE RLLEAFRRYT PFDPTSEAQK
LRGSSRGPTN LAKVREVMQG PNEPPSVFLE RLMEAFRRFT PFDPTSEAQK
LRCASRWPTN LAKVREVMQG QNEPPLVFLE RLMEAFRWFT PFDPTSEAQK
LRCASRWPTN LAKVREVMQG QNEPPLVFLE RLMEAFRWFT PFDPTSEAQK
L-------TN LAKVKEVMQG PNELPSIFLE RLMEAFRRFT PFDPTSEAQK
--------TN LAKVKEVMQG PNELPSIFLE RLMEAFRRFT PFDPTSEAQK
--------TN LAKVKEVMQG PNELPSIFLE RLMEAFRRFT PFDPTSEAQK
L-------TN LAKVKEVMQG PNELPSIFLE RLMEAFRRFT PFDPTSEAQK
L-------TN LAKVKEVMQG PNELPSIFRW RLMEAFRRFT PFDPTSEAQK
--------TN LAKVKEVMQG PNELPSIFLE RLMEAFRRFT PFDPTSEAQK
--------TN LAKVKEVMQG PNELPSIFLE RLMEAFRRFT PFDPTSEAQK
L-------TN LAKVKEVMQG PNELPSIFLE RLMEAFRRFT PFDPTSEAQK
LRGASRRPTN LAKVREVMQG PNEPPSVFLE RLMEAFRRFT TFDPTSEAQK
LRGASRRPTN LAKVREVMQG PNEPPSVFLE RLMEAFRRFT TFDPTSEAQK
LRGASRRPTN LAKVREVMQG PNEPPSVFLE RLMEAFRRFT TFDPTSEAQK
LRGASRRPTN LAKVREVMQG PNEPPSVFLE RLMEAFRRFT TFDPTSEAQK
LRGASRRPTN LAKVREVMQG PNEPPSVFLE RLMEAFRRFT TFDPTSEAQK
LRGASRRPTN LAKVREVMQG PNEPPSVFLE RLMEAFRRFT TFDPTSEAQK
LRGASRRPT- LAKVREVMQG PNEPPSVFLE RLMEAFRRFT PFDPTSEAQK
LRGASRRPTN LTKVKKVIQG PNEPPSVFLE RLLKAFRRYT PFDPTSEAQK
LRGASRRPTN LA--REVMQG PNEPPSVFLE RLMEAFRRFT PFDPTSEAQK
LRGASRRPTN LA--REVMQG PSEPPSVFLE RLMEAFRRFT PFDPTSEAQK
LRGASRRPTN LA--REVMQG PNEPPSVFLE RLMEAFRRFT PFDPTSEAQK
LRGASRRPTN LAKVREVMQG PTESPSMFLE RLMKAFRWFT PFDPTSEIQK
LQDASRRPPR LARVREMMQG PTESPSMFLE RLMEAFWRFT PFDPTSEIQK
LRGAACRPTN LAKVREVTQG PTEAPSVFLE RIIDAF-RYT PFDPTSEGQG
IRGAARYPTN LAKVREVTQG PTEAPSVFLE RIIDAF-RYT SFDPTSEGQR
LRGAARRPTN LAKVREVLQG QTEPPSVFLE RLMEAYRRYT PFDPSSEGQK
LKGAARRPTN LAKVREVLQG PTEPPSVFLE RLMEAYRRYT PFDPSSEGQK
LKGAARRPTN LAKVREVLQG PAEPPSVFLE RLMEAYRRYT PFDPSSEGQQ
LHGAARRPTN LAKVREVTQG PQESPTVFLE RLMEAFRRFT PYDPTSEEHR
LKGAGKRPTN LAKVRTIIQG KEESPAAFME RLLEGFRMYT PFTPEAPEHK
LKGAGKRPTN LAKVRTITQG KDESPAAFME RLLEGFRMYT PFDPEAPEHK
LRGAARRPTN LAQVKQVVQG KEETPAAFLE RLKEAYRMYT PYDPEDPGQA
LQNAGRSPTN LAKVKGITQG PNESPSAFLE RLKEAYRRYT PYDPEDPGQE
LQNAGRSPTN LAKVKGITQG PNESPSAFLE RLKEAYRRYT PYDPEDPGQE
LQNAGRSPTN LAKVKGITQG PNESPSAFLE RLKEAYRRYT PYDPEDPGQE
LQNAGRSPTN LAKVKGITQG SNESPSAFLE RLKEAYRRYT PYDPEDPGQE
LRAAARRPTN LAKVKAIMQG DNESPAVFLE RLYDAYRQYT PLDPLAEENQ
LRAAARKPTN LSKITEVRQG ADESPTAYLE RLYQAYRTWS PIDPRAPENQ

ASVALAFIGQ SALDIRKKLQ RLEGLQEAEL RDLVKEAEKV YYKRETEEER

ASVALAFIGQ SALDIRKKLQ RLEGLQEAEL RDLVKEAEKV YYKRETEEER
ASVALAFIGQ SALDIRKKLQ RLEGLQEAEL RDLVKEAEKV YYKRETEEER
ASVALAFIGQ SALDIRKKLQ RLEGLQEAEL RDLVKEAEKV YYKRETEEER
ASVALAFIGQ SALDIRKKLQ RLEGLQEAEL RDLVKEAEKV YYKRETEEER
ASVALAFIGQ SALDIRKKLQ RLEGLQEAEL RDLVKEAEKV YYKRETEEER
ASVALAFIGQ SALDIRKKLQ RLEGLQEAEL RDLVKEAEKV YYKRETEEER
ASVALAFIGQ SALDIRKKLQ RLEGLQEAEL RDLVKEAEKV YYKRETEEER
ASVALAFIGQ SALDIRKKLQ RLEGLQEAEL RDLVKEAEKV YYKRETEEER
ASVALAFIGQ SALDIRKKLQ RLEGLQEAEL RDLVKEAEKV YYKRETEEER
ASVALAFIGQ SALDIRKKLQ RLEGLQEAEL RDLVKEAEKV YYKRETEEER
ASVALAFIGQ SALDIRKKLQ RLEGLQEAEL RDLVKEAEKV YYKRETEEER
ASVALAFIGQ SALDIRKKLQ RLEGLQEAEL RDLVKEAEKV YYKRETEEER
ASVALAFIGQ SALDIRKKLQ RLEGLQEAEL RDLVKEAEKV YYKRETEEER
ASVALAFIGQ SALDIRKKLQ RLEGLQEAEL HDLVKEAEKV YYKRETEEER
ASVALAFIGQ SALDIKKKLQ RLEGLQEAEL RDLVKEAEKV YYKRETEEER
ASVALAFIGQ SALDIRKKLQ RLEGLQEAEL RDLVKEAEKV YYKRETEEER
ASMALAFIGQ SALDIRKKLQ RLEGLQEAEL RDLVKEAKKV YYKRETEEER
ASMALAFIGQ SALDIRKKLQ RLEGLQEAEL HDLVKEAEKV YYKRETEEER
ASVALAFIGQ SALNIRKKLQ RLEGLQEAEL RDLVKEAEKV YYKRETEEER
ASVALAFIGQ SALDIRKKLQ RLEGLQEAEL RDLVKEAEKV YYKRETEEER
ASVALAFIGQ SALDIRKKLQ RLEGLQEAEL RDLVKEAEKV YYKRETEEER
ASVALAFIGQ SALDIRKKLQ RLEGLQEAEL RDLVKEAEKV YYKRETEEER
ASVALAFIGQ SALDIRKKLQ RLEGLQEAEL RDLVKEAEKV YYKRETEEER
ASVALAFIGQ SALDIRKKLQ RLEGLQEAEL RDLVKEAEKV YYKRETEEER
ASVALAFIGQ SALDIGKKLQ RLEGLQEAEL RDLVKEAEKV YYKRETEEER
ASVALAFIGQ SALDIRKKLQ RLEGLQEAEL RDLVKEAEKV YYKRETEEER
ASVALAFIGQ SALDIRKKLQ RLEGLQEAKL RDLVKEAEKV YYKRETEEER
ASVALAFIGQ SALDIRKKLQ RLEGLQEAEL RDLVREAEKV YYRRETEEEK
ASVALAFIGQ SALDIRKKLQ RLEGLQEAEL RDLVREAEKV YYRRETEEEK
ASVALAFIGQ SALDIRKKLQ RLEGLQEAEL RDLVREAEKV YYRRETEEEK
ASVALAFIGQ SALDIRKKLQ RLEGLQEAEL RDLVREAEKV YYRRETEEEK
ASVALAFIGQ SALDIRKKLQ RLEGLQEAEL RDLVREAEKV YYRRETEEEK
ASVALAFIGQ SALDIRKKLQ RLEGLQEAEL RDLVREAEKV YYRRETEEEK
ASVALAFIGQ SALDIRKKLQ RLEGLQEAEL RDLVREAEKV YYRRETEEEK
ASVALAFIGQ SALDIRKKLQ RLEGLQEAEL RDLVREAEKV YYRRETEEEK
ASVALAFIGQ SALDIRKKLQ RLEGLQEAEL RDLVREAEKV YYRRETEEEK
ASVALAFIGQ SALDIRKKLQ RLEGLQEAEL RDLVREAEKV YYRRETEEEK
ASVALAFIGQ SALDIRKKLQ RLEGLQEAEL RDLVKEAEKV YYRRETEEEK
ASVALAFIGQ SALDIRKKLQ RLEGLQEAEL RDLVREAEKV YYRRETEEEK
ASVALAFIGQ SALDIRKKLQ RLEGLQEAEL RDLVREAEKV YYRRETEEEK
ASVALAFIGQ SALDIRKKLQ RLEGLQEAEL RDLVREAEKV YYRRETEEEK
ASVALAFIGQ SALDIRKKLQ RLEGLQEAEL RDLVREAEKV YYRRETEEEK
ASVALAFIGQ SALDIRKKLQ RLEGLQEAEL RDLVREAEKV YYRRETEEEK
ASVALAFIGQ SALDIRKKLQ RLEGLQEAEL RDLVKEAEKV YYRRETEEEK

ASVALAFIGQ SALDIRKKLQ RLEGLQEAEL RDLVKEAEKV YYRRETEEEK

ASVALAFIGQ SALDIRKKLQ RLEGLQEAEL RDLVKEAEKV YYRRETEEEK

ASVALAFIGQ SALDIRKKLQ RLEGLQEAEL RDLVKEAEKV YYRSETEEEK

ASVALAFIGQ SALDIRKKLQ RLEGLQEAEL RDLVKEAEKV YYRRETEEEK

ASVALAFVGQ SALDIRKKLQ RLEGLQEAEL RDLVKEAEKV YYRRETEEEK

ASVALAFIGQ SALDIRKKLQ RLEGLQEAEL RDLVKEAEKV YYKRETEEER

ASVALAFIGQ SALDIRRKLQ RLEGLQEAEL RDLVKEAEKV YYKRETEEER

ASVALAFIGQ SALDIRKKLQ RLEGLQEAEL HDLVREAEKV YYRRETEEEK

ASVALAFIGQ SALDIRKKLQ RLEGLQEAEL RDLVREAEKV YYRRETEEEK

ASVALVFIGQ SALDIRKKLQ RLEGLQEAEL RDLVKEAEKV YYRRETEEEK

TSVALAFIGQ SALDIRKKLQ RLEGLQEAEL RDLVREAEKV YYRRETEEEK

ASVALAFIGQ SALDIRKKLQ RLEGLQEAEL RDLVKEAEKV YYKRETEEER

ASMALAFIGQ SALDIRKKLQ RLEGLQEAKL RDLVKEAEKV YYKRETDEER

ASMALAFIGQ SALDIRKKLQ RLEGLQEAEL RDLVKEAEKV YYKRETEEER

ASMALAFIGQ AALDIRKKCQ RLEGLQEAEL RDLVKEAEKV YYKRETEEER

ASMALAFIGQ AALDIRKKCQ RLEGLQEAEL RDLVKEAEKV YYKRETEEER

ASMALAFIGQ AALDIRKKCQ RLEGLQEAEL RDLVKEAEKV YYKRETEEER

ASVALAFIGQ SAQDIRRKLQ RLEGLQEAEL HDLVKEAEKV YYKRETEEER

ASVALDFIGQ SALDIRKKLR RLEGLQEA-L HDLVKEADKV YYRRETEEEK

ASVALDFIGQ SALDIRKKLR RLEGLQEA-L HDLVKEADKV YYRRETEEEK

ASVALDFIGQ SALDIRKKLR RLEGLQEA-L HDLVKEADKV YYRRETEEEK

ASVALDFIGQ SALDIRKKLR RLEGLQEA-L HDLVKEADKV YYRRETEEEK

ASVALDFIGQ SALDIRKKLR RLEGLQEA-L HDLVKEADKV YYRRETEEEK

ASVALDFIGQ SALDIRKKLR RLEGLQEA-L HDLVKEADKV YYRRETEEEK

ASMALAFIGQ AALDIRKKCQ RLEGLQEAEL RDLVKEAEKV YYRRETEEEK

ASMALAFIGQ SALDIKKKLQ RLEGLQEAEL RDLVKEAEKV YYKRETEEER

ASMALAFIGQ SALDIKKKLQ RLEGLQEAEL RDLVKEAEKV YYKRETEEER

ASVALAFIGQ SALDIKKKLQ RLKGLQEAEL RDLVKETEKV YYKRETEEER

ASVALAFIGQ SALDIRKKLQ RLEGLQEAEL HDIMKETEKV YYKRETEEER

ASVALAFIGQ SALDIRKKLQ RLEGLQEAEL HDLVKEAEKV YYKRETEEER

ASVALAFIGE LALDIRKKLQ RLEGLQEAEL CDLVKEAEKV YYRRETEEEK

ASGALAFIGQ LALDIRKKLQ RLEGLQEAEL CDLVKEAEKV YYRRETEEEK

ASGALAFIGQ LALDIRKKLQ RLEGLQEAEL RDLVKEAEKV YYKRETEEER

ASVALAFIGQ LAQNIRKKLQ RLEGLQEAEL RDLVKEAEKV YYKRETEEER

ASVALAFIGQ LAQNIRKKLQ RLEGLQEAEL RDLVKEAEKV YYKRETEEER

ASVALAFIGQ LAQNIRKKLQ RLEGLQEAEL RDLVKEAEKV YYKRETEEER

ASVALAFIGQ LAQNIRKKLQ RLEGLQEAEL RDLVKEAEKV YYKRETEEER

ASVALAFIGQ LAQNIRKKLQ RLEGLQEAEL RDLVKEAEKV YYKRETEEER

ASVALAFIGQ LAQNIRKKLQ RLEGLQEAEL RDLVKEAEKV YYKRETEEER

ASVALAFIGQ LAQNIRKKLQ RLEGLQEAEL RDLVKEAEKV YYKRETEEER

ASVALAFIGQ LAQNIRKKLQ RLEGLQEAEL RDLVKEAEKV YYKRETEEER

ASVALAFIGQ SALDIRRKLQ RLEGLQEAEL RDLVKEAEKV YYKRETEEER

ASVALAFIGQ SALDIRRKLQ RLEGLQEAEL RDLVKEAEKV YYKRETEEER

ASVALAFIGQ SALDIRRKLQ RLEGLQEAEL RDLVKEAEKV YYKRETEEER

ASVALAFIGQ SALDIRRKLQ RLEGLQEAEL RDLVKEAEKV YYKRETEEER
ASVALAFIGQ SALDIRRKLQ RLEGLQEAEL RDLVKEAEKV YYKRETEEER
ASVALAFIGQ SALDIRRKLQ RLEGLQEAEL RDLVKEAEKV YYKRETEEER
ASVALAFIGQ -ALDIRKKLQ RLEGLQEAEL RDLVKEAE-V YYKRETEEEE
ASMALAFIGQ STLDIKKKLQ RLKGLQEAEL RDLVKEAEKV YYKRET-KKK
ASMALAFVGQ AALDIRKKRQ RLEGLQEAEL RDLVKEAEKV YYRRETKEEK
ASMALAFVGQ AALDIRKKRQ RLEGLQEAEL RDLVKEAEKV YYRRETKEEK
ASMALAFVGQ AALDIRKK-Q RLEGLQEAEL RDLVKEAEKV YYRRETKEEK
GSVALAFIGQ SASDIRKKLQ RLEGLQEAES CDLVKEAEKV YYKRETEEEK
GSVALAFIG- SAPDIRKKLQ RLEGLQEAEL HDLVKEAEKV YYKRETEEEK
ASVTLAFIGQ AAPDIKRKLQ RLEGLQDLSL QDLIKEAEKV FYKRETEEEK
ASVALAFIGQ AAPDIKRKLQ RLEGLQDLSL QDLVKEAEKV FYKREMEEEK
AAVAMAFIGQ SAPDIKKKLQ RLEGLQDYTL QDLVKEAEKV YHKRETEEER
AAVAMSFIGQ SAPDIKKKLQ RLEGLQDHSL QDLIKEAEKV YHKRETEEEK
AAVAMAFIGQ SAPDIKKKLQ RLEGLQDYSL QDLVKEAEKV YHKRETEEER
ATIAMAFIDQ AAPDIKKKLQ SLDGLQGFSL QELVKEADKV YNKRETEEEK
ATVAMSFIDQ AASDIKGKLQ RLDGIQTYGL QELVREAEKV YNKRETPEEK
ATVAMSFIDQ AALDIKGKLQ RLDGIQTHGL QELVREAEKV YNKRETPEER
ASVILSFIYQ SSPDIRNKLQ RLEGLQGFTL SDLLKEAEKI YNKRETPEER
TNVSMSFIWQ SAPDIGRKLE RLEDLKNKTL GDLVREAEKI FNKRETPEER
TNVAMSFIWQ SAPDIGRKLE RLEDLKSKTL GDLVREAEKI FNKRETPEER
TNVSMSFIWQ SAPDIGRKLE RLEDLKSKTL GDLVREAEKI FNKRETPEER
TNVSMSFIWQ SAPDIGRKLE RLEDLKNKTL GDLVREAERI FNKRETPEER
SAVIMSFINQ AAPDIRKKLY KQEGLGEMSI RDLMKVAERV FNTRETPEER
AAIVIQFVSQ SAPDIRKKIQ KIDGFQGKSL SELVAIAQKV FDQREATHEL

EQRKEREREE REERRNKRQE KNLTK
EQRKEREREE REERRNKRQE KNLTK
EQRKEREREE REERRNKRQE KNLTK
EQRKEREREE REERRNKRQE KNLTK
EQRKEREREE REERRNKRQE KNLTK
EQRKEREREE REERRNKRQE KNLTK
EQRKEREREE REERRNKRQE KNLTK
EQRKEREREE REERRNKRQE KNLTK
EQRKEREREE REERRNKRQE KNLTK
EQRKEREREE REERRNKRQE KNLTK
EQRKEREREE REERRNKRQE KNLTK
EQRKEREREE REERRNKRQE KNLTK
EQRKKREREE REERRNKRQE KNLTK
EQKKEKEREE REERRNKRQE KNLTK
EQRKEREREE REERRNKRQE KNLTK
EQRKEREREK REERRNKRQE KNLTK
EQRKEREREE REERRNKRQE KNLTK
EQRKEREREE REERRNKRQE KNLTK

```
EQRKEREREE REERRNKRQE KNLTK
EQRKEREREE REERRNKRQE KNLTK
EQRKEREREE REERRNKRQE KNLTK
EQRKEREREE REERRNKRQE KNLTK
EQRKEREREE REERRNKRQE KNLTK
EQRKEREREE REERRNKRQE KNLTK
EQRKEREREE REERRNKRQE KNLTK
EQRKEREREE REERRNKRQE KNLTK
EQRKEREREE REERRNKRQE KNLTK
EQRKEREREE REERRNKRQE KNLTK
EQRKEKEREE REERRDRRQE KNLTK
EQRKEKEREE REERRDRRQE KNLTK
EQRKEKEREE REERRDRRQE KNLTK
EQRKEKEREE REERRDRRQE KNLTK
EQRKEKEREE REERRDRRQE KNLTK
EQRKEKEREE REERRDRRQE KNLTK
EQRKEKEREE REERRDRRQE KNLTK
EQRKEKEREE REERRDRRQE KNLTK
EQRKEKEREE REERRDRRQE KNLTK
EQRKEKEREE REERRDRRQE KNLTK
EQRKEKEREE REERRDRRQE KNLTK
EQRKEKEREE REERRDRRQE KNLTK
EQRKEKEREE REERHDRRQE KNLTK
EQRKEKEREK REKRRDRRQE KNLTK
EQRKEKEREE REERRDRRQE KNLTK
EQRKEKEREE REERRDRRQE KNLTK
EQRKEKEKEE REERCDRQQE KNLTK
EQRKEKEKEE REERCDRQQE KNLTK
EQRKEKEREE REERRDRRQE KNLTK
EQRKEKEREE REERRDRRQE KNLTK
EQRKEKEREE REERRDRRQE KNLTK
EQRKEKEREE REERRDRRQE KNLTK
EQRKEREREE REERRNKRQE KNLTK
EQKKEREKEK REERRDRRQE KNLTK
EQRKEKEREE REERRDRRQE KNLTK
EQRKEKEKEE REERRDRQQE KNLTK
EQRKEKEKEE REERCDRQQE KNLTK
EQRKEKEREE REERRDRRQE KNLTK
EQRKEREREE REERRNKRQE KNLTK
EQRKEREREE REERRNKRQE KNLTK
EQKKEREKEE REKRRNKRQE KNLTK
EQRKEREREE REERRNKRQE KNLTK
EQKKAREREE REERRNKRQE KNLTK
EQKKARERE- -EERRNKRQE K-LTK
```

```
EQRKEREREE REERRDRRQE KNLTK
EPRKEKEREE REERRDRRQE KNLTK
EPRKEKEREE REERRDRRQE KNLTK
EPRKEKEREE REERRDRRQE KNLTK
EPRKEKEREE REERRDRRQE KNLTK
EQSKEKEREE REERRDRRQE KNLTK
EPRKEKEREE REERRDRRQE KNLTK
EQRKEKEREE REERRDRWQE KNLTK
EQKKER-KKE KKKRRNKRQE KNLTK
EQKKER-KKE KKKRRNKRQE KNLTK
EQKKERKKEE REKRRNKRQE KNLTK
EQKKEREKEE REERRNKRQE KNLTK
KQRK-KKREK REKRRNKWQE KNLTK
EQRKEKEREE REERRDRRQE KNLTK
EQRKEKEREE REERRDRRQE KNLTK
EQRKEREREE REERRNKRQE KNLTK
EQRKEREREK REKRRDRR-E KNLTK
EQRKEREREK REKRRDRR-E KNLTK
EQRKEREREK REKRRDRR-E KNLTK
EQRKEREREK REKRRDRR-E KNLTK
EQRKEREREK REKRRDRR-E KNLTK
EQRKEREREK REKRRVPG-E KNLTK
EQRKEREREK REKRRDRR-E KNLTK
EQRKEREREE REERRNKRQE KNLTK
EQRKER---E KEERRDRRQE KNLTK
EQRKER---E KEERRDRRQE KNLTK
EQRKER---E KEERRDRRQE KNLTK
EQRKER---E KEERRDRRQE KNLTK
EQRKER---E KEERRDRRQE KNLTK
EQRKER---E KEERRDRRQE KNLTK
QRKK---REE REERR-IRQE KNLTK
GTKKEREREE REKRHNKQQE KNLTK
EQRKEKEREE REERRYRWQE KNLTK
EQRKEKEREE REERRYRWQE KNLTK
EQRKEKEREE REERRYRWQE KNLTK
EQRKEREREE REDRRNRRQK KNLNR
EQRKEREREE RKDRRNRRQE KNLTR
EQRREKEREE REEKHDRKRE KNLSK
EQRKEKEREE REEKCDRKWE KNLSK
QEREKKEVEE RENRRDRRQE RNLSK
QEREKKETEE RERRRDRRQE KNLTK
QEREKKEAEE KERRRDRPKK KNLTK
EERKQKEQEA REIRRDKRQE RNLSK
EARLAKEQEA REERRDRKRD KHLTK
```

| | | | |
|---|---|---|---|
| | EARLIKEQEE | REDRRDRKRD | KHLTK |
|  | EERLWQRQEE | R----DKKRH | KEMTK |
| | EERIRRETEE | KEERRDRRRH | REMSK |
| | EERIRRETEE | KEERRDRRRH | REMSK |
| | EDRIRRETEE | KEERRDRRRH | REMSK |
| | EERIKRETEE | KEERRDRRRQ | REMSK |
| | EDRIRKENQE | LQERINRRQQ | REMAK |
| | TRKMAKAQES | RAERGSKKTP | PGKGR |

Pol

76 567

| | | | | | |
|---|---|---|---|---|---|
| AOCR01031838.1 | EGIRPHVQRL | IQQGILVPVQ | SPWNTPLLPV | RKPGTNDYRP | VQDLREVNKR |
| AOCR01107158.1 | EGIRPHVQRL | IQQGILVPVQ | SPWNTPLLPV | RKPGTNDYRP | VQDLREVNKR |
| AOCR01098665.1 | EGIRPHVQRL | IQQGILVPVQ | SPWNTPLLPV | RKPGTNDYRP | VQDLREVNKR |
| PERV-C | EGIRPHVQRL | IQQGILVPVQ | SPWNTPLLPV | RKPGTNDYRP | VQDLREVNKR |
| CM000825.5_b | EGIRPHVQRL | IQQGILVPVQ | SPWNTPLLPV | RKPGTNDYRP | VQDLREVNKR |
| LUXS01031219.1 | EGIRPHVQRL | IQQGILVPVQ | SPWNTPLLPV | RKPGTNDYRP | VQDLREVNKR |
| LUXX01045907.1 | EGIRPHVQRL | IQQGILVPVQ | SPWNTPLLPV | RKPGTNDYRP | VQDLREVNKR |
| LT634572.1_d | EGIRPHVQRL | IQQGILVPVQ | SPWNTPLLPV | RKPGTNDYRP | VQDLREVNKR |
| LUXX01006697.1 | EGIRPHVQRL | IQQGILVPVQ | SPWNTPLLPV | RKPGTNDYRP | VQDLREVNKR |
| LUXQ01071671.1 | EGIRPHVQRL | IQQGILVPVQ | SPWNTPLLPV | RKPGTNDYRP | VQDLREVNKR |
| CM000816.5_b | EGIRPHVQRL | IQQGILVPVQ | SPWNTPLLPV | RKPGTNDYRP | VQDLREVNKR |
| PERV-A | EGIRPHVQRL | IQQGILVPVQ | SPWNTPLLPV | RKPGTNDYRP | VQDLREVNKR |
| LUXX01090830.1 | EGIRPHVQRL | IQQGILVPVQ | SPWNTPLLPV | RKPGTNDYRP | VQDLREVNKR |
| LUXU01038095.1 | EGIRPHVQRL | IQQGILVPVQ | SPWNTPLLPV | RKPGTNDYRP | VQDLREVNKR |
| LUXV01020898.1 | EGIWPHVQRL | IQQGILVPVQ | SPWNTPLLPV | RKPGTNDYRP | VQDLREVNKR |
| CM000812.5_b | EGIRPHVQRL | IQQGILVPVQ | SPWNTPLLPV | RKPGTNDYRP | VQDLREVNKR |
| CM000826.5_c | EGIRPHVQRL | IQQGILVPVQ | SPWNTPLLPV | RKPGTNDYRP | VQDLREVNKR |
| CM000823.5_b | EGIRPHVQRL | IQQGILVPVQ | SPWNTPLLPV | RKPGTNDYRP | VQDLREVNKR |
| AOCR01000130.1 | EGIQPHVQRL | IQQGILVPVQ | SPWNTPLLPV | RKPGTNDYRP | VQDLREVNKR |
| LUXW01074277.1 | EGIRPHVQRL | IQQGILVPVQ | SPWNTPLLPV | RKPGTNDYRP | VQDLREVNKR |
| LUXS01060324.1 | EGIQPHVQRL | IQQGILVPVQ | SPWNTPLLPV | RKPGTNDYRP | VQDLREVNKR |
| AOCR01079940.1 | EGIRPHVQRL | IQQGILVPVQ | SPWNTPLLPV | RKPGTNDYRP | VQDLREVNKR |
| AEMK02000197.1 | EGIWPHVQRL | IQQGILVPVQ | SPWNTPLLPV | RKPGTNDYRP | VQDLREVNKR |
| LUXR01004647.1 | EGIWPHVQRL | IQQGILVPVQ | SPWNTPLLPV | RKPGTNDYRP | VQDLREVNKR |
| LUXR01088996.1 | EGIWPHVQRL | IQQGILVPVQ | SPWNTPLLPV | RKPGTNDYRP | VQDLREVNKR |
| CM000815.5 | EGIWPHVQRL | IQQGILVPVQ | SPWNTPLLPV | RKPGTNDYRP | VQDLREVNKR |
| CM000822.5_b | EGIWPHVQRL | IQQGILVPVQ | SPWNTPLLPV | RKPGTNDYRP | VQDLREVNKR |
| CM000820.5 | EGIWPHVQRL | IQQGILVPVQ | SPWNTPLLPV | RKPGTNDYRP | VQDLREVNKR |
| PERV-B | EGIWPHVQRL | IQQGILVPVQ | SPWNTPLLPV | RKPGTNDYRP | VQDLREVNKR |
| CM000814.5_c | EGIWPHVQRL | IQQGILVPVQ | SPWNTPLLPV | RKPGTNDYRP | VQDLREVNKR |
| CM000827.5 | EGIWPHVQRL | IQQGILAPVQ | SPWNTPLLPV | RKPGTNDYRP | VQDLREVNKR |
| CM000826.5_a | EGIWPHVQRL | IQQDILVPVQ | SPWNTPLLPV | RKPGTNDYRP | VQDLREVNKR |
| CM000825.5_a | EGIWPHVQRL | IQQGILVPVQ | SPWNTPLLPV | RKPGTNDYRP | VQDLREVNKR |

| | | | | | |
|---|---|---|---|---|---|
| CM000819.5_a | EGIWPHVQRL | IQQGILVPVQ | SPWNTPLLPV | RKPGTNDYRP | VQDLREVNKR |
| CM000824.5_b | EGIWPHVQRL | IQQGILVPVQ | SPWNTPLLPV | RKPGTNDYRP | VQDLREVNKR |
| AEMK02000137.1 | EGIWPHVQRL | IQQGILVPVQ | SPWNTPLLPV | RKPGTNDYRP | VQDLREVNKR |
| CM000828.5_a | EGIRPHVQKL | IQQDILVPVQ | SPWNTPLLPV | RKPGTNDYRP | VQDLREVNKR |
| AEMK02000476.1 | EGIWPHVQRL | IQQGILVPVQ | SPWNTPLLLV | RKPGTNDYRP | VQDLREVNKR |
| CM000824.5_a | EGIRSHVQRL | IQQGILVPVQ | SPWNTPLLPV | RKPGTNDYRP | VQDLREVNKR |
| CM000819.5_b | EGIRPHVQRL | IQQGILVPVQ | SPWNTPLLPV | RKPGTNDYRP | VQDLREVN-R |
| CM000830.5_d | EGIRPHVQRL | IQQGILVPVQ | SPWNTPLLPV | RKPGTNDYRP | VQDLREVNKR |
| LT634572.1_b | EGIWPHVQRL | IQQGILVPVQ | SPWNTPLLPV | RKPGTNDYRP | VQDLREVNKR |
| CM000828.5_b | EGIWPHVQRL | I-QGILVPIQ | SPWNTPLLPV | RKPGTNDYRP | VQDLREVNKR |
| LT634572.1_a | EGIWPHVQRL | IQQGILVPVQ | SPWNTPLLPV | RKPGTNDYRP | VQDLREVNKR |
| LUXY01101100.1 | EGIWPHVQRL | IQQGILVPVQ | SPWNTPLLPV | RKPGTNDYRP | VQDLREVNKR |
| LUXQ01076544.1 | EGIWPHVQRL | IQQGILVPVQ | SPWNTPLLPV | RKPGTNDYRP | VQDLREVNKR |
| CM000828.5_c | EGIWPHVQRL | IQQGILVPVR | SP-NTPLLLV | RKPGTNDYRP | VQDLREVNKR |
| LIDP01000017.1 | EGIWPHVQRL | IQQGILVPVR | SP-NTPLLPV | RKPGTNDYRP | VQDLREVNKR |
| CM000814.5_b | EEIWPHVQRL | IQQGILVPVR | SPWNTPLLPV | RKPGTNDYRP | VQDLREVNKR |
| CM000813.5 | EGIRPHVQRL | IQQGILVPVQ | SPWNTPLLPV | RKPGTNDYRP | VQDLREVNKW |
| AJKK01232762.1 | EGIQPHVQRL | IQQGILVPVQ | SPWNTPLLPV | RKPGTNDYRP | VQDLREVNKR |
| CM000814.5_a | EGIRPHVQRL | IQQGILVPVQ | SP-NTPLLPV | RKPGTNDYRP | VQDLREVNKR |
| LUXY01013808.1 | EGIWPHVQRL | IQQGILVPVQ | SPWNTPLLPV | RKPGTNDYRP | VQDLREVNKR |
| AEMK02000393.1_a | EGIWPHVQRL | IQQGILVPVR | SPWNTPLLPV | RKPGTNDYRP | VQDLREVNKR |
| LUXW01069599.1 | EGIWPHVQRL | IQQGILVPVR | SPWNTPLLPV | RKPGTNDYRP | VQDLREVNKR |
| AEMK02000133.1 | EGIRPHVQRL | IQQGILVPVQ | SPWNTPLLPV | RKPGTNDYRP | VQDLREVNKW |
| CM000818.5_b | EGIWPHVQRL | IQQGILVPVQ | SPWNTPLLPV | RKPGTNDYCP | VQDLREVNKR |
| eJJRV_NW_004504375.1 | TGIWPHILRL | IQQGILVPIQ | SPWNTPLLPV | RKPGTNDYCP | VQDLREVNKR |
| eJJRV_NW_004504334.1 | TGIWPHIQRL | IQQGILVPVQ | SPWNTPLLPV | RKPGTNDYHP | VQDLKK-NKR |
| MDEV | EGIRPHIRRL | LDQGILVACQ | SPWNTPLLPV | RKPGTNDYRP | VQDLREVNKR |
| KoRV | EGIRPHIQRF | LDLGILVPCQ | SPWNTPLLPV | KKPGTNDYRP | VQDLREVNKR |
| GALV | EGIRPHIQKF | LDLGVLVPCR | SPWNTPLLPV | KKPGTNDYRP | VQDLREINKR |
| AEMK02000536.1 | EGIRPHVQRL | IQQGILVPVQ | SPWNTPLLPV | RKPGTNDYRP | VQDLREVNKR |
| ePCRV_KN680906.1 | EGIKPHIQRL | LDLGILIKCQ | SPWNTPLLPV | KKPGTGDYHP | VQDLREVNKR |
| ePCRV_KN676905.1 | SGIKPHIQRL | LDLGILIRYQ | SPWNTPLLPV | KKPGTGDYRP | VQDLRGISKR |
| ePCRV_KN676491.1 | TGIKPHIQRL | LNLGILIRCQ | SPWNTPLLPV | KKPGTGDYRP | VQDLREVNKR |
| M-CRV | LGIKPHIQRL | LDQGILVPCQ | SPWNTPLLPV | KKPGTNDYRP | VQDLREVNKR |
| R-MuLV | LGIKPHIQRL | LDQGILVPCQ | SPWNTPLLPV | KKPGTHDYRP | VQDLREVNKR |
| F-MuLV | LGIKPHIQRL | LDQGILVPCQ | SPWNTPLLPV | KKPGTNDYRP | VQDLREVNKR |
| M-MuLV | LGIKPHIQRL | LDQGILVPCQ | SPWNTPLLPV | KKPGTNDYRP | VQDLREVNKR |
| FeLV | QGIKPHIRRM | LDQGILKPCQ | SPWNTPLLPV | KKPGTEDYRP | VQDLREVNKR |
| RD114 | MGIQPHITRF | LELGVLRPCR | SPWNTPLLPV | KKPGTRDYRP | VQDLREVNKR |
| BaEV | MGIRQHIIKF | LELGVLRPCR | SPWNTPLLPV | KKPGTQDYRP | VQDLREINKR |
| OOEV | DGIRPHIQRL | LELGILVRCQ | STWNTPLLPV | KKPGTNDYQP | VQDLREVNKR |
| RfRV | KGIAPHINRL | LEAGILKPCH | SAWNTPLLPV | KKPGGKDYRP | VQDLREVNKR |
| REV | RSLRETIHKF | RAAGILRPVH | SPWNTPLLPV | RKSGTSEYRM | VQDLREVNKR |

VQDIHPTVPN PYNLLCALPP QRSWYTVLDL KDAFFCLRLH PTSQPLFAFE
VQDIHPTVPN PYNLLCALPP QRSWYTVLDL KDAFFCLRLH PTSQPLFAFE
VQDIHPTVPN PYNLLCALPP QRSWYTVLDL KDAFFCLRLH PTSQPLFAFE
VQDIHPTVPN PYNLLCALPP QRSWYTVLDL KDAFFCLRLH PTSQPLFAFE
VQDIHPTVPN PYNLLCALPP QRSWYTVLDL KDAFFCLRLH PTSQPLFAFE
VQDIHPTVPN PYNLLCALPP QRSWYTVLDL KDAFFCLRLH PTSQPLFAFE
VQDIHPTVPN PYNLLCALPP QRSWYTVLDL KDAFFCLRLH PTSQPLFAFE
VQDIHPTVPN PYNLLCALPP QRSWYTVLDL KDAFFCLRLH PTSQPLFAFE
VQDIHPTVPN PYNLLCALPP QRSWYTVLDL KDAFFCLRLH PTSQPLFAFE
VQDIHPTVPN PYNLLCALPP QRSWYTVLDL KDAFFCLRLH PTSQPLFAFE
VQDIHPTVPN PYNLLCALPP QRSWYTVLDL KDAFFCLRLH PTSQPLFAFE
VQDIHPTVPN PYNLLCALPP QRSWYTVLDL KDAFFCLRLH PTSQPLFAFE
VQDIHPTVPN PYNLLCALPP QWSWYTVLDL KDVFFCLRLH PTSQPLFAFE
VQDIHPTVPN PYNLLCALPP QWSWYTVLDL KDVFFCLRLH PTSQPLFAFE
VQDIHPTVPN PYNLLCALPP QWSWYTVLDL KDVFFCLRLH PTSQPLFAFE
VQDIHPTVPN PYNLLCALPP QRSWYTVLDL KDAFFCLRLH PTSQPLFAFE
VQDIHPTVPN PYNLLCALPP QRSWYTVLDL KDAFFCLRLH PTSQPLFAFE
VQDIHPTVPN PYNLLCALPP QRSWYTVLDL KDAFFCLRLH PTSQPLFAFE
VQDIHPTVPN PYNLLCALPP QRSWYTVLDL KDAFFCLRLH PTSQPLFAFD
VQDIHPTVPN PYNLLCALPP QRSWYTVLDL KDAFFCLRLH PTSQPLFAFE
VQDIHPTVPN PYNLLCALPP QRSWYTVLDL KDAFFCLRLH PTSQPLFAFD
VQDIHPTVPN PYNLLCALPP QRSWYTVLDL KDAFFCLRLH PTSQPLFAFE
VQDIHPTVPN PYNLLSALPP ERNWYTVLDL KDAFFCLRLH PTSQPLFAFE
VQDIHPTVPN PYNLLSALPP ERNWYTVLDL KDAFFCLRLH PTSQPLFAFE
VQDIHPTVPN PYNLLSALPP ERNWYTVLDL KDAFFCLRLH PTSQPLFAFE
VQDIHPTVPN PYNLLSALPP ERNWYTVLDL KDAFFCLRLH PTSQPLFAFE
VQDIHPTVPN PYNLLSALPP ERNWYTVLDL KDAFFCLRLH PTSQPLFAFE
VQDIHPTVPN PYNLLSALPP ERNWYTVLDL KDAFFCLRLH PTSQPLFAFE
VQDIHPTVPN PYNLLSALPP ERNWYTVLDL KDAFFCLRLH PTSQPLFAFE
VQDIHPTVPN PYNLLSALPP ERNWYTVLDL KDAFFCLRLH PTSQPLFAFE
VQDIHPTVPN PYNLLSALPP ERNWYTVLDL KDAFFCLRLH PTSQPLFAFE
VQDIHPTVPN PYNLLSALPP ERNWYTVLDL KDAFFCLRLH PTSQPLFAFK
VQDIHPTVPN PYNLLSALPP KRNWYTVLDL KDAFFCLRLH PTSQPLFAFE
VQDIHPTVPN PYNLLSALPP ERNWYTVLDL KDAFFCLRLH PTSQPLFAFE
VQDIHPTVPN PYNLLSALPP ERNWYTVLDL KDAFFCLRLH PTSQPLFAFE
VQDIHPTVPN PYNLLSALPP ERNWYTVLDL KDAFFCLRLH PTSQPLFAFE
VQDIHPTVPN PYNLLCALPP QRSWYTVLDL KDAFFCLRLH PTSQPLFAFE
VQDIHPTVPN PYNLLSALPP ERNWYTVLDL KDAFFCLRLH PTSQPLFAFK
VQDIHPTVPN PYNLLCALPP QRSWYTVLDL KDAFFCLRLH PTSQPLFAFE
VQDIHPTVPN PYNLLCALPP QRSWYTVLDL KDAFFCLRLH PTSQPLFAFE
VQDIHPTVPN PYNLLCALPP QRSWYKVLDL KDAFFCLRLH PTSQPLFAFE
VQDIHPTVPN PYNLLSALPP ERNWYTVLDL KDAFFCLRLH PTSQPLFAFE
VQDIHPTVPN PYNLLSALLP ERNWYTVLDL KDAFFCLRLH PTSQPLFAFE
VQDIHPTVPN PYNLLSALPP ERNWYTVLDL KDAFFCLRLH PTSQPLFAFE

VQDIHPTVPN PYNLLSALPP ERNWYTVLDL KDAFFCLRLH PTSQPLFAFE
VQDIHPTVPN PYNLLSALPP ERNWYTVLDL KDAFFCLRLH PTSQPLFAFE
VQDIYPTVPN PYNLLSALPP ERNWYTVLDL KDAFFCLRLH PSSQPLFAFE
V-DIYPTVPN PYNLLSALPP E-NWYTVLDL KDAFFCLRLH PTSQPLFAFE
VQDIHPMVPN PYNLLSALPP KRNWYTVLDL KNAFFCLRLH PTSQPLFAFE
VQDIHPTVPN PYNLLCALPP QRSWYTVLDL KDAFFCLRLH PTSQPLFAFE
VQDIHPTVPN PYNLLCALPP QRSWYTVLDL KDAFFCLRLH PTSQPLFAFD
VQDIHPTVPN PYNLLCALPP QRSWYTVLDL KDAFFCLRLH PTSQPLFAFK
VQDIHPTVPN PYNLLSALPP ERNWYTVLDL KDAFFCLRLH PTSQPLFAFE
VQDIHPTVPN SYNLLSALPP ERNWYTVLDL KDAFFCLRLH PTSQPLFAFE
VQDIHPTVPN SYNLLSALPP ERNWYTVLDL KDAFFCLRLH PTSQPLFAFE
VQDIHPTVPN PYNLLCALPP QWSWYTVLDL KDVFFCLRLH PTSQPLFAFE
VQDIHPTVPN PYNLLSALPP E--WYTVLDL KD-FFCLRLH PTSQPLFAFE
VQDIHPTVPN PCNLLSALPP ERIWYTVLDL KDAFFCLRLH LDSQPLFAFE
VQDIHPTVPN PYNLLSALPP KRS-YTVLDL KDAFFCLRLH LDSQSLFAFE
VLDIHPTVPN PYNLLSSLPP ERTWYTVLDL KDAFFCLRLH PKSQLLFAFE
VQDIHPTVPN PYNLLSSLPP SHTWYSVLDL KDAFFCLKPH PNSQPLFAFE
VQDIHPTVPN PYNLLSSLPP SYTWYSVLDL KDAFFCLRLH PNSQPLFAFE
VQDIHPT-PN -YNLLSALLP ERNWYTV-DL KDAFFCLRLP P-TQPLLF-E
VLDIHPTVPN PYNLLSSLPP SHVWYTVLDF KDAFFCLRLH PTSQPMFAFE
VQDIHPTVPN PYNLLSSLPP SHVWYTVLDL KDAFFCLRLH PTSQPMFAFE
VQDIHPTVPN PYNLLSSLPP SHVWYTVLDL KDAFFCLRLH PTSQPMFAFE
VEDIHPTVPN PYNLLSGLPP SHQWYTVLDL KDAFFCLRLH PTSQPLFAFE
VEDIHPTVPN PYNLLSGLPP SHQWYTVLDL KDAFFCLRLH PTSQSLFAFE
VEDIHPTVPN PYNLLSGLPP SHQWYTVLDL KDAFFCLRLH PTSQSLFAFE
VEDIHPTVPN PYNLLSGLPP SHQWYTVLDL KDAFFCLRLH PTSQPLFAFE
VEDIHPTVPN PYNLLSTLPP SHPWYTVLDL KDAFFCLRLH SESQLLFAFE
TMDIHPTVPN PYNLLSTLSP DRTWYTVLDL KDAFFCLPLA PQSQELFAFE
TVDIHPTVPN PYNLLSTLKP DYSWYTVLDL KDAFFCLPLA PQSQELFAFE
VADLHPTVPN PYNLLSTLPP QHTWYTVLDL KDAFFCLRLS PLSQPYFAFE
VEDIHPTVPN PYTLLSHLPP SHVWYTTLDL KDAFFSIALA PSSQHIFAFE
VETIHPTVPN PYTLLSLLPP DRIWYSVLDL KDAFFCIPLA PESQLIFAFE

WRDPGTGRTG QLTWTRLPQG FKNSPTIFDE ALHRDLANFR IQHPQVTLLQ
WRDPGTGRTG QLTWTRLPQG FKNSPTIFDE ALHRDLANFR IQHPQVTLLQ
WRDPGTGRTG QLTWTRLPQG FKNSPTIFDE ALHRDLANFR IQHPQVTLLQ
WRDPDTGRTG QLTWTRLPQG FKNSPTIFDE ALHRDLANFR IQHPQVTLLQ
WRDPGTGRTG QLTWTRLPQG FKNSPTIFDE ALHRDLANFR IQHPQVTLLQ
WRDPGAGRTG QLTWTRLPQG FKNSPTIFDE ALHRDLANFR IQHPQVTLLQ
WRDPGAGRTG QLTWTRLPQG FKNSPTIFDE ALHRDLANFR IQHPQVTLLQ
WRDPGAGRTG QLTWTRLPQG FKNSPTIFDE ALHRDLANFR IQHPQVTLLQ
WRDPGAGRTG QLTWTRLPQG FKNSPTIFDE ALHRDLANFR IQHPQVTLLQ
WRDPGAGRTG QLTWTRLPQG FKNSPTIFDE ALHRDLANFR IQHPQVTLLQ
WRDPGAGRTG QLTWTRLPQG FKNSPTIFDE ALHRDLANFR IQHPQVTLLQ

WRDPGAGRTG QLTWTRLPQG FKNSPTIFDE ALHRDLANFR IQHPQVTLLQ
WRDPGAGRTG QLTWTRLPQG FKNSPTIFDE ALHRDLANFR IQHPQVTLLQ
WRDPGAGRTG QLTWTRLPQG FKNSPTIFDE ALHRDLANFR IQHPQVTLLQ
WRDPGTGRTG QLTWTRLPQG FKNSPTIFDE ALHRDLANFR IQHPQVTLLQ
WRDPGAGRTG QLTWTRLPQG FKNSPTIFDE ALHRDLANFR IQHPQVTLLQ
WRDPGAGRTG QLTWTRLPQG FKNSPTIFDE ALHRDLANFR IQHPQVTLLQ
WRDPGAGRTG QLTWTRLPQG FKNSPTIFDE ALHRDLANFR IQHPQVTLLQ
WKDPGTGRTG QLTWTRLPQR FKNAPTIFDE ALHRDLANFR IQHPQVTLLQ
WRDPGAGRTG QLTWTRLPQG FKNSPTIFDE ALHRDLANFR IQHPQVTLLQ
WKDPGTGRTG QLTWTRLPQR FKNAPTIFDE ALHRDLANFR IQHPQVTLLQ
WRDPGTGRTG QLTWTRLPQG FKNSPTIFDE ALHRDLANFR IQHPQVTLLQ
WRDPGTGRTG QLTWTRLPQG FKNSPTIFDE ALHRDLANFR IQHPQVTLLQ
WRDPGTGRTG QLTWTRLPQG FKNSPTIFDE ALHRDLANFR IQHPQVTLLQ
WRDPGTGRTG QLTWTRLPQG FKNSPTIFDE ALHRDLANFR IQHPQVTLLQ
WRDPGTGRTG QLTWTRLPQG FKNSPTIFDE ALHRDLANFR IQHPQVTLLQ
WRDPGTGRTG QLTWTRLPQG FKNSPTIFDE ALHRDLANFR IQYPQVTLLQ
WRDPGTGRTG QLTWTRLPQG FKNSPTIFDE ALHRDLANFR IQHPQVTLLQ
WRDPGTGRTG QLTWTRLPQG FKNSPTIFDE ALHRDLANFR IQHPQVTLLQ
WRDPGTGRTG QLTWTRLPQG FKNSPTIFDE ALHRDLANFR IQHPQVTLLQ
WRDPGTGRTG QLTWTRLPQG FKNSPTIFDE ALHRDLANFR IQHPQVTLLQ
WRDPGTGRTG QLTWTRLPQG FKNSPTIFDE ALHRDLANFR IQHPQVTLLQ
WRDPGTGRTG QLTWTRLPQG FKNSPTIFDE ALHRDLANFR IQHPQVTLLQ
WRDPGTGRTG QLTWTRLPQG FKNSPTIFDE ALHRDLANFR IQHPQVTLLQ
WRDPGTGRTG QLTWTRLPQG FKNSPTIFDE ALHRDLANFR IQHPQVTLLQ
WRDPGTGRTG QLTWTRLPQG FKNSPTIFDE ALHRDLANFR IQHPQVTLLQ
WRDPGAGRTG QLTWTRLPQG FKNSPTIFDE ALHRDLANFR IQHPQVTLLQ
WRDPGTGRTG QLTWTRLPQG FKNSPTIFDE ALHRDLANFR IQHPQVTLLQ
WRDPGAGRTG QLTWTRLPQG FKNSPTIFDE ALHRDLANFR IQHPQVTLLQ
WRDPGAGRTR QLTWTRLPQG FKNSPTIFDE ALHRDLANFR IQHPQVTLLQ
WRDPGAGRTG QLTWTRLPQG FKNSPTIFDE ALHRDLANFR IQHAQVTLLQ
WRDPGTGRTR QLTWTRLPQG FKNSPTIFDE ALHRDLANFR IQHPQVTLLQ
WRDPGTGRTG QLTWTRLPQG FKNSPTIFDE ALHRDLANFR IQHPQVTLLQ
WRDPGTGRTR QLTWTRLPQG FKNSPTIFDE ALHRDLANFR IQHPQVTLLQ
WRDPGTGRTG QLTWTRLPQG FKNSPTIFDE ALHRDLANFR IQHPQVTLLQ
WRDPGTGRTG QLTWTRLPQG FKNSPTIFDE ALHRDLANFR IQHPQVTLLQ
WRDPGTGRTG QLTWTRLPQG FKNSPTIFDE ALHRDLANFR IQHPQVTLLQ
WRDPGTGRTG QLTWTRLPQG FKNSPTIFDE ALHRDLANFR IQHPQVTLLQ
WRDPGTGRTG QLTWTRLPQG FKNSPTIFDE ALHRDLANFR IQHPQVTLLQ
WRDPGAGRTG QLTWTRLPQG FKNSPTIFDE ALHRDLANFR IQHPQVTLLQ
WKDPGTGRTG QLTWTRLPQR FKNAPTIFDE ALHRDLANFR IQHPQVTLLQ
WRDPGAGRTG QLTWTRLPQG FKNSPTIFDE ALHRDLANFR IQHPQVTLLQ
WRDPGTGRTG QLTWTRLPQG FKNSPTIFDE ALHRDLANFR IQHPQVTLLQ
WRDPGTGRTG QLTWTRLPQG FKNSPTIFDE ALHRDLANFR IQHPQVTLLQ
WRDPGTGRTG QLTWTRLPQG FKNSPTIFDE ALHRDLANFR IQHPQVTLLQ

```
WRDPGTGRTG QLTWTRLPQG FKNSPTIFDE ALHRDLANFR IQHPQVTLLQ
WRDPGTGRTG QLTWT-LPQG FKNSPTIFDK ALHKDLANFR IQHPQVTLLQ
WRDPNTERTG QLTWTRLPQG FKNSPTIFDE ALHRDLANFR VQHPQVTLLQ
WRDPDTGRTG -VNWTHLP-G FKNSPTIFDK ALHRDLANFR VQHPHLTLLQ
WRDPEGGQTG QLTWTRLPQG FKNSPTLFDE ALHRDLAPFR AQNPQLTLLQ
WRDPEKGNTG QLTWTRLPQG FKNSPTLFDE ALHRDLASFR ALNPQVVMLQ
WKDPEKGNTG QLTWTRLPQG FKNSPTLFDE ALHRDLAPFR ALNPQVVLLQ
-RDPGT-EEG QLTWTRLPQG -KNSPTIFDE ALHRD-ANF- IQHPT---PQ
W-DPQDGITG QLTWTRLPQG FKNSPTIFDE ALHQDLMPFR VSHPQVTLLK
WKDPQDGTTG QLTWTRLPQG FKNSPTIFDE ALHQDLTPFH ASHPQVTLLQ
WKDPQDGTTG QLMWTRLPQG FKNSPTIFDE ALHQDLTPFR ASHPQVTLLQ
WRDPEMGISG QLTWTRLPQG FKNSPTLFDE ALHRDLADFR IQHPDLILLQ
WRDPEMGISG QLTWTRLPQG FKNSPTLFDE ALHRDLADFR IQHPDLILLQ
WRDPEMGISG QLTWTRLPQG FKNSPTLFDE ALHRDLADFR IQHPDLILLQ
WRDPEMGISG QLTWTRLPQG FKNSPTLFDE ALHRDLADFR IQHPDLILLQ
WRDPEIGLSG QLTWTRLPQG FKNSPTLFDE ALHSDLADFR VRYPALVLLQ
WRDPERGISG QLTWTRLPQG FKNSPTLFDE ALHRDLTDFR TQHPEVTLLQ
WKDPERGISG QLTWTRLPQG FKNSPTLFDE ALHRDLTDFR TQHPEVTLLQ
WKDPASGISG QLTWTCLPQE FKNSPTIFDE ALHQDLALYR ESNPQVTLLQ
WNNGNTGTPG QLTWTRLPQG FKNSPTLFNE ALNQDLDSFR QSHNSVTLLQ
WADAEEGESG QLTWTRLPQG FKNSPTLFDV ALNRDLQGFR LDHPSVSLLQ
```

```
YVDDLLLAGA TKQDCLEGTK ALLLELSDLG YRASAKKAQI CRREVTYLGY
YVDDLLLAGA TKQDCLEGTK ALLLELSDLG YRASAKKAQI CRREVTYLGY
YVDDLLLAGA TKQDCLEGTK ALLLELSDLG YRASAKKAQI CRREVTYLGY
YVDDLLLAGA TKQDCLEGTK ALLLELSDLG YRASAKKAQI CRREVTYLGY
YVDDLLLAGA TKQDCLEGTK ALLLELSDLG YRASAKKAQI CRREVTYLGY
YVDDLLLAGA TKQDCLEGTK ALLLELSDLG YRASAKKAQI CRREVTYLGY
YVDDLLLAGA TKQDCLEGTK ALLLELSDLG YRASAKKAQI CRREVTYLGY
YVDDLLLAGA TKQDCLEGTK ALLLELSDLG YRASAKKAQI CRREVTYLGY
YVDDLLLAGA TKQDCLEGTK ALLLELSDLG YRASAKKAQI CRREVTYLGY
YVDDLLLAGA TKQDCLEGTK ALLLELSDLG YRASAKKAQI CRREVTYLGY
YVDDLLLAGA TKQDCLEGTK ALLLELSDLG YRASAKKAQI CRREVTYLGY
YVDDLLLAGA TKQDCLEGTK ALLLELSDLG YRASAKKAQI CRREVTYLGY
YVDDLLLAGA TKQDCLEGTK ALLLELSDLG YRASAKKAQI CRREVTYLGY
YVDDLLLAGA TKQDCLEGTK ALLLELSDLG YRASAKKAQI CRREVTYLGY
YVDDLLLAGA TKQDCLEGTK ALLLELSDLG YRASAKKAQI CRREVTYLGY
YVDDLLLAGA TKQDCLEGTK ALLLELSDLG YRASAKKAQI CRREVTYLGY
YVDDLLLAGA TKQDCLEGTK ALLLELSDLG YRASAKKAQI CRREVTYLGY
YVDDLLLAGA TKQDCLEGTK ALLLELSDLG YRASAKKAQI CRREVTYLGY
YVDDLLLAGA TKQDCLEGTK TLLLELSDIG YRASTKKAQI CRREVTYLGY
YVDDLLLAGA TKQDCLEGTK ALLLELSDLG YRASAKKAQI CRREVTYLGY
YVDDLLLAGA TKQDCLEGTK ALLLELSDLG YRASAKKAQI CRREVTYLGY
YVDDLLLAGA TKQDCLEGTK ALLLELSDLG YRASAKKAQI CRREVTYLGY
```

```
YVDDLLLAGA TKQDCLEGTK ALLLELSDLG YRASAKKAQI CRREVTYLGY
YVDDLLLAGA TKQDCLEGTK ALLLELSDLG YRASAKKAQI CRREVTYLGY
YVDDLLLAGA TKQDCLEGTK ALLLELSDLG YRASAKKAQI CRREVTYLGY
YVDDLLLAGA TKQDCLEGTK ALLLELSDLG YRASAKKAQI CRREVTYLGY
YVDDLLLAGA TKQDCLEGTK ALLLELSDLG YRASAKKAQI CRREVTYLGY
YVDDLLLAGA TKQDCLEGTK ALLLELSDLG YRASAKKAQI CRREVTYLGY
YVDDLLLAGA TKQDCLEGTK ALLLELSDLG YRASAKKAQI CRREVTYLGY
YVDDLLLAGA TKQDCLEGTK ALLLKLSDLG YRASAKKAQI CRREVTYLGY
YVDDLLLAGA TKQDCLEGTK ALLLELSDLG YRASAKKAQI CRREVTYLGY
YVDDLLLAGA TKQDCLEGTK ALLLELSDLG YRASAKKAQI CRREVTYLGY
YVDDLLLAGA TKQDCLEGTK ALLLELSDLG YRASAKKAQI CRREVTYLGY
YVDDLLLAGA TKQDCLEGTK ALLLELSDLG YRASAKKAQI CRREVTYLGY
YVDDLLLAGA TKQDCLEGTK ALLLELSDLG YRASAKKAQI CRREVTYLGY
YVDDLLLAGA TKQDCLEGTK ALLLELSDLG YRASAKKAQI CRREVTYLGY
YVDDLLLAGA TKQDCLEGTK ALLLELSDLG YRASAKKAQI CRREVTYLGY
YVDDLLLAGA TKQDCLEGTK ALLLELSDLG YRASAKKAQI CRREVTYLGY
YVDDLLLAGA TKQDCLEGTK ALLLELSDLG YRASAKKAQI CRREVTYLGY
YVDDLLLAGA TKQDCLEGTK ALLLELSDLG YRASAKKAQI CRREVTYLGY
YVDDLLLVGA TKQDCLEGTK -LLLELSDLG YRASAKKAQI CRREVTYLGY
YVDDLLLAGA TKQDCLEGTK ALLLELSDLG YRASAKKAQI CRKEVTYLGY
YVDDLLLAGA TKQDCLEGTK ALLLELSDLG YRASAKKAQI CRREVTYLGY
YVDDLLLAGA TKQDCLEGTK ALLLELSDLG YRASAKKAQI CRKEVTYLGY
YVDDLLLAGA TKQDCLEGTK ALLLELSDLG YRASAKKAQI CRREVTYLGY
YVDDLLLAGA TKQDCLEGTK ALLLELSDLG YRASAKKAQI CRREVTYLGY
YVDDLLLAGA TKQDCLEGTK ALLLKLSDLG YRASAKKAQI CRREVTYLGY
YVDDLLLAGA TKQDCLEGTK ALLLKLSDLG YRASAKKAQI CRREVTYLGY
YVDDLLLAGA TKQDCLEGTK ALLLELSDLG YRASAKKAQI CRREVTYLGY
YVDDLLLAGD TKQDCLKGTK ALLLELSDLG YRASAKKAQI CRREVTYLGY
YVDDLLLAGA TKQDCLEGTK TLLLELSDIG YRASTKKAQI CRREVTYLGY
YVDDLLLAGA TKQDCLEGTK ALLLELSDLG YRASAKKAQI CRREVTYLGY
YVDDLLLAGA TKQDCLEGMK ALLLELSDLG YRASAKKAQI CRREVTYLGY
YVDDLLLAGA TKQDCLEGTK ALLLELSYLG YRASAKKAQI CRREVTYLG-
YVDDLLLAGA TKQDCLEGTK ALLLELSYLG YRASAKKAQI CRREVTYLG-
YVDDLLLAGA TKQDCLECTK ALLLELSDLG YRASAKKAQI CRREVTYLGY
FEDDLLLAGA TKQDCLEGTK ALLLELSDLG YRASAKKAQI CRREVTYLGY
YVDDLLLAGA TQQDCLKGTK ALLLELTDLG YRASAKKAQI CKREVTYLGY
YVDDLLLAGA TQQNCLKGMK AQLLELTDLG Y-ASAKKAQI CKREVTYLGY
YVDDLLIAAA SKELCQQGTE RLLTELGNLG YRVSAKKAQI CQTEVIYLGY
YVDDLLVAAP TYRDCKEGTR RLLQELSKLG YRVSAKKAQL CREEVTYLGY
YVDDLLVAAP TYEDCKKGTQ KLLQELSKLG YRVSAKKAQL CQREVTYLGY
YVE---LLGA TK-DC-EGTK ALLLELSP-S YRA-AKKAQI CRREVTYLGY
YVDDLLIAGA TETECREATM ALLTELSRLG Y-ASAKKALL CQREVTYLGY
YVDDLLIARA SEAECQEATM ALLTELSPLG YRASAKKAQL CRQEVTYLGY
YVDDLLIAGA SEAECREATM ALLTELSRLG YRASAKKAQL CRQEVTYLGY
```

YVDDLLLAAT SELDCQQGTR ALLQTLGDLG YRASAKKAQI CQKQVKYLGY
YVDDLLLAAT SELDCQQGTR ALLQTLGDLG YRASAKKAQI CQKQVKYLGY
YVDDLLLAAT SELDCQQGTR ALLQTLGDLG YRASAKKAQI CQKQVKYLGY
YVDDLLLAAT SELDCQQGTR ALLQTLGNLG YRASAKKAQI CQKQVKYLGY
YVDDLLLAAA TRTECLEGTK ALLETLGNKG YRASAKKAQI CLQEVTYLGY
YVDDLLLAAP TEEACTRGTK HLLRELGDKG YRASAKKAQI CQTKVTYLGY
YVDDLLLAAP TKKACTQGTR HLLQELGEKG YRASAKKAQI CQTKVTYLGY
YVDDILLAAE TQEDCIKGTE KLLTELRTLG YRASAKKAQI CQQQVSYLGY
YVDDLLLAAP SEAECRQATG DLLQELGQLG YRASAKKAQI CRQTVTYLGY
YVDDLLIAAD TQAACLSATR DLLMTLAELG YRVSGKKAQL CQEEVTYLGF

SLRDGQRWLT EARKKTVVQI PAPTTAKQMR EFLGTAGFCR LWIPGFATLA
SLRDGQRWLT EARKKTVVQI PAPTTAKQMR EFLGTAGFCR LWIPGFATLA
SLRDGQRWLT EARKKTVVQI PAPTTAKQMR EFLGTAGFCR LWIPGFATLA
SLRDGQRWLT EARKKTVVQI PAPTTAKQMR EFLGTAGFCR LWIPGFATLA
SLRDGQRWLT EARKKTVVQI PAPTTAKQMR EFLGTAGFCR LWIPGFATLA
SLRGGQRWLT EARKRTVVQI PAPTTAKQVR EFLGTAGFCR LWIPGFATLA
SLRGGQRWLT EARKRTVVQI PAPTTAKQVR EFLGTAGFCR LWIPGFATLA
SLRGGQRWLT EARKRTVVQI PAPTTAKQVR EFLGTAGFCR LWIPGFATLA
SLRGGQRWLT EARKRTVVQI PAPTTAKQVR EFLGTAGFCR LWIPGFATLA
SLRGGQRWLT EARKRTVVQI PAPTTAKQVR EFLGTAGFCR LWIPGFATLA
SLRGGQRWLT EARKRTVVQI PAPTTAKQVR EFLGTAGFCR LWIPGFATLA
SLRGGQRWLT EARKRTVVQI PAPTTAKQVR EFLGTAGFCR LWIPGFATLA
SLRGGQRWLT EARKRTVVQI PAPTTAKQVR EFLGTAGFCR LWIPGFATLA
SLRGGQRWLT EARKRTVVQI PAPTTAKQVR EFLGTAGFCR LWIPGFATLA
SLRGGQRWLT EARKKTVVQI PAPTTAKQVR EFLGTAGFCR LWIPGFATLA
SLRGGQRWLT EARKRTVVQI PAPTTAKQVR EFLGTAGFCR LWIPGFATLA
SLRGGQRWLT EARKRTVVQI PAPTTAKQVR EFLGTAGFCR LWIPGFATLA
SLRGGQRWLT EARKRTVVQI PAPTTAKQVR EFLGTAGFCR LWIPGFATLA
SLRDGQRWLT EARKKTVVQI PAPTTAKQMR EFLGTAGFCR LWIPGFATLA
SLRGGQRWLT EARKRTVVQI PAPTTAKQVR EFLGTAGFCR LWIPGFATLA
SLRGGQRWLT EARKRTVVQI PAPTTAKQVR EFLGTAGFCR LWIPGFATLA
SLRDGQRWLT EARKKTVVQI PAPTTAKQMR EFLGTAGFCR LWIPGFATLA
SLRGGQRWLT EARKKTVVQI PAPTTAKQVR EFLGTAGFCR LWIPGFATLA
SLRGGQRWLT EARKKTVVQI PAPTTAKQVR EFLGTAGFCR LWIPGFATLA
SLRGGQRWLT EARKKTVVQI PAPTTAKQVR EFLGTAGFCR LWIPGFATLA
SLRGGQRWLT EARKKTVVQI PAPTTAKQVR EFLGTAGFCR LWIPGFATLA
SLRGGQRWLT EARKKTVVQI PAPTTAKQVR EFLGTAGFCR LWIPGFATLA
SLRGGQRWLT EARKKTVVQI PAPTTAKQVR EFLGTAGFCR LWIPGFATLA
SLRGGQRWLT EARKKTVVQI PAPTTAKQVR EFLGTAGFCR LWIPGFATLA
SLRGGQRWLT EARKKTVVQI PAPTTAKQVR EFLGTAGFCR LWIPGFATLA
SLRGGQRWLT EARKKTVVQI PAPTTAKQVR EFLGTAGFCR LWIPGFATLA
SLRGGQRWLT EARKKTVVQI PAPTTAKQVR EFLGTAGFCR LWIPGFATLA
SLRGGQRWLT EARKKTVVQI PAPTTAKQVR EFLGTVGFCR LWIPGFATLA

SLRGGQRWLT EAWKKTVVQI PAPTTAKQVR EFLGTAGFCR LWIPGFATLA

SLRDGQRWLT EARKKTVVQI PAPTTAKQVR EFLGTAGFCR LWIPGFATLA

SLRGGQRWLT EARKKTVVQI PAPTTAKQVR EFLGTAGFCR LWIPGFATLA

SLRGGQRWLT EARKRTVVQI PAPTTAKQVR EFLGTAGFCR LWIPGFATLA

SLRGGQRWLT EARKKTVVQI PAPTTAKQVR EFLGTAGFCR LWIPGFATLA

SLRGGQRWLT EARKKTVVQI PAPTTAKQVR EFLGTAGFCR LWIPGFATLA

SLRGGQRWLT EARKRTVVQI PAPTTAKQVR EFLGTAGFCR LWIPGFATLA

SLQGGQRWLT EARKKTVVQI PAPTTAKQVR EFLGTAGFCR LWIPGFATLA

SLRDGQRWLT EARKKTVVQI PAPTTAKQVR EFLGTAGFCR LWIPGFATLA

SLRDGQRWLT EARKKTVVQI PAPTTAKQVR EFLGTAGFCR LWIPGFATLA

SLRDRQRWLT EARKKTVVQI LAPTTAKQVR EFLGTAGFCR LWIPGFATLE

SLRGGQRWLT EARKRTVVQI PAPTTAKQVR EFLGTAGFCR LWIPGFATLA

SLRGGQRWLT EARKKTVVQI PAPTTAKQVR EFLGTAGFCR LWIPGFATLA

SLRDGQRWLT EARKRTVVQI LAPTTAKQVR EFLGTAGFCR LWIPGFTTLA

SLRDEQRWLT EARKRTVVQI LAPTTAKQVR EFLGTAGFCR LWIPGFTTLA

SLRDGQRWLT EARKRTVVQI LAPTTAKQVR EFLGTAGFCR LWIPGFATLA

SLQDGQRWLT EARKRTVAQI PTPTTAKQVR EFLGTA---- ----GFATLA

SLRGGQRWLT EARKKTVVQI PAPTTAKQVR EFLGTAGFCR LWILGFATLT

SLRGGQRWLT EARKRTVVQI PAPTTAKQVR EFLGTAGFCR LWIPGFATLT

SLRGGQRWLT EARKKTVVQI PAPTTAKQVR EFLGTAGFCR LWIPGFATLA

SLRDGQRWLT EARKRTVVQI LVPTTAKQVR EFLGTAGFC- LWIPGFATLA

SLRDGQRWLT EARKRTVVQI LVPTTAKQVR EFLGTAGFC- LWIPGFATLA

SLWGGQRWLT GAWKKTVVQI PAPTTAKQVR E-LGTAGFCR LWIPRFATLA

SLRD-QRWLT EARKRTVVQI PAPTTAKQVR EFLGTAGFCR LWIPGFSTLA

SLRDGQRWLT EARKRTVVQI LVPTMSRQFR EFLGTAGFCR LWILGFATLA

SLWDGKRWLA EAQKRTVVQI LVPTMARQVR EFLGTAGFCR LWILGFATLA

TLRGGKRWLT EARKKTVMMI PPPTTPRQVR EFLGTAGFCR LWIPGFATLA

LLKGGKRWLT PARKATVMKI PTPTTPRQVR EFLGTAGFCR LWIPGFASLA

LLKEGKRWLT PARKATVMKI PVPTTPRQVR EFLGTAGFCR LWIPGFASLA

SLRDGQRWLT EARKKTVVQI PAPTTAKQVE RVLGTAGFCR LWIPGFATL-

ILKDGKRWLT EARKKTVTQI PVPSTHRQVR EFLGTAGFCR LWIPRYATLA

ILKDGKRWLT KALKKTVTQI PVPSTHRQVR EFLSTAGFCQ LWIPRYASLV

ILKDGKRWLT EARKKTVTQI PVPSTHRQVR EFLGTAGFCR LWIPRYASL-

LLREGQRWLT EARKETVMGQ PVPKTPRQLR EFLGTAGFCR LWIPGFAEMA

LLKEGQRWLT EARKETVMGQ PTPKTPRQLR EFLGTAGFCR LWIPGFAEMA

LLKEGQRWLT EARKETVMGQ PTPKTPRQLR EFLGTAGFCR LWIPGFAEMA

LLKEGQRWLT EARKETVMGQ PTPKTPRQLR EFLGTAGFCR LWIPGFAEMA

SLKDGQRWLT KARKEAILSI PVPKNSRQVR EFLGTAGYCR LWIPGFAELA

ILSEGKRWLT PGRIETVARI PPPQSPREVR EFLGTAGFCR LWIPGFAELA

ILSEGKRWLT PGRIETVARI PPPRNPREVR EFLGTAGFCR LWIPGFAELA

LLKGGQRWLT ESRKDTVAQI PAPKNARQVR EFWGMAGFCR LWIPGFAKLT

KLKEGTRWLT EAMKETILRL PVPTSAREVR EFLGTTGYCR LWILGYAEIA

KIHKGSRSLS NSRTQAILQI PVPKTKRQVR EFLGTIGYCR LWIPGFAELA

APLYPLTKEK GEFSWAPEHQ KAFDAIKKAL LSAPALALPD VTKPFTLYVD
APLYPLTKEK GEFSWAPEHQ KAFDAIKKAL LSAPALALPD VTKPFTLYVD
APLYPLTKEK GEFSWAPEHQ KAFDAIKKAL LSAPALALPD VTKPFTLYVD
APLYPLTKEK GEFSWAPEHQ KAFDAIKKAL LSAPALALPD VTKPFTLYVD
APLYPLTKEK GEFSWAPEHQ KAFDAIKKAL LSAPALALPD VTKPFTLYVD
APLYPLTKEK GEFSWAPEHQ KAFDAIKKAL LSAPALALPD VTKPFTLYVD
APLYPLTKEK GEFSWAPEHQ KAFDAIKKAL LSAPALALPD VTKPFTLYVD
APLYPLTKEK GEFSWAPEHQ KAFDAIKKAL LSAPALALPD VTKPFTLYVD
APLYPLTKEK GEFSWAPEHQ KAFDAIKKAL LSAPALALPD VTKPFTLYVD
APLYPLTKEK GEFSWAPEHQ KAFDAIKKAL LSAPALALPD VTKPFTLYVD
APLYPLTKEK GEFSWAPEHQ KAFDAIKKAL LSAPALALPD VTKPFTLYVD
APLYPLTKEK GEFSWAPEHQ KAFDAIKKAL LSAPALALPD VTKPFTLYVD
APLYPLTKEK GEFSWAPEHQ KAFDAIKKAL LSAPALALPD VTKPFTLYVD
APLYPLTKEK GEFSWAPEHQ KAFDAIKKAL LSAPALALPD VTKPFTLYVD
APLYPLTKEK GEFSWAPEHQ KAFDAIKKAL LSAPALALPD VTKPFTLYVD
APLYPLTKEK GEFSWAPEHQ KAFDAIKKAL LSAPALALPD VTKPFTLYVD
APLYPLTKKK GEFSWAPEHQ KAFDAIKKAL LSAPALALPD VTKPFTLYVD
APLYPLTKEK G-FSWAPEHQ KAFDAIKKAL LSAPALALPD VTKHFTLYVD
APLYPLTKEK GEFSWAPEHQ KAFDAIKKAL LSAPALALPD VTKPFTLYVD
APLYPLTKEK GEFSWAPEHQ KAFDAIKKAL LSAPALALPD VTKPFTLYVD
APLYPLTKEK GEFSWAPEHQ KAFDAIKKAL LSAPALALPD VTKPFTLYVD
APLYPLTKEK GEFSWAPEHQ KAFDAIKKAL LSAPALALPD VTKPFTLYVD
APLYPLTKEK GEFSWAPEHQ KAFDAIKKAL LSAPALALPD VTKPFTLYVD
APLYPLTKEK GEFSWAPEHQ KAFDAIKKAL LSAPALALPD VTKPFTLYVD
APLYPLTKEK GEFSWAPEHQ KAFDAIKKAL LSAPALALPD VTKPFTLYVD
APLYPLTKEK GEFSWAPEHQ KAFDAIKKAL LSAPALALPD VTKPFTLYVD
APLYPLTKEK GEFSWAPEHQ KAFDAIKKAL LSAPALALPD VTKPFTLYVD
APLYPLTKEK GEFSWAPEHQ KAFDAIKKAL LSAPALALPD VTKPFTLYVD
APLYPLTKEK GGFSWAPEHQ KAFDAIKKAL LSAPALALPD VTKPFTLYVD
APLYPLTKEK GEFSWAPEHQ KAFDAIKKAL LSAPALALPD VTKPFTLYVD
APLYPLTKEK GEFSWAPEHQ KAFDAIKKAL LSAPALALPD VTKPFTLYVD
APLYPLTKEK GEFSWAPEHQ KAFDAIKKAL LSAPALALPD VTKPFTLYVD
APLYPLTKEK GEFSWAPEHQ KAFDAIKKAL LSAPALALPD VTKPFTLYVD
APLYPLTKEK GEFSWAPEHQ KAFDAIKKAL LSAPALALPD VTKPFTLYVD
APLYPLTKEK GEFSWAPEHQ KAFDAIKKAL LSAPALALPD VTKPFTLYVD
APLYPLTKEK GEFSWAPEHQ KAFDAIKKAL LSAPALALPD VTKPFTLYVD
APLYPLTKEK GEFSWAPEHQ KTFDAIKKAL LSTPALALPD VTKPFTLYVN
APLYQLTKEK GEFSWAPEHQ KAFDAIKKAL LSAPALALPD VTKPFTLYVD
APLYPLTKEK GEFSWAPEHQ KAFDAIKKAL LSAPALALPD VTKPFTLYVE
APLYPLTKEK REFSWAPEHQ KAFDAIKKAL LSAPALALPD VTKPFTLYVD
APLYPLTKEK GEFSWAPEHQ KAFDAIKKAL LSAPALALPD VTKPFTLYVD
APLYPLTKEK GEFSWAPEHQ NAFDAIKKAL LSAPALALSD VTKPFTLYVD
APLYPLTKEK GEFSWAPEHQ KAFDAIKKAL LSAPALALPD VTKPFTLYVD
APLYPLSKEK GEFSWAPEHQ KAFDAIKKAL LSTPALALPD ITKPFTLYVD

APLYPLTKEK GEFSWAPEHQ KAFDAIKKAL LSTPALALPD VTKPFTLYVD
APLYPLTKEK GEFSWAPEHQ KAFDAIKKAL LSAPALALPD VTKPFTLYVD
APLYPLTKEK GEFSWAPEHQ KAFDAIKKAL LSTPALALPD VTKPFTLYVD
APLYPLTKEK GEFSWAPEHQ KAFDAIKKAL LSTPALALPD VTKPFTLYVN
APLYPLTKEK GEFSWAPEHQ KAFDAIKKAL LSAPALALPD VTKPFTLYVD
APLYPLTKEK GEFSWAPEHQ KAFDAIKKAL LSAPALALPD VTKPFTLYVD
APLYPLTKEK GEFSWAPEHQ KAFDASKKAL LSAPALALPD VTKPFTLYVD
APLYPLTKEK GKFSWAPEHQ KTFDAIKKAL LSAPALALPD VTKPFTLYVD
APLYPLTKEK GEFSWAPEHQ KAFDAIKKAL LSAPALALPD VTKPFTLYVD
APLYPLTKEK GEFSCAPKHQ KAFDAIKKAL LSAPALALPD VTKPFTLYVD
APLYPLTKEK GEFSCAPEHQ KAFDAIKKAL LSAPALALPD VTKPFTLYVD
APLYPLTKEK GEFSWAPEHQ KAFDAIKKAL LSAPALALPD VTKPFTLYVD
APLYPLTKEK GEFSWAPEHQ KAFDAIKKAL LSAPALALPD VTKPFTLYVD
APFYSLTKEK GKFTWTSEHQ GAFDAVKKAL LSVPALALLD MTKPFTLYVD
APLYSLTKEK EGFTWTSKHQ RVFDTIKKAL LSAPALALSD VTKPFTLYVD
APLYPLTREG IPFEWKEEHQ RAFEAIKSSL MTAPALALPD LTKSFVLYVD
APLYPLTREK VPFTWTEAHQ EAFGRIKEAL LSAPALALPD LTKPFALYVD
APLYPLTKES IPFIWTEEHQ QAFDHIKKAL LSAPALALPD LTKPFTLYID
APLYPLTKEK GEFS-APEHQ KA-DAIKKAL LSAPALALPD VT-PFTLY-D
APLYPLTKNS TPFIWGPGQQ EAFDAIKQAL LTAPALALPD VTKPFTLFID
APLYPLTKIS TPFTWGPEQQ EAFDTIKQAL LTALALALPD V----TLFVD
APLYPLTKNS TPFARGPEQQ EAFDTIKQA- LTAPALALPD VTKPFTLFVD
APLYPLTKTG TLFNWGPDQQ KAYQEIKQAL LTAPALGLPD LTKPFELFVD
APLYPLTKTG TLFNWGPDQQ KAYQEIKQAL LTAPALGLPD LTKPFELFVD
APLYPLTKTG TLFEWGPDQQ KAYQEIKQAL LTAPALGLPD LTKPFELFVD
APLYPLTKTG TLFNWGPDQQ KAYQEIKQAL LTAPALGLPD LTKPFELFVD
APLYPLTRPG TLFQWGTEQQ LAFEDIKKAL LSSPALGLPD ITKPFELFID
APLYALTKES APFTWQEKHQ SAFEALKEAL LSAPALGLPD TSKPFTLFID
APLYALTKES TPFTWQTEHQ LAFEALKKAL LSAPALGLPD TSKPFTLFLD
APLYPLTKNS TPFVWGDKEQ RAFDQIKRAL LSAPALGLPD VTKPFHLYVA
KPLYEATKDK VPWAWGSDQQ KAYDELKVAL LRAPALALPD PLKPFTLFVD
QPLYAATRGG NPLVWGEKEE EAFQSLKLAL TQPPALALPS LDKPFQLFVE

ERKGVARGVL TQTLGPWRRP VAYLSKKLDP VASGWPICLK AIAAVAILVK
ERKGVARGVL TQTLGPWRRP VAYLSKKLDP VASGWPICLK AIAAVAILVK
ERKGVARGVL TQTLGPWRRP VAYLSKKLDP VASGWPICLK AIAAVAILVK
ERKGVARGVL TQTLGPWRRP VAYLSKKLDP VASGWPVCLK AIAAVAILVK
ERKGVARGVL TQTLGPWRRP VAYLSKKLDP VASGWPICLK AIAAVAILVK
ERKGVARGVL TQTLGPWRRP VAYLSKKLDP VASGWPVCLK AIAAVAILVK
ERKGVARGVL TQTLGPWRRP VAYLSKKLDP VASGWPVCLK AIAAVAILVK
ERKGVARGVL TQTLGPWRRP VAYLSKKLDP VASGWPVCLK AIAAVAILVK
ERKGVARGVL TQTLGPWRRP VAYLSKKLDP VASGWPVCLK AIAAVAILVK
ERKGVARGVL TQTLGPWRRP VAYLSKKLDP VASGWPVCLK AIAAVAILVK
EHKGVARGVL TQTLGPWRRP VAYLSKKLDP VASGWPVCLK AIAAVAILVK

ERKGVARGVL TQTLGPWRRP VAYLSKKLDP VASGWPICLK AIAAVAILVK
ERKGVARGVL TQTLGPWRRP VAYLSKKLDP VASGWPVCLK AIAAVAILVK
ERKGVARGVL TQTLGPWRRP VAYLSKKLDP VASGWPVCLK AIAAVAILVK
ERKGVARGVL TQTLGPWRRP VAYLSKKLDP VASGWPVCLK AIAAVAILVK
ERKGVARGVL TQTLGPWRRP VAYLSKKLDP VASGWPICLK AIAAVAILVK
ERKGVARGIL TQTLGPWRRP VAYLSKKLDP VASGWPICLK AIAAVAILVK
ERKGVARGVL TQTLGPWRRP VAYLSKKLDP VASGWPVCLK AIAAVAILVK
ERKGVARGVL TQTLGPWRRP VAYLSKKLDP VASGWPICLK AIAAVAILVK
ERKGVARGVL TQTLGPWRRP VAYLSKKLDP VASGWPVCLK AIAAVAILVK
ERKGVARGVL TQTLGPWRRP VAYLSKKLDP VASGWPVCLK AIAAVAILVK
ERKGVARGVL TQTLGPWRRP VAYLSKKLDP VASGWPICLK AIAAVAILVK
ERKGVARGVL TQTLGPWRRP VAYLSKKLDP VASGWPVCLK AIAAVAILVK
ERKGVARGVL TQTLGPWRRP VAYLSKKLDP VASGWPVCLK AIAAVAILVK
ERKGVARGVL TQTLGPWRRP VAYLSKKLDP VASGWPVCLK AIAAVAILVK
ERKGVARGVL TQTLGPWRRP VAYLSKKLDP VASGWPVCLK AIAAVAILVK
ERKGVARGVL TQTLGPWRRP VAYLSKKLDP VASGWPVCLK AIAAVAILVK
ERKGVARGVL TQTLGPWRRP VAYLSKKLDP VASGWPVCLK AIAAVAILVK
ERKGVARGVL TQTLGPWRRP VAYLSKKLDP VASGWPVCLK AIAAVAILVK
ERKGVARGVL TQTLGPWRRP VAYLSKKLDP VASGWPVCLK AIAAVAILVK
ERKGVARGVL TQTLGPWRRP VAYLSKKLDP VASGWPVCLK AIAAVAILVK
ERKGVARGVL TQTLGPWRRP VAYLSKKLDP VASGWPVCLK AIAAVAILVK
ERKGVAQGVL TQTLGPWRRP VAYLSKKLDP VASGWPVCLK AIAAVAILVK
ERKGVARGVL TQTLGPWRRP VAYLSKKLDP VASGWPVCLK AIAAVAILVK
ERKGVARGVL TQTLGPWRRP VAYLSKKLDP IASGWPVCLK AIAAVAILVK
ERKGVARGVL TQTLGPWRRP VAYLSKKLDP VASGWPICLK AIAAVAILVK
ERKGVARGVL TQTLGPWRRP VAYLSKKLDP VASGWPVCLK AIAAVAILVK
ERKGVARGVL TQTLGPWRRP VAYLSKKLDP VASGWPVCLK AIAAVAILVK
ERKGVARGVL TQTLGPWRRP VAYLSKKLDP VASGWPVCLK AIAAVAILVK
ERKGVARGVL TQTLGPWRRP VAYLSKKLDP VASGWPVCLK AIAAVAILVK
ERKGVARGVL TQTLGPWRRP VAYLSKKLDP VASGWPVCLK AIAAVAILVK
EHKGVARGVL TQTLGPWRRP VAYLSKKLDP VASGWPVCLK AIAAVAILVK
ERKGVARGVL T-TLGPWRRP VAYLSKKLDP IASGWPVCLK AIAAVAILVK
ECNGVARGVL TQTLGPWRRP VAYLSKKLDP VASGWPVCLK AIAAVAILVK
ERKGVARGVL TQTLGPWRRP VAYLSKKLDP VASGWPICLK AIAAVAILVK
ERKGVARGVL TQTLGPWRRP VAYLSKKLDP VASGWPVCLK AIAAVAILVK
ERKGVARGVL TQSLGPWRRP VAYLSKKLDP VASGWPICLK AIAAVAILVK
ERKGVARGVL TQSLGPWRRP VAYLSKKLDP VASGWPICLK AIAAVAILVK
EHKGVARGVL TQSLGPWRRP VAYLSKKLDP VASGWPVCLK AIAAVAILVK
ERKGIARGVL TQTLGPWRRP VAYLSKKLDP VASGWPVCLK AIAAVAILVK
ERKGVARGVL TQTLGPWRRP VAYLSKKLDP VASGWPICLK AIAALAILVK
EHKEIARGVL TQTLGPWRRP VAYLSKKLDP VASGWPVCLK AIAAVAILVK
ERKGVAQGVL TQSLGPWRRP VDYLSKKLDP VASGWPVCLK AIAAMAILVK
ERKGVAQGVL TQSLGPWRRP VDYLSKKLDP VASGWPVCLK AIAAMAILVK
ERKGVAQGVL TQSLGPWRRP VDYLSKKLDP VASGWPVCLK AIAAMAILVK

ERKGVARGVL TQTLGPWRRP VAYLSKKLDP VASGWPVCLK AIAAVAILVK
ECKGVAQGVL TQSLGPWRRP VAYLSKKLDP VASGWPVCLK AIAAVAILVK
ERKGIARGVL TQSLGPWRRP IAYLSKKLDP VASGWPVCLK AIVAVSTLIK
EHKGIAQGVL TQSLGPWRRP VAYLSKKLDS VASGWPVCLK SIVAMPTLIK
ERAGIARGVL TQALGPWKRP VAYLSKKLDP VASGWPTCLK AIAAVALLIK
EKEGVARGVL TQTLGPWRRP VAYLSKKLDP VASGWPTCLK AIAAVALLLK
ERAGVARGVL TQTLGPWRRP VAYLSKKLDP VASGWPTCLK AVAAVALLLK
ERKGVARGVL T-TLGPWRRP VAYLSKKLDP IASGWPVCLK AIAAVAILVK
ERKGVARGVL TQSLGPWKRP VAYLSKKLDP VASGWPTCLQ AIAAVASLVR
ERKGVARGVL TQPLGPWKRP VAYLSKKLDL VASGWPTCLR AIAAVASLVK
ERKGVARGVL TQLLGPWKRP VAYLSKKLDP VASGWPTCLR AIAAVASLVK
EKQGYAKGVL TQKLGPWRRP VAYLSKKLDP VAAGWPPCLR MVAAIAVLTK
EKQGYAKGVL TQKLGPWRRP VAYLSKKLDP VAAGWPPCLR MVAAIAVLTK
EKQGYAKGVL TQKLGPWRRP VAYLSKKLDP VAAGWPPCLR MVAAIAVLTK
EKQGYAKGVL TQKLGPWRRP VAYLSKKLDP VAAGWPPCLR MVAAIAVLTK
ENSGFAKGVL VQKLGPWKRP VAYLSKKLDT VASGWPPCLR MVAAIAILVK
EKQGIAKGVL TQKLGPWKRP VAYLSKKLDP VAAGWPPCLR IMAATAMLVK
ERQGIAKGVL TQKLGPWKRP VAYLSKKLDP VAAGWPPCLR IMAATAMLVK
ENKGIAKGVL TQKLGPWNRP VAYLSKKMDP VVSGWPTCLK IIAAVAVLVK
ERRGIAKGVL MQRLGPWKRP VAYLSKKLDP VAAGWPPCLR IIAAVALMVK
ETSGAAKGVL TQALGPWKRP VAYLSKRLDP VAAGWPRCLR AIAAAALLTR

DADKLTLGQN ITVTAPHALE NTVRQPPDRW MTNARMTHYQ SLLLTERVTF
DADKLTLGQN ITVIAPHALE NIVRQPPDRW MTNARMTHYQ SLLLTERVTF
DADKLTLGQN ITVIAPHALE NIVRQPPDRW MTNARMTHYQ SLLLTERVTF
DADKLTLGQN ITVIAPHALE NIVRQPPDRW MTNARMTHYQ SLLLTERVTF
DADKLTLGQN ITVIAPHALE NIVRQPPDRW MTNARMTHYQ SLLLTERVTF
DADKLTLGQN ITVIAPHALE NIVRQPPDRW MTNARMTHYQ SLLLTERVTF
DADKLTLGQN ITVIAPHALE NIVRQPPDRW MTNARMTHYQ SLLLTERVTF
DADKLTLGQN ITVIAPHALE NIVRQPPDRW MTNARMTHYQ SLLLTERVTF
DADKLTLGQN ITVIAPHALE NIVRQPPDRW MTNARMTHYQ SLLLTERVTF
DADKLTLGQN ITVIAPHALE NIVRQPPDRW MTNARMTHYQ SLLLTERVTF
DADKLTLGQN ITVIAPHALE NIVRQPPDRW MTNARMTHYQ SLLLTERVTF
DADKLTLGQN ITVIAPHALE NIVRQPPDRW MTNARMTHYQ SLLLTERVTF
DADKLTLGQN ITVIAPHALE NIVRQPPDRW MTNARMTHYQ SLLLTERVTF
DADKLTLGQN ITVIAPHALE NIVRQPPDRW MTNARMTHYQ SLLLTERVTF
DADKLTLGQN ITVIAPHALE NIVRQPPDRW MTNARMTHYQ SLLLTERVTF
DADKLTLGQN ITVI-PHALE NIVRQPPDRW MTNARMTHYQ SLLLTERVTF
DADKLTLGQN ITVIAPHTLE NIVRQPPDRW MTNARMTHYQ SLLLTERVTF
DADKLTLGQN ITVIAPHALE NIVRQPPDRW MTNARMTHYQ SLLLTERVTF
DADKLTLGQN ITVIAPHALE NIVRQPPDRW MTNARMTHYQ SLLLTERVTF
DADKLTLGQN ITVIAPHALE NIVRQPPDRW MTNARMTHYQ SLLLTERVTF
DADKLTLGQN ITVIAPHALE NIVRQPPDRW MTNARMTHYQ SLLLTERVTF
DADKLTLGQN ITVIAPHALE NIVRQPPDRW MTNARMTHYQ SLLLTERVTF

DADKLTLGQN ITVIAPHALE NIVRQPPDRW MTNARMTHYQ SLLLTERVTF
DADKLTLGQN ITVIAPHALE NIVRQPPDRW MTNARMTHYQ SLLLTERVTF
DADKLTLGQN ITVIAPHALE NIVRQPPDRW MTNARMTHYQ SLLLTERVTF
DADKLTLGQN ITVIAPHALE NIIRQPPDRW MTNARMTHYQ SLLLTERVTF
DADKLTLGQN ITVIAPHALE NIVRQPPDRW MTNARMTHYQ SLLLTERVTF
DADKLTLGQN ITVIAPHALE NIVRQPPDRW MTNARMTHYQ SLLLTERVTF
DADKLTLGQN ITVIAPHALE NIVRQPPDRW MTNARMTHYQ SLLLTERVTF
DADKLTLGQN ITVIAPHALE NIVRQPPDRW MTNARMTHYQ SLLLTERVTF
DADKLTLRQN ITVIAPHALE NIVRQPPDRW MTNARMTHYQ SLLLTERVTF
DADKLTLGQN ITVIAPHALE NIVRQPPDRW MTNARMTHYQ SLLLTERVTF
DADKLTLGQN ITVIAPHALE NIVRQSPDRW MTNARMTHYQ SLLLTERVTF
DADKLTLGQN ITVIAPHALE NIVRQPPDRW MTNARMTHYQ SLLLTERVTF
DADKLTLGQN ITIIAPHALE NIVRQPPDRW MTNARMTHYQ SLLLTERIMF
DADKLTLGQN ITVIAPHALE NIVRQPPDRW MTNARMTHYQ SLLLTERVTF
DADKLTLGQN ITVIAPHALE NIVRQPPDRW MTNARMTHYQ SLLLTERVTF
DADKLTLGQN ITVIAPHALE NIVRQPPDRW MTNARMTHYQ SLLLTERVTF
DADKLTLGQN ITVIAPHALE NIVRQPPDRW MTNA-MTHYQ SLLLTERVTF
DADKLTLGQN ITVIAPHALE NIVRQPPDRW MTNARMTHYQ SLLLTERVTF
DADKLTLGQN ITVIAPHALE NIVRQPPDRW MTNARMTHYQ SLLLTERVTF
DADKLTLGQN ITIIAPHALE NIIRQPPDRW MTNACMSHYQ SLLLTERITF
DADKLTLGQN ITIIAP-ALE NIVRQPPDRW MTNARMTHYQ SLLLTERITF
DADKLTLGQN ITIIAPHALE NIVQQPPDRW MTNARMSHYQ SLLLTERITF
DADKLTLGQN ITVIAPHALE NIVRQPPDRW MTNARMTHYQ SLLLTERITF
DADKLTLGQN ITVIAPHALE NIVRQPPDRW MTNARMTHYQ SLLLTERITF
DADKLTLGQN ITVIAPHALE NIVRQPPDRW MTNARMTHYQ SLLLTERITF
DADKLTLGQN ITVIAPHALE NIVRQPPDRW MTNARMTHYQ SLLLTERITF
DADKLTLGQN ITVIAPHALE NIVRQPPDRW MTNARMTHYQ SLLLTERITF
DADKLTLGQN ITVIAPHALE NIVRQPPDRW MTNARMTHYQ SLLLTERVTF
DADKLTLGQN ITVIAPHALE NIIRQPPD-W MTNARMTHYQ SLLLTERVTF
DADKLTLGQN ITVIAPHTLE NIVRQPPDRW MTNAHMTHYQ SLLLTERVTF
DADKLTLGQN ITIIAPHALE NIVRQPPDRW MTNARMTHYQ SLLLTERITF
DADKLTLGQN ITIIAPHALE NIVRQPPDRW MTNACMTHYQ SLLLTERITF
DADKLTLGQN ITIIAPHALE NIVRQPPDRW MTNACMTHYQ SLLLTERITF
DADKLTLGQN ITVIAPHALE NIVRQPPDRW MTNVCMTRYQ SLLLTERVTF
DTDKLTLGQN ITIIAPHALE NIVRQPPDRW MTNARMTHYQ SLLLTERITF
DADKLTLEDN ---LPPHALE SIVRQPPDRW ITNAHMTHYQ SLLLTNRITF
DADKLTLGQK ITVIAPHAL- STVQQPPDQW MTNAQMAHYQ SLLLTDRVMF
DADKLTMGQQ VTVVAPHALE SIVRQPPDRW MTNARMTHYQ SLLLNDRVTF
DADKLTLGQN VLVIAPHNLE SIVRQPPDRW MTNARMTHYQ SLLLNERVSF
DADKLTLGQN VTVIASHSLE SIVRQPPDRW MTNARMTHYQ SLLLNERVSF
DADKLTLGQN -TIIAPHAL- NIVRQPPDRW MTNARMTHYQ SLLLTERVTF
DADKLTLGQR APCIRPHALE SIIRQPPERW MSNARMTHYQ SLLLDDRIGF
DADRLTLGQK LTVIAPHTLE SIIRQPPERW MSNAQMTHYQ SLLLNDRIEF
DADKLTLGQK LTVIAPHTLE SIIRQPPERW MSNARMTHYQ SLLLNDRIEF

DAGKLTMGQP LVILAPHAVE ALVKQPPDRW LSNARMTHYQ AMLLDDRVQF
DAGKLTMGQP LVILAPHAVE ALVKQPPDRW LSNARMTHYQ ALLLDDRVQF
DAGKLTMGQP LVILAPHAVE ALVKQPPDRW LSNARMTHYQ ALLLDDRVQF
DAGKLTMGQP LVILAPHAVE ALVKQPPDRW LSNARMTHYQ ALLLDDRVQF
DAGKLTLGQP LTILTSHPVE ALVRQPPNKW LSNARMTHYQ AMLLDERVHF
DSAKLTLGQP LTVITPHALE AIVRQPPDRW ITNARLTHYQ ALLLDDRVQF
DSAKLTLGQP LTVITPHTLE AIVRQPPDRW ITNARLTHYQ ALLLDDRVQF
DADKLTLGQN LTITAPHALE NVIRQPPDRW LTNARMTHYQ TLLLNDRIKF
DADKLTFGQH LKVVTPHAIE GVLKYPPGRW MTNARLTHYQ GLLLDPRIIF
EASKLTFGQD IEITSSHNLE SLLRSPPDKW LTNARITQYQ VLLLDPRVRF

APPAALNPAT LLPEETDEPV THDCHQLLIE ETGVRKDLTD IPLTGEVLTW
APPAALNPAT LLPEETDEPV THDCHQLLIE ETGVRKDLTD IPLTGEVLTW
APPAALNPAT LLPEETDEPV THDCHQLLIE ETGVRKDLTD IPLTGEVLTW
APPAALNPAT LLPEETDEPV THDCHQLLIE ETGVRKDLTD IPLTGEVLTW
APPAALNPAT LLPEETDEPV THDCHQLLIE ETGVRKDLTD IPLTGEVLTW
APPAALNPAT LLPEETDEPV THDCHQLLIE ETGVRKDLTD IPLTGEVLTW
APPAALNPAT LLPEETDEPV THDCHQLLIE ETGVRKDLTD IPLTGEVLTW
APPAALNPAT LLPEETDEPV THDCHQLLIE ETGVRKDLTD IPLTGEVLTW
APPAALNPAT LLPEETDEPV THDCHQLLIE ETGVRKDLTD IPLTGEVLTW
APPAALNPAT LLPEETDEPV THDCHQLLIE ETGVRKDLTD IPLTGEVLTW
APPAALNPAT LLPEETDEPV THDCHQLLIE ETGVRKDLTD IPLTGEVLTW
APPAALNPAT LLPEETDEPV THDCHQLLIE ETGVRKDLTD IPLTGEMLTW
APPAALNPAT LLPEETDEPV THDCHQLLIE ETGVRKDLTD IPLTGEVLTW
APPAALNPAT LLPEETDEPV THDCHQLLIE ETGVRKDLTD IPLTGEVLTW
APPAALNPAT LLPEETDEPV THDCHQLLIE ETGVRKDLTD IPLTGEVLTW
APPAALNPAT LLPEETDEPV THDCHQLLIE ETGVRKDLTD IPLTGEMLTW
APPAALNPAT LLPEETDKPV THDCHQLLIE ETGVRKDLTD IPLTGEVLTW
APPAALNPAT LLPEETDEPV THDCHQLLIE ETGVRKDLTD IPLTGEVLTW
APPAALNPAT LLPEETDEPV THDCHQLLIE ETGVRKDLTD IPLTGEVLTW
APPAALNPAT LLPEETDEPV THDCHQLLIE ETGVRKDLTD IPLTGEVLTW
APPAALNPAT LLPEETDEPV THDCHQLLIE ETGVRKDLTD IPLTGEVLTW
APPAALNPAT LLPEETDEPV THDCHQLLIE ETGVRKDLTD IPLTGEVLTW
APPAALNPAT LLPEETDEPV THDCHQLLIE ETGVRKDLTD IPLTGEVLTW
APPAALNPAT LLPEETDEPV THDCHQLLIE ETGVRKDLTD IPLTGEVLTW
APPAALNPAT LLPEETDEPV THDCHQLLIE ETGVRKDLTD IPLTGEVLTW
APPAALNPAT LLPEETDEPV THDCHQLLIE ETGVRKDLTD IPLTGEVLTW
APPAALNPAT LLPEETDEPV THDCHQLLIE ETGVRKDLTD IPLTGEVLTW
APPAALNPAT LLPEETDEPV THDCHQLLIE ETGVRKDLTD IPLTGEVLTW
APPAALNPAT LLPEETDEPV THDCHQLLIE ETGVRKDLTD IPLTGEVLTW
APPAALNPAT LLPEETDEPV THDCHQLLIE ETGVRKDLTD IPLTGEVLTW
APPAALNPAT LLPEETDEPV THDCHQLLIE ETGVRKDLTD IPLTGEVLTW
APPAALNPAT LLPEETDEPV THDCHQLLIE ETGVRKDLTD IPLTGEVLTW
APPAALNPAT LLPEETDEPV THDCHQLLIE ETGVRKDLTD IPLTGEVLTW

APPAALNPAT LLPEETDEPV THDCHQLLIE ETGVRKDLTD IPLTGEVLTW

APPAALNPAT LLPEETDEPV THDCHQLLIE ETGVRKDLID IPLTGEVLTW

APPAALNPAT LLPEETDEPV THDCHQLLIE ETGVRKDLTD IPLTGEVLTW

APPAALNPAT LLPEETDEPV THDCHQLLIE ETGVRKDLTD IPLTGKVLTW

APPATLNPAT LLPEETDEPV THDCHQLLIE ETGVHKDLTD IPLTGEVLTW

APPAALNPAT LLPEEADEPV THDCHQLLIE ETGVRKDLTD IPLTGEVLTW

APPAALNPAT LLPEETDEPV THDCHQLLIE ETGVRKDLTD IPLTGEVLTW

APPAALNPAT LLPEETDESV THDCHQLLIE ETGVRKDLTD IPLTGEVLTW

APPAALNPAT LLPEETDEPV THDCHQLLIK ETGVRKDLTD IPLTGEVLTW

APPAALNPAT LLPEETDEPV THDCHQLLIE ETGVRKDLID IPLTGEVLTW

APPATLNPAT LLPEETDEPV THDCHQLLIE ETGVRKDLTD IPLTGEVLTW

TLPAALNPAT LLPEETDEPL THDCHQLLIE E--VRKDLTD IPLTGEVLTW

TLPAALNPAT LLPEETDEPL THDCHQLLIE E--VRKDLTD IPLTGEVLTW

TLPAALNPAT LLPEETDEPL THDCHQLLIE E--VRKDLTD IPLTGEVLTW

TLPAALNPAI LLPEETDEPL THDCHQLLIE E--VRKDLTD IPLTGEVLTW

APPAALNPAT LLPEETDEPV THDCHQLLIE ETGVRKDLTD IPLTGEVLTW

APPAALNPAT LLPEETDEPV THDCHQLLIE ETGVRKDLTD IPLTGEVLTW

APPATLNPAT LLPEETDEPV THDCHQLLIE ETGVRKDLTD IPLTGEVLTW

APPAALNPAT LLPEETDEPV THDCHQLLIE ETGVRKDLTD IPLTGEMLTW

APPAALNPAT LLPEETDEPV THDCQQLLIE ETGVHKDLTD IPLTGEVLTW

TPPAALNPAT LLPEETDEPV THDCQQLLIE ETGVHKDLTD IPLTGEVLTW

TPPAALNPAT LLPEETDEPV THDCQQLLIE ETGVHKDLTD IPLTGEVLTW

APRAALNPAT LLPEETDEPV THDCHQLLIE ETGVRKDLTD IPLTGEVLTW

APPAALNPAT LLPEETDEP- THDCHQLLIE EIGVHKDLPD IPLTGEVLTW

APPAILNPAT LLPEETDEPA THDCHQMLVE EIGI-KDLTD VSLTGGTLTW

TPPAILNPAT LLPEENDEPV THDFHQLLVE ETGIRKDLTD VPLTRGTLTW

APPAILNPAT LLPLTNDSVP VHRCADILAE EIGTRKDLTD QPWPG-APSW

APPAILNPAT LLPVESDDTP IHICSEILAE ETGTRPDLRD QPLPG-VPAW

APPAVLNPAT LLPVESEATP VHRCSEILAE ETGTRRDLED QPLPG-VPTW

APPAASNPAT LLPEETDEPV THDCHQVI-E ET-VRKDLID IPLTGEVLTW

GTPATLNPAT LLPETSVN-V THSCQEILAE ETGTWKDLKD QPLKESLLTW

GTPVTLNPAT LLPDISAE-V THSCQEILAE EAGTRRDLRD QPLKGSLLTW

VTPVTLNPAT LLPDISTE-V THSCQEILAE EAGTRQDLRD QPLKGNLLTW

GPVVALNPAT LLPLPEKG-A PHDCLEILAE THGTRPDLTD QPIPDADHTW

GPIVTLNPAT LLPLPEEG-L QHDCLDILAE AHGTRPDLTD QPLPDADHTW

GPIVALNPAT LLPLPEEG-L QHDCLDILAE AHGTRPDLTD QPLPDADHTW

GPVVALNPAT LLPLPEEG-L QHNCLDILAE AHGTRPDLTD QPLPDADHTW

GPTVSLNPAT LLPLPSGG-N HHDCLQILAE THGTRPDLTD QPLPDADLTW

GPPVTLNPAT LLPAPKDQQS AHDCRQVLAE THGTREDLKD QELPDADHSW

GPPVTLNPAT LLPVPENQPS PHDCRQVLAE THGTREDLKD QELPDADHTW

APATGLNPAT LLPDPDLEGS THDCQEVLAA AHGSRPDLTD LPLPDADFTW

AEPTALNPAT LLPTPDLRAP LHDCQEIMAE VTQVRPDLQD TALPNSELVW

KQTAALNPAT LLPETDDTLP IHHCLDTLDS LTSTRPDLTD QPLAQAEATL

FTDGSSYVVE GKRMAGPAVV AGTRPIWASS LPEGTSAQKA ELMALTQALR
FTDGSSYVVE GKRMAGPAVV AGTRPIWASS LPEGTSAQKA ELMALTQALR
FTDGSSYVVE GKRMAGPAVV AGTRPIWASS LPEGTSAQKA ELMALTQALR
FTDGSSYVVE GKRMAGAAVV DGTRTIWASS LPEGTSAQKA ELMALTQALR
FTDGSSYVVE GKRMAGAAVV DGTRTIWASS LPEGTSAQKA ELMALTQALR
FTDGSSYVVE GKRMAGAAVV DGTRTIWASS LPEGTSAQKA ELMALTQALR
FTDGSSYVVE GKRMAGAAVV DGTRTIWASS LPEGTSAQKA ELMALTQALR
FTDGSSYVVE GKRMAGAAVV DGTRTIWASS LPEGTSAQKA ELMALTQALR
FTDGSSYVVE GKRMAGAAVV DGTRTIWASS LPEGTSAQKA ELMALTQALR
FTDGSSYVVE GKRMAGAAVV DGTRTIWASS LPEGTSAQKA ELMALTQALR
FTDGSSYVVE GKRMAGAAVV DGTRTIWASS LPEGTSAQKA ELMALTQALR
FTDGSSYMVE GKRMAGAAVV DGTRTIWASS LPEGTSAQKA ELMALTQALR
FTDGSSYVVE GKRMAGAAVV DGTRTIWASS LPEGTSAQKA ELMALTQALR
FTDGSSYVVE GKRMAGAAVV DGTRTIWASS LPEGTSAQKA ELMALTQALR
FTDGSSYVVE GKRMAGAAVV DGTRTIWASS LPEGTSAQKA ELMALTQALR
FTDGSSYVVE GKRMAGAAVV DGTRTIWASS LPEGTSAQKA ELMALTQALR
FTDGSSYVVE GKRMAGAAMV DGTRTIWASS LPEGTSAQKA ELMALTQALR
FTDGSSYVVE GKRMAGAAVV DGTRTIWASS LPEGTSAQKA ELMALTQALR
FTDGSSYVVE GKRMAGAAVV DGTRTIWASS LPEGTSAQKA ELMALTQALR
FTDGNNYVVE DKRMAGAVVV DGTRKIWASS LPEGTSAQKA ELMALTQALR
FTDGNNYVVE DKRMAGAVVV DGTRKIWASS LPEGTSAQKA ELMALTQALR
FTDGSSYVVE GKRMAGAAVV DGTRMILASS LPEGTSAQKA ELVALTQ---
FTDGSSYVVE GKRMAGAAVV DGTRTIWASS LPEGTSAQKA ELMALTQALR
FTDGSSYVVE GKRMAGAAVV DGTRTIWASS LPEGTSAQKA ELMALTQALR
FTDGSSYVVE GKRMAGAAVV DGTRTIWASS LPEGTSAQKA ELMALTQALR
FTDGSSYVVE GKRMAGAAVV DGTRTIWASS LPEGTSAQKA ELMALTQALR
FTDGSSYVVE GKRMAGAAVV DGTRTIWASS LPEGTSAQKA ELMALTQALR
FTDGSSYVVE GKRMAGAAVV DGTHTIWASS LPEGTSAQKA ELMALTQALR
FTDGSSYVVE GKRMAGAAVV DGTRTIWASS LPEGTSAQKA ELMALTQALR
FTDGSSYVVE GKRMAGTAVV DGTHTIWASS LPEGTSAQKA ELMALTQALR
FTDGSSYVVE GKRMAGAAVV DGTRTIWASS LPEGTSAQKA ELMALTQALR
FTDGSSYVVE GKRMAGAAVV DGTRTIWASS LPEGTSAQKA ELMALTQALR
FTDGSSYVVE GKRMAGAAVV DGTRTIWASS LLEGTSAQKA ELMALTQALR
FTDGSSYVVE GKRMAGAAVV DGTRTIWASS LPEGTSAQKA -LMALTQALR
FTDGSSYVVE GKRMAGAAVV DGTRTIWASS LPEGTSAQKA ELMALTQALR
FTDGSSYVV- GKRMAGAAVV DGTRTI-ASS LPEGTSAQKA ELMALTQALR
FTDGNSYVVE GKRMAGAAVV DGTRTIWASS LPEGTSAQKA ELMALTQALR
FTDGSSYVVE CKRMAGAAVV DGTRTIWASS LPEGTSAQKA ELMALTQALR
FTDRSSYVVE GKRMAGAAVV DGTHTIWASS LPEGTSAQKA ELMALTQALR
FTDGSSYVVE GKRMAGAAVV DGTHTIWASS LPEGTSAQKA ELMALTQALW
FTDGSSYVVE GKRMAGVAVV DGTRTIWASS LPEGTSAQKA ELMALTQALG
FTDGSSYVVE GKRMAGAAVV DGTRTIWASS LPEGTSVQKA ELMALTQALR
FTDGSSYVVE GKRMAGAAVV DGTRTIWASS LPEGTSAQKA ELMALTQALR
FTDGSSYVVE GKRMAGAAVV DGTRTIWATS LPEGTSVQKA ELMALTQALR

FTDGSSYVVE GKRVAGAAVV DGTRTIWASS LPEGTSAQKA ELVALTQALR
FTDGSSYVVE GKRVAGAAVV DGTRTIWASS LPEGTSAQKA ELVALTQALR
FTDGSSYVVE GKRVAGAAVV DGTRTIWASS LPEGTSAQKA ELVALTQALR
FTDGSSYVVE GKRVAGAAVV DGTRMIWASS LPEGTSAQKA ELVALTQALR
FTDGSSYVVE GKRMARAAVV DGTRTIWASS LSEGTSAQKA ELVALTQALR
FTDGSSYVVE GKRMAGAAVV DGTRTIWASS LPEGTSAQKA ELMALTQALR
FTDGNNYVVE DKRMAGAVVV DGTRKIWASS LPEGTSAQKA ELMALTQALR
FTDGSSYVVE GKRMAGAAMV DGTRTIWASS LPEGTSAQKT ELMALTQALR
FTDGSSYVVE GKRMAGGAVV GGTRTIWARS LPEGISAQKA ELVALMQALR
FTDGSSYVVE GKRMAGGAVV GGTRTIWARS LPEGISAQKA ELVALMQALR
FTDGSSYVVE GKRMAGGAVV GGTRTIWARS LPEGISAQKA ELVALMQALR
FTDGSSYVVE GKRMAGAAVV DGTHTIWASS LLEGTSAQKA ELMALTQALQ
FTDGSSYVVE GKRMAGAAVV DRTRMIWASS LPEGTSAQKA ELVALMQALR
FTDGSSYIVE GKRMAGVAIV DGM-TVWASS LPEGTLAQKA ELMALTQALQ
FTDGSSYMVE GKRMAGAAIV DG--TVWASS LPEGTAAQKA ELVALTQAL-
YTDGSSFLIE GKRRAGAAVV DGKKVIWASA LPEGTSAQKA ELIALTQALR
YTDGSSFIMD GRRQAGAAIV DNTRTVRASN LPEGTSAQKA ELIALTQALR
YTDGSSFITE GKRRAGAPIV DGKRTVWASS LPEGTSAQKA ELVALTQALR
FTDGSSYVVE GKRMAGAA-V DGTRTIWASS LPEGTSAQKA ELMALTQALR
YSDGSSYMTE GKRVAGAAVV DDSTTIWASS LPGGISAQWA ELIALTQALK
YTDGSSCIAE GKQVAGAAVV DDSTIILASR LPEGTSAQWA ELIALTQALK
YTDGRSYIVE GKRVAGAAVV DDSTIIWASR LPEGTSAQRA ELIALTQALK
YTDGSSFLQE GQRKAGAAVT TETEVIWARA LPAGTSAQRA ELIALTQALK
YTDGSSFLQE GQRKAGAAVT TETEVIWAKA LPAGTSAQRA ELIALTQALK
YTDGSSFLQE GQRKAGAAVT TETEVVWAKA LPAGTSAQRA ELIALTQALK
YTDGSSLLQE GQRKAGAAVT TETEVIWAKA LPAGTSAQRA ELIALTQALK
YTDGSSFIRN GEREAGAAVT TESEVIWAAP LPPGTSAQRA ELIALTQALK
YTDGSSYIDS GTRRAGAAVV DGHHIIWAQS LPPGTSAQKA ELIALTKALE
YTDGSSYLDS GTRRAGAAVV DGHNTIWAQS LPPGTSAQKA ELIALTKALE
FTDGSSFLEG GKRRAGAAVV DGKQVIWAAA LPQGTSAQRA ELIAMTRALE
YTDGSSFVID GVRRAGAAVV DGGNIIWSAS LSPGTSAQKA ELIALAEALE
FTDGSSYVRD GKRYAGAAVV TLDSVIWAEP LPIGTSAQKA ELIALTKALE

LAEGKSINIY TDSRYAFATA HVHGAIYKQR GLLTSAGREI KNKEEILSLL
LAEGKSINIY TDSRYAFATA HVHGAIYKQR GLLTSAGREI KNKEEILSLL
LAEGKSINIY TDSRYAFATA HVHGAIYKQR GLLTSAGREI KNKEEILSLL
LAEGKSINIY TDSRYAFATA HVHGAIYKQR GLLTSAGREI KNKEEILSLL
LAEGKSINIY TDSRYAFATA HVHGAIYKQR GLLTSAGREI KNKEEILSLL
LAEGKSINIY TDSRYAFATA HVHGAIYKQR GLLTSAGREI KNKEEILSLL
LAEGKSINIY TDSRYAFATA HVHGAIYKQR GLLTSAGREI KNKEEILSLL
LAEGKSINIY TDSRYAFATA HVHGAIYKQR GLLTSAGREI KNKEEILSLL
LAEGKSINIY TDSRYAFATA HVHGAIYKQR GLLTSAGREI KNKEEILSLL
LAEGKSINIY TDSRYAFATA HVHGAIYKQR GLLTSAGREI KNKEEILSLL
LAEGKSINIY TDSRYAFATA HVHGAIYKQR GLLTSAGREI KNKEEILSLL

LAEGKSINIY TDSRYAFATA HVHGAIYKQR GLLTSAGREI KNKEEILSLL
LAEGKSINIY TDSRYAFATA HVHGAIYKQR GLLTSAGREI KNKEEILSLL
LAEGKSINIY TDSRYAFATA HVHGAIYKQR GLLTSAGREI KNKEEILSLL
LAEGKSINIY TDSRYAFATA HVHGAIYKQR GLLTSAGREI KNKEEILSLL
LAEGKSINIY TDSRYAFATA HVHGAIYKQR GLLTSAGREI KNKEEILSLL
LAEGKSINIY TDNRYAFATA HVHGAIYKQR GLLTSAGREI KNKEEILSLL
LAEGKSINIY TDSRYAFATA HVHGAIYKQR GLLTSAGREI KNKEEILSLL
LAEGKSINIY TDSRYAFATA HVHGAIYKQR GLLTSAGREI KNKEEILSLL
LAKGKSINIY TDSRYAFATA HVHGAIYKQR GLLTSAGREI KNKEEILSLL
LAKGKSINIY TDSRYAFATA HVHGAIYKQR GLLTSAGREI KNKEEILSLL
LAEGKSINIY TDSRYAFATV HVHGAIYKQR GLLTSAGREI KNKEEILSLL
LAEGKSINIY TDSRYAFATA HVHGAIYKQR GLLTSAGREI KNKEEILSLL
LAEGKSINIY TDSRYAFATA HVHGAIYKQR GLLTSAGREI KNKEEILSLL
LAEGKSINIY TDSRYAFATA HVHGAIYKQR GLLTSAGREI KNKEEILSLL
LAEGKSINIY TDSRYAFATA HVHGAIYKQR GLLTSAGREI KNKEEILSLL
LAEGKSINIY TDSRYAFATA HVHGAIYKQR GLLTSAGREI KNKEEILSLL
LAEGKSINIY TDSRYAFATA HVHGAIYKQR GLLTSAGREI KNKEEILSLL
LAEGKSINIY TDSRYAFATA HVHGAIYKQR GLLTSAGREI KNKEEILSLL
LAEGKSINIY TDSRYAFATA HVHGAIYKQR GLLTSAGREI KNKEEILSLL
LAEGKSINIY TDSRYAFATA HVHGAIYKQR GLLTSAGREI KNKEEILSLL
LAEGKSINIY TDSRYAFATA HVHGAIYKQR GLLTSAGREI KNKEEILSLL
LAEGKSINIY TDSRYAFATA HVHGAIYKQR GLLTSAGREI KNKEEILSLL
LAEGKSINIY TDSRYAFATA HVHGAIYKQR GLLTSAGREI KNKEEILSLL
LAEGKSINIS TDSRYAFATA HVHGAIYKQR GLLTSAGREI KNKEEILSLL
LAEGKSINIY TDSRYAFATA HVHGAIYKQR GLLTSAGREI KNKDEILSLL
LAEGKSINIY TDSRYAFATA HVHGAIYKQR GLLTSAGREI KNKEKILSLL
LAEGKSINIY TDSRYAFATA HVHGAIYKQR GLFTSAGREI KNKEEILSLL
LAEGKSINIY TDSRYAFATA HVHRAIYKQR GLLTSAGREI KNKEEILSLL
LAEGKSINIY TDSRYAFATA HVHGAIYKQR VLLTSAGREI KNKEEILSLL
LAEGKSINIY TDSRYAFATV HIHGAIYKQR GLLTSAGREI KNKEEILSLL
LAEGKSINIY TDSRYAFATA HVHGAIYKQR GLLTSAGREI KNKEEILSLL
LAEGKSINIY TDSRYAFATA HVHGAIYKQR GLLTSAGREI KNKEEILSLL
LAEGKSINIY TDSRYAFATA HVHGAIYKQK GLLTSAGREI KNKEEILSLL
LAEGKSINIY TYSRYALATA HVHGAIYKQR GLLTSAGREF KNKEEILSLL
LAEGKSINIY TYSRYALATA HVHGAIYKQR GLLTSAGREF KNKEEILSLL
LAEGKSINIY TYSRYAFATA HVHGAIYKQR GLLTSAGREV KNKEEILSLL
LAEGKSINIY TYSRYAFATA HVHGAIYKQR GLLTSAGREV KNKEEILSLL
LAEGKSINIY TDSRYAFATA HVHGAIYKQR GLLTSAGREV KNKEKILSLL
LAEGKSINIY TDSRYAFATA HIHGAIYKQR GLLTSAGREI KNKEEILSLL
LAKGKSINIY TDSRYAFATA HVHGAIYKQR GLLTSAGREI KNKEEILSLL
LAEGKSINIY TDSRYAFATA HVHGAIYKQR GLLTSTGRKI KNKEEILSLL
LAEGKSINIY TDSRYAFVTA QVHGAIYKQR GLLTSTGREV KNKEEILSLL
LAEGKSINIY TDSRYAFVTA QVHGAIYKQR GLLTSTGREV KNKEEILSLL
LAEGKSINIY TDSRYAFVTA QVHGAIYKQR GLLTSTGREV KNKEEILSLL

LAEGKSIDIY TDSRYAFATA HVHGAIYKQR GLLTSAGREI KNKEEILSLL
LAKGKSINIY TDSRYAFATA HVHGAIYKQR GLLTSAGREV KNKEEILSLL
LAEGKSVNIY TDSR-AFATA HVHGAIYKQR GLLTSAGKEI KNKEEFLSLL
LAEGKSVNIY MDSRYAFATA HVHGAIYQQR GLLTSAGKEV KNKEEILCLL
EAEGKIINIY TDSRYAFATA HIHGAIYRQR GLLTSAGKDI KNKEEILALL
LAEGKSINIY TDSRYAFATA HVHGAIYKQR GLLTSAGKDI KNKEEILALL
LAEGKNINIY TDSRYAFATA HIHGAIYKQR GLLTSAGKDI KNKEEILALL
LARRESINIY TAGRYAFATA HVHGAIYKQR GLLTSAGREI KNKEEILSLL
LAEGKRLNLY TDSCYAFATA HVHGAIYQWR GLLTSAGKDI KNRQEILELL
LAEKKRLNLY TDSRYAFATA HVHGAIYRQR GLLTSAGKDI KNKQEILELL
LAEKKRLNVY TDSRYAFATA HVHRAIYRQR GLLTSAGKDI KSKQDIIGLL
MAEGKKLNVY TDSRYAFATA HVHGEIYRRR GLLTSEGKEI KNKSEILALL
MAEGKKLNVY TDSRYAFATA HIHGEIYRRR GLLTSEGKEI KNKDEILALL
MAEGKKLNVY TDSRYAFATA HIHGEIYRRR GLLTSEGKEI KNKDEILALL
MAEGKKLNVY TDSRYAFATA HIHGEIYRRR GLLTSEGKEI KNKDEILALL
MAEGKKLTVY TDSRYAFATT HVHGEIYRRR GLLTSEGKEI KNKNEILALL
LSEGKKANIY TDSRYAFATA HTHGSIYERR GLLTSEGKEI KNKAEIIALL
LSKGKKANIY TDSRYAFATA HTHGSIYERR GLLTSEGKEI KNKAEIIALL
MAENKKVNIY TDSRYAFATA HIHGTIYQQR GLLTSSGREI KNKDEIVALL
RAEGRRVTVY TDSRYAFGTV HVHGAIYRER GFVTAEGKAL RNLPEVRRLL
WSKDKSVNIY TDSRYAFATL HVHGMIYRER GLLTAGGKAI KNAPEILALL

EAVHLPKRLA IIHCPGH
EAVHLPKRLA IIHCPGH
EAVHLPKRLA IIHCPGH
EALHLPKRLA IIHCPGH
EAVHLPKRLA IIHCPGH
EALHLPKRLA IIHCPGH
EALHLPKRLA IIHCPGH
EALHLPKRLA IIHCPGH
EALHLPKRLA IIHCPGH
EALHLPKRLA IIHCPGH
EALHLPKRLA IIHCPGH
EALHLPKRLA IIHCPGH
EALHLPKRLA IIHCPGH
EALHLPKRLA IIHCPGH
EALHLPKRLA IIHCPGH
EALHLPKRLA IIHCPGH
EALHLPKRLA IIHCPGH
EALHLPKRLA IIHCPGH
EAVHLPKRLA IIHCPGH
EALHLPKRLA IIHCPGH
EALHLPKRLA IIHCPGH
EAVHLPKRLA IIHCPGH

EALHLPKRLA IIHCPGH
EALHLPKRLA IIHCPGH
EALHLPKRLA IIHCPGH
EALHLPKRLA IIHCPGH
EALHLPKRLA IIHCPGH
EALHLPKRLA IIHCPGH
EALHLPKRLA IIHCPGH
EALHLPKRLA IIHCPGH
EALHLPKRLA IIHCPGH
EALHLPKRLA IIHCPGH
EALHLPKRLA IIHCPGH
EALHLPKRLA IIHCPGH
EALHLPKRLA IIHCPGH
EALHLPKRLA IIHCPGH
KALHLPKRLA IIHCPGH
EALHLPERLA IIHCPGH
EALHLPKRLA IIHCPGH
EALHLPKRLA IIHCPGH
EALHLPKRLA IIHCPGH
EALHLPKRLA IIHCPGH
EALHLPKRLA IIHCPGH
EALHLPKRLA IIHCPGH
EALHLPKRPA IIHCLGH
EALHLPKRLA IIHCLGH
EALHLPKRLA IIHCLGH
EALHLPKRLA IIHCLGH
EALHLPKRLA IIHCPGH
EALHLPKRLA IIHCPGH
EALHLPKRLA IIHCPGH
EALHLPKRLA IIHCPGH
EALHLPKRLA IIHCLGH
EALHLPKRLA IIHCLGH
EALHLPKRLA IIHCLGH
EALHLPKRLA IIHCPGH
EALHLPKRLA IIHCLGH
EALYLPKRLA IMHCPGH
EALHLPKRLA IMHCPGH
EAIHAPKKVA IIHCPGH
EAIHLPKRVA IIHCPGH
EAIHLPRRVA IIHCPGH
EALHLPKRLV RPHSPLH
TALHLPAELA IIHCPGH
TAIHLPAELA IIHCPGH
TAIHLPAELA IIHCPGH

| | |
|---|---|
| | KALFLPKRLS IIHCPGH |
| | KALFLPKRLS IIHCPGH |
| | KALFLPKRLS IIHCPGH |
| | KALFLPKRLS IIHCPGH |
| | EALFLPKRLS IIHCPGH |
| | KALFLPRRVA IIHCPGH |
| | KALFLPQEVA IIHCPGH |
| | VALMLPTKVS IIHCPGH |
| | MAVQMPRAVA VVHIPGH |
| | TAVWLPKRVA VMHCKGH |

Env

121 308

| | |
|---|---|
| AEMK02000133.1 | PVAIGPNKCL AEQGPPIQEQ RPSLNPSDYN TASGSVPTEP NITIKTGAKL |
| CM000812.5_c | PVAIGPNKGL AEQGPPIQEQ RPSPNPSDYN TTSGSVPTEP NITIKTGAKL |
| LT634572.1_d | PVAIGPNKGL AEQGPPIQEQ RPSPNPSDYN TTSGSVPTEP NITIKTGAKL |
| CM000817.5 | PVAIGPNKGL AEQGPPIQEQ RPSPNPSDYN TTSGSVPTEP NITIKTGAKL |
| CM000818.5_b | PVAIGPNKGL AEQGPPIQEQ RPSPNPSDYN TTSGSVPTEP NITIKTGAKL |
| CM000812.5_b | PVAIGPNKGL AEQGPPIQEQ RPSPNPSDYN TTSGSVPTEP NITIKTGAKL |
| LUXX01006697.1 | PVAIGPNKGL AEQGPPIQEQ RPSPNPSDYN TTSGSVPTEP NITIKTGAKL |
| PERV-A | PVAIGPNKGL AEQGPPIQEQ RPSPNPSDYN TTSGSVPTEP NITIKTGAKL |
| LUXS01031219.1 | PVAIGPNKGL AEQGPPIQEQ RPSPNPSDYN TTSGSVPTEP NITIKTGAKL |
| LUXX01090830.1 | PVAIGPNKGL AEQGPPIQEQ RPSPNPSDYN TTSGSVPTEP NITIKTGAKL |
| LUXU01038095.1 | PVAIGPNKGL AEQGPPIQEQ RPSPNPSDYN TTSGSVPTEP NITIKTGAKL |
| CM000819.5_b | PVAIGPNKGL AEQGPPIQEQ RPSPNPSDYN TTSGSVPTEP NITIKTGAKL |
| CM000823.5_b | PVAIGPNKGL AEQGPPIQEQ RPSPNPSDYN TTSGSVPTEP NITIKTGAKL |
| CM000830.5_c | LVAIGPNKGL AEQGPPIQEQ RPSPNPSDYN TTSGSVPTEP NITIKTGAKL |
| CM000824.5_b | PIAIGPNKGL AEQGPPIQEQ RPSPNPSDYN TTSGSVPTEP NITIKTGAKL |
| CM000816.5_b | PVAIGPNKGL AEQGPPIQEQ RPSPNPSDYN TTSGSVPTEP NITIKTGAKL |
| LUXV01020898.1 | PVAIGPNKGL AEQGPPIQEQ RPSPNPSVYN TTSGLVPPEP NFTIKTGAKL |
| CM000828.5_a | PVAIGPNKGL AEQGPPIQEQ RPSPNPSVYN TTSGLVPPEP NFTIKTGAKL |
| CM000826.5_a | PVAIGPNKGL AEQGPPIQEQ RPSPNPSDYN TTSGSVPTEP NITIKTGTKL |
| CM000814.5_a | PVAIRPNKGL AEQGPPIQEQ RPSPNPSDYN TTSGSVPTEP NITIKTGAKL |
| LT634572.1_a | PVAIGPNKGL AEQGPLIQEQ RPSPNPSDYN TTSGSVPTEP NITIKTVVKL |
| CM000828.5_c | PVTIGPNKGL AEQGPPIQEQ RPSPNPSDYN TTSGSVPTEP NITIKTGAKL |
| LT634572.1_b | PVAIGPNKGL AEQGPPIQEQ RPSPNPSDYN TTSGSVPTEP NITIKTAAKL |
| AJKK01232762.1 | PVAIGPNKGL AEQGPPIQEQ RPSPNPSDYN TTSGSVPTEP NITIKTGAKL |
| LUXW01074277.1 | PVAIGPNKGL AEQGPPIQEQ RPSPNPSDYN TTSGSVPTEP NITIKTGAKL |
| AOCR01193540.1 | PVAVGPNKVL TEQGPPIQEQ RPSPNPSDFN TTSGSVPTEP NITIKTGAKL |
| CM000814.5_b | PVAVGPNKVL TEQGPPIQEQ RPSPNPSDFN TTSGSVPTEP NVPVKTGQRL |
| LUXT01066854.1 | PVAVGPNKVL TEQGPPIQEQ RPSPNPSDFN TTSGSVPTEP NVPVKTGQRL |
| LUXX01079974.1 | PVAVGPNKVL TEQGPPIQEQ RPSPNPSDFN TTSGSVPTEP NVPVKTGQRL |
| KQ003728.1 | PVAVGPNKVL TEQGPPIQEQ RPSPNPSDFN TTSGSVPTEP NVPVKTGQRL |
| LUXY01106894.1 | PVAVGPNKVL TEQGPPIQEQ RPSPNPSDFN TTSGSVPTEP NVPVKTGQRL |

| | | | | | |
|---|---|---|---|---|---|
| LUXV01069975.1 | PVAVGPNKVL | TEKGPPIQEQ | RPSPNPSDFN | TTSGSVPTEP | NVPVKTGQRL |
| LUXS01071149.1 | PVAVGPNKVL | TEQGPPIQEQ | RPSPNPSDFN | TTSGSVPTEP | NVPVKTGQRL |
| LUXQ01024825.1 | PVAVEPNKVL | TEQGPPIQEQ | RPSPNPSDFN | TTSGSVPTEP | NVPVKTGQRL |
| LUXR01116336.1 | PVAVGPNKVL | TEQGPPIQEQ | RPSPNPSDFN | TTSGSVPTEP | NVPVKTGQRL |
| LUXW01019135.1 | PVAVGPNKVL | TEQGPPIQEQ | RPSPNPSDFN | TTSGSVPTEP | NVPVKTGQRL |
| LIDP01000003.1_a | PVAVGPNKVL | TEQGPPIQXQ | RPSPNPSDFN | TTSGXVPTEP | NVPVKTGQRL |
| AOCR01017097.1 | PMAIGPNTVL | TGQRPPTQGP | GPS-----SN | ITSGSDPTES | NSTTKMGAKL |
| AOCR01011636.1 | PMAIGPNTVL | TGQRPPTQGP | GPS-----SN | ITSGSDPTES | NSTTKMGAKL |
| AOCR01098665.1 | PMAIGPNTVL | TGQRPPTQGP | GPS-----SN | ITSGSDPTES | NSTTKMGAKL |
| AOCR01116418.1 | PMAIGPNTVL | TGQRPPTQGP | GPS-----SN | ITSGSDPTES | NSTTKMGAKL |
| AOCR01214488.1 | PMAIGPNTVL | TGQRPPTQGP | GPS-----SN | ITSGSDPTES | NSTTKMGAKL |
| AOCR01214680.1 | PMAIGPNTVL | TGQRPPTQGP | GPS-----SN | ITSGSDPTES | NSTTKMGAKL |
| PERV-C | PMAIGPNTVL | TGQRPPTQGP | GPS-----SN | ITSGSDPTES | NSTTKMGAKL |
| KQ004162.1 | PMAVGPNTVL | TGQRTPT--P | GPS-----SD | ITSELDPTES | NSTTKTGAKL |
| LUXQ01128968.1 | PMAVGPNTVL | TGQRTPT--P | GPS-----SD | ITSELDPTES | NSTTKTGAKL |
| LUXR01088996.1 | PMAVGPNTVL | TGQRTPT--P | GPS-----SD | ITSELDPTES | NSTTKTGAKL |
| CM000824.5_a | PMAIGPNTVL | TGQRSPAQGP | GPF-----FN | ITSGSDPTES | NSTTKTGTKL |
| LUXU01026052.1 | PMAIGPNTVL | TGQRSPAQGP | GPF-----FN | ITSGSDPTES | NSTTKTGTKL |
| KQ003791.1 | PMAIGPNTVL | TGQRSPAQGP | GPF-----FN | ITSGSDPTES | NSTTKTGTKL |
| LUXT01031517.1 | PMAIGPNTVL | TGQRSPAQGP | GPF-----FN | ITSGSDPTES | NSTTKTGTKL |
| LUXQ01071671.1 | PMAIGPNTVL | TGQRPPAQGP | GPF-----FN | ITSGSDPTES | NSTTKTGTKL |
| LUXV01065321.1 | PMAIGPNTVL | TGQRPPAQGP | GPF-----FN | ITSGSDPTES | NSTTKTGTKL |
| LUXT01067056.1 | PMAIGPNTVL | TGQRTPT--P | GPS-----SD | ITSKLDPTES | NSTTKTGTKL |
| LUXU01073709.1 | PMAIGPNTVL | TGQRTPT--P | GPS-----SD | ITSKLDPTES | NSTTKTGTKL |
| AEMK02000197.1 | PVAVGPDKVL | AEQGPPALEP | RPDITQPPSN | STTGLIPTSP | GVPVKTGQRL |
| CM000812.5_a | PVAVGPDKVL | AEQGPPALEP | RPDITQPPSN | GTTGLIPTSP | GVPVKTGQRL |
| CM000819.5_a | PVAVGPDKVL | AEQGPPALEP | RPDITQPPSN | GTTGLIPTSP | GVPVKTGQRL |
| LUXR01004647.1 | PVAVGPDKVL | AEQGPPALEP | RPDITQPPSN | GTTGLIPTSP | GVPVKTGQRL |
| CM000822.5_b | PVAVGPDKVL | AEQGPPALEP | RPDITQPPSN | GTTGLIPTSP | GVPVKTGQRL |
| PERV-B | PVAVGPDKVL | AEQGPPALEP | RPDITQPPSN | GTTGLIPTSP | GVPVKTGQRL |
| CM000820.5 | PVAVGPDKVL | AEQGPPGLEP | RPDITQPPSN | GTTGLIPTSP | GVPVKTGQRL |
| CM000830.5_d | PVAVGPDKIL | AEQGPPALEP | RPDITQPPSN | GTTGLIPTSP | GVPVKTGQRL |
| CM000827.5 | PVAVGPDKVL | AEQGPPALEP | RPDITQPPSN | GTTGLIPTSP | GVPVKTGQRL |
| AEMK02000137.1 | PVAVGPDKVL | AEQGPPALEP | RPDITQPPSN | GTTGLIPTSP | GVPVKTGQRL |
| CM000815.5 | PVAVGPDKVL | AEQGPPALEP | RPDITQPPSN | GTTGLIPTSP | GVPVKTGQRL |
| CM000814.5_c | PVAVGPDKVL | AEQGPPALEP | RPDITQPPSN | GTTGLIPTSP | GVPVKTGQRL |
| CM000825.5_b | PLAVGPDKVL | AEQGPPALEP | RPDITQTPSN | GTTGLIPTSP | GVPVKTGQRL |
| LUXQ01076544.1 | PVAVGPDKVL | AEQGPPALEP | RPDITQPPSN | GTTGLIPTSP | GVPVKTGQRL |
| LUXY01101100.1 | PVAVGPDKVL | AEQGPPALEP | RPDITQPPSN | GTTGLIPTSP | GVPVKTGQRL |
| AEMK02000476.1 | PVAVGPDKVL | AEQGPPALEP | RPDITQPPSN | GTTGLIPTSP | GVPVKTGQRL |
| AOCR01107158.1 | PVAVGPDKVL | AEQGPPALEP | RPDITQPPSN | GTTGLIPTSP | GVPVKTGQRL |
| CM000826.5_b | PVAVGPDKVL | AEQGPPALEP | RPDITQPPSN | GTTGLIPTSP | GVPVKTGQRL |
| LUXW01069599.1 | PVAVGSNKVL | AEQGPPALEP | RPDITQLPDN | GTTGLIPTSP | GVPVKTGQRL |
| LUXR01068772.1 | PVAVGSNKVL | AEQGPPALEP | RPDITQLPDN | GTTGLIPTSP | GVPVKTGQRL |

```
LUXS01081675.1        PVAVGSNKVL AEQGPPALEP RPDITQLPDN GTTGLIPTSP GVPVKTGQRL
LUXY01013808.1        PVAVGSNKVL AEQGPPALEP RPDITQLPDN GTTGLIPTSP GVPVKTGQRL
LUXX01033707.1        PVAVGSNKVL AEQGPPALEP RPDITQLPDN GTTGLIPTSP GVPVKTGQRL
LUXV01015952.1        PVAVGSNKVL AEQGPPALEP RPDITQLPDN GTTGLIPTSP GVPVKTGQRL
LUXU01085042.1        PVAVGSNKVL AEQGPPALEP RPDITQLPDN GTTGLIPTSP GVPVKTGQRL
LUXX01080744.1        PVAVGPNKVL AEQGPPALEP RPDITQPPGN GTTGLIPTSP GVPVKTGQRL
CM000818.5_a          PVAVGPNKVL AEQGPPALEP RPDITQPPGN GTTGLIPTSP GVPVKTGQRL
CM000828.5_d          PVAVGPNIVL AEQGPLALEP RPDITQPPDN STTGLIPTSP GVPVKTGQRL
LUXR01022139.1        PVAVGPDKVL AEQGPPALEP RPDITQPPDN STTGLIPTSP GVPVKTGQRL
LUXX01045907.1        PVAVGPDKVL AEQGPPALEP RPDITQPPDN STTGLIPTSP GVPVKTGQRL
LUXW01068183.1        PVAVGPNIVL AEQGPPALEP RPDITQPPDN STTGLIPTSP GVPVKTGQRL
KQ003125.1            PVAVGPNKVL AEQGPPALE- RPNTTQPPGN STAGLIPT-- -----TGQRL
LUXR01063446.1        PVAVGPNKVL AEQGPPALE- RPNTTQPPGN STAGLIPT-- -----TGQRL
AORO02006834.1        PVAVGPNKVL AEQGPPALE- RPNTTQPPGN STAGLIPT-- -----TGQRL
LUXV01061131.1        PVAVGPNKVL AEQGPPALE- RPNTTQPPGN STAGLIPT-- -----TGQRL
CM000823.5_a          PVAVGPNKVL AEQGPPALE- RPNTTQPPGN STAGLIPT-- -----TGQRL
LUXS01013084.1        PVAVGPNKVL AEQGPPALE- RPNTTQPPGN STAGLIPT-- -----TGQRL
LUXW01070394.1        PVAVGPNKVL AEQGPPALE- RPNTTQPPGN STAGLIPT-- -----TGQRL
LUXX01073178.1        PVAVGPNKVL AEQGPPALE- RPNTTQPPGN STAGLIPT-- -----TGQRL
LUXU01008622.1        PVAVGPNKVL AEQGPPALE- RPNTTQPPGN STAGLIPT-- -----TGQRL
LUXT01085568.1        PVAVGPNKVL AEQGPPALE- RPNTTQPPGN STAGLIPT-- -----TGQRL
LUXQ01001340.1        PVAVGPNKVL AEQGPPALE- RPNTTQPPGN STAGLIPT-- -----TGQRL
LUXY01039945.1        PVAVGPNKVL AEQGPPALE- RPNTTQPPGN STAGLIPT-- -----TGQRL
LIDP01000012.1        PVAVGPNKVL AEQGPPALE- RPNTTQPPGN STAGLIPT-- -----TGQRL
AOCR01201355.1        PVAGGPNKVL AEQGPPA--- RPDITQPPSN GTTGLIPTSP GVPVKTGQRL
eJJRV_NW_004504375.1 PRPVGPNKVL VDQGPPCGIA APTHSGSQDI MTARTTSTNP EAPHRTRQRL
eJJRV_NW_004504378.1 PRPVGPNKVL VDQGPPN--V ALVSAKTRRI MTARTTLTSP EAPHGKGQRL
eJJRV_NW_004504334.1 PSSIGPNNVL ADQGLPN--A ALVAAGTCRV TTTRITPTSP VAPHKTGQRL
KoRV                  PVVVGPDPVL AEQGPPRKIP SPPAS----- ----PIPTSP TPSPTTGDRL
GALV                  AVAVGPDLVL VEQGPPRTSL APPPSLPDSN STSAQTPTTI TPPPTTGDRL
MDEV                  VTSIGPNKVL TEQAPPVAPA VPAPPTSRPY ----TVPSTL PPLLDTENRL
ePCRV_KN678005.1      AQSIGPNKVL S--GPRP--- LPEPTLSPS- ----SIPVTP PAPINTEQRL
ePCRV_KN680906.1      AQSIGPNKVL S--GPRP--- LPEPTLSPS- ----SIPVTP SAPINTEQRL
ePCRV_KN676491.1      AQSVGPNKVL S--GQKP--- LPALTLSPS- ----SIPITP PAPTNTEQRL
ePCRV_KN677924.1      AQSVGPNKVL S--GQKP--- LPAPTLSPS- ----SIPIMP PTSTNTEQRL
ePCRV_KN676182.1      AQSVGPNKVL S--GQGP--- LPALTLSPS- ----SIPITP PAPTNTEQRL
ePCRV_KN678690.1      AQSVGPNKVL S--GRGP--- LPALT----- ---------- --PTNTEQRL
ePCRV_KN676905.1      AQLVGLNKVL S--GQKP--- LPTLTLSPS- ----SIPNNP LHQHRAE--I
OOEV                  PVSIGPNPVL ---GPPPLPP PLLPPAPKNN GTSGLKET-- IKPGTRQDRM
R-MuLV                RVPIGPNPVL ADQLSFPLPN KPAKSPPVSN ----STPTTP PPPAGTGDRL
F-MuLV                RIPIGPNPVL ADQLSFPLPN KPAKSPPASS ----STPTTP PPPAGTGDRL
M-MuLV                RVPIGPNPVL ADQQPLSKPK SPSVTKPPS- ----GTPLSP LPPAGTENRL
M-CRV                 RVPIGPNPVI TEQLPPSQPV LPRPPHPPPS GAASMVPGPP SQQPGTGDRL
FeLV                  PQAMGPNLVL PDQKPPSRQS SKVATQRLQT --TESAPRAP PKRIGTGDRL
```

| | |
|---|---|
| RfRV | STQVGPNKVL APLVP----- TKNPGSRDKD TTAGTQPKTP PATQTTEDSL |
| REV | SYEDGPNKLL QAGKPVCWDP GPTDMIREES VRRHSYPSHP PRGVDLDPQT |
| | FSLIQGAFQA LNSTTPEATS SCWLCLASGP PYYEGTARGG KFNVTKEHRD |
| | FNLIQGAFQA LNSTTPEATS SCWLCLASGP PYYEGMARGG KFNVTKEHRD |
| | FNLIQGAFQA LNSTTPEATS SCWLCLASGP PYYEGMARGG KFNVTKEHRD |
| | FNLIQGAFQA LNSTTPEATS SCWLCLASGP PYYEGMARGG KFNVTKEHRD |
| | FNLIQGAFQA LNSTTPEATS SCWLCLASGP PYYEGMARGG KFNVTKEHRD |
| | FNLIQGAFQA LNSTTPEATS SCWLCLASGP PYYEGMARGG KFNVTKEHRD |
| | FSLIQGAFQA LNSTTPEATS SCWLCLASGP PYYEGMARGG KFNVTKEHRD |
| | FSLIQGAFQA LNSTTPEATS SCWLCLASGP PYYEGMARGG KFNVTKEHRD |
| | FSLIQGAFQA LNSTTPEATS SCWLCLASGP PYYEGMARGG KFNVTKEHRD |
| | FSLIQGAFQA LNSTTPEATS SCWLCLASGP PYYEGMARGG KFNVTKEHRD |
| | FSLIQGAFQA LNSTTPEATS SCWLCLASGP PYYEGMARGG KFNVTKEHRD |
| | FSLIQGAFQA LNSTTPEATS SCWLCLASGP PYYEGMARGG KFNVTKEHRD |
| | FSLIQGAFQA LNSTTPEATS SCWLCLASGP PYYEGMARGG KFNVTKEHRD |
| | FSLIQGAFQA LNSTTPEATS SCWLCLASGP PYYEGMARGG KFNVTKEHRD |
| | FSLIQGAFQA LNSKTPEATS SCWLCLASGP PYYEGMARGG KFNVTKEHRD |
| | FSLIQGAFQA LNSTTPEATS SCWLCLASGP PYYEGMAKGG KFNVTKEHRD |
| | FSLIQGAFQA LNSTTPEATS SCWLCLASGP PYYEGMARGG KFNVTKEHRD |
| | FSLIQGAFQA LNSTTPEATS SCWLCLALSP PYYEGMARGG KFNVTKEHRD |
| | FSLIQGAFQA LNSTTPEATS SCWLCLASGP PYYEGMARGG KFNVTKEHRD |
| | FNLIQGAFQA LNSTTPEATS SCWLCLASGP PYYEEMAREG KFNVTKEHRD |
| | FSLIQGDFQA LNSTTPEATS SCWLCLASGP PYYEGMARGG KFNVTKEHRD |
| | FNLIQGAFQA LNSTTPEATS SCWLCLTSGP PYYEGMARGG KFNVTKEHRD |
| | FSLIQGDFQA LNSTTPEATS SCWLCLASGP PYYEGMARGG KFNVTKEHRD |
| | FSLIQGAFQA LNSTTPEATS SCWLCLASGP PYYEKMARGG KFNVSKKHRD |
| | FSLIQGAFQA LNSTTPEATS SCWLCLASGP PYYEKMARGG KFNVTKKHRD |
| | FSLIQGAFQA LNSKAPEATS SCWLCLASGP PYYEGMARGG KFNVTKEHRD |
| | FSLIQGAFQA LNSTTPEATS SCWLCLASGP PYYGGMAREG KFNVTKEHRD |
| | FSLIQGAFQA LNSTTPEATS SCWLCLASGP PYYGGMAREG KFNVTKEHRD |
| | FSLIQGAFQA LNSTTPEATS SCWLCLASGP PYYGGMAREG KFNVTKEHRD |
| | FSLIQGAFQA LNSTTPEATS SCWLCLASGP PYYGGMAREG KFNVTKEHRD |
| | FSLIQGAFQA LNSTTPEATS SCWLCLASGP PYYGGMAREG KFNVTKEHRD |
| | FSLIQGAFQA LNSTTPEATS SCWLCLASGP PYYGGMAREG KFNVTKEHRD |
| | FSLIQGAFQA LNSTTPEATS SCWLCLASGP PYYEGMAREG KFNVTKEHRD |
| | FSLIQGAFQA LNSTTPEATS SCWLCLASGP PYYEGMAREG KFNVTKEHRD |
| | FSLIQGAFQA LNSTTPEATS SCWLCLASGP PYYGGMAREG KFNVTKEHRD |
| | FSLIQGAFQA LNSTTPEATS SCWLCLASGP PYYXGMARRG KFNVTKEHRD |
| | FSLIQGAFQA LNSTTPEATS SCWLCLALGP PYYEGMARRG KFNVTKEHRD |
| | FSLIQGAFQA LNSTTPEATS SCWLCLALGP PYYEGMARRG KFNVTKEHRD |
| | FSLIQGAFQA LNSTTPEATS SCWLCLALGP PYYEGMARRG KFNVTKEHRD |
| | FSLIQGAFQA LNSTTPEATS SSWLCLASGP PYYEGMARRG KFNVTKEHRD |

FSLIQGAFQA LNSTTPEATS SSWLCLASGP PYYEGMARRG KFNVTKEHRD
FSLIQGAFQA LNSTTPEATS SSWLCLASGP PYYEGMARRG KFNVTKEHRD
FSLIQGAFQA LNSTTPEATS SCWLCLASGP PYYEGMARRG KFNVTKEHRD
FSLIQGAFQA LKSTSPEATS SCWLCLASGP PYYEGMAKRG KFNVTKEHRD
FSLIQGAFQA LKSTSPEATS SCWLCLASGP PYYEGMARGG KFNVTKEHRD
FSLIQGAFQA LKSTSPEATS SCWLCLALSP PYYEGMARGG KFNVTKEHRD
FSLIQGAFQA LNSTTPEATS SCWLCLASGP PYYEGMARGG KFNVTKEYRD
FSLIQGAFQA LNSTTPEATS SCWLCLASGP PYYEGMARGG KFNVTKEYRD
FSLIQGAFQA LNSTTPEATS SCWLCLASGP PYYEGMARGG KFNVTKEYRD
FSLIQGDFQA LNSTTPEATS SCWLCLASGP PYYEGMARGG KFNVTKEYRD
FSLIQGAFQA LNSTTPEATS SCWLCLASGP PYYEGMARGG KFNVTKEHRD
FSLIQGAFQA LNSTTPEATS SCWLCLASGP PYYGGMAREG KFNVTKEHRD
FSLIQGAFQA LNSTTPEATS SCWLCLTSGP PYYKKMAKRE KFNVTKKHRD
FSLIQGAFQA LNSTTPEATS SCWLCLASGP PYYEGMARGG KFNVTKEHRD
FSLIQGAFQA INSTDPDATS SCWLCLSSGP PYYEGMAKEG KFNVTKEHRN
FSLIQGAFQA INSTDPDATS SCWLCLSSGP PYYEGMAKEG KFNVTKEHRN
FSLIQGAFQA INSTDPDATS SCWLCLSSGP PYYEGMAKEG KFNVTKEHRN
FSLIQGAFQA INSTDPDATS SCWLCLSSGP PYYEGMAKEG KFNVTKEHRN
FSLIQGAFQA INSTDPDATS SCWLCLSSGP PYYEGMAKEG KFNVTKEHRN
FSLIQGAFQA INSTDPDATS SCWLCLSSGP PYYEGMAKEG KFNVTKEHRN
FSLIQGAFQA INSTDPDATS SCWLCLSSGP PYYEGMAKEG KFNVTKEHRN
FSLIQGAFQA INSTDPDATS SCWLCLSSGP PYYEGMAKEG KFNVTKEHRN
FSLIQGAFQA INSTDPDATS SCWLCLSSGP PYYEGMAKEG KFNVTKEHRN
FSLIQGAFQA INSTDPDATS SCWLCLSSGP PYYEGMAKEG KFNVTKERRN
FSLIQGAFQA INSTDPDATS SCWLCLSSGP PYYEGMAKEG KFNVTKEHRN
FSLIQGAFQA INSTDPDATS SCWLCLSSGP PYYEGMAKEG KFNVTKEHRN
FSLIQGAFQA INSTDPDATS SCWLCLSSGP PYYEGMAKEG KFNVTKEHRN
FSLIQGAFQA INSTDPDATS SCWLCLSSGP PYYEGMAKEG KFNVTKEHRN
FSLIQGAFQA INSTDPDATS SCWLCLSSGP PYYEGMAKEG KFNVTKEHRN
FSLIQGAFQA INSTDPDATS SCWLCLSSGP PYYEGMAKEG KFNVTKEHRN
FSLIQGAFQA INSTDPDATS SCWLCLSSGP PYYEGMAKEG KFNVTKEHRN
FSLIQGAFQA INSTDPDATS SCWLCLSSGP PYYEGMAKEE KFNVTKEHRN
FRLIQGAFQA ITSTDPDATS SCWLCLSSGP PYYEGMAREG KFDVTKKHRN
FRLIQGAFQA ITSTDPDATS SCWLCLSSGP PYYEGMAREG KFDVTKKHRN
FRLIQGAFQA ITSTDPDATS SCWLCLSSGP PYYEGMAREG KFDVTKKHRN
FRLIQGAFQA ITSTDPDATS SCWLCLSSGP PYYEGMAREG KFDVTKKHRN
FRLIQGAFQA ITSTDPDATS SCWLCLSSGP PYYEGMAREG KFDVTKKHRN
FRLIQGAFQA ITSTDPDATS SCWLCLSSGP PYYEGMAREG KFDVTKKHRN
FRLIQGAFQA ITSTDPDATS SCWLCLSSGP PYYEGMAREG KFDVTKEHRN
FSLIQGVFQA INSTNSDATS FCWLCLSSGP PYYEGMAREG KFNVTKEHRN
FSLIQGVFQA INSTDPDATS SCWLCLSSGL LIMRNGRR-- KFNVTKEHRN
FSLIQGAFQA INSTDPDATS SCWLCLSSGP PYYEGMAREG KFNVTKEHRN
FSLIQGAFQA INSTDPDASS SCWLCLSSGP PYYEGMAREG KFNVTKEHRN
FSLIQGAFQA INSTDPDATS SCWLCLSSGP PYYEGMAREG KFNVTKEHRN

```
FSLIQGAFQA INSTDPDASS SCWLCLSSGP PYYEGMAREG KFNVTKEHRN
FSLIQGAFQV LNSTDPDATS SCWLCLSSGP PYYEGMAREG KFNVTKEHRN
FSLIQGAFQV LNSTDPDATS SCWLCLSSGP PYYEGMAKEG KFNVTKEHRN
FSLIQGAFQV LNSTDPDATS SCWLCLSSGP PYYEGMAREG KFNVTKEHRN
FSLIQGAFQV LNSTDPDATS SCWLCLSSGP PYYEGMAREG KFNVTKEHRN
FSLIQGAFQV LNSTDPDATS SCWLCLSSGP PYYEGMAREG KFNVTKEHRN
FSLIQGAFQV LNSTDPDATS SCWLCLSSGP PYYEGMAREG KFNVTKEHRN
FSLIQGAFQV LNSTDPDATS SCWLCLSSGP PYYEGMAREG KFNVTKEHRN
FSLIQGAFQV LNSTDPDATS SCWLCLSSGP PYYEGMAREG KFNVTKEHRN
FSLIQGAFQV LNSTDPDATS SCWLCLSSGP PYYEGMAREG KFNVTKEHRN
FSLIQGAFQV LNSTDPDATS SCWLCLSSGP PYYEGMAREG KFNVTKEHRN
FSLIQGAFQV LNSTDPDATS SCWLCLSSGP PYYEGMAKEG KFNVTKEHRN
FSLIQGAFQV LNSTDPDATS SCWLCLSSGP PYYEGMAKEG KFNVTKEHRN
FSLIQGAFQV LNSTDPDATS SCWLCLSSGP PYYEGMAREG KFNVTKEHRN
FSLIQGAFQA INSTDPDATS SCWLCLSSGP PYYEGMAKEG KFNVTKEHRN
FNLVQGAFLA INATNSNATS ACWLCLSSGP SYYEGMATMG EFNVTKEHNR
FNLIQGAFLA INATNPNATS ACWLCLSLGP PYYEGMATMG EFNVTKEYNS
LNLIQGAFLT INATNPNATS ACWLCLSSGP PNYEGVANVG EFNVTKEHNR
FGLVQGAFLA LNATNPEATE SCWLCLALGP PYYEGIATPG QVTYASTD-S
FDLVQGAFLT LNATNPGATE SCWLCLAMGP PYYEAIASSG EVAYSTDL-D
VSLVQGAFLV LNRTNPNMTQ SCWLCYASNP PYYEGIAQTR TYNITSDH-S
LNLVQGAFSV LNTSDPSITN SCWLCLASRP PYYEGVAFSG NFNNATSH-S
LNLVQGAFNV LNTSDPSITN SCWLCLASRP PYYEGVAFSE NFNNATSH-S
LNLVQGAFSV LNTSNPSITK SCWLCLASRP PYYEGVAFSG NFNNTTSH-S
LNLVQ-AFSV LNTSNSSITK SCWLCLASRP PYYEGVALSG NFNNTTSH-S
LNLVQGAFSV LNTSNPSIIK SCWLCLASRP PYYEGVAFSG NFNNTTSH-S
LNLVQGAFSV LNTSNPSITK SCWLCLASRP PYYEGVAFSG NFSNTTSH-S
AEFSTGSFQV LNTSNPSITN SCWLYLASRP PYYEGVAFSG NFNNTTSH-S
LSLVQAAFTV LNATNPEATK SCWLCYAAPP PYYDAIGYSS NYTNVSSP-D
LNLVQGAYQA LNLTNPDKTQ ECWLCLVSGP PYYEGVAVLG TYSNHTSAPT
LNLVQGAYQA LNLTNPDKTQ ECWLCLVSGP PYYEGVAVLG TYSNHTSAPA
LNLVDGAYQA LNLTSPDKTQ ECWLCLVAGP PYYEGVAVLG TYSNHTSAPA
LNLVKGAYQA LNLTSPDRTQ GCWLCLVSGP PYYEGVAVLG TYSNHTSAPA
INLVQGTYLA LNATDPNKTK DCWLCLVSRP PYYEGIAILG NYSNQTNPPP
RKLVRTVYET LNATSPHLTT SCWLCYDVKP PFYEAIGLNA TYNASNKNPS
SDILEATHQV LNATNPRLAE NCWLCMTLGT PIPAAIPANG EVTLD----G

QCTWGSQNKL TLTEVSGKGT CIGRVPPSHQ HLCNHTEAFN RTSESQYLVP
QCTWGSQNKL TLTEVSGKGT CIGMVPPSHQ HLCNHTKAFN RTSESQYLVP
QCTWGSQNKL TLTEVSGKGT CIGMVPPSHQ HLCNHTEAFN RTSESQYLVP
QCTWGSQNKL TLTEVSGKGT CIGMVPPSHQ HLCNHTEAFN RTSESQYLVP
QCTWGSQNKL TLTEVSGKGT CIGMVPPSHQ HLCNHTEAFN RTSESQYLVP
QCTWGSQNKL TLTEVSGKGT CIGMVPPSHQ HLCNHTEAFN RTSESQYLVP
QCTWGSQNKL TLTEVSGKGT CIGMVPPSHQ HLCNHTEAFN RTSESQYLVP
```

QCTWGSQNKL TLTEVSGKGT CIGMVPPSHQ HLCNHTEAFN RTSESQYLVP
QCTWGSQNKL TLTEVSGKGT CIGMVPPSHQ HLCNHTEAFN RTSESQYLVP
QCTWGSQNKL TLTEVSGKGT CIGMVPPSHQ HLCNHTEAFN RTSESQYLVP
QCTWGSQNKL TLTEVSGKGT CIGMVPPSHQ HLCNHTEAFN RTSESQYLVP
QCTWGSQNKL TLTEVSGKGT CIGRVPPSHQ HLCNHTEAFN RTSESQYLVP
QCTWGSQNKL TLTEVSGKGT CIGRVPPSHQ HLCNHTEAFN RTSESQYLVP
QCTWGSQNKL TLTEVSGKGT CIGRVPPSHQ HLCNHTEAFN QTSESQYLVP
QCTWGSQNKL TLTEVSGKGT CIGRVPPSHQ HLCNHTEAFN RTSESQYLVP
QCTWGSQNKL TLTEVSGKGT CIGRVPPSHQ HLCNHTEAFN RTSESQYLVP
QCTWGSQNKL TLTEVSGKGT CIGRVPPSHQ HLCNHTEAFN RTSESQYLVP
QCTWGSQNKL TLTEVSGKGT CIGRVPPSHQ HLCNHTEAFN RTSESQYLVP
QCTWGSQNKL TLTEVSEKGT CIGRVPPSHQ HLCNHTEAFN RTSESQYLVP
QCTWGSQNKL TLTEVSGKGT CIGMVPPSHQ HLCNHTEAFN RTSESQYLVP
QCTWGSQNKL TLTEVSGKGT CIGRVPPSHQ HLCNHTEAFN -TSESQYLVP
QCTWGSQNKL TLTEVSEKDT CIGMVPPSHQ HLCNHTEAFN QTSESQYLVP
QCTWGSQNKL TLTEVSGKGT CIGRVPPSHQ HLCNHTEAFN RTSESQ--VS
QCT-GSQNKL TLTEVSGKDT CIGRVPPSHQ HLCNHTEAFN RTSESQ--VT
QCT-GSQNKL TLTEVSGKDT CIGRVPPSHQ HLCNHTEAFN RTSKSQVSGT
QCTWGSQNKL TLTEVSGKGT CIGRVPPSHQ HLCNHTEAFN RTSENQYLVP
QCTWGSQNKL TLTEVSGKGT YIGRVPPSHQ HLCNHTEAFN RTSENQYLVP
QCTWGSQNKL TLTEVSGKGT YIGRVPPSHQ HLCNHTEAFN RTSENQYLVP
QCTWGSQNKL TLTEVSGKGT YIGRVPPSHQ HLCNHTEAFN RTSENQYLVP
QCTWGSQNKL TLTEVSGKGT YIGRVPPSHQ HLCNHTEAFN RTSENQYLVP
QCTWGSQNKL TLTEVSGKGT YIGRVPPSHQ HLCNHTEAFN RTSESQYLVP
QCTWGSQNKL TLTEVSGKGT YIGRVPPSHQ HLCNHTEAFN RTSESQYLVP
QCTWGSQNKL TLTEVSGKGT YIGMVPPSHQ HLCNHTEAFN RTSESQYLVP
QCTWGSQNKL TLTEVSGKGT CIGRVPPSHQ HLCNHTEAFN RTSESQYLVP
QCTWGSQNKL TLTEVSGKGT CIGKVPPSHQ HLCNHTEAFN QTSESQYLVP
QCTWGSQNKL TLTEVSGKGT YIGRVPPSHQ HLCNHTEAFN RTSENQYLVP
QCTWGSQNKL TLTEVSGKGT XIGRVPPSHQ HLCNHTEAFN RTSENQYLVP
QCTWGSQNKL TLTEVSGKGT CIGKVPPSHQ HLCNHTEAFN QTSESQYLVP
QCTWGSQNKL TLTEVSGKGT CIGKVPPSHQ HLCNHTEAFN QTSESQYLVP
QCTWGSQNKL TLTEVSGKGT CIGKVPPSHQ HLCNHTEAFN QTSESQYLVP
QCTWGSQNKL TLTEVSGKGT CIGKVPPSHQ HLCNHTEAFN QTSESQYLVP
QCTWGSQNKL TLTEVSGKGT CIGKVPPSHQ HLCNHTEAFN QTSESQYLVP
QCTWGSQNKL TLTEVSGKGT CIGKVPPSHQ HLCNHTEAFN QTSESQYLVP
QCTWGSQNKL TLTEVSGKGT CIGKVPPSHQ HLCNHTEAFN QTSESQYLVP
QCTWGSQNKL TLTEVSGKGT CIGKVPPSHQ HLCNHTEAFN QTSESQYLVP
QCTWGSQNKL TLTEVSGKGT CIGRVPPSHQ HLCNHTEAFN RTSESQYLVP
QCTWGSQNKL TLTEVSGKGT CIGKVPPSHQ HLCNHTEAFN QTSESQYLVP
QCTWGSQNKL TLTEVSGKGT YIGRVPPSHQ HLCNHTEAFN RTSESQYLVP
QCTWGSQNKL TLTEVSGKGT YIGRVPPSHQ HLCNHTEAFN RTSESQYLVP
QCTWGSQNKL TLTEVSGKGT YVGRVPPSHQ HLCNHTEAFN RTSESQYLVP
QCTWGSQNKL TLTEVSGKGT CIGRVPPSHQ HLYNHTEAFN RTSESQYLVP

QCTWGSQNKL TLTEVSGKGT CIGRVPPSHQ HLCNHTEAFN RTSESQYLVP
QCTWGSQNKL TLTEVSGKGT YIGRVPPSHQ HLCNHTEAFN RTSESQYLVP
QCTWGSQNKL TLTEVSGKDT CIKKVPPSHQ HLYNHTEAFN QTSESQYLVP
QCTWGSQNKL TLTEVSGKDT CIKKVPPSHQ HLYNHTEAFN QTSESQYLVP
QCTWGSRNKL TLTEVSGKGT CIGKAPPSHQ HLCYSTVVYE QASENQYLVP
QCTWGSRNKL TLTEVSGKGT CIGKAPPSHQ HLCYSTVVYE QASENQYLVP
QCTWGSRNKL TLTEVSGKGT CIGKAPPSHQ HLCYSTVVYE QASENQYLVP
QCTWGSRNKL TLTEVSGKGT CIGKAPPSHQ HLCYSTVVYE QASENQYLVP
QCTWGSRNKL TLTEVSGKGT CIGKAPPSHQ HLCYSTVVYE QASENQYLVP
QCTWGSRNKL TLTEVSGKGT CIGKAPPSHQ HLCYSTVVYE QASENQYLVP
QCTWGSRNKL TLTEVSGKGT CIGKAPPSHQ HLCYSTVVYE QASENQYLVP
QCTWGSRNKL TLTEVSGKGT CIGKAPPSHQ HLCYSTVVYE QASENQYLVP
QCTWGSRNKL TLTEVSGKGT CIGKAPPSHQ HLCYSTVVYE QASENQYLVP
QCTWGSRNKL TLTEVSGKGT CIGKAPPSHQ HLCYSTVVYE QASENQYLVP
QCTWGSRNKL TLTEISGKGT CIGKAPPSHQ HLCYSTVVYE QASENQYLVP
QCTWGSQNKL TLTEVSGKGT CIGKAPPSHQ HLCYSTVVYE QASENQYLVP
QCTWGSRNKL TLTEVSGKGT CIGKAPPSHQ HLCYSTVVYE QASENQYLVP
QCTWGSRNKL TLTEVSGKGT CIGKAPPSHQ HLCYSTVVYE QASENQYLVP
QCTWGSRNKL TLTEVSGKGT CIGKAPPSHQ HLCYSTVVYE QASENQYLVP
QCTWGSRNKL TLTEVSGKGT CIGKAPPSHQ HLCYSTVVYE QASENQYLVP
QCTWGSRNKL TLTEVSGKGT CIGKAPPSHQ HLCYSTVVYE QASENQYLVP
QCTWGSRNKL TLTEVSEKGT CIGKAPPSHQ HLCYSTVVYE QASENQYLVP
QCTWGSRNKL TLTEVSGKGT CIGKAPPSHQ HLCNSTMVYE QASENQYLVP
QCTWGSRNKL TLTEVSGKGT CIGKAPPSHQ HLCNSTMVYE QASENQYLVP
QCTWGSRNKL TLTEVSGKGT CIGKAPPSHQ HLCNSTMVYE QASENQYLVP
QCTWGSRNKL TLTEVSGKGT CIGKAPPSHQ HLCNSTMVYE QASENQYLVP
QCTWGSRNKL TLTEVSGKGT CIGKAPPSHQ HLCNSTMVYE QASENQYLVP
QCTWGSRNKL TLTEVSGKGT CIGKAPPSHQ HLCNSTMVYE QASENQYLVP
QCTWGSRNKL TLTEVSGKGT CIGKAPPSHQ HLCNSTMVYE QASENQYLVP
QCTWGSRNKL TFTEVSGKGT CIGKAPPSHQ HLCNSTMVYE QASENQYLVP
QCTWGSRNKL TFTEVSGKGT CIGKAPPSHQ HLCNSTMVYE QASENQYLVP
QCTWGSRNKL TLTEVSGKGT CIGKAPPSHQ HLCNSTMVYE QALENQYLVP
QCTWGSRNKL TLTEVSGKGT CIGKAPPSHQ HLCNSTMAYE QALENQYLVP
QCTWGSRNKL TLTEVSGKGT CIGKAPPSHQ HLCNSTMVYE QALENQYLVP
QCTWGSRNKL TLTEVSGKGT CIGKAPPSHQ HLCNSTMVYE QASENQYLVP
QCTWGSRYKL TLTEVSGKGT CIGKAPPSHQ HLCNSTMVYK QASENQYLVP
QCTWGSRNKL TLTEVSGKGT CIGKAPPSHQ HLCYSTVVYE QASENQYLVP
QCTRGSRNKL TLTEVSGKGT CIGKAPPSHQ HLCNSTMVYK QASENQYLVP
QCTRGSRNKL TLTEVSGKGT CIGKAPPSHQ HLCNSTMVYE QASENQYLVP
QCTRGSRNKL TLTEVSGKGT CIGKAPPSHQ HLCNSTMVYK QASENQYLVP
QCTRGSRNKL TLTEVSGKGT CIGKAPPSHQ HLCNSTMVYK QASENQYLVP
QCTRGSRNKL TLTEVSGKGT CIGKAPPSHQ HLCNSTMVYK QASENQYLVP
QCTRGSRNKL TLTEVSGKGT CIGKAPPSHQ HLCNSTMVYK QASENQYLVP
QCTRGSRNKL TLTEVSGKGT CIGKAPPSHQ HLCNSTMVYK QASENQYLVP

QCTRGSRNKL TLTEVSGKGT CIGKAPPSHQ HLCNSTMVYK QASENQYLVP
QCTWGSRNKL TLTEVSGKGT CIGKAPPSHQ HLCNSTMVYE QASENQYLVP
QCTWGSRNKL TLTEVSGKGT CIGKAPPSHQ HLCIVLWFYE QASENQYLVP
QCTRGSRXKL TXTEVSGKGT CIGKAPPSHQ HLCNSTMVYK QASENQYLVP
QCTRGSRNKL TLTEVSGKGT CIGKAPPSHQ HLCYSTVVYE QASENQYLVP
QCSWGTKNKL TPPEVSGRGT CIEKPPTSHQ HLCNITLTYN QTSDNQYLVP
RCPWGTKNKL TLPEVSGRGT CIK--PPSHQ HLCNVTLTYS QTSDNQYLVP
QCSWGIKNKL TFPDISGSGT CIGSAPPSHQ HLCNNTLIYN QTSDNQYLMP
QCRWGGKGKL TLTEVSGLGL CIGKVPPTHQ HLCNLTIPLN ASHTHKYLLP
RCRWGTQGKL TLTEVSGHGL CIGKVPFTHQ HLCNQTLSIN SSGDHQYLLP
QCLWGENRKL TLTAVSGNGL CLGQVPQDKW HLCNQTQNIR PNKGGQYLVP
LCTWGTQNKL TLTEVTGKGT CLGNVPADKK HLCNQTLTSP HTSTTYYLVP
LCTWGTQNKL TLTEVTGKGT CLGNVPADKK QLCNQTLTSP HTSTTYYLVP
LCTWGTQNKL TLTEVTGKGT CLGNVPTSRK HLCNQTLASP HTSTTYYLVP
LCTWGTQNKL TLTEVTGKGT CLGNVPTSRK HLCNQTLASP HTSTTYYLVP
LCTWGTQNKL TLTEVTGKGT CLGNVPTSRK HLCNQTLASP HTSTTYYLVP
LCTWGTQNKL TLTEVTGKGT CLGNVPTSRE HLCNQTLASP HTSTTYYLVP
LCTWGTQNKL TLTEVTGKGT CLGNVLTSRK HLCNQTLASP HSSTSYYLVP
HCRWKQESKL TLSSVFGNGT CVGTPPQSHR HLCHD--FS- LPSGSGYLVP
NCSVASQHKL TLSEVTGRGL CIGTVPKTHQ ALCNTTLKT- -NKGSYYLVA
NCSVASQHKL TLSEVTGRGL CIGTVPKTHQ ALCNTTLKA- -GKGSYYLVA
NCSVASQHKL TLSEVTGQGL CIGAVPKTHQ ALCNTTQTS- -SRGSYYLVA
NCSVASQHKL TLSEVTGQGL CVGAVPKTHQ ALCNTTQKA- -SDGSYYLAA
SCLSTPQHKL TISEVSGQGL CIGTVPKTHQ ALCNETQQG- -HTGAHYLAA
QCSWGDRIGL TLQLVSGNGT CLGKVPQAKQ SLCASIDSSS WKSDTKWLIP
NCSLRVQGSV DVNCYAGEAD NRTGIPVGYV HFTNCTSIQE VSNETSHLCP

GYDRWWACNT GLTPCVSTLV FNQTKDFCVM VQIVPRVYYY PEKAVLDEYD
GYDRWWACNT GLTPCVSTLV FNQTKDFCVM VQIVPRVYYY PEKAVLDEYD
GYDRWWACNT GLTPCVSTLV FNQTKDFCVM VQIVPRVYYY PEKAVLDEYD
GYDRWWACNT GLTPCVSTLV FNQTKDFCVM VQIVPRVYYY PEKAVLDEYD
GYDRWWACNT GLTPCVSTLV FNQTKDFCVM VQIVPRVYYY PEKAVLDEYD
GYDRWWACNT GLTPCVSTLV FNQTKDFCVM VQIVPRVYYY PEKAVLDEYD
GYDRWWACNT GLTPCVSTLV FNQTKDFCVM VQIVPRVYYY PEKAVLDEYD
GYDRWWACNT GLTPCVSTLV FNQTKDFCVM VQIVPRVYYY PEKAVLDEYD
GYDRWWACNT GLTPCVSTLV FNQTKDFCVM VQIVPRVYYY PEKAVLDEYD
GYDRWWACNT GLTPCVSTLV FNQTKDFCVM VQIVPRVYYY PEKAVLDEYD
GYDRWWACNT GLTPCVSTLV FNQTKDFCVM VQIVPRVYYY PEKAVLDEYD
GYDRWWACNT GLTPCVSTLV FNQTKDFCFM VQIVPRVYYY PEKAVLDEYD
GYDRWWACNT GLTPCVSTLV FNQTKDFCVM VQIVPRVYYY PEKAVLDEYD
GYDRWWACNT GLTPCVSTLV FNQTKDFCVM VQIVPRVYYY PEKAVLDEYD
GYDRWWACNT GLTPCVSTLV FNQTKDFCVM VQIVPRVYYY PEKAVLDEYD
GYDRWWACNT GLTPCVSTLV FNQTKDFCVM VQIVPRVYYY PEKAVLDEYN
GYDRWWACNT GLTPCVSTLV FNQTKDFCVM VQIVPRVYYY PEKAVLDEYD

GYDRWWACNT GLTPCVSTLV FNQTKDFCVM VQIVPRVYYY PEKAVLDEYD
GYDRWWACNT GLTPCVSTLV FNQTKDFCVM VQIVPRVYYY PEKAVLDEYD
GYDRWWACNT GLTPCVSTLV FNQTKDFCVM VQIVPRVYYY PEKAVLNKYD
GYDRWWACNT GLTPCVSTLV FNQTKDFCFM VQIVPRVYYY PKKAVLDEYD
GYDRWWTCNT GLTPCVSTLV FNQTKDFCVM VQIVPRVYYY PEKTVLDKYD
GT---WLQVV GMTPCVSTLV FNQTKDFCVM VQIVPRVYYY PKKAVLDEYG
GT---WLQVV GMTPCFSTLV FNQTKDFCVM VQIVPRVYYY PKKAVLDEYG
RLDRWWTCNT GLTPCVSTLV FNQTKDFCVM VQIVPRVYYY PEKTVLDKYD
GYDRWWACNT GLTPCVSTLV FDQTKGFCVM VQIVPRVYYH PEKAALDKYD
GYDRWWACNT GLTPCVSTLV FDQTKGFCVM VQIVPRVYYH PEKAALDKYD
GYDRWWACNT GLTPCVSTLV FDQTKGFCVM VQIVPRVYYH PEKAALDKYD
GYDRWWACNT GLTPCVSTLV FDQTKGFCVM VQIVPRVYYH PEKAALDKYD
GYDRWWACNT GLTPCVSTLV FDQTKGFCVM VQIVPRVYYH PEKAALDKYD
GYDRWWACNT GLTPCVSTLV FDQTKGFCVM VQIVPRVYYH PEKAALDKYD
GYDRWWACNT GLTPCVSTLV FDQTKGFCVM VQIVPRVYYH PEKAALDKYD
GYDRWWACNT GLTPCVSTLV FDQTKGFCVM VQIVPRVYYH PEKAALDKYD
GYDRWWACNT GLTPCVSTLV FDQTKGFCVM VQIVPRVYYH PEKAVLDKYD
GYDRWWACNT GLTPCVSTLV FNQTKDFCVM VQIVPRVYYH PEKAALDKYD
GYDRWWACNT GLTPCVSTLV FDQTKGFCVM VQIVPRVYYH PEKAALDKYD
GYDRWWACNT GLTPCVSTLV FDQTKGFCVM VQIVPRVYYH PEKAALDKYD
GYDRWWACNT GLTPCVSTLV FNQTKDFCIM VQIVPRVYYY PEKAILDEYD
GYDRWWACNT GLTPCVSTLV FNQTKDFCIM VQIVPRVYYY PEKAILDEYD
GYDRWWACNT GLTPCVSTLV FNQTKDFCIM VQIVPRVYYY PEKAILDEYD
GYDRWWACNT GLTPCVSTLV FNQTKDFCIM VQIVPRVYYY PEKAILDEYD
GYDRWWACNT GLTPCVSTLV FNQTKDFCIM VQIVPRVYYY PEKAILDEYD
GYDRWWACNT GLTPCVSTLV FNQTKDFCIM VQIVPRVYYY PEKAILDEYD
GYDRWWACNT GLTPCVSTLV FNQTKDFCIM VQIVPRVYYY PEKAILDEYD
GYDRWWACNT GLTPCVSTLV FNQTKDFCIM VQIVPRVYYY PEKAILDEYD
GYDRWWACNT GLTPCVSTLV FNQTKDFCIM VQIVPRVYYY PEKAILDEYD
GYDRWWACNT GLTPCVSTLV FNQTKDFCIM VQIVPRVYYY PEKAILDEYD
GYDRWWACNT GLTPCVSTLV FNQTKDFCVM VQIVPRVYYY PEKTVLDEYD
GYDRWWACNT GLTPCVSTLV FNQTKDFCVM VQIVPRVYYY PEKTVLDEYD
GYDRWWACNT GLTPCVSTLV FNETKDFCVM VQIVPRVYYY PEKTVLDEYD
GYDRWWACNT GLTPCVSTLV FNKTKDFCVM VQIVPRVYYY PEKTVLDEYD
GYDRWWACNT RLTPCVSTLV FNQTKDFYIV VQIVPRVYYY PEKTILDEYN
GYDRWWACNT RLTPCVSTLV FNQTKDFYIV VQIVPRVYYY PEKTILDEYN
GYDRWWACNT GLTPCVSTLV FNQTKDFYIM VQIVPRVYYY PKKTILDEYD
GYDRWWACNT GLTPCVSTLV FNQTKDFYIM VQIVPRVYYY PKKTILDEYD
GYNRWWACNT GLTPCVSTSV FNQSKDFCVM VQIVPRVYYH PEEVVLDEYD
GYNRWWACNT GLTPCVSTSV FNQSKDFCVM VQIVPRVYYH PEEVVLDEYD
GYNRWWACNT GLTPCVSTSV FNQSKDFCVM VQIVPRVYYH PEEVVLDEYD
GYNRWWACNT GLTPCVSTSV FNQSKDFCVM VQIVPRVYYH PEEVVLDEYD
GYNRWWACNT GLTPCVSTSV FNQSKDFCVM VQIVPRVYYH PEEVVLDEYD
GYNRWWACNT GLTPCVSTSV FNQSKDFCVM VQIIPRVYYH PEEVVLDEYD

GYNRWWACNT GLTPCVSTSV FNQSKDFCVM VQIVPRVYYH PEEVVLDEYD
GYNRWWACNT GLTPCVSTSV FNQSKDFCVM VQIVPRVYYH PEEVVLDEYD
GYNRWWACNT GLTPCVSTSV FNQSKDFCVM VQIVPRVYYH PEEVVLDEYD
GYNRWWACNT GLTPCVSTSV FNQSKDFCVM VQIVPRVYYH PEEVVLDEYD
GCNRWWACNT GLTPCVSTSV FNQSKDFCVM VQIVPRVYYH PEEVVLDEYD
GYNRWWACNT GLTPCVSTSV FNQSKNFCVM VQIVPRVYYH PEKVVLDEYD
GYNRWWACNT GLTPCVSTSV FNQSKDFCVM VQIVPRVYYH PEEVVLDEYD
GYNRWWACNT GLTPCVSTSV FNQSKDFCVM VQIIPRVYYH PEEVVLDEYD
GYNRWWACNT GLTPCVSTSV FNQSKDFCVM VQIVPRVYYH PEEVVLDEYD
GYNRWWACNT GLTPCVSTSV FNQSKDFCVM VQIIPRVYYH PEEVVLDEYD
GYNRWWACNT GLTPCVSTSV FNQSKDFCVM VQIIPRVYYH PEEVVLDEYD
GYNRWWAYNT GLTPCVSTSV FNQSKDFCVM VQIVPRVYYH PEKVVLDEYD
GYNRWWACNT GLTPCVSTTV FNQSKDFCVM VQLIPQVYYH PEEVVIDEYD
GYNRWWACNT GLTPCVSTTV FNQSKDFCVM VQLIPQVYYH PEEVVIDEYD
GYNRWWACNT GLTPCVSTTV FNQSKDFCVM VQLIPQVYYH PEEVVIDEYD
GYNRWWACNT GLTPCVSTTV FNQSKDFCVM VQLIPQVYYH PEEVVIDEYD
GYNRWWACNT GLTPCVSTTV FNQSKDFCVM VQLIPQVYYH PEEVVIDEYD
GYNRWWACNT GLTPCVSTTV FNQSKDFCVM VQLIPQVYYH PEEVVIDEYD
GYNRWWACNT GLTPCVSTTV FNQSKDFCVM VQLIPQVYYH PEEVVIDEYD
GYNRWWACNT GLTPCVSTTV FNQSKDFCVM VQLVPRVYYH PEEVVIDEYD
GYNRWWACNT GLTPCVSTTV FNQSKDFCVM VQLVPRVYYH PEEVVIDEYD
GYN-WWACNT GLTPCVSTTV FNQSKDFCVM VQLVPWVHYH PEEVVIDEYD
GYN-WWACNT GLTPCVSTTV FNQSKDFCVM VQLVPWVHYH PEEVVIDEYD
GYN-WWACNT GLTPCVSTTV FNQSKDFCVM VQLVPWVHYH PEEVVIDEYD
GYN-WWACNT GLTPCVSTTV FNQSKDFCVM VQLVPWVHYH PEEVVIDEYD
GYNRWWACNT GLTPRVSTTV FNQSKDFCVM VQ-------- PEEVVIDKYD
GYNRWWACNT GLTPCVSTSV FNQSKDFCVM VQ-------- PEEVVIDKYD
GYNRWWACNT GLTPCVSTTV FNQSKDFCVM VQLVPRVYYH PEEVVIDEYD
GYNRWWACNT GLTPCVSTTV FNQSKDFCVM VQLVPRVYYH PEEVVIDEYD
GYNRWWACNT GLTPRVSTTV FNQSKDFCVM VQLVPRVYYH PEEVVIDEYD
GYNRWWACNT GLTPRVSTTV FNQSKDFCVM VQLVPRVYYH PEEVVIDEYD
GYNRWWACNT GLTPRVSTTV FNQSKDFCVM VQLVPRVYYH PEEVVIDEYD
GYNRWWACNT GLTPRVSTTV FNQSKDFCVM VQLVPRVYYH PEEVVIDEYD
GYNRWWACNT GLTPRVSTTV FKQSKDFCVM VQLVPRVYYH PEEVVIDEYD
GYNRWWACNT GLTPRVSTTV FNQSKDFFVM VQLVPRLYYH PEEVVIDEYD
GYNRWWACNT GLTPCVSTSV FNQSKDFCVM VQLVPRVYYH PEEVVIDEYD
GYNRWWACNT GLTPCVSTSV FNQSKDFCVM VQLVPRVYYH PEEVVIDEYD
GYNRWWACNT GLTPCVSTTV FNQSKDFCVM VQLVPRVYYH PEEVVIDXYD
GYNRWWACNT GLTPCVSTTV FNQSKDFCVM VQ-------- PEEVVIDKYD
GYNRWWACDT GLTPCVSTSV FNPSKNFCIM VQLVPRVYYH PEEVVIDEYD
EYNRWWACDT GLTPCVSTSV FNPSKNFCIM VQLVPWVYYH PEEVVIDEYD
GYNRWWACDT GLTPCVSPSV FKQSKNLYIM VRLVPQVYYH PEEVVVDEYD
SNHSWWACNS GLTPCLSTSV FNQSNDFCIQ IQLVPRIYYH PDGTLLQAYE
SNHSWWACST GLTPCLSTSV FNQTRDFCIQ VQLIPRIYYY PEEVLLQAYD

PIDTVWACNT GLTPCISMSV FNSSKDFCIL VQLIPRLLYH DDSSFLDKFE
PLEGWWACST GLTPCVATSV FNNSKDFCIL IQLVPRIIYH DSKSFEDQFD
PLEGWWAYST RLTPCVATSV FNNSKDFCIL IQLVPRIIYH DSKSLEDKFD
PLEGWWACST GLTPCVATSV FNSSRDFCIL VQLVPRVIYH DSKSFEAQFD
PLEGWWACST GLTPCVATSV FNNSKDFCIL VQLVPRVIYH DSKSFEDQFD
PLEGWWACST GLTPCVATSV FNSSKDFCIL VQLVPRVIYH SKINLIPKYD
PLEGWWACST GLTPCVATSV FNSSRDFCIL VQLVPRVIYH DSKSFEDQFD
PLEGWWACST GLTPCVATSV FNSSKDFCIL VQLVPRVIFH DSKSFEDQFD
PQDGWWACNT GLTPCVSLEV LNTSADFCVL VQLVPRLIYH SDPSFLDEYE
PAGTTWACNT GLTPCLSATV LNRTTDYCVL VELWPRVTYH PPSYVYSQFE
PTGTMWACNT GLTPCLSATV LNRTTDYCVL VELWPRVTYH PPSYVYSQFE
PTGTMWACST GLTPCISTTI LNLTTDYCVL VELWPRVTYH SPSYVYGLFE
PAGTIWACNT GLTPCLSTTV LNLTTDYCVL VELWPKVTYH SPGYVYDQFE
PNGAYWACNT GLTPCISMAV LNWTSDFCVL IELWPRVTYH QPEYVYTHFA
RTDGWWICST GLTPCLSTSV FNAANEFCVL VTVLPRILYH PEESMYSHWD
PPGHVFVCGN NMA---YTAL PNKWIGLCIL ASIVPSIISG EEPIPLPSIE

YRYNQPKREP ISLTLAVMLG LGVAAGVGTG TAALITGPQQ LEKGLSDLHR
YRYNRPKREP ISLTLAVMLG LGVAAGVGTG TAALITGPQQ LEKGLSNLHR
YRYNRPKREP ISLTLAVMLG LGVAAGVGTG TAALITGPQQ LEKGLSNLHR
YRYNRPKREP ISLTLAVMLG LGVAAGVGTG TAALITGPQQ LEKGLSNLHR
YRYNRPKREP ISLTLAVMLG LGVAAGVGTG TAALITGPQQ LEKGLSNLHR
YRYNRPKREP ISLTLAVMLG LGMAAGVGTG TAALITGPQQ LEKGLSNLHR
YRYNRPKREP ISLTLAVMLG LGVAAGVGTG TAALITGPQQ LEKGLSNLHR
YRYNRPKREP ISLTLAVMLG LGVAAGVGTG TAALITGPQQ LEKGLSNLHR
YRYNRPKREP ISLTLAVMLG LGVAAGVGTG TAALITGPQQ LEKGLSNLHR
YRYNRPKREP ISLTLAVMLG LGVAAGVGTG TAALITGPQQ LEKGLSNLHR
YRYNRPKREP ISLTLAVMLG LGVAAGVGTG TAALITGPQQ LEKGLSNLHR
YRYNRPKREP ISLTLAVMLG LGVAAGVGTG TAALITGPQQ LEKGLSNLHR
YRYNRPKREP ISLTLAVMLG LGMAAGVGTG TAALITGPQQ LEKGLSNLHR
YRYNRPKREP ISLTLAVMLG LGVAAGVGTG TAALITGPQQ LEKGLSDLHR
YRYNWPKREP ISLTLAVMLG LGVAAGVGTG TAALITGPQQ LEKGLSNLHR
YRYNRPKREP ISLTLAVMLG LGMATGVGTG TAALITGPQQ LEKGLSNLHR
YRYNRPKREP ISLTLAVMLG LGVAAGVGTG TAALITGPQQ LEKGLSDLHR
YRYNRPKREP ISLTLAVMLG LGVAAGVGTG TAALITGPQQ LEKGLSDLHR
YRYNRPKREP ISLTLAVMLG LGVATGVGTG TAALITGPQQ LEKGLSNLHR
YRYNRPKREP ISLTLAVMLG LGVAAGVGTG TAALITGPQQ LEKELSNLHR
YRYNRPKREP ISLTLAVMLG LGVAAGMGTG TAALITGPQQ LEKGLSDLHR
YRYNRPKKEP ISLTLAVMLG LGVAAGVGTG TAALITGPQQ LEKELNNLHR
YRYNRPKREP ISLTLAVMLG LGVAAGVGTG MAALITGPQQ LEKGLSDLHR
YRYNRPKREP ISLTLAVMLG LGVAAGVGTG TAALITGPQQ LEKGISDLHR
YRYNRPKREP ISLTLAVMLG LGVAAGVGTG TAALITGPQQ LEKGISDLHR
YKYNRPKRKP ISLTLAAMLG LGVAAGVGTG TAALITGPRQ LEKGLGELHR
YKYNRPKRKP ISLTLAAMLG LGVAAGVGTG TAALITGPRQ LEKGLGELHR

YKYNRPKRKP ISLTLAAMLG LGVAAGVGTG TAALITGPRQ LEKGLGELHR
YKYNRPKRKP ISLTLAAMLG LGVAAGVGTG TAALITGPRQ LEKGLGELHR
YKYNRPKRKP ISLTLAAMLG LGVAAGVGTG TAALITGPRQ LEKGLGELHR
YKYNRPKRKP ISLTLAAMLG LGVAAGVGTG TAALITGPRQ LEKGLGELHR
YKYNRPKRKP ISLTLAAMLG LGVAAGVGTG TAALITGPRQ LEKGLGELHR
YKYNRPKRKP ISLTLAAMLG LGVAAGVGTG TAALITGPRQ LEKGLGELHR
YKYNRPKRKP ISLTLAAMLG LGVAAGVGTG TAALITGPRQ LEKGLGELHR
YKYNRPKRKP ISLTLAAMLG LGVAAGVGTG TAALITGPRQ LEKGLGELHR
YKYNRPKRKP ISLTLAAMLG LGVAAGVGTG TAALITGPRQ LEKGLGELHR
YKYNRPKRKP ISLTLAAMLG LGVAAGVGTG TAALXTGPRQ LEKGLGELHR
YRNHRQKREP ISLTLAVMLG LGVAAGVGTG TAALVTGPQQ LETGLSNLHR
YRNHRQKREP ISLTLAVMLG LGVAAGVGTG TAALVTGPQQ LETGLSNLHR
YRNHRQKREP ISLTLAVMLG LGVAAGVGTG TAALVTGPQQ LETGLSNLHR
YRNHRQKREP ISLTLAVMLG LGVAAGVGTG TAALVTGPQQ LETGLSNLHR
YRNHRQKREP ISLTLAVMLG LGVAAGVGTG TAALVTGPQQ LETGLSNLHR
YRNHRQKREP ISLTLAVMLG LGVAAGVGTG TAALVTGPQQ LETGLSNLHR
YRNHRQKREP ISLTLAVMLG LGVAAGVGTG TAALVTGPQQ LETGLSNLHR
YRNHRQKREP ISLTLAVMLG LGMVAGVGTG TAALVTGPQQ LETGLSNLHR
YRNHRQKREP ISLTLAVMLG LGMVAGVGTG TAALVTGPQQ LETGLSNLHR
YRNHRQKREP ISLTLAVMLG LGMVAGVGTG TAALVTGPQQ LETGLSNLHR
YKYHRQKREP ISLTLAVMLG LGVAAGVGTG TAALVTGPQQ LETGLSNLHR
YKYHRQKREP ISLTLAVMLG LGVAAGVGTG TAALVTGPQQ LETGLSNLHR
YKYHRQKREP ISLTLAVMLG LGVAAGVGTG TAALVTGPQQ LEKGLSNLHR
YRYHRQKREP ISLTLTVMLG LGVAAGVGTG TAALVTGPQQ LETGLSNLHR
YRNHRQKRKP ISLTLAVMLG LGVAAGVGTG TAALVTGPQQ LETGLNNLHQ
YRNHRQKRKP ISLTLAVMLG LGVAAGVGTG TAALVTGPQQ LETGLNNLHQ
YRNHRQKKKP ISLTLAVMLG LGVITGVRTG TAALVTGPQQ LETGLSNLHQ
YRNHRQKKKP ISLTLAVMLG LGVITGVRTG TAALVTGPQQ LETGLSNLHQ
YRYNRPKREP VSLTLAVMLG LGTAVGVGTG TAALITGPQQ LEKGLGELHA
YRYNRPKREP VSLTLAVMLG LGTAVGVGTG TAALITGPQQ LEKGLGELHA
YRYNRPKREP VSLTLAVMLG LGTAVGVGTG TAALITGPQQ LEKGLGELHA
YRYNRPKREP VSLTLAVMLG LGTAVGVGTG TAALITGPQQ LEKGLGELHA
YRYNRPKREP VSLTLAVMLG LGTAVGVGTG TAALITGPQQ LEKGLGELHA
YRYNRPKREP VSLTLAVMLG LGTAVGVGTG TAALITGPQQ LEKGLGELHA
YRYNRPKREP VSLTLAVMLG LGTAVGVGTG TAALITGPQQ LEKGLGELHA
YRYNRPKREP VSLTLAVMLG LGTAVGVGTG TAALITGPQQ LEKGLGELHA
YRYNRPKREP VSLTLAVMLG LGTAVGVGTG TTALITGPQQ LEKGLGELHA
YRYNRPKREP VSLTLAVMLG LGTAVGIGTG TAALITGPQQ LEKGLGELHA
YRYNRPKREP VSLTLAVMLG LGTAVGVGTG TAALITGPQQ LEKGLGELHA
YRYNRPKREP VSLTLAVMLG LGTAVGVGTG TAALITGPQQ LEKGLGELHA
YRYNQPKREP ISLTLAVMLG LGTAVGVGTG TAALITGPQQ LEKGLGELHA
YRYNRPKREP VSLTLAVMLG LGTAVGVGTG TAALITGPQQ LEKGLGELHA
YRYNRPKREP VSLTLAVMLG LGTAVGVGTG TAALITGPQQ LEKGLGELHA
YRYNRPKREP VSLTLAVMLG LGTAVGVGTG TAALITGPQQ LEKGLGELHA

```
YRYNRPKREP VSLTLAVMLG LGTAVGVGTG TAALITGPQQ LEKGLGELHA
YRYNRPKREP VSLTLAVMLG LGTAVGVGTG TAALITGPQQ LEKGLGELHA
YRPTRSKREP VTLTLAIILG LGMA-GVGTG TAALITGPQQ LEKGLGELHA
YRPTRSKREP VTLTLAIILG LGMA-GVGTG TAALITGPQQ LEKGLGELHA
YRPTRSKREP VTLTLAIILG LGMA-GVGTG TAALITGPQQ LEKGLGELHA
YRPTRSKREP VTLTLAIILG LGMA-GVGTG TAALITGPQQ LEKGLGELHA
YRPTRSKREP VTLTLAIILG LGMA-GVGTG TAALITGPQQ LEKGLGELHA
YRPTRSKREP VTLTLAIILG LGMA-GVGTG TAALITGPQQ LEKGLGELHA
YRPTRSKREP VTLTLAIILG LGMA-GVGTG TAALITGPQQ LEKGLGELHA
YRPTRSKREP VTLTLAVILG LGMA-GVGTG TAALITEPQQ LEKGLGELHA
YRPTRSKREP VTLTLAVILG LGMA-GVGTG TAALITGPQQ LEKGLGELHA
YRPTRSKREP VTL------- ---------- -------PQQ LETGLGELHA
YRPTRSKREP VTL------- ---------- -------PQQ LEKGLGELHA
YRPTRSKREP VTL------- ---------- -------PQQ LEKGLGELHA
YRPTRSKREP VTL------- ---------- -------PQQ LEKGLGELHA
HRPTRSKREP MSLTLAVMLG LGVTAGVGTG AAALITGPQQ LEKGLGAPHA
HRPTRSKREP MSLTLAVMLG LGVTAGVGTG AAALITGPQQ LEKGLGAPHA
HRPTRSKREP VSLTLAVMLG LGVTAGVGTG AAALITGPQQ LEKGLGAPHA
HRPTRSKREP VSLTLAVMLG LGVTAGVGTG AAALITGPQQ LEKGLGAPHA
HRPTRSKREP VSLTLAVMLG LGVTAGVGTG AAALITGPQQ LEKGLGAPHA
HRPTRSKREP VSLTLAVMLG LGVTAGVGTG AAALITGPQQ LEKGLGAPHA
HRPTRSKREP VSLTLAVMLG LGVTAGVGTG AAALITGPQQ LEKGLGAPHA
HRPTRSKREP VSLTLAVMLG LGVTAGVGTG AAALITGPQQ LEKGLGAPHA
HRPTRSKREP VSLTLAVMLG LGVTAGVGTG AAALITGPQQ LEKGLGAPHA
HRPTRSKREP VSLTLAVMLG LGVTAGVGTG AAALITGPQQ LEKGLGAPHA
HRPTRSKREP VSLTLAVMLG LGVTAGVGTG AAALITGPQQ LEKGLGAPHA
HRPTRSKREP VSLTLAVMLG LGVTAGVGTG AAALITGPQQ LEKGLGAPHA
HRPTRSKREP VSLTLAVMLG LGVTAGVGTG AAALITGPQQ LEKGLGAPHA
HRPTRSKREP MSLTLAVMLG LGVTAGVGTG AAALITGPQQ LEKGLGAPHA
YHSPRSKREP ------VMLG LGIAAGVGSG TTALVMGPQQ LERGLGKLHA
YHPTRSKREP VTLTLAVILG LGIAAGVGTG TTALVTEPQQ LERGVGELHA
Y-----KREP VTLTLAVMLG LGIATGVGTG TTALITGPQQ LEKGLGELHA
SPHPRNKREP VSLTLAVLLG LGVAAGIGTG STALIKGPID LQQGLTSLQI
NSHPRTKREA VSLTLAVLLG LGITAGIGTG STALIKGPID LQQGLTSLQI
HR-VRWKREP ITLTLAVLLG LGVAAGVGTG TAALIQTPRY ----FEELRT
WRIYRVRREP VSMALAILLG AGVVARMGTG TTALIQGSQR ----YEKLRA
SRIYRVRREP VSVTLAILLG AGVVA-MGTG TTALIQGSQR ----YEKLRA
SKIHRTRREP VSMTLAVLLG LGVAAGVRTG TTALIQGPLH ----YEKLRA
SKIHRIQRDP VSMTLAVLLG LGVAAGVGTG TTALIQGPHH ----YEKLRA
SKIHRTQREP VSMTLAVLLG LGVAAGVGTR TTALIQGSHH ----YEKFRA
PKIHRTRREP VSMTLAVLLG SGVAAKVGTG TTALIQGPHH ----YEKLRA
SKIHRTRREP VSMTLAVLLG LGVAARIETG TTALIQGPHH ----YEKLRA
GR----KKKK ITLTLAMLLG LGITAGIGTG TTALIQQPQY ----YASLRQ
-KSYRHKREP VSLTLALLLG GGIAAGVGTG TTALVATQQ- ----FQQLHA
```

-KSHRHKREP VSLTLALLLG GGIAAGVGTG TTALVATQQ- ----FQQLHA
-RSNRHKREP VSLTLALLLG GGIAAGIGTG TTALMATQQ- ----FQQLQA
-RKTKYKREP VSLTLALLLG GGIAAGVGTG TTALVATKQ- ----FEQLQA
-KAVRFRREP ISLTVALMLG GGIAAGVGTG TKALLETAQ- ----FRQLQM
STSTRSKREP IALTIATLFS LRI-AGAGTG IASLA-T-Q- -QSGITSLRA
YTAGRHKRAV QFIPLPV--G LGITGATLTG GTGLGVSVHT YH----KLSN

IVTEDLQALE KSVSNLEESL TSLSEVVLQN RRGLDLLFLK EGGLCVALKE
IVTEDLQALE KSVSNLEESL TSLSEVVLQN RRGLDLLFLK EGGLCVALKE
IVTEDLQALE KSVSNLEESL TSLSEVVLQN RRGLDLLFLK EGGLCVALKE
IVTEDLQALE KSVSNLEESL TSLSEVVLQN RRGLDLLFLK EGGLCVALKE
IVTEDLQALE KSVSNLEESL TSLSEVVLQN RRGLDLLFLK EGGLCVALKE
IVTEDLQALE KSVSNLEESL TSLSEVVLQN RRGLDLLFLK EGGLCVALKE
IVTEDLQALE KSVSNLEESL TSLSEVVLQN RRGLDLLFLK EGGLCVALKE
IVTEDLQALE KSVSNLEESL TSLSEVVLQN RRGLDLLFLK EGGLCVALKE
IVTEDLQALE KSVSNLEESL TSLSEVVLQN RRGLDLLFLK EGGLCVALKE
IVTEDLQALE KSVSNLEESL TSLSEVVLQN RRGLDLLFLK EGGLCVALKE
IVTEDLQALE KSVSNLEESL TSLSEVVLQN RRGLDLLFLK EGGLCVALKE
IVTEDLQALE KSVSNLEESL TSLSEVVLQN RRGLDLLFLK EGGLCVALKE
IVTEDLQALE KSVSNLEESL TSLSEVVLQN RRGLDLLFLK EGGLYVALKE
IVTEDLQALE KSVSNLEESL TSLSEVVLQN RRGLDLLFLK EGGLCVALKE
IVTEDLQALE KSVSNLEESL TSLSEVVLQN RRGLDLLFLK EGGLCVALKE
IVTEDLQALE KSVSNLEESL TSLSEVVLQN RRGLDLLFLK EEGLCVALKE
IVTEDLQALE KSVSNLEESL TSLSEVVLQN RRGLDLLFLK EGGLCVALKE
IVTEDLQALE KSVSNLEESL TSLSEVVLQN RRGLDLLFLK EGGLCVALKE
IVTEDLQALE KSVSNLEESL TSLSEMVLQN RRGLDLLFLK KGGLCVALKE
IVTEDLQALK KSVSNLEESL TSLSKVVLQN RRGLDLLFLK KGELCVALKE
IVMEDLQALE KSVSNLEESL TSLSEVVLQN RRGLDLLFLK EGGLCVALKE
IVTEDLQALE KSVSNLKKSL TSLSEVVLQN RRGLDLLFLK EGGLCVALKE
IVMEDL-ALE KSVCNQEESL TSLSEVVLQN RRGLDLLFLK EGGLCVALKE
IVTKDLQALK QSVSNLEESL TSLSEVVLQN RRGLDLLFLK ESGLCVALKE
IVTKDLQALK QSVSNLEESL TSLSEVVLQN RRGLDLLFLK ESGLCVALKE
IITEDLQALE KSVSNLEEFL TSLSEVVLQN RRGLDLLFLK KGGLCVALKE
IITEDLQALE KSVSNLEEFL TSLSEVVLQN RRGLDLLFLK KGGLCVALKE
IITEDLQALE KSVSNLEEFL TSLSEVVLQN RRGLDLLFLK KGGLCVALKE
IITEDLQALE KSVSNLEEFL TSLSEVVLQN RRGLDLLFLK KGGLCVALKE
IITEDLQALE KSVSNLEEFL TSLSEVVLQN RRGLDLLFLK KGGLCVALKE
IITEDLQALE KSVSNLEEFL TSLSEVVLQN RRGLDLLFLK KGGLCVALKE
IITEDLQALE KSVSNLEEFL TSLSEVVLQN RRGLDLLFLK KGGLCVALKE
IITEDLQALE KSVSNLEEFL TSLSEVVLQN RRGLDLLFLK KGGLCVALKE
IITEDLQALE KSVSNLEEFL TSLSEVVLQN RRGLDLLFLK KGGLCVALKE
IITEDLQALE KSVSNLEEFL TSLSEVVLQN RRGLDLLFLK KGGLCVALKE
IITEDLQALE KSVSNLEEFL TSLSEVVLQN RRGLDLLFLK KGGLCVALKE
IITEDLQALE KSVSNLEEFL TSLSEVVLQN RRGLDLLFLK KGGLCVALKE

IVTEDLQALE KSVSNLEESL TSLSEVVLQN RRGLDLLFLK EGGLCVALKE
IVTEDLQALE KSVSNLEESL TSLSEVVLQN RRGLDLLFLK EGGLCVALKE
IVTEDLQALE KSVSNLEESL TSLSEVVLQN RRGLDLLFLK EGGLCVALKE
IVTEDLQALE KSVSNLEESL TSLSEVVLQN RRGLDLLFLK EGGLCVALKE
IVTEDLQALE KSVSNLEESL TSLSEVVLQN RRGLDLLFLK EGGLCVALKE
IVTEDLQALE KSVSNLEESL TSLSEVVLQN RRGLDLLFLK EGGLCVALKE
IVTEDLQALE KSVSNLEESL TSLSEVVLQN RRGLDLLFLK EGGLCVALKE
IVTEDLQALE KSVSNLEESL TSLSEVVLQN RRGLDLLFLK EGGLCVALKE
IVTEDLQALE KSVSNLEESL TSLSEVVLQN RRGLDLLFLK EGGLCVALKE
IVTEDLQALE KSVSNLEESL TSLSEVVLQN RRGLDLLFLK EGGLCVALKE
IVTEDLQALE KSVSNLEESL TSLSEVVLQN RRGLDLLFLK ERGLCVALKE
IVTEDLQALE KSVSNLEESL TSLSEVVLQN RRGLDLLFLK ERGLCVALKE
IVTEDLQALE KSVSNLEESL TSLSEVVLQN RRGLDLLFLK EGGLCVALKE
IVTEDLQALE KSVSNLEESL TSLSEVVLQN RRGLDLLFLK EGGLCVALKE
IVTEDLQALE KSVSNLEESL TSLSEVVLQN RRGLDLLFLK EGGLCVALKE
IVTEDLQALE KSVSNLEKSL TSLSEVVLQN RRGLDLLFLK EGRLCVALKE
IVTGNLQALK KSVSNLEKSL TSLSEVVLQN KKGLDLLFLK KRRLCVALKE
IVTGNLQALK KSVSNLEKSL TSLSEVVLQN KKGLDLLFLK KRRLCVALKE
AMTEDLRALE ESVSNLEESL TSLSEVVLQN RRGLDLLFLR EGGLCAALKE
AMTEDLRALE ESVSNLEESL TSLSEVVLQN RRGLDLLFLR EGGLCAALKE
AMTEDLRALE ESVSNLEESL TSLSEVVLQN RRGLDLLFLR EGGLCAALKE
AMTEDLRALE ESVSNLEESL TSLSEVVLQN RRGLDLLFLR EGGLCAALKE
AMTEDLRALE ESVSNLEESL TSLSEVVLQN RRGLDLLFLR EGGLCAALKE
AMTEDLRALE ESVSNLEESL TSLSEVVLQN RRGLDLLFLR EGGLCAALKE
AMTEDLRALE ESVSNLEESL TSLSEVVLQN RRGLDLLFLR EGGLCAALKE
AMTEDLRALE EAVSNLEESL TSLSEVVLQN RRGLDLLFLR EGGLCAALKE
AMTEDLRALE ESVSNLEESL TSLSEVVLQN RRGLDLLFLR EGGLCAALKE
AMTEDLRALE ESVSNLEESL TSLSEVVLQN RRGLDLLFLR EGGLCAALKE
AMTEDLRALE ESVSNLEESL TSLSEVVLQN RRGLDLLFLR EGGLCAALKE
AMTEDLRALE ESVSNLEESL TSLSEVVLQN RRGLDLLFLR EGGLCAALKE
AMTEDLRALE ESVSNLEESL TSLSEVVLQN RRGLDLLFLR EGGLCAALKE
AMTEDLRALE ESVNNLEESL TSLSEVVLQN WRGLDLLFLK EGGLCAALRE
AMTEDLRALE ESVNNLEESL TSLSEVVLQN WRGLDLLFLK EGGLCAALRE
TMTEDL-ALE ESVSNLEESL TSLSEVVLQN RRGLDLLFLR EGGLCAALKE
AMTEDLRALE ESVSNLEETL TSWSEVVLQN QRGLDLLFLK EGGLCAALKE
AMTEDLRALE ESVSNLEESL TSLSEVVLQN RRGLDLLFLR EGGLCAALKE
AMTEDLRALE ESVSNLEETL TSWSEVVLQN QRGLDLLFLK EGGLCAALKE
AMTEDLRALE ESVSNLEETL TSWSEVVLQN QRGLDLLFLK EGGLCAALKE
AMTEDLRALE ESVSNLEETL TSWSEVVLQN QRGLDLLFLK EGGLCAALKE
AMTEDLRALE ESVSNLEETL TSWSEVVLQN QRGLDLLFLK EGGLCAALKE
AMTEDLRALE ESVSNLEETL TSWSEVVLQN QRGLDLLFLK EGGLCAALKE
AMTEDLRALE ESVSNLEETL TSWSEVVLQN QRGLDLLFLK EGGLCAALKE
AMTEDLRALE ESVSNLEETL TSWSEVVLQN QRGLDLLFLK EGGLCAALKE
AMTEDLRALE ESVSNLEESL TSLSEVVLQN WRGLDLVFLK EGGLCAALKE

```
AMTEDLRALE ESVSNLEESL TSLSEVVLQN RTGGDIWFLK EGGLCAALKE
AITEDLRALE ESVNNLEESL TSLSEVVLQN WRGLDLLFLK EGGLCAALRK
AITEDLRALE ESVNNLEESL TSLSEVVLQN WRGLDLLFLK EGGLCAALRE
AITEDLRALE ESVNNLEESL ISLSEVVLQN WRGLDLLFLK EGGLCAALRE
AITEDLRALE ESVNNLEESL ISLSEVVLQN WRGLDLLFLK EGGLCAALRE
AMTEDLRALE ESVSNLEESL TPLSEVVLQN RRGLDLLFLK EGGLCAALKE
AMTEDLRALE ESVSNLEESL TPLSEVVLQN RRGLDLLFLR EGGLCAALKE
AMTEDLRALE ESVSNLEESL TPLSEVVLQN RRGLDLLFLK EGGLCAALKE
AMTEDLRALE ESVSNLEESL TPLSEVVLQN RRGLDLLFLK EGGLCAALKE
AMTEDLRALE ESVSNLEESL TPLSEVVLQN RRGLDLLFLK EGGLCAALKE
AMTEDLRALE ESVSNLEESL TPLSEVVLQN RRGLDLLFLK EGGLCAALKE
AMTEDLRALE ESVSNLEESL TPLSEVVLQN RRGLDLLFLK EGGLCAALKE
AMTEDLRALE ESVSNLEESL TPLSEVVLQN RRGLDLLFLK EGGLCAALKE
AMTEDLRALE ESVSNLEESL TPLSEVVLQN RRGLDLLFLK EGGLCAALKE
AMTEDLRALE ESVSNLEESL TPLSEVVLQN RRGLDLLFLK EGGLCAALKE
AMTEDLRALE ESVSNLEESL TPLSEVVLQN RRGLDLLFLK EGGLCAALKE
AMTEDLRALE ESVSNLEESL TPLSEVVLQN RRGLDLLFLK EGGLCAALKE
AMTEDLRALE ESVSNLEESL TPLSEVVLQN RRGLDLLFLX EGGLCAALKE
AMTEDLRALE ESVSNLEESL TPLSEVVLQN RRGLDLLFLK EGGLCAALKE
VITEDLQALE KSISNLEESL TSLSEVVLQN RRGLDLLFLK EGGLCAALKK
VITEDLQALE KSVSNLEESL TSLSEVVLQN WRGLDLLFLK EGGLCAALKE
VITEDLQALQ KSISNLEETL TSLSEVVLQN WRELDLLFLK EGGLCAALKE
AMDTDLRALQ DSVSKLENSL TSLSEVVLQN RRGLDLLFLK EGGLCAALKE
AIDADLRALQ DSVSKLEDSL TSLSEVVLQN RRGLDLLFLK EGGLCAALKE
AMDTDLRAIE HSITKLEESL TSLSEVVLQN RRGLDLLFLK EGGLCAALKE
TMDEDLKTIE NSISKLEESL TSLSEVVLQN RRGLDLLFLK DGGLRAAIKE
AMDEDLKTIE NSISKLEESL TSLSEVVLQN RRGLDLLFLK DGGLCAARKE
AMDEDLKAIE NSITKLEESL TSLSEVVLQN RRGLDLLFLK DGGLCAAIKE
AMDEDLKAIE NSITKLEESL TSLSEVVLQN RRGLDLLFLK DGGLCAAIKE
AMDEDLKAIE NSITKLEESL TSLSEVVLQN RRGLDLLFLK DRGLCAAIKE
AMDEDLKAIE NS----EESL TSRSEVVLQN RRGLDLSSLK DGGLCAAIKE
AMYEDLKAIE NSITKLEESL TSLSEVVLQN RRGLDLLFLK DGGLCAAIKE
AVDIDLRALE SSITQLKESL TSLSEVVLQN RRGLDLLFLK EGGLCAALKE
AVQDDFKEVE KSITNLEKSL TSLSEVVLQN RRGLDLLFLK EGGLCAALKE
AVQDDLKEVE KSITNLEKSL TSLSEVVLQN RRGLDLLFLK EGGLCAALKE
AVQDDLREVE KSISNLEKSL TSLSEVVLQN RRGLDLLFLK EGGLCAALKE
AIHTDLGALE KSVSALEKSL TSLSEVVLQN RRGLDLLFLK EGGLCAALKE
AMHTDIQALE ESISALEKSL TSLSEVVLQN RRGLDILFLQ EGGLCAALKE
AIDEDIERLE TSISHLEKSL TSLSEVVLQN RRGLDLLFLQ QGGLCVALGE
QLIEDVQALS GTINDLQDQI DSLAEVVLQN RRGLDLLTAE QGGICLALQE

ECCFYVDH
ECCFYVDH
ECCFYVDH
```

ECCFYVDH
ECCFYVDH
ECCFYVDH
ECCFYVDH
ECCFYVDH
ECCFYVDH
ECCFYVDH
ECCFYVDH
ECCFYVDH
ECCFYVDH
ECCFYVDH
ECCFYVDH
ECCFYVDH
ECCFYVDH
ECCFYVDH
KCCFYVNH
ECCFYVDH
ECCFYVDH
KCCFYVDH
ECCFYVDH
ECCFYVDH
ECCFYVDH
EC--WSSH
EC--WSSH
EC--WSSH
EC--WSSH
EC--WSSH
EC--WSSH
EC--WSSH
EC--WSSH
EC--WSSH
EC--WSSH
ECCFYVDH
EC--WSSH
ECCFYVDH
ECCFYVDH
ECCFYVDH
ECCFYVDH
ECCFYVDH
ECCFYVDH
ECCFYVDH
ECCFYVDH
ECCFYVDH
ECCFYVDH

ECCFYVDH
ECCFYVDH
ECCFYVDH
ECCFYVDH
ECCFYVDH
ECCFYVDH
KCCFYVDH
KCCFYVDH
ECCFYVDH
ECCFYVDH
ECCFYVDH
ECCFYVDH
ECCFYVDH
ECCFYVDH
ECCFYVDH
ECCFYVDH
ECCFYVDH
ECCFYVDH
ECCFYVDH
ECCFYVDH
ECCFYVDH
ECCFYVDH
ECCFYVDH
ECCFYVDH
E-CFYVDH
ECYFYVDH
E-CFYVDH
E-CFYVDH
E-CFYVDH
E-CFYVDH
E-CFYVDH
E-CFYVDH
E-CFYVDH
ECCFYADH
ECCFYADH
ECCFYVDH
ECCFYVDH
ECCFYVDH
ECCFYVDH
ECCFYVDH
ECCFYVDH
ECCFYIDH
ECCFYVDH
ECCFYVDH

ECCFYVDH
ECCFYVDH
ECCFYVDH
ECCFYVDH
ECCFYVDH
ECCFYVDH
ECCFYVDH
ECCFYVDH
ECCFYVDH
ECCFYVVH
ECCFYVDH
ECCFYVDH
ECCFYVDH
ECCFYIDH
ECCFYVDH
ECCFYVDH
ECCFYVDH
ECCFYVDH
ECCFYVDH
ECCFYVDH
ECCFYVDH
ECCFYVNH
ECCFYIDH
ECCFYADH
ECCFYADH
ECCFYADH
ECCFYADH
ECCFYADH
ECCFYADH
KCCFYANK

**Dataset S2.** The alignments used to build the phylogenetic trees of Gag, Pol and Env represented in Fig. S7.

```
Gag
139 225
LUXW01011341.1       NHPPFSEDPQ RLTGLVESLM FSHQPTWDDC QQLLQTLFTT EERERILLEA
LUXS01031219.1       NHPPFSEDPQ RLTGLVESLM FSHQPTWDDC QQLLQTLFTT EERERILLEA
LUXX01090830.1       NHPPFSEDPQ RLTGLVESLM FSHQPTWDDC QQLLQTLFTT EERERILLEA
LT634572.1_d         NHPPFSEDPQ RLTGLVESLM FSHQPTWDDC QQLLQTLFTT EERERILLEA
CM000818.5_a         NHPPFSEDPQ RLTGLVESLM FSHQPTWDDC QQLLQTLFTT EERERILLEA
LUXS01011274.1       NHPPFSEDPQ RLTGLVESLM FSHQPTWDDC QQLLQTLFTT EERERILLEA
LUXU01062474.1       NHPPFSEDPQ RLTGLVESLM FSHQPTWDDC QQLLQTLFTT EERERILLEA
LUXX01045907.1       NHPPFSEDPQ RLTGLVESLM FSHQPTWDDC QQLLQTLFTT EERERILLEA
LUXX01080744.1       NHPPFSEDPQ RLTGLVESLM FSHQPTWDDC QQLLQTLFTT EERERILLEA
LUXU01038095.1       NHPPFSEDPQ RLTGLVESLM FSHQPTWDDC QQLLQTLFTT EERERILLEA
PERV_C               NHPPFSEDPQ RLTGLVESLM FSHQPTWDDC QQLLQTLFTT EERERILLEA
LUXW01089741.1       NHPPFSEDPQ RLTGLVESLM FSHQPTWDDC QQLLQTLFTT EERERILLEA
LUXT01005398.1       NHPPFSEDPQ RLTGLVESLM FSHQPTWDDC QQLLQTLFTT EERERILLEA
CM000826.5_c         NHPPFSEDPQ RLTGLVESLM FSHQPTWDDC QQLLQTLFTT EERERILLEA
CM000812.5_b         NHPPFSEDPQ RLTGLVESLM FSHQPTWDDC QQLLQTLFTT EERERILLEA
CM000814.5_a         NHPPFSEDPQ RLTGLVESLM FSHQPTWDDC QQLLQTLFTT EERERILLEA
CM000824.5_a         NHPPFSEDPQ RLTGLVESLM FSHQPTWDDC QQLLQTLFTT EERERILLEA
CM000812.5_a         NHPPFSEDPQ RLTGLVESLM FSHQPTWDDC QQLLQTLFTT EERERILLEA
PERV-A               NHPPFSEDPQ RLTGLVESLM FSHQPTWDDC QQLLQTLFTT EERKRILLEA
CM000816.5_b         NHPPFSEDPQ RLTGLVESLM FSHQPTWDDC QQLLQTLFTT EERERILLEA
AOCR01107158.1       NHPPFSEDPQ RLTGLVESLM FSHQPTWDDC QQLLQTLFTT EERERILLEA
CM000825.5_b         NHPPFSEDPQ RLTGLVESLM FSHQPTWDDC QQLLQTLFTT EERERILLEA
AOCR01079940.1       NHPPFSEDPQ RLTGLVESLM FSHQPTWDDC QQLLQTLFTT EERERILLEA
AOCR01098665.1       NHPPFSEDPQ RLTGLVESLM FSHQPTWDDC QQLLQTLFTT EERERILLEA
KQ004162.1           NHPPFSEDPQ RLTALVESLM FSHQPTWDDC QQLLQTLFTT EERERILLEA
CM000819.5_b         NHPPFSEDPQ RLTGLVESLM FSHQPTWDDC QQLLQTLFTT EERERILLEA
CM000823.5_b         NHPPFSEDPQ RLTGLVESLM FSHQPTWDDC QQLLQTLFTT EERERILLEA
CM000828.5_a         NHPPFSEDPQ RLTGLVESLM FSHQPTWDNC QQLLQTLFTT EERERILLEA
AEMK02000137.1       NHPPFSEDPQ RLTGLVESLM FSHQPTWDDC QQLLQTLFTT EERERILLEA
PERV_B               NHPPFSEDPQ RLTGLVESLM FSHQPTWDDC QQLLQTLFTT EEQERILLEA
CM000820.5           NHPPFSEDPQ RLTGLVESLM FSHQPTWDDC QQLLQTLFTT EERERILLEA
CM000814.5_c         NHPPFSEDPQ RLTGLVESLM FSHQPTWDDC QQLLQTLFTT EERERILLEA
CM000812.5_c         NHPPFSEDPQ RLTGLVESLM FSHQPTWDDC QQLLQTLFTT EERERILLEA
AEMK02000197.1       NHPPFSEDPQ RLTGLVESLM FSHQPTWDDC QQLLQTLFTT EERERILLEA
CM000827.5           NHPPFSEDPQ RLTGLVESLM FSHQPTWDDC QQLLQTLFTT EERERILLEA
LUXR01057621.1       NHPPFSEDPQ RLTGLVESLM FSHQPTWDDC QQLLQTLFTT EERERILLEA
LUXV01020898.1       NHPPFSEDPQ RLTGLVESLM FSHQPTWDDC QQLLQTLFTT EERERILLEA
LUXS01071149.1       NHPPFSEDPQ RLTGLVESLM FSHQPTWDDC QQLLQTLFTT EERERILLEA
```

| | | | | | |
|---|---|---|---|---|---|
| LUXR01088996.1 | NHPPFSEDPQ | RLTGLVESLM | FSHQPTWDDC | QQLLQTLFTT | EERERILLEA |
| CM000825.5_a | NHPPFSEDPQ | RLTGLVESLM | FSHQPTWDDC | QQLLQTLFTT | EERERILLEA |
| CM000819.5_a | NHPPFSEDPQ | RLTGLVESLM | FSHQPTWDDC | QQLLQTLFTT | EERERILLEA |
| CM000826.5_a | NHPPFSEDPQ | RLTGLVESLM | FSHQPTWDDC | QQLLQTLFTT | EERERILLEA |
| CM000815.5 | NHPPFSEDPQ | RLTGLVESLM | FSHQPTWDDC | QQLLQTLFTT | EE-ERILLEA |
| CM000822.5_b | NHPPFSEDPQ | RLTGLVESLM | FSHQPTWDDC | QQLLQTLFTT | E--ERILLEA |
| LUXV01022316.1 | NHPPFSEDPQ | RLTGLVESLM | FSHQPTWDDC | QQLLQTLFTT | EERERILLEA |
| LUXY01101100.1 | NHPPFSEDPQ | RLTGLVESLM | FSHQPTWDDC | QQLLQTLFTT | EERERILLEA |
| CM000828.5_b | NHPPFSEDPQ | CLTGLVESLM | FSHQPTWDDC | QQLLQTLFTT | EERERILLEA |
| CM000824.5_b | NHPPFSEDPQ | RLTGLVESLM | FSHQPTWDDC | QQLLQTLFTT | EERERILLEA |
| LUXR01004647.1 | NHPPFSEDPQ | RLTGLVESLM | FSHQPTWDDC | QQLLQTLFTT | EERERILLEA |
| CM000818.5_b | NHPPFSEDPQ | RLTGLVESLM | FSHQPTWDDC | QQLLQTLFTT | EERERILLEA |
| CM000830.5_d | NHPPFSEDPQ | RLMGLVESLM | FSHQPTWDDC | QQLLQTLFTT | EE-ERILLEA |
| CM000814.5_b | NHPPFSEDPQ | RLTGLVESLM | FSHQPTWDNC | QQLLQTLFTT | EERERILLEA |
| AEMK02000476.1 | NHPPFSEDPQ | RLTGLVESLM | FSHQPTWDDC | QQLLQTLFTT | EE-ERILLEA |
| LUXS01081946.1 | NHPPFSEDPQ | RLTGLVESLM | FSHQPTWDDF | QQLLQTLFTT | EGRERILLEA |
| CM000828.5_c | NHPPFSEDPQ | RLTGLVESLM | FSHQPTWDDC | QQLLQTLFTT | EGRERILLEA |
| LUXQ01076544.1 | NHPPFSEDPQ | RLTGLVESLM | FSHQPTWDDC | QQLLQTLFTT | EGRERILLEA |
| LIDP01000002.1_a | NHPPFSEDPQ | R---LVESLM | FSHQPTWDDC | QQLLQTLFTT | EERERILLEA |
| CM000816.5_a | NHPPFSEDPQ | RLTALVESLM | FSHQPTWNDC | QQLLQTLFTT | EKRERILLEA |
| AEMK02000141.1 | NHPPFSEDPQ | RLTALVESLM | FSHQPTWDDC | QQLLQTLFTT | EERERILLET |
| AOCR01031838.1 | NHPPFSEDPQ | CLTGLVESLM | FSHQPTWDDC | QQLLQTLFTT | EERERILLEA |
| CM000813.5 | NHPPFSEDPQ | CLTGLVESLM | FSHQPTWDDC | QQLLQTLSTT | GERERILLEA |
| LIDP01000002.1_b | NHPPFSEDPQ | CLTGLVESLM | FSHQPTWDDC | QQLLQTLSTT | GERERILLEA |
| AEMK02000133.1 | NHPPFSEDPQ | HLTGLVESLM | FSHQPTWDDC | QQLLQTLFTT | EERERILLEA |
| LUXU01085042.1 | NHPPFSEDPQ | RLTGLVESLM | FSHQPTWDDC | QQLLQTLFTT | EERERILLEA |
| AEMK02000393.1_a | NHPPFSEDPQ | CLVGLVESLM | FSHQPTWDDC | QQLLQTLFTT | EERERILLEA |
| LUXV01069687.1 | NHPPFSEDPQ | CLVGLVESLM | FSHQPTWDDC | QQLLQTLFTT | EERERILLEA |
| LUXY01013808.1 | NHPPFSEDPQ | CLVGLVESLM | FSHQPTWDDC | QQLLQTLFTT | EERERILLEA |
| LUXX01033707.1 | NHPPFSEDPQ | CLVGLVESLM | FSHQPTWDDC | QQLLQTLFTT | EERERILLEA |
| LUXS01081675.1 | NHPPFSEDPQ | RLTGLVESLM | FSHQPTWDDC | QQLLQTLFTT | EERERILLEA |
| CM000830.5_b | NHPPFSEDPQ | CLTGLVESLM | FSHQPTWDDC | QQLLQTLFTT | EE-ERILLEA |
| LT634572.1_f | NHPPFSEDPQ | RLTALVESLM | FSHQPTWDNC | QQLLQTLFTT | EKRERILLEA |
| LT634572.1_g | NHPPFSEDPQ | RLTALVESLM | FSHQPTWDNC | QQLLQTLFTT | EKRERILLEA |
| LUXY01042539.1 | NHPPFSEDPQ | RLTALVESLM | FSHQPT-DNC | QQLLQTLFTT | EEREKILLEA |
| LT634572.1_c | NHPPFSEDPQ | RLTALVESLM | FSHQPTWDDC | QQLLQTHFTT | EERERILLKA |
| CM000822.5_a | NHPPFSEDPQ | CLTALVESLM | FSHQPTWDDC | QQLLQTLFTT | EERERILLEA |
| LT634572.1_a | NHPPSSEDPQ | HLTGLVESLM | FSHQPTWYDC | QQLLQTLFTT | EERERILLEA |
| LT634572.1_b | NHPPSSEDPQ | HLTGLVESLI | FSHQPTWYDC | QQLLQTLFTT | EE-ERILLEA |
| AJKK01232762.1 | NHPPSSEDPQ | HLTGLVESLI | FSHQPTWYDC | QQLLQTLFTT | EERERILLEA |
| LUXW01074277.1 | NHPPFSEDPQ | RLTGLVESLM | FSHQPTWDDC | QQLLQTLFTT | EERERILLEA |
| LUXX01006697.1 | NHPPFSEDPQ | RLTGLVESLM | FSHQPTWDDC | QQLLQTLFTT | EERERILLEA |
| LUXT01031363.1 | NHPPFSEDPQ | RLTGLVESLM | FSHQPTWDDC | QQLLQTLFTT | EERERILLEA |
| LUXY01052675.1 | NHPPFSEDPQ | RLTGLVESLM | FSHQPTWDDC | QQLLQTLFTT | EERERILLEA |

| | | | | | |
|---|---|---|---|---|---|
| LUXQ01112116.1 | NHPPFSEDPQ | RLTGLVESLM | FSHQPTWDDC | QQLLQTLFTT | EERERILLET |
| LUXV01092473.1 | NHPPFSEDPQ | RLTGLVESLM | FSHQPTWDDC | QQLLQTLFTT | EERERILLEA |
| LUXS01060324.1 | NHPPFSEDPQ | RLTGLVESLM | FSHQPTWDDC | QQLLQTLFTT | EE-ERILLET |
| KQ001904.1 | NYPPFSEDPQ | CLTGLVESLM | FSHQPTWDDC | QQLLQTLFTT | EERERILLEA |
| CM000830.5_c | NHPPFSENPQ | SLMGLVESLM | FSHQPTWDDC | QQQLQTLFTT | EEREKILLEA |
| LUXU01061831.1_a | NHPPFSEDPQ | SLMGLVESLM | FSHQPTWDDC | QQQLQTLFTT | EEREKILLEA |
| LUXU01061831.1_b | NHPPFSEDPQ | SLMGLVESLM | FSHQPTWDDC | QQQLQTLFTT | EEREKILLEA |
| LUXW01038847.1 | NHPPFSENPQ | SLMGLVESLM | FSHQPTWDDC | QQQLQTLFTT | EEREKILLEA |
| LUXX01056080.1 | NHPPFSENPQ | SLMGLVESLM | FSHQPTWDDC | QQQLQTLFTT | EEREKILLEA |
| LUXS01056853.1 | NHPPFSENPQ | SLMGLVESLM | FSHQPTWDDC | QQQLQTLFTT | EEREKILLEA |
| AEMK02000536.1 | NHPPFSEDPQ | RLTGLVESLM | FSHQPTWDDC | QQLLQTLFTT | EERERILLEA |
| LUXT01067056.1 | NHPPFSEDPQ | RLTALVESLM | FSHQPTWDDC | QQLLQTLFTT | EERERILLET |
| CM000826.5_b | NHPPFSEDPQ | CLTGL-GSLM | FSHQPTWDDC | QQLLQTLSTT | EERERILLEA |
| LUXY01106724.1 | NHPPFLEDPQ | CLTGLGESLM | FSHQPTWDDC | QQLLQTLSTT | EERERILLEA |
| LIDP01000015.1 | NHPPFSEDPQ | CLTGLGESLM | FSHQPTWDDC | QQLLQTLSTT | EERERILLEA |
| eJJRV_NW_004504375.1 | NHPPFSEEPW | HLTGLIESLM | FSHQPTWDDC | QQLLQTLFTT | KEQERILIEA |
| eJJRV_NW_004504334.1 | KYPPFSEEPR | RLTVLIESLM | FSHQPTWDD- | QQLLQTLFTT | EEREGILIEA |
| RNERV_6 | NHPSFSENPS | GLTGLLESLM | FSHQPTWDDC | QQLLQALFTT | EEKERILLEA |
| MMERV_35 | NHPPFSENPS | GLTGLLESLM | FSHQPTWDDC | QQLLQVLFTT | EERERILMEA |
| MMERV_11 | NHPPFSENPS | GLTGLLESLM | FSHQPTWDDC | QQLLQVLFTT | EERERILMEA |
| MDEV | NHPPFSENPS | GLTGLLESLM | FSHQPTWDDC | QQLLQVLFTT | EERERILMEA |
| MMERV_19 | NHPPFSENPP | GLTGLLESLM | FSHQPTWDDC | QQFLQVLFTT | EERERILMEA |
| MMERV_18 | NHPPFSENPP | GLTGLLESLM | FSHQPTWDDC | QQFLQVLFTT | EERERILMEA |
| MMERV_5 | NHPPFSENHS | GLTGLLESLM | FSHQPTWDDC | QQLLQVLFTI | EERERILMEA |
| KoRV | NHPSFSENPT | GLTGLLESLM | FSHQPTWDDC | QQLLQVLFTT | EERERILLEA |
| GALV | NHPSFSENPA | GLTGLLESLM | FSHQPTWDDC | QQLLQILFTT | EERERILLEA |
| MMERV_39 | NHASFSENPT | SLVGLVESLM | FSHQPTWDDC | QQLLQVLFTT | EEKERILLEA |
| MMERV_8 | NHASFSENPT | SLVGLVESLM | FSHQPTWDDC | QQLLQVLFTT | EEKERILLEA |
| musMus5656 | NHASFSENPT | SLVGLVESLM | FSHQPTWDDC | QQLLQVLFTT | EEKERILLEA |
| MCERV_9 | NHPTFSENPA | GLTSLVESLM | FSHQPT-DDC | QQLLQVLFTT | EEKERILVEA |
| MMERV_21 | HHPPFSENPA | GLTGLVESLM | YSHQPTWDDC | QQLLQTLFTT | EERERILLEA |
| MMERV_10 | HHPPFSENPA | GLTGLVESLM | YSHQPTWDDC | QQLLQTLFTT | EERERILLEA |
| MMERV_22 | HHPPFSENPA | GLTGLVESLM | YSHQPTWDDC | QQLLQTLFTT | EERERILLEA |
| MMERV_3 | NHPPFSENPA | GLTGLVESLM | YSHQPTWDDC | HQLLQTLFTT | EERERILLEA |
| MSERV_1 | NHSTFSENPA | GLTCLVESLM | FSHQPTWDDC | LQLLQVLFTT | EEKERILLEA |
| MCERV_12 | NHPPFSENPA | SLTGLVGSLM | FSHQPTWDDC | QQ-LQVLFTT | EKKERILHEA |
| MPERV_8 | NHPPFSENPA | SLTGLVESLM | FSHQTTWDDC | Q---QVLFTT | EEKEKILHEA |
| ratNor293 | NHPSFSENPA | GLTGLLESLM | FSHQPMWDGC | QQLLQVLFTT | EEEERILLEA |
| ratNor2288 | NHPFSSADLY | NKTNHIESLM | FSHQPTWDDC | QQLLQVLFTT | EEKERILLEA |
| OOEV | HNPTFSENPQ | ALTALIESLV | FSHQPTWDDC | QQLLQTLLTT | EERQRVLLEA |
| RD114 | HNPSFSQEPQ | ALTSLIESIL | LTHQPTWDDC | QQLLQVLLTT | EERQRVLLEA |
| BaEV | HNPSFSQDPQ | ALTSLIESIL | LTHQPTWDDC | QQLLQVLLTT | EERQRVLLEA |
| FeLV | HNPPFSQDPV | ALTNLIESIL | VTHQPTWDDC | QQLLQALLTG | EERQRVLLEA |
| M-MuLV | NNPSFSEDPG | KLTALIESVL | ITHQPTWDDC | QQLLGTLLTG | EEKQRVLLEA |

```
F-MuLV              NNPSFSEDPA KLTALIESVL LTHQPTWDDC QQLLGTLLTG EEKQRVLLEA
R-MuLV              NNPSFSEDPG KLTALIESVL LTHQPTWDDC QQLLGTLLTG EEKQRVLLEA
M-CRV               NNPSFSEDPG KLTALIESVL TTHQPTWDDC QQLLGTLLTG EEKQRVLLEA
RfRV                QNPPFSEDPK GLTDLFESVM HTHSPTWDDC QQLLKTLFTT EERERILTEA
ratNor338           NHPSFSENPS ILTGLMESLM NSHQPMWGDC QQLLQILFTT EECERILLEA
musMus7492          NHAPFSENPS SLTNLIKSLM FFHQPTWDDC QQLLQVLFTT EEKEKILLRQ
musMus9005          NHALFSENPS SLTNIMYSLM FFHQPNWDDC QQLLQVLFST EE-NRILLEA
ratNor4542          NHLSFSENPA SLTGLVESLI LSHQPTWDDV QQLLLVLFTT EEKERILLEA
MMERV_33            NHPSFSENPA SLTGLVESLM YSHQPTWDDC QQLLQVLYTT EEREKILLEA
REV                 QNPSFSQAPD EVISLLESVF YTHQPTWDDC QQLLRTLFTT EERERVRTES
ePCRV_KN678005.1    NHPSFSDDPQ KLTSLMESLM FSHQPTWDDC QQLLQVLFTT EEHERILLEA
ePCRV_KN680906.1    NLPSFSDDPQ KLTSLMESLM FSHQPTWDDC QQLLQVLFTT EERERILLEA
ePCRV_KN676905.1    NHPSFSDDPQ KLTSLMESLM FSHQPTWEDC QQLLQVLFTT EERERILLKA

                    RKNVPGADGR PTQLQNEIDM GFPLTRPGWD YNTAEGRESL KIYRQALVAG
                    RKNVPGADGR PTQLQNEIDM GFPLTRPGWD YNTAEGRESL KIYRQALVAG
                    RKNVPGADGR PTQLQNEIDM GFPLTRPGWD YNTAEGRESL KIYRQALVAG
                    RKNVPGADGR PTQLQNEIDM GFPLTRPGWD YNTAEGRESL KIYRQALVAG
                    RKNVPGADGR PTQLQNEIDM GFPLTRPGWD YNTAEGRESL KIYRQALVAG
                    RKNVPGADGR PTQLQNEIDM GFPLTRPGWD YNTAEGRESL KIYRQALVAG
                    RKNVPGADGR PTQLQNEIDM GFPLTRPGWD YNTAEGRESL KIYRQALVAG
                    RKNVPGADGR PTQLQNEIDM GFPLTRPGWD YNTAEGRESL KIYRQALVAG
                    RKNVPGADGR PTQLQNEIDM GFPLTRPGWD YNTAEGRESL KIYRQALVAG
                    RKNVPGADGR PTQLQNEIDM GFPLTRPGWD YNTAEGRESL KIYRQALVAG
                    RKNVPGADGR PTQLQNEIDM GFPLTRPGWD YNTAEGRESL KIYRQALVAG
                    RKNVPGADGR PTQLQNEIDM GFPLTRPGWD YNTAEGRESL KIYRQALVAG
                    RKNVPGADGR PTQLQNEIDM GFPLTRPGWD YNTAEGRESL KIYRQALVAG
                    RKNVPGADGR PTQLQNEIDM GFPLTRPGWD YNTAEGRESL KIYRQALVAG
                    RKNVPGADGR PTQLQNEIDM GFPLTRPGWD YNTAEGRESL KIYRQALVAG
                    RKNVPGADGR PTQLQNEIDM GFPLTRPGWD YNTAEGRESL KIYRQALVAG
                    RKNVPGADGR PTQLQNEIDM GFPLTRPGWD YNTAEGRESL KIYRQALVAG
                    RKNVPGADGR PTQLQNEIDM GFPLTRPGWD YNTAEGRESL KIYRQALVAG
                    RKNVPGADGR PTQLQNEIDM GFPLTRPGWD YNTAEGRESL KIYRQALVAG
                    RKNVPGADGR PTQLQNEIDM GFPLTRPGWD YNTAEGRESL NIYRQALVAG
                    RKNVPGADGR PTRLQNEIDM GFPLTRPGWD YNTAEGRESL KIYRQALVAG
                    RKNVPGADGR PTRLQNEIDM GFPLTRPGWD YNTAEGRESL KIYRQALVAG
                    RKNVPGADGR PTRLQNEIDM GFPLTRPGWD YNTAEGRESL KIYRQALVAG
                    RKNVPGADGR PTRLQNEIDM GFPLTRPGWD YNTAEGRESL KIYRQALVAG
                    RKNVPGADGR PTQLQNEIDM GFPLTRPGWD YNTAEGRESL KIYRQALVAG
                    RKNVPGADGR PTQLQNEIDM GFPLTRPGWD YNTAEGRESL KIYRQALVAG
                    RKNVPGADGR PTQLQNEIDM GFPLTRPGWD YNTAEGRESL KIYRQALVAG
                    KKNVPGADGR PTQLQNEIDM GFPLTRPGWD YNTAEGRESL KIYRQALVAG
                    RKNVPGADGR PTQLQNEIDM GFPLTRPGWD YNTAEGRESL KIYCQALVAG
                    RKNVPGADGR PTQLQNEIDM GFPLTRPGWD YNTAEGRESL KIYRQALVAG
```

```
RKNVPGADGR PTQLQNEIDM GFPLTRPGWD YNTAEGRESL KIYRQALVAG
RKNVPGADGR PTQLQNEIDM GFPLTRPGWD YNTAEGRESL KIYRQALVAG
RKNVPGADGR PTQLQNEIDM GFPLTRPGWD YNTAEGRESL KIYRQALVAG
RKNVPGADGR PTQLQNEIDM GFPLTRPGWD YNTAEGRESL KIYRQALVAG
RKNVPGADGR PTQLQNEIDM GFPLTRPGWD YNTAEGRESL KIYRQALVAG
RKNVPGADGR PTQLQNEIDM GFPLTRPGWD YNTAEGRESL KIYRQALVAG
RKNVPGADGR PTQLQNEIDM GFPLTRPGWD YNTAEGRESL KIYRQALVAG
RKNVPGADGR PTQLQNEIDM GFPLTRPGWD YNTAEGRESL KIYRQALVAG
RKNVPGADGR PTQLQNEIDM GFPLTRPGWD YNTAEGRESL KIYRQALVAG
RKNVPGADGR PTQLQNEIDM GFPLTRPGWD YNTAEGRESL KIYRQALVAG
RKNVPGADGR PTQLQNEIDM GFPLTRPGWD YNTAEGRESL KIYRQALVAG
RKNVPGADGR PTQLQNEIDM GFPLTRPGWD YNTAEGRESL KIYRQALVAG
RKNVPGADGR PTQLQNEIDM GFPLTRPGWD YNTAEGRESL KIYRQALVAG
RKNVPGADGR PTQLQNEIDM GFPLTRPGWD YNTAEGRESL KIYRQALVAG
RKNVPGADGR PTQLQNEIDM GFPLTRPGWD YNTAEGRESL KIYRQALVAG
RKNVPGADGR PTQLQNEIDM GFPLTRPGWD YNTAEGRESL KIYRQALVAG
RKNVPGADGR PTQLQNEIDM GFPLTRPSWD YNTAEGRESL KIYRQALVAG
RKNVPGADGR PTQLQNEIDM GFPLTRPSWD YNTAEGRESL KIYRQALVAG
RKNVPGADGR PTQLQNETDM GFPLTPPGWD YNTAEGRESL KIYGQALVAG
RKNVPGADGR PTQLQNETDM GFPLTRPGWD YNTAEGRESL KIYRQALVAG
RKNVPGADGR PTQLQNEIDM GFPLTRPGWD YNTAEGRESL KIYRQALVAG
RKNVPGADGR PTQLQNEIDM GFPLTRPSWD YNTAEGRESL KIYHQALVAG
RKNVPGADG- PTQLQNEIDM GFPLTRPGWD YNTAEGRESL KIYRQALVAG
RKNVPGADGR PTQLQNEIDM GFPLTRPGWD YNTAEGRESL KIYCQALVAG
RKNVPGADGR PTQLQNEIDM GFPLTRPGWD YNTAEGRESL KIYRQALVAG
RKNVPGADGR PTQLQNEIDM GFPLTRPGWD YNTAEGRESL KIYRQALVAG
RKNVPGADGR PTQLQNEIDM RFPLTRPG-D YNTAEGRESL KIYRQALVAG
RKNVPGADGR PTQLQNEIDM GFPLTRPG-D YNTAEGRESL KIYRQALVAG
KKNVPGADGR PTQLQNKIDM GFPLTRPG-D YNTAEGRESL KIYRQALVAG
RKNVPGADGR PTQLQNEIDM GFPLTRPDWD YNTAEGRESL KIYRQALVVG
KKNVPGADGR PTQLQNEIDM GFPLTRP-WD YNTAEGRESL KIYRQALVVG
KKNVPGADGR PTQLQNEIDM GFPLTRP-WD YNTAEGRESL KIYRQALVVG
RKNVPGADGR PTQLQNEIDM GFP-THPGWD YNTAEGRESL KIYRQALVAG
RKNVPGADGR PTQLQNEIDM GFPLTRPSWD YNTAEGRESL KIYHQALVAG
RKNVPGANGR PTQLQNEIDM GFPLTRPSWD YNTAEGRESL KIYHQALVAG
RKNVPGANGR PTQLQNEIDM GFPLTRPSWD YNTAEGRESL KIYRQALVAG
RKNVPGANGR PTQLQNEIDM GFPLTRPSWD YNTAEGRESL KIYRQALVVG
RKNVPGANGR PTQLQNEIDM GFPLTRPSWD YNTAEGRESL KIYHQALVAG
RKNVPGADGR PTQLQNEIDM GFPLTRPGWD YNTAEGRESL KIYRQALVAG
RKNVPGADGQ PTQLQNEIDM GFPLTRPGWD YNTAEGRESL KIYRQALVAG
RKNVPGADGR PTQLQNEIDM GFPLTRPG-D YNTAEGRESL KIYRQALVAD
RKNVPGADGR PTQLQNEIDM GFPLTRPG-D YNTAEGRESL KIYRQALVAD
RKNVPGADGR PTQLQNEIDI GFPLTRPG-D YNTAEGRESL KIYRQALVAG
KKNVPGADGR PTQLQNEIDM RFPLTRPG-N YNTAEGRESL KIYRQALMAG
```

RKNVPGADGR PTQLQNEIDM GFPLTHPG-D YNTAEGRESL KIYRQALVAG
RKNIPGADGR PTQLQNEIDM GFPLTRHGWD YNTAKGRESL KIYRQALVAG
RKNIPGADGR PTQLQNEIDM GFPLTRPGWD HNTAVGRESL KIYRQALLAG
RKNIPGADGR PTQLQNEIDM GFPLTRPGWD YNTAVGRESL KIYRQALLAG
RKNVPGADGR PTQLQNEIDM GFPLTRPGWD YNTAEGRESL KIYHQALVAG
RKNVPGADGR PTQLQNEIDM GFPLTRPGWD YNTAEGRESL KIYHQALVAG
RKNVPGADGR PTQLQNEIDM GFPLTRPGWD YNTAEGRESL KIYHQALVAG
RKNVPGADG- PTQLQNEIDM GFPLTRPGWD YNTAEGRESL KIYHQALVAG
RKNVPGADGR PTQLQNEIDM GFPLTRPGWD YNTAEGRESL KIYHQALVAG
RKNVPGADGR PTQLQNEIDM GFPLTRPGWD YNTAEGRESL KIYHQALVAG
RRNVPGADG- PTQLQNEIDM GFPLTRPGWD YNTAEGRESL KIYHQALVAG
RKNVPGADGR PTQLQNEIDM GFPLTRPGWD YNTAEGRESL KIYHQALVAG
RKNVPGADGR PTHLQNEIDM GFPLTRPGWD YNTAEGRESV KIYCQSLVAG
RKNVPGADGR PTHLQNEIDM GFPLTRPGWD YNTAEGRESV KIYCQSLVAG
RKNVPGADGR PTHLQNEIDM GFPLTRPGWD YNTAEGRESV KIYCQSLVAG
RKNVPGADGQ PTHLQNEIDM GFPLTRPGWD YNTAEGRESV KIYCQALVAG
RKNVPGADGQ PTHLQNEIDM GFPLTRPGWD YNTAEGRESV KIYCQALVAG
RKNVPGADGQ PTHLQNEIDM GFPLTRPGWD YNTAEGRESV KIYCQALVAG
RKNVPGADGR PTQLQNEIDM GFPLTRPGWD YNTAEGRESL KIYRQALVAG
KKNVPGADGR PTQLQNEIDM GFPLTRPGWD YNTAEGRESL KIYHQALVAG
KKKVPGANGR PTQLQNEIDM EFPLTRPSWD YNTAEGRESL KIYRQALVAG
KKKVPGANGR PTQLQNEIDM EFPLTRPSWD YNTAEGRESL KIYRQALVAG
KKKVPGANGR PTQLQNEIDM EFPLTRPSWD YNTAEGRESL KIYRQALVAG
RKNVPGANRQ PTQLQHEIDM GFPPTCPVWD YNTAEGRESL KICLQALVAG
RKNVPGA-GQ PSQLQHEIDM GFPLTHPAWE YNTAEGRQSL KIYCQALVAD
RKNVPGPDGT PTNLPNLIDE TFPLTRPAWD FNTAEGRERL TVYRRTLVAG
RKNVLGEDGT PTALPNLVDE AFPLNRPNWD YNTAEGRGRL LVYRRTLVAG
RKNVLGEDGT PTALPNLVDE AFPLNRPNWD YNTAEGRGRL LVYRRTLVAG
RKNVLGEDGT PTALPNLVDE AFPLNRPNWD YNTAEGRGRL LVYRRTLVAG
RKNVLGEDGT PTALPNLVDE AFPLNRPNWD YNTAEGRGCL LVYRRTLVAG
RKNVLGEDGT PTALPNLVDE AFPLNRPNWD YNTAEGRGCL LVYRRTLVAG
RKNVLGEDGT PTALPNLVDE AFPLNRPNWD YNTAEGRGRL LVYRRTLVAG
RKNVLGVNGA PTQLENLINE AFPLNRPQWD HNTAEGRERL LVYRRTLVAG
RKNVLGDNGA PTQLENLINE AFPLNRPHWD YNTAAGRERL LVYRRTLVAG
QKNVPGVNGA PTNLPNEINA GFPLDRPDWN YNTAQGRERL AVYRRALVAG
QKNVPGVNGA PTNLPNEINA GFPLDRPDWN YNTAQGRERL AVYRRALVAG
QKNVPGVNRA PTNLPNEINA GFPLDRPDWN YNTAQGRERL AVYRRALVAG
RKNVPGINGA QTNLPNEINA GFPLDRPNWD YNTAQGRERL TVYRQALVAG
RKNVRDEAGR PVQTPAEIDE GFPLTRPRWD YNTASGRERL SNYRRVLVAG
RKNVRDEAGR PVQTPAEIDE GFPLTRPRWD YNTASGRERL SNYRRVLVAG
RKNVRDEAGR PVQTPAEIDE GFPLTRPRWD YNTASGRERL SNYRRVLVAG
RKNVRDEAGR PVQTPAEIDK GFPLTQPRWD YNTASGRERL SNYRRVLVAG
RKNILGINGA PINLPNEIYA GFPLDRPDWD YNTDQGREQL TVYHQALVAG
RKNVPGPDRF PTTLPNEIDA GFPLDRPNWD FNMAAGRERL TVYCQTLVAG

RKNVPGPNGF PTILLNEIDA GFPLDFPNWD FNTAAVRE-L AIYCQTLVAG
RKNVPGNGGV PTNLPNEMDA GFPLNRLNWD YNTAEGRERS TVYRRALVAG
RKHVPGPDGT PTQMPNLIDA AFPLDRPPWD YNTAEGRERL STYRRALVAG
RKNVLGANGQ PTQLPNEIDV GFPLVRPNWD FNAPEGRERL KMYRQALVVG
RKNVPGPGGF PTQLPNEIDE GFPLTRPDWD YETAPGRESL RIYRQALLAG
RKNVPGPGGL PTQLPNEIDE GFPLTRPDWD YETAPGRESL RIYRQALLAG
RKQVPGEDGR PTQLPNVIDE TFPLTRPNWD FATPAGREHL RLYRQLLLAG
RKAVRGDDGR PTQLPNEVDA AFPLERPDWD YTTQAGRNHL VHYRQLLLAG
RKAVRGEDGR PTQLPNDIND AFPLERPDWD YNTQRGRNHL VHYRQLLLAG
RKAVRGEDGR PTQLPNDIND AFPLERPDWD YNTQRGRNHL VHYRQLLLAG
RKAVRGDDGR PTQLPNEIEA AFPLERPDWD YTTLRGRNHL VLYRQLLLAG
RKNVPGDNGR PTTLPNLIDE RFPLNRPDWD FGNAEGRERL RVYRQTLMAG
HKNVPA-DVL LTQLSNVIEA GFPLDQPNWD FNTAAGRERL TVH-QALVAS
ERTCWGPDGW PTQLPNEIDA GFPLTHPDW- ---GEGREQL TVFRQALVAG
RKNVLGAKGL PTQF-NEIDA GFPLTCPDWD RNRPEGREQL TVFHWALVAG
WKNIPGTNGA PTNLRNEIDA GFPLNRPDWD YNTSAGMEWL TSYRQALMAG
RKNVPRANGV PTNLPNETDA GFPLN-PAWD FNMTADRAWL TVYHQALMAG
RREVRNDQGV QVTDEREIEA QFPATRPDWD PNTGRGNDNL ERYRQILLRG
RKNVPGTDGR PTQLPNEIDQ GFPLTRPLWD FNTPEGRERL TIYRQALVAG
RKNMPGTDGR PTQLPNEIDR GFPLTRPLWD FNTPEGRERL TIYRQALVAG
RKNVPGTDGR PTQFTNKIDR GFPLTRHLWD FNTPEGRERQ TTYHQALVAG

LRGASRRPTN LAKVREVMQG PNEPPSVFLE RLMEAFRRFT PFDPTSEAQK
LRGASRRPTN LAKVREVMQG PNEPPSVFLE RLMEAFRRFT PFDPTSEAQK
LRGASRRPTN LAKVREVMQG PNEPPSVFLE RLMEAFRRFT PFDPTSEAQK
LRGASRRPTN LAKVREVMQG PNEPPSVFLE RLMEAFRRFT PFDPTSEAQK
LRGASRRPTN LAKVREVMQG PNEPPSVFLE RLMEAFRRFT PFDPTSEAQK
LRGASRRPTN LAKVREVMQG PNEPPSVFLE RLMEAFRRFT PFDPTSEAQK
LRGASRRPTN LAKVREVMQG PNEPPSVFLE RLMEAFRRFT PFDPTSEAQK
LRGASRRPTN LAKVREVMQG PNEPPSVFLE RLMEAFRRFT PFDPTSEAQK
LRGASRRPTN LAKVREVMQG PNEPPSVFLE RLMEAFRRFT PFDPTSEAQK
LRGASRRPTN LAKVREVMQG PNEPPSVFLE RLMEAFRRFT PFDPTSEAQK
LRGASRRPTN LAKVREVMQG PNEPPSVFLE RLMEAFRRFT PFDPTSEAQK
LRGASRRPTN LAKVREVMQG PNEPPSVFLE RLMEAFRRFT PFDPTSEAQK
LRGASRRPTN LAKVREVMQG PNEPPSVFLE RLMEAFRRFT PFDPTSEAQK
LRGASRRPTN LAKVREVMQG PNEPPSVFLE RLMEAFRRFT PFDPTSEAQK
LRGASRRPTN LAKVREVMQG PNEPPSVFLE RLMEAFRRFT PFDPTSEAQK
LRGASRRPTN LAKVREVMQG PNEPPSVFLE RLMEAFRRFT PFDPTSEAQK
LRGASRRPTN LAKVREVMQG LNEPPSVFLE RLMEAFRRFT PFDPTSEAQK
LRGASRRPTN LAKVREVMQG PNEPPSVFLE RLMEAFRRFT PFDPTSEAQK
LRGASRRPTN LAKVREVMQG PNEPPSVFLE RLMEAFRRFT PFDPTSEAQK
LRGASRRPTN LAKVREVMQG PNEPPSVFLE RLMEAFRRFT PFDPTSEAQK
LRGASRRPTN LAKVREVMQG PNEPPSVFLE RLLEAFRRYT PFDPTSEAQK
LRGASRRPTN LAKVREVMQG PNEPPSVFLE RLLEAFRRYT PFDPTSEAQK

LRGASRRPTN LAKVREVMQG PNEPPSVFLE RLLEAFRRYT PFDPTSEAQK
LRGASRRPTN LAKVREVMQG PNEPPSVFLE RLLEAFRRYT PFDPTSEAQK
LRGASRRPTN LAKVREMMQG PNEPPSVFLE RLLEAFRRYT PFDPTSEAQK
LQGASRRPTN LAKVREVMQG PNEPPSVFLE RLMEAFRRFT PFDPTSEAQK
LRGASRR-TN LAKVREVMQG PNEPPSVFLE RLMEAFRRFT PFDPTSEAQK
LRGASRRPTN LAKVREVMQG PNKPPSVFLE RLMEAFRRFT PFDPTSEAQK
LRGASRRPTN LAKVREVMQG PNKPPSVFLE RLMEAFRRFT PFDPTSEAQK
LRGASRRPTN LAKVREVMQG PNEPPSVFLE RLMEAFRRFT PFDPTSEAQK
LRGASRRPTN LAKVREVMQG PNEPPSVFLE RLMEAFRRFT PFDPTSEAQK
LRGASRRPTN LAKVREVMQG PNEPPSVFLE RLMEAFRRFT PFDPTSEAQK
LRGASRRPTN LAKVREVMQG PNEPPSVFLE RLMEAFRRFT PFDPTSEAQK
LRGASRRPTN LAKVREVMQG PNEPPSVFLE RLMEAFRRFT PFDPTSEAQK
LRGASRRPTN LAKVREVMQG PNEPPSVFLE RLMEAFRRFT PFDPTSEAQK
LRGASRRPTN LAKVREVMQG PNEPPSVFLE RLMEAFRRFT PFDPTSEAQK
LRGASRRPTN LAKVREVMQG PNEPPSVFLE RLMEAFRRFT PFDPTSEAQK
LRGASRRPTN LAKVREVMQG PNEPPSVFLE RLMEAFRRFT PFDPTSEAQK
LRGASRRPTN LAKVREVMQG PNEPPSVFLE RLMEAFRRFT PFDPTSEAQK
LRGASRRPTN LAKVREVMQG ANEPPSVFLE RLMEAFRRFT PFDPTSEAQK
LRGASRRPTN LAKVREVMQG PNEPPSVFLE RLMEAFRRFT PFDPTSEAQK
LRGASRRPTN LAKVREVMQG PNEPPSVFLE RLMEAFRRFT PFDPTSEAQK
LRGASRRPTN LAKVREVMQG PNEPPSVFLE RLMEAFRRFT PFDPTSEAQK
LRGASRRPTN LAKVREVMQG PNEPPSVFLE RLMEAFRRFT PFDPTSEAQK
LRGASRRPTN LAKVREVMQG PNEPPSVFLE RLMEAFRRFT PFDPTSEAQK
LRGASRRPTN LAKVREVMQG PNEPPSVFLE RLMEAFRRFT PFDPTSEAQK
LRGASRRPTN LAKVREVMQG PNEPPSVFLE RLMEAFRRYT PFDPTSEAQK
LRGASRRPTN LAKVREMMQG PNEPPSVFLE RLMEAFRRYT PFDPTSEAQK
LRGASKRSTN LAKVREVMQG PKEPPSVFLE RLMEAFRRFT PFDPTSEAQK
LRGASKRSTN LAKVREVMQG PKEPPSVFLE RLMEAFRRFT PFDPTSEAQK
LRGASRRPTN LAKVREVKQG PNEPPSVFLE RLMEAFRRFT PFDPTSEAQK
LRGASRRPTN LAKVREVMQG PNEPPSVFLE RLMEAFRRFT PFDPTSEAQK
LRGASRWPTN LAKVREVMQG PNEPPSVFLE RLMEAFRRFT PFDPTSEAQK
LRGASRRPTN LAKVREMMQG PHEPPSVFLE RLMEAFRRFT PFDPTSEAQK
LRGASRRPTN LAKVREMMQG PHEPP--FLE RLMEAFRRFT PFDPTSEAQK
LRGASRRPTN LAKVREMMQG PHEPP--FLE RLMEAFRRFT PFDPTSEAQK
LRGASRRPTN LAKVREVMQG PNEPPSVFLE RLLEAFRRYT PFDPTSEAQK
LRGASRRPTN LAKVKKVMQG PNEPPSVFLE RLLEAFRRYT PFDPTSEAQK
LRGASKRPTN LAKVK-VMQG PNEPPSVFLE RLLEAFRRYT PFDPTSEAQK
LRGASRRPTN LAKVREVMQG PNEPPSVFLE RIMEAFRWFT PFDPTSEAQK
LWGASRRPTN LAKVREVMQG PNEPPSVFLE RIMEAFRWFT PFDPTSEAQK
LWGASRRPTN LAKVREVMQG PNEPPSVFLE RIMEAFRWFT PFDPTSEAQK
LRGASRRPTN LAKVREVMQG PNEPPSVFLE RLMEAFRRFT PFDPTSEAQK
LWGTSRWPTN LAKVREVMQG LNEPPLVFLE RLMEAFRRFT PFDPTSEAQK
LWGTSRWPTN LAKVREVMQG LNEPPLVFLE RLMEAFRRFT PFDPTSEAQK
LWGASRWPTN LAKVREVMQG LNEPPLVFLE RLMEAFRRFT PFDPTSEAQK

LWGTSRWPTN LAKVREVMQG LNEPPLVFLE RLMEAFRRFT PFDPTSEAQK

LRGTSRWPTN LAKVREVMQG LNEPPLVFLE RLMEAFRRFT PFDPTSEAQK

LRGASRRPTN LAKVREVMQG PNEPPSVFLE RLMEAFRRFT PFDPTSEAQK

LWGASRWPTN LAKVREVMQG PNEPPSVFLE RLMEAFRRFT PFDPTSEAQK

LRGASRRPTN LAKVREVMQG PNEPPSVFLE RLLEAFRRYT PFDPTSEAQK

LRGASRRPTN LAKVREVMQG PNEPPSVFLE RLLEAFRRYT PFDPTSEAQK

LRGASRRPTN LAKVREVMQG PNEPPSVFLE RLLEAFRRYT PFDPTSEAQK

LQGASRRPTN LAKVRKVMQG PNEPPSVFLE RLLEAFRRYT PFDPTSEAQK

LRGASRRPTN LAKVREVMQG PNEPPSVFLE RLLEAFRRYT PFDPTSEAQK

LRGSSRGPTN LAKVREVMQG PNEPPSVFLE RLMEAFRRFT PFDPTSEAQK

LRCASRWPTN LAKVREVMQG QNEPPLVFLE RLMEAFRWFT PFDPTSEAQK

LRCASRWPTN LAKVREVMQG QNEPPLVFLE RLMEAFRWFT PFDPTSEAQK

L-------TN LAKVKEVMQG PNELPSIFLE RLMEAFRRFT PFDPTSEAQK

--------TN LAKVKEVMQG PNELPSIFLE RLMEAFRRFT PFDPTSEAQK

--------TN LAKVKEVMQG PNELPSIFLE RLMEAFRRFT PFDPTSEAQK

L-------TN LAKVKEVMQG PNELPSIFLE RLMEAFRRFT PFDPTSEAQK

L-------TN LAKVKEVMQG PNELPSIFRW RLMEAFRRFT PFDPTSEAQK

--------TN LAKVKEVMQG PNELPSIFLE RLMEAFRRFT PFDPTSEAQK

--------TN LAKVKEVMQG PNELPSIFLE RLMEAFRRFT PFDPTSEAQK

L-------TN LAKVKEVMQG PNELPSIFLE RLMEAFRRFT PFDPTSEAQK

LRGASRRPTN LAKVREVMQG PNEPPSVFLE RLMEAFRRFT TFDPTSEAQK

LRGASRRPTN LAKVREVMQG PNEPPSVFLE RLMEAFRRFT TFDPTSEAQK

LRGASRRPTN LAKVREVMQG PNEPPSVFLE RLMEAFRRFT TFDPTSEAQK

LRGASRRPTN LAKVREVMQG PNEPPSVFLE RLMEAFRRFT TFDPTSEAQK

LRGASRRPTN LAKVREVMQG PNEPPSVFLE RLMEAFRRFT TFDPTSEAQK

LRGASRRPTN LAKVREVMQG PNEPPSVFLE RLMEAFRRFT TFDPTSEAQK

LRGASRRPT- LAKVREVMQG PNEPPSVFLE RLMEAFRRFT PFDPTSEAQK

LRGASRRPTN LTKVKKVIQG PNEPPSVFLE RLLKAFRRYT PFDPTSEAQK

LRGASRRPTN LA--REVMQG PNEPPSVFLE RLMEAFRRFT PFDPTSEAQK

LRGASRRPTN LA--REVMQG PSEPPSVFLE RLMEAFRRFT PFDPTSEAQK

LRGASRRPTN LA--REVMQG PNEPPSVFLE RLMEAFRRFT PFDPTSEAQK

LRGASRRPTN LAKVREVMQG PTESPSMFLE RLMKAFRWFT PFDPTSEIQK

LQDASRRPPR LARVREMMQG PTESPSMFLE RLMEAFWRFT PFDPTSEIQK

LKGAARRPTN LAKVREVLQG PAEPPSVFLE RLMEAYRRYT PFDPSSEGQQ

LRGAARRPTN LAKVREVLQG QTEPPSVFLE RLMEAYRRYT PFDPLSEGQR

LRGAARRPTN LAKVREVLQG QTEPPSVFLE RLMEAYRRYT PFDPLSEGQR

LRGAARRPTN LAKVREVLQG QTEPPSVFLE RLMEAYRRYT PFDPSSEGQK

LRGAARRPTN LAKVREVLQG QTEPPSVFLE RLMEAYRRYT PFDPLSEGQR

LRGAARRPTN LAKVREVLQG QTEPPSVFLE RLMEAYRRYT PFDPLSEGQR

LRGAARRPTN LAKVREVLQG QIEPPSVFLE RLMEAYRRYT PFDPSSEGQR

LKGAARRPTN LAKVREVLQG PTEPPSVFLE RLMEAYRRYT PFDPSSEGQK

LKGAARRPTN LAKVREVLQG PAEPPSVFLE RLMEAYRRYT PFDPSSEGQQ

LKGAARRPTN LAKVREVMQG PTEPPSVFLE RLMEAYRRYT PFDPTSESQQ

LKGAARRPTN LAKVREVMQG PTEPPSVFLE RLMEAYRRYT PFDPTSESQQ

LKGAARRPTN LAKVREVMQG PTEPPSVFLE RLMEAYRRYT PFDPTSESQQ
LKGAARCPTN LSKVREVMQG PTEPPSVFLE RLMEAYRCYT PFDPTSEGQQ
LRGAARQPTN LAKVREVMQG ATEPPSVFLE RLMEAYRRYT PFDPTSEGQR
LRGAARQPTN LAKVREVMQG ATEPPSVFLE RLMEAYRRYT PFDPTSEGQR
LRGAAWQPTN LAKVREVMQG ATEPPSVFLE RLMEAYRRYT PFDPTSEGQR
LRGAARQPTN LAKVREVMQG ATEPPSVFLK RLIEAYKRYT PFDPTSEGQR
LKGVARCPTN LAKIREVMLG ANQPPICF-E RLMEAYRHYT PFDPSSEGQQ
LQGAARRPTN LAKVREVLQG PVEPPSVFLE -LVEAYRRYT PFDPSSEGQQ
LQGAARCPTN LAKVREVPQG PIEPPSVFLE -LKEAYRRYT AFDSSSEGQQ
LKGAARCPTN LAKVREVMQR PAEPPSVFLE RLMEAYRRYT PFDPTAEGQA
LKGAARRPTN LAKVREVLQG PTEAPSLFLE RLLEAYRRYT PFDPTSEGQQ
LHGAARRPTN LAKVREVTQG PQESPTVFLE RLMEAFRRFT PYDPTSEEHR
LKGAGKRPTN LAKVRTIIQG KEESPAAFME RLLEGFRMYT PFTPEAPEHK
LKGAGKRPTN LAKVRTITQG KDESPAAFME RLLEGFRMYT PFDPEAPEHK
LRGAARRPTN LAQVKQVVQG KEETPAAFLE RLKEAYRMYT PYDPEDPGQA
LQNAGRSPTN LAKVKGITQG PNESPSAFLE RLKEAYRRYT PYDPEDPGQE
LQNAGRSPTN LAKVKGITQG PNESPSAFLE RLKEAYRRYT PYDPEDPGQE
LQNAGRSPTN LAKVKGITQG PNESPSAFLE RLKEAYRRYT PYDPEDPGQE
LQNAGRSPTN LAKVKGITQG SNESPSAFLE RLKEAYRRYT PYDPEDPGQE
LRAAARRPTN LAKVKAIMQG DNESPAVFLE RLYDAYRQYT PLDPLAEENQ
LKGALRHPTN LAKVREILQG PVELPSAFLE CLMEAYRRYN PFDPASPGHP
LRGAGKRPTN LAKVREILQE EKEPPAVFLE HLLEAYRCYT PFDPMSEGQQ
LRGAGRCPTN LAKIREILQE GKEPPAVFLE HLLEEQRHYT PFDPTSEGQQ
LKGAARCPTN LAKAKEVLQG PVEPPSVFLE HLMEAYRRCT PFDPLPL-RL
LKGAARLPTN LAKIREVLQG SVELLSVFVE RLLEAYWYFI PFDPTSEGQQ
LRAAARKPTN LSKITEVRQG ADESPTAYLE RLYQAYRTWS PIDPRAPENQ
LRGAACRPTN LAKVREVTQG PTEAPSVFLE RIIDAFR-YT PFDPTSEGQG
LRGAACRPTN LAKVREVTQG PTEAPSVFLE RIIDAFR-YT PFDPTSEGQR
IRGAARYPTN LAKVREVTQG PTEAPSVFLE RIIDAFR-YT SFDPTSEGQR

ASVALAFIGQ SALDIRKKLQ RLEGLQEAEL RDLVKEAEKV YYKRETEEER
ASVALAFIGQ SALDIRKKLQ RLEGLQEAEL RDLVKEAEKV YYKRETEEER
ASVALAFIGQ SALDIRKKLQ RLEGLQEAEL RDLVKEAEKV YYKRETEEER
ASVALAFIGQ SALDIRKKLQ RLEGLQEAEL RDLVKEAEKV YYKRETEEER
ASVALAFIGQ SALDIRKKLQ RLEGLQEAEL RDLVKEAEKV YYKRETEEER
ASVALAFIGQ SALDIRKKLQ RLEGLQEAEL RDLVKEAEKV YYKRETEEER
ASVALAFIGQ SALDIRKKLQ RLEGLQEAEL RDLVKEAEKV YYKRETEEER
ASVALAFIGQ SALDIRKKLQ RLEGLQEAEL RDLVKEAEKV YYKRETEEER
ASVALAFIGQ SALDIRKKLQ RLEGLQEAEL RDLVKEAEKV YYKRETEEER
ASVALAFIGQ SALDIRKKLQ RLEGLQEAEL RDLVKEAEKV YYKRETEEER
ASVALAFIGQ SALDIRKKLQ RLEGLQEAEL RDLVKEAEKV YYKRETEEER
ASVALAFIGQ SALDIRKKLQ RLEGLQEAEL RDLVKEAEKV YYKRETEEER
ASVALAFIGQ SALDIRKKLQ RLEGLQEAEL RDLVKEAEKV YYKRETEEER
ASVALAFIGQ SALDIRKKLQ RLEGLQEAEL RDLVKEAEKV YYKRETEEER

ASVALAFIGQ SALDIRKKLQ RLEGLQEAEL HDLVKEAEKV YYKRETEEER
ASVALAFIGQ SALDIKKKLQ RLEGLQEAEL RDLVKEAEKV YYKRETEEER
ASVALAFIGQ SALDIRKKLQ RLEGLQEAEL RDLVKEAEKV YYKRETEEER
ASMALAFIGQ SALDIRKKLQ RLEGLQEAEL RDLVKEAKKV YYKRETEEER
ASMALAFIGQ SALDIRKKLQ RLEGLQEAEL HDLVKEAEKV YYKRETEEER
ASVALAFIGQ SALNIRKKLQ RLEGLQEAEL RDLVKEAEKV YYKRETEEER
ASVALAFIGQ SALDIRKKLQ RLEGLQEAEL RDLVKEAEKV YYKRETEEER
ASVALAFIGQ SALDIRKKLQ RLEGLQEAEL RDLVKEAEKV YYKRETEEER
ASVALAFIGQ SALDIRKKLQ RLEGLQEAEL RDLVKEAEKV YYKRETEEER
ASVALAFIGQ SALDIRKKLQ RLEGLQEAEL RDLVKEAEKV YYKRETEEER
ASVALAFIGQ SALDIRKKLQ RLEGLQEAEL RDLVKEAEKV YYKRETEEER
ASVALAFIGQ SALDIGKKLQ RLEGLQEAEL RDLVKEAEKV YYKRETEEER
ASVALAFIGQ SALDIRKKLQ RLEGLQEAEL RDLVKEAEKV YYKRETEEER
ASVALAFIGQ SALDIRKKLQ RLEGLQEAKL RDLVKEAEKV YYKRETEEER
ASVALAFIGQ SALDIRKKLQ RLEGLQEAEL RDLVREAEKV YYRRETEEEK
ASVALAFIGQ SALDIRKKLQ RLEGLQEAEL RDLVREAEKV YYRRETEEEK
ASVALAFIGQ SALDIRKKLQ RLEGLQEAEL RDLVREAEKV YYRRETEEEK
ASVALAFIGQ SALDIRKKLQ RLEGLQEAEL RDLVREAEKV YYRRETEEEK
ASVALAFIGQ SALDIRKKLQ RLEGLQEAEL RDLVREAEKV YYRRETEEEK
ASVALAFIGQ SALDIRKKLQ RLEGLQEAEL RDLVREAEKV YYRRETEEEK
ASVALAFIGQ SALDIRKKLQ RLEGLQEAEL RDLVREAEKV YYRRETEEEK
ASVALAFIGQ SALDIRKKLQ RLEGLQEAEL RDLVREAEKV YYRRETEEEK
ASVALAFIGQ SALDIRKKLQ RLEGLQEAEL RDLVREAEKV YYRRETEEEK
ASVALAFIGQ SALDIRKKLQ RLEGLQEAEL RDLVREAEKV YYRRETEEEK
ASVALAFIGQ SALDIRKKLQ RLEGLQEAEL RDLVKEAEKV YYRRETEEEK
ASVALAFIGQ SALDIRKKLQ RLEGLQEAEL RDLVREAEKV YYRRETEEEK
ASVALAFIGQ SALDIRKKLQ RLEGLQEAEL RDLVREAEKV YYRRETEEEK
ASVALAFIGQ SALDIRKKLQ RLEGLQEAEL RDLVREAEKV YYRRETEEEK
ASVALAFIGQ SALDIRKKLQ RLEGLQEAEL RDLVREAEKV YYRRETEEEK
ASVALAFIGQ SALDIRKKLQ RLEGLQEAEL RDLVREAEKV YYRRETEEEK
ASVALAFIGQ SALDIRKKLQ RLEGLQEAEL RDLVKEAEKV YYRRETEEEK
ASVALAFIGQ SALDIRKKLQ RLEGLQEAEL RDLVKEAEKV YYRRETEEEK
ASVALAFIGQ SALDIRKKLQ RLEGLQEAEL RDLVKEAEKV YYRRETEEEK
ASVALAFIGQ SALDIRKKLQ RLEGLQEAEL RDLVKEAEKV YYRSETEEEK
ASVALAFIGQ SALDIRKKLQ RLEGLQEAEL RDLVKEAEKV YYRRETEEEK
ASVALAFVGQ SALDIRKKLQ RLEGLQEAEL RDLVKEAEKV YYRRETEEEK
ASVALAFIGQ SALDIRKKLQ RLEGLQEAEL RDLVKEAEKV YYKRETEEER
ASVALAFIGQ SALDIRRKLQ RLEGLQEAEL RDLVKEAEKV YYKRETEEER
ASVALAFIGQ SALDIRKKLQ RLEGLQEAEL HDLVREAEKV YYRRETEEEK
ASVALAFIGQ SALDIRKKLQ RLEGLQEAEL RDLVREAEKV YYRRETEEEK
ASVALVFIGQ SALDIRKKLQ RLEGLQEAEL RDLVKEAEKV YYRRETEEEK
TSVALAFIGQ SALDIRKKLQ RLEGLQEAEL RDLVREAEKV YYRRETEEEK
ASVALAFIGQ SALDIRKKLQ RLEGLQEAEL RDLVKEAEKV YYKRETEEER
ASMALAFIGQ SALDIRKKLQ RLEGLQEAKL RDLVKEAEKV YYKRETDEER

```
ASMALAFIGQ SALDIRKKLQ RLEGLQEAEL RDLVKEAEKV YYKRETEEER
ASMALAFIGQ AALDIRKKCQ RLEGLQEAEL RDLVKEAEKV YYKRETEEER
ASMALAFIGQ AALDIRKKCQ RLEGLQEAEL RDLVKEAEKV YYKRETEEER
ASMALAFIGQ AALDIRKKCQ RLEGLQEAEL RDLVKEAEKV YYKRETEEER
ASVALAFIGQ SAQDIRRKLQ RLEGLQEAEL HDLVKEAEKV YYKRETEEER
ASVALDFIGQ SALDIRKKLR RLEGLQEA-L HDLVKEADKV YYRRETEEEK
ASVALDFIGQ SALDIRKKLR RLEGLQEA-L HDLVKEADKV YYRRETEEEK
ASVALDFIGQ SALDIRKKLR RLEGLQEA-L HDLVKEADKV YYRRETEEEK
ASVALDFIGQ SALDIRKKLR RLEGLQEA-L HDLVKEADKV YYRRETEEEK
ASVALDFIGQ SALDIRKKLR RLEGLQEA-L HDLVKEADKV YYRRETEEEK
ASVALDFIGQ SALDIRKKLR RLEGLQEA-L HDLVKEADKV YYRRETEEEK
ASMALAFIGQ AALDIRKKCQ RLEGLQEAEL RDLVKEAEKV YYRRETEEEK
ASMALAFIGQ SALDIKKKLQ RLEGLQEAEL RDLVKEAEKV YYKRETEEER
ASMALAFIGQ SALDIKKKLQ RLEGLQEAEL RDLVKEAEKV YYKRETEEER
ASVALAFIGQ SALDIKKKLQ RLKGLQEAEL RDLVKETEKV YYKRETEEER
ASVALAFIGQ SALDIRKKLQ RLEGLQEAEL HDIMKETEKV YYKRETEEER
ASVALAFIGQ SALDIRKKLQ RLEGLQEAEL HDLVKEAEKV YYKRETEEER
ASVALAFIGE LALDIRKKLQ RLEGLQEAEL CDLVKEAEKV YYRRETEEEK
ASGALAFIGQ LALDIRKKLQ RLEGLQEAEL CDLVKEAEKV YYRRETEEEK
ASGALAFIGQ LALDIRKKLQ RLEGLQEAEL RDLVKEAEKV YYKRETEEER
ASVALAFIGQ LAQNIRKKLQ RLEGLQEAEL RDLVKEAEKV YYKRETEEER
ASVALAFIGQ LAQNIRKKLQ RLEGLQEAEL RDLVKEAEKV YYKRETEEER
ASVALAFIGQ LAQNIRKKLQ RLEGLQEAEL RDLVKEAEKV YYKRETEEER
ASVALAFIGQ LAQNIRKKLQ RLEGLQEAEL RDLVKEAEKV YYKRETEEER
ASVALAFIGQ LAQNIRKKLQ RLEGLQEAEL RDLVKEAEKV YYKRETEEER
ASVALAFIGQ LAQNIRKKLQ RLEGLQEAEL RDLVKEAEKV YYKRETEEER
ASVALAFIGQ LAQNIRKKLQ RLEGLQEAEL RDLVKEAEKV YYKRETEEER
ASVALAFIGQ LAQNIRKKLQ RLEGLQEAEL RDLVKEAEKV YYKRETEEER
ASVALAFIGQ SALDIRRKLQ RLEGLQEAEL RDLVKEAEKV YYKRETEEER
ASVALAFIGQ SALDIRRKLQ RLEGLQEAEL RDLVKEAEKV YYKRETEEER
ASVALAFIGQ SALDIRRKLQ RLEGLQEAEL RDLVKEAEKV YYKRETEEER
ASVALAFIGQ SALDIRRKLQ RLEGLQEAEL RDLVKEAEKV YYKRETEEER
ASVALAFIGQ SALDIRRKLQ RLEGLQEAEL RDLVKEAEKV YYKRETEEER
ASVALAFIGQ SALDIRRKLQ RLEGLQEAEL RDLVKEAEKV YYKRETEEER
ASVALAFIGQ -ALDIRKKLQ RLEGLQEAEL RDLVKEAE-V YYKRETEEEE
ASMALAFIGQ STLDIKKKLQ RLKGLQEAEL RDLVKEAEKV YYKRET-KKK
ASMALAFVGQ AALDIRKKRQ RLEGLQEAEL RDLVKEAEKV YYRRETKEEK
ASMALAFVGQ AALDIRKKRQ RLEGLQEAEL RDLVKEAEKV YYRRETKEEK
ASMALAFVGQ AALDIRKK-Q RLEGLQEAEL RDLVKEAEKV YYRRETKEEK
GSVALAFIGQ SASDIRKKLQ RLEGLQEAES CDLVKEAEKV YYKRETEEEK
GSVALAFIG- SAPDIRKKLQ RLEGLQEAEL HDLVKEAEKV YYKRETEEEK
AAVAMAFIGQ SASDIKRKLQ RLEGLQDYTL RDLVKEAEKV YHKRETEEEK
AAVAMAFIGQ SAPDIKKKLQ RLEGLQDHML QDLVKEAEKV YHKRETEEER
ATVAMAFIGQ SAPDIKKKLQ RLEGLQDHTL QDLVKEAEKV YHKRETEEER
```

AAVAMAFIGQ SAPDIKKKLQ RLEGLQDYTL QDLVKEAEKV YHKRETEEER

AAVAMAFIGQ SAPDIKKKLQ RLEGLQDHTL QDLVKEAEKV YHKRETEEER

AAVAMAFIGQ SAPDIKKKLQ RLEGLQDHTL QDLVKEAEKV YHKRETEEER

AAVAMAFIGQ SAPDIKKKLQ RLEGLQDHTL QDLVKEAEKV YHKRETEEER

AAVAMSFIGQ SAPDIKKKLQ RLEGLQDHSL QDLIKEAEKV YHKRETEEEK

AAVAMAFIGQ SAPDIKKKLQ RLEGLQDYSL QDLVKEAEKV YHKRETEEER

ATVAMAFIGQ SALDIKRKLQ RLEELHAMAL QDLVKEAEKV YHKRETEEEK

ATVAMAFIGQ SALDIKRKLQ RLEELHAMAL QDLVKEAEKV YHKRETEEEK

ATVAMAFIGQ SALDIKRKLQ RLEELHAMAL QDLVKEAEKV YHKRETEEEK

ATVAMAFIGQ SASDIKRKLQ RSEGLHAMAL QNLVREAEKM F---------

ASVIMAFIGQ SAPDIRKKLQ RIEGLQDYTI RDVVREAEKV YHRRETEDEK

ASVIMAFIGQ SAPDIRKKLQ RIEGLQDYTI RDVVREAEKV YHRRETEDEK

ASVIMAFIGQ SAPDIRKKLQ RIEGLQDYTI RDVVREAEKV Y---------

ASVIMAFIGQ SAPDIKKKLQ QIEGLQDYTI RDVVREAEKV YHRRKTEDKK

AMGAMTFIGQ SASDIKRKLQ RLKGLHAMAL QDLVREAEKV F---------

AAVTMAFIGQ SASNIKKKLQ RLEGLQDLAL SDLVKEAEKV YHKRETEEEK

AAVAMAFTGH SALDIKKKLQ RLEGLQDLAL SDLVKEAEKV Y---------

AAVAMAFIGQ SA-DIRKKLQ KIVGLQDYTL QDLVKEAEKV YHKRETKEER

AAVAMSFIGQ SYPDI-KRLQ RLEGLQDLTV RDLVKGAEKV YHKRETEEEK

ATIAMAFIDQ AAPDIKKKLQ SLDGLQGFSL QELVKEADKV YNKRETEEEK

ATVAMSFIDQ AASDIKGKLQ RLDGIQTYGL QELVREAEKV YNKRETPEEK

ATVAMSFIDQ AALDIKGKLQ RLDGIQTHGL QELVREAEKV YNKRETPEER

ASVILSFIYQ SSPDIRNKLQ RLEGLQGFTL SDLLKEAEKI YNKRETPEER

TNVSMSFIWQ SAPDIGRKLE RLEDLKNKTL GDLVREAEKI FNKRETPEER

TNVAMSFIWQ SAPDIGRKLE RLEDLKSKTL GDLVREAEKI FNKRETPEER

TNVSMSFIWQ SAPDIGRKLE RLEDLKSKTL GDLVREAEKI FNKRETPEER

TNVSMSFIWQ SAPDIGRKLE RLEDLKNKTL GDLVREAERI FNKRETPEER

SAVIMSFINQ AAPDIRKKLY KQEGLGEMSI RDLMKVAERV FNTRETPEER

TAMIIAFIGL -------KWQ HLEGLQDYTL QDFVKEAEKV YHKRETKEKR

AAVAMAFIGQ SASDIRRRLQ RLESLQTLSL QDPVKEAEKV YHN-------

VAVAMAFIWQ SASDIRRRVQ HLEGLQTLSL QDLVKEADKV YHKSETEEVK

NKLQLWPLYD SQRQIKRKLQ RLEGPQDYTL RDLVKEAEKC FS--------

AAVGGYGLFR AVSIRYEKVT AARRTAGLYL TGFGGGRESV -S---AGDGR

AAIVIQFVSQ SAPDIRKKIQ KIDGFQGKSL SELVAIAQKV FDQREATHEL

ASVTLAFIGQ AAPDIKRKLQ RLEGLQDLSL QDLIKEAEKV FYKRETEEEK

ASVALAFIGQ AAPDIKRKLQ RLEGLQDLSL EDLIKEAEKV FY--------

ASVALAFIGQ AAPDIKRKLQ RLEGLQDLSL QDLVKEAEKV FYKREMEEEK

EQRKERERE REERRNKRQE KNLTK

EQRKERERE REERRNKRQE KNLTK

EQRKERERE REERRNKRQE KNLTK

EQRKERERE REERRNKRQE KNLTK

EQRKERERE REERRNKRQE KNLTK

EQRKERERE REERRNKRQE KNLTK

EQRKEREREE REERRNKRQE KNLTK
EQRKEREREE REERRNKRQE KNLTK
EQRKEREREE REERRNKRQE KNLTK
EQRKEREREE REERRNKRQE KNLTK
EQRKEREREE REERRNKRQE KNLTK
EQRKEREREE REERRNKRQE KNLTK
EQRKKREREE REERRNKRQE KNLTK
EQKKEKEREE REERRNKRQE KNLTK
EQRKEREREE REERRNKRQE KNLTK
EQRKEREREK REERRNKRQE KNLTK
EQRKEREREE REERRNKRQE KNLTK
EQRKEREREE REERRNKRQE KNLTK
EQRKEREREE REERRNKRQE KNLTK
EQRKEREREE REERRNKRQE KNLTK
EQRKEREREE REERRNKRQE KNLTK
EQRKEREREE REERRNKRQE KNLTK
EQRKEREREE REERRNKRQE KNLTK
EQRKEREREE REERRNKRQE KNLTK
EQRKEREREE REERRNKRQE KNLTK
EQRKEREREE REERRNKRQE KNLTK
EQRKEREREE REERRNKRQE KNLTK
EQRKEREREE REERRNKRQE KNLTK
EQRKEKEREE REERRDRRQE KNLTK
EQRKEKEREE REERRDRRQE KNLTK
EQRKEKEREE REERRDRRQE KNLTK
EQRKEKEREE REERRDRRQE KNLTK
EQRKEKEREE REERRDRRQE KNLTK
EQRKEKEREE REERRDRRQE KNLTK
EQRKEKEREE REERRDRRQE KNLTK
EQRKEKEREE REERRDRRQE KNLTK
EQRKEKEREE REERRDRRQE KNLTK
EQRKEKEREE REERRDRRQE KNLTK
EQRKEKEREE REERRDRRQE KNLTK
EQRKEKEREE REERRDRRQE KNLTK
EQRKEKEREE REERHDRRQE KNLTK
EQRKEKEREK REKRRDRRQE KNLTK
EQRKEKEREE REERRDRRQE KNLTK
EQRKEKEREE REERRDRRQE KNLTK
EQRKEKEKEE REERCDRQQE KNLTK
EQRKEKEKEE REERCDRQQE KNLTK
EQRKEKEREE REERRDRRQE KNLTK
EQRKEKEREE REERRDRRQE KNLTK
EQRKEKEREE REERRDRRQE KNLTK
EQRKEKEREE REERRDRRQE KNLTK

EQRKEREREE REERRNKRQE KNLTK
EQKKEREKEK REERRDRRQE KNLTK
EQRKEKEREE REERRDRRQE KNLTK
EQRKEKEKEE REERRDRQQE KNLTK
EQRKEKEKEE REERCDRQQE KNLTK
EQRKEKEREE REERRDRRQE KNLTK
EQRKEREREE REERRNKRQE KNLTK
EQRKEREREE REERRNKRQE KNLTK
EQKKEREKEE REKRRNKRQE KNLTK
EQRKEREREE REERRNKRQE KNLTK
EQKKAREREE REERRNKRQE KNLTK
EQKKARERE- -EERRNKRQE K-LTK
EQRKEREREE REERRDRRQE KNLTK
EPRKEKEREE REERRDRRQE KNLTK
EPRKEKEREE REERRDRRQE KNLTK
EPRKEKEREE REERRDRRQE KNLTK
EPRKEKEREE REERRDRRQE KNLTK
EQSKEKEREE REERRDRRQE KNLTK
EPRKEKEREE REERRDRRQE KNLTK
EQRKEKEREE REERRDRWQE KNLTK
EQKKER-KKE KKKRRNKRQE KNLTK
EQKKER-KKE KKKRRNKRQE KNLTK
EQKKERKKEE REKRRNKRQE KNLTK
EQKKEREKEE REERRNKRQE KNLTK
KQRK-KKREK REKRRNKWQE KNLTK
EQRKEKEREE REERRDRRQE KNLTK
EQRKEKEREE REERRDRRQE KNLTK
EQRKEREREE REERRNKRQE KNLTK
EQRKEREREK REKRRDRR-E KNLTK
EQRKEREREK REKRRDRR-E KNLTK
EQRKEREREK REKRRDRR-E KNLTK
EQRKEREREK REKRRDRR-E KNLTK
EQRKEREREK REKRRDRR-E KNLTK
EQRKEREREK REKRRVPG-E KNLTK
EQRKEREREK REKRRDRR-E KNLTK
EQRKEREREE REERRNKRQE KNLTK
EQRKER---E KEERRDRRQE KNLTK
EQRKER---E KEERRDRRQE KNLTK
EQRKER---E KEERRDRRQE KNLTK
EQRKER---E KEERRDRRQE KNLTK
EQRKER---E KEERRDRRQE KNLTK
EQRKER---E KEERRDRRQE KNLTK
QRKK---REE REERR-IRQE KNLTK
GTKKEREREE REKRHNKQQE KNLTK

```
EQRKEKEREE REERRYRWQE KNLTK
EQRKEKEREE REERRYRWQE KNLTK
EQRKEKEREE REERRYRWQE KNLTK
EQRKEREREE REDRRNRRQK KNLNR
EQRKEREREE RKDRRNRRQE KNLTR
QEREKKEAEE RENRRDRRQE RNLTK
QEREKKEIEE RENRRDRRQE RNLSK
QEREKKEMEE RENRRDRRQE RNLSK
QEREKKEVEE RENRRDRRQE RNLSK
QEREKKEMEE RENRRDRRQE RNLSK
QEREKKEMEE RENRRDRRQE RNLSK
QEREKKEMEE RENRRDRRQE RNLSK
QEREKKETEE RERRRDRRQE KNLTK
QEREKKEAEE KERRRDRPKK KNLTK
REREKREEEE REKQKERRQE RNLSR
REREKREEEE REKQKERRQE RNLSR
REREKREEEE REKQKERRQE RNLSR
---------- ---------- -NLSR
LEREKREKRE EEDRRDRRQE KVLTR
LEREKREKRE EEDRRDRRQE KVLTR
---------- ---------- --LTR
LEREKREKRE EEDRRDRRQE KVLTM
---------- ---------- -NLSR
KGK--KEAKK REEKRDRRQE RNLTR
---------- ---------- --LTR
MEREKREKKE KT--KGKEKE KRLTK
REREKKEREE REDERDRCRD QNLTR
EERKQKEQEA REIRRDKRQE RNLSK
EARLAKEQEA REERRDRKRD KHLTK
EARLIKEQEE REDRRDRKRD KHLTK
EERLWQRQEE ----RDKKRH KEMTK
EERIRRETEE KEERRDRRRH REMSK
EERIRRETEE KEERRDRRRH REMSK
EDRIRRETEE KEERRDRRRH REMSK
EERIKRETEE KEERRDRRRQ REMSK
EDRIRKENQE LQERINRRQQ REMAK
QEREKREAEE REEQRDHRQT RNLTR
---------- ---------- --LAR
RERV-KEAEA RE-SRYRRQE RNLRR
---------- ---------- --LTR
EARRRKER-G RKRRQDRRQE RNLTR
TRKMAKAQES RAERGSKKTP PGKGR
EQRREKEREE REEKHDRKRE KNLSK
---------- ---------- -NLSK
```

| | |
|---|---|
| | EQRKEKEREE REEKCDRKWE KNLSK |
| Pol | |
| 116 567 | |
| AOCR01031838.1 | EGIRPHVQRL IQQGILVPVQ SPWNTPLLPV RKPGTNDYRP VQDLREVNKR |
| AOCR01107158.1 | EGIRPHVQRL IQQGILVPVQ SPWNTPLLPV RKPGTNDYRP VQDLREVNKR |
| AOCR01098665.1 | EGIRPHVQRL IQQGILVPVQ SPWNTPLLPV RKPGTNDYRP VQDLREVNKR |
| PERV-C | EGIRPHVQRL IQQGILVPVQ SPWNTPLLPV RKPGTNDYRP VQDLREVNKR |
| CM000825.5_b | EGIRPHVQRL IQQGILVPVQ SPWNTPLLPV RKPGTNDYRP VQDLREVNKR |
| LUXS01031219.1 | EGIRPHVQRL IQQGILVPVQ SPWNTPLLPV RKPGTNDYRP VQDLREVNKR |
| LUXX01045907.1 | EGIRPHVQRL IQQGILVPVQ SPWNTPLLPV RKPGTNDYRP VQDLREVNKR |
| LT634572.1_d | EGIRPHVQRL IQQGILVPVQ SPWNTPLLPV RKPGTNDYRP VQDLREVNKR |
| LUXX01006697.1 | EGIRPHVQRL IQQGILVPVQ SPWNTPLLPV RKPGTNDYRP VQDLREVNKR |
| LUXQ01071671.1 | EGIRPHVQRL IQQGILVPVQ SPWNTPLLPV RKPGTNDYRP VQDLREVNKR |
| CM000816.5_b | EGIRPHVQRL IQQGILVPVQ SPWNTPLLPV RKPGTNDYRP VQDLREVNKR |
| PERV-A | EGIRPHVQRL IQQGILVPVQ SPWNTPLLPV RKPGTNDYRP VQDLREVNKR |
| LUXX01090830.1 | EGIRPHVQRL IQQGILVPVQ SPWNTPLLPV RKPGTNDYRP VQDLREVNKR |
| LUXU01038095.1 | EGIRPHVQRL IQQGILVPVQ SPWNTPLLPV RKPGTNDYRP VQDLREVNKR |
| LUXV01020898.1 | EGIWPHVQRL IQQGILVPVQ SPWNTPLLPV RKPGTNDYRP VQDLREVNKR |
| CM000812.5_b | EGIRPHVQRL IQQGILVPVQ SPWNTPLLPV RKPGTNDYRP VQDLREVNKR |
| CM000826.5_c | EGIRPHVQRL IQQGILVPVQ SPWNTPLLPV RKPGTNDYRP VQDLREVNKR |
| CM000823.5_b | EGIRPHVQRL IQQGILVPVQ SPWNTPLLPV RKPGTNDYRP VQDLREVNKR |
| AOCR01000130.1 | EGIQPHVQRL IQQGILVPVQ SPWNTPLLPV RKPGTNDYRP VQDLREVNKR |
| LUXW01074277.1 | EGIRPHVQRL IQQGILVPVQ SPWNTPLLPV RKPGTNDYRP VQDLREVNKR |
| LUXS01060324.1 | EGIQPHVQRL IQQGILVPVQ SPWNTPLLPV RKPGTNDYRP VQDLREVNKR |
| AOCR01079940.1 | EGIRPHVQRL IQQGILVPVQ SPWNTPLLPV RKPGTNDYRP VQDLREVNKR |
| AEMK02000197.1 | EGIWPHVQRL IQQGILVPVQ SPWNTPLLPV RKPGTNDYRP VQDLREVNKR |
| LUXR01004647.1 | EGIWPHVQRL IQQGILVPVQ SPWNTPLLPV RKPGTNDYRP VQDLREVNKR |
| LUXR01088996.1 | EGIWPHVQRL IQQGILVPVQ SPWNTPLLPV RKPGTNDYRP VQDLREVNKR |
| CM000815.5 | EGIWPHVQRL IQQGILVPVQ SPWNTPLLPV RKPGTNDYRP VQDLREVNKR |
| CM000822.5_b | EGIWPHVQRL IQQGILVPVQ SPWNTPLLPV RKPGTNDYRP VQDLREVNKR |
| CM000820.5 | EGIWPHVQRL IQQGILVPVQ SPWNTPLLPV RKPGTNDYRP VQDLREVNKR |
| PERV-B | EGIWPHVQRL IQQGILVPVQ SPWNTPLLPV RKPGTNDYRP VQDLREVNKR |
| CM000814.5_c | EGIWPHVQRL IQQGILVPVQ SPWNTPLLPV RKPGTNDYRP VQDLREVNKR |
| CM000826.5_a | EGIWPHVQRL IQQDILVPVQ SPWNTPLLPV RKPGTNDYRP VQDLREVNKR |
| CM000827.5 | EGIWPHVQRL IQQGILAPVQ SPWNTPLLPV RKPGTNDYRP VQDLREVNKR |
| CM000825.5_a | EGIWPHVQRL IQQGILVPVQ SPWNTPLLPV RKPGTNDYRP VQDLREVNKR |
| CM000819.5_a | EGIWPHVQRL IQQGILVPVQ SPWNTPLLPV RKPGTNDYRP VQDLREVNKR |
| CM000824.5_b | EGIWPHVQRL IQQGILVPVQ SPWNTPLLPV RKPGTNDYRP VQDLREVNKR |
| AEMK02000137.1 | EGIWPHVQRL IQQGILVPVQ SPWNTPLLPV RKPGTNDYRP VQDLREVNKR |
| CM000828.5_a | EGIRPHVQKL IQQDILVPVQ SPWNTPLLPV RKPGTNDYRP VQDLREVNKR |
| AEMK02000476.1 | EGIWPHVQRL IQQGILVPVQ SPWNTPLLLV RKPGTNDYRP VQDLREVNKR |
| CM000824.5_a | EGIRSHVQRL IQQGILVPVQ SPWNTPLLPV RKPGTNDYRP VQDLREVNKR |
| CM000819.5_b | EGIRPHVQRL IQQGILVPVQ SPWNTPLLPV RKPGTNDYRP VQDLREVN-R |
| CM000830.5_d | EGIRPHVQRL IQQGILVPVQ SPWNTPLLPV RKPGTNDYRP VQDLREVNKR |

| | | | | | |
|---|---|---|---|---|---|
| LT634572.1_b | EGIWPHVQRL | IQQGILVPVQ | SPWNTPLLPV | RKPGTNDYRP | VQDLREVNKR |
| CM000828.5_b | EGIWPHVQRL | I-QGILVPIQ | SPWNTPLLPV | RKPGTNDYRP | VQDLREVNKR |
| LT634572.1_a | EGIWPHVQRL | IQQGILVPVQ | SPWNTPLLPV | RKPGTNDYRP | VQDLREVNKR |
| LUXY01101100.1 | EGIWPHVQRL | IQQGILVPVQ | SPWNTPLLPV | RKPGTNDYRP | VQDLREVNKR |
| LUXQ01076544.1 | EGIWPHVQRL | IQQGILVPVQ | SPWNTPLLPV | RKPGTNDYRP | VQDLREVNKR |
| CM000828.5_c | EGIWPHVQRL | IQQGILVPVR | SP-NTPLLLV | RKPGTNDYRP | VQDLREVNKR |
| LIDP01000017.1 | EGIWPHVQRL | IQQGILVPVR | SP-NTPLLPV | RKPGTNDYRP | VQDLREVNKR |
| CM000814.5_b | EEIWPHVQRL | IQQGILVPVR | SPWNTPLLPV | RKPGTNDYRP | VQDLREVNKR |
| CM000813.5 | EGIRPHVQRL | IQQGILVPVQ | SPWNTPLLPV | RKPGTNDYRP | VQDLREVNKW |
| AJKK01232762.1 | EGIQPHVQRL | IQQGILVPVQ | SPWNTPLLPV | RKPGTNDYRP | VQDLREVNKR |
| CM000814.5_a | EGIRPHVQRL | IQQGILVPVQ | SP-NTPLLPV | RKPGTNDYRP | VQDLREVNKR |
| LUXY01013808.1 | EGIWPHVQRL | IQQGILVPVQ | SPWNTPLLPV | RKPGTNDYRP | VQDLREVNKR |
| AEMK02000393.1_a | EGIWPHVQRL | IQQGILVPVR | SPWNTPLLPV | RKPGTNDYRP | VQDLREVNKR |
| LUXW01069599.1 | EGIWPHVQRL | IQQGILVPVR | SPWNTPLLPV | RKPGTNDYRP | VQDLREVNKR |
| AEMK02000133.1 | EGIRPHVQRL | IQQGILVPVQ | SPWNTPLLPV | RKPGTNDYRP | VQDLREVNKW |
| CM000818.5_b | EGIWPHVQRL | IQQGILVPVQ | SPWNTPLLPV | RKPGTNDYCP | VQDLREVNKR |
| eJJRV_NW_004504375.1 | TGIWPHILRL | IQQGILVPIQ | SPWNTPLLPV | RKPGTNDYCP | VQDLREVNKR |
| MMERV_21 | EGIRPQINKL | LQQGILVPCK | SPWNTPLLPV | KKPGTRDFRP | VQDLREVNKR |
| MMERV_22 | EGIRPQINKL | LQQGILVPCK | SPWNTPLLPV | KKPGTRDFRP | VQDLREVNKR |
| MMERV_10 | EGIHPQINKL | LQQGILVPCK | SPWNTPLLPV | KKPGTRDFHT | VQDLREVNKR |
| MMERV_3 | EGIRPQINKL | LQQGILVPCK | SPWNTPLLPV | KKPGTRDYRP | VQDLREVNKR |
| MMERV_1 | EGIRPHIQRL | LGQGVLVACQ | SPWNTPLLPV | RKPGTNDYRP | VQDLWEVNKR |
| MMERV_11 | EGIRPHIQRL | LGQGVLVACQ | SPWNTPLLPV | RKPGTNDYRP | VQDLREVNKR |
| MMERV_27 | EGIRPHIQRL | LGQGVLVACQ | SPWNTPLLPV | RKPGTNDYRP | VQDLREVNKR |
| MMERV_5 | EGIRPHIQRL | LDQGVLVACQ | SPWNTPLLPV | RKPGTNDYRP | VQDLREVNKR |
| MMERV_35 | EGIQPHIQRL | LGQGVLVACQ | SPWNTPLLPV | QKPGTNDYRP | VQDLREVNKR |
| MMERV_18 | EGIRPHIQRL | LDQGVLVACQ | SPWNTPLLPV | RKPGTNDYRP | VQDLREVNKR |
| MDEV | EGIRPHIRRL | LDQGILVACQ | SPWNTPLLPV | RKPGTNDYRP | VQDLREVNKR |
| RNERV_6 | EGIRPHIQRL | LQLGILVPCQ | SPWNTPLLPV | RKPGTNDYRP | VQDLREVNKR |
| KoRV | EGIRPHIQRF | LDLGILVPCQ | SPWNTPLLPV | KKPGTNDYRP | VQDLREVNKR |
| GALV | EGIRPHIQKF | LDLGVLVPCR | SPWNTPLLPV | KKPGTNDYRP | VQDLREINKR |
| MMERV_31 | DGIRPHIQRL | LQLGILVPCQ | SPWNTPLLPV | KKPGTSDYRP | VQDLREVNKR |
| MMERV_9 | DGIRPHIQRL | LQLGILVPCQ | SPWNTPLLPV | KKPGTSDYRP | VQDLREVNKR |
| musMus5656 | DGIRPHIQRL | LQLGILVPCQ | SPWNTPLLPV | KKPGTSDYRP | VQDLREVNKR |
| MMERV_39 | DGIRPHIQRL | LQLGILVPCQ | SPWNTPLLPV | KKPGTSDYRP | VQDLREVNKR |
| MMERV_8 | DGIRPHIQRL | LQLGILVPCQ | SPWNTPLLPV | KKPGTSDYRP | VQDLREVNKR |
| MCERV_7 | DGIRPHIQRL | LQQGILVPYQ | SPWNTPLLLV | KKPGSGDYSP | VQDLREVNKR |
| eJJRV_NW_004504334.1 | TGIWPHIQRL | IQQGILVPVQ | SPWNTPLLPV | RKPGTNDYHP | VQDLKK-NKR |
| ratNor2288 | EGIRPHINKF | LQLGILTPCQ | SPWNTPLLPV | KKPGTQDYRP | VQDLREVNKR |
| musMus7492 | EGIRPHIQRL | IELGVLAPCQ | SSWNTPLLPV | KKPDTNDYQP | VQDLREVNKR |
| ratNor1430 | DGI-PHIQRL | LSLGILVPCQ | SP-NIPLLPV | RKPGTNDYRP | VQDLREVNKR |
| ratNor293 | EGIRPHIQRL | LAQGILVPCK | SPWNTPLLPV | KKPGTSDYQP | VQDLQEVNKR |
| ePCRV_KN680906.1 | EGIKPHIQRL | LDLGILIKCQ | SPWNTPLLPV | KKPGTGDYHP | VQDLREVNKR |
| ePCRV_KN676905.1 | SGIKPHIQRL | LDLGILIRYQ | SPWNTPLLPV | KKPGTGDYRP | VQDLRGISKR |

| | | | | | |
|---|---|---|---|---|---|
| ePCRV_KN676491.1 | TGIKPHIQRL | LNLGILIRCQ | SPWNTPLLPV | KKPGTGDYRP | VQDLREVNKR |
| MMERV_13 | EGICPHITRL | HQQGFLVP-K | SP-NTPLLPV | KKLGTNNYRP | VQDLREVNKR |
| MPERV_4 | EGIRPHILRL | LQLGILVPCQ | SPWNTPLLPV | KKPGTRDYRP | VQDLREVNKR |
| MMERV_6 | LGIKPHIQRL | LDQGILVPCQ | SPWNTPLLPV | KKPGTNDYRP | VQDLREVNKR |
| MMERV_16 | LGIKPHIQRL | LDQGILVPCQ | SPWNTPLLPV | KKPGTNDYRP | VQDLREVNKR |
| MMERV_26 | LGIKPHIQRL | LDQGILVPCQ | SPWNTPLLPV | KKPGTNDYRP | VQDLREVNKR |
| MMERV_32 | LGIKPHIQRL | LDQGILVPCQ | SPWNTPLLPV | KKPGTNDYRP | VQDLREVNKR |
| MMERV_12 | LGIKPHIQRL | LDQGILVPCQ | SPWNTPLLPV | KKPGTNDYRP | VQDLREVNKR |
| MMERV_15 | LGIKPHIQRL | LDQGVLVPCQ | SPWNTPLLPV | KKPGTNDYRP | VQDLRKVNKR |
| M-CRV | LGIKPHIQRL | LDQGILVPCQ | SPWNTPLLPV | KKPGTNDYRP | VQDLREVNKR |
| R-MuLV | LGIKPHIQRL | LDQGILVPCQ | SPWNTPLLPV | KKPGTHDYRP | VQDLREVNKR |
| F-MuLV | LGIKPHIQRL | LDQGILVPCQ | SPWNTPLLPV | KKPGTNDYRP | VQDLREVNKR |
| M-MuLV | LGIKPHIQRL | LDQGILVPCQ | SPWNTPLLPV | KKPGTNDYRP | VQDLREVNKR |
| RNERV_3 | QGMRPHITRL | LDQGILVPCR | SPWNTPLLPV | KKPGTGDYRP | VQDLREVNKR |
| RNERV_2 | QGMRPHITRL | LDQGILVPCR | SPWNTPLLPV | KKPGTGDYRP | VQDLREVNKR |
| FeLV | QGIKPHIRRM | LDQGILKPCQ | SPWNTPLLPV | KKPGTEDYRP | VQDLREVNKR |
| RD114 | MGIQPHITRF | LELGVLRPCR | SPWNTPLLPV | KKPGTRDYRP | VQDLREVNKR |
| BaEV | MGIRQHIIKF | LELGVLRPCR | SPWNTPLLPV | KKPGTQDYRP | VQDLREINKR |
| OOEV | DGIRPHIQRL | LELGILVRCQ | STWNTPLLPV | KKPGTNDYQP | VQDLREVNKR |
| ratNor338 | DGIRPHIQ-L | LSLGILVPRQ | LPWNTPLLPV | RKPGTNDYRT | VQDLREVNKR |
| AEMK02000536.1 | EGIRPHVQRL | IQQGILVPVQ | SPWNTPLLPV | RKPGTNDYRP | VQDLREVNKR |
| MCERV_1 | RDQTPHPESL | TRY------- | --FDTLPVPM | EHPPPASQKA | RDKLQTSTRR |
| ratNor4542 | EGIRPNIQRL | LQQGILVPCQ | SPWNTLLLPV | KKPGTNNYRP | VQDLKEVNMR |
| RfRV | KGIAPHINRL | LEAGILKPCH | SAWNTPLLPV | KKPGGKDYRP | VQDLREVNKR |
| MCERV_6 | DGMRPHIQKL | LQLDILVPCQ | SPWNTPLLLV | KKPGTSDYPP | VQDLREVNKX |
| REV | RSLRETIHKF | RAAGILRPVH | SPWNTPLLPV | RKSGTSEYRM | VQDLREVNKR |
| musMus10244 | DRIRPYIHRL | LQLCILVPFL | SPWNTPLLLA | KKPGTSDYRP | AQDLREVNKR |
| musMus10318 | DRIRPHIQRL | LQLGILVPFQ | SPWNTPLLLA | KKPGTSDYRP | AQDLREVNKR |
| musMus9012 | EGIRPHIQRI | LELGVLVPCQ | SPWNTLLLLV | KKPGTNDNQP | VQSLKR--KR |
| musMus9830 | SAERPHYSQL | KSQALLFICQ | ---------- | -------YKT | IK----VNER |
| musMus9837 | EGIRPYIQRL | IELGVLVPCQ | SPWNTPLLPV | KKPGTN---P | VQDLIRVNER |

| | | | | | |
|---|---|---|---|---|---|
| | VQDIHPTVPN | PYNLLCALPP | QRSWYTVLDL | KDAFFCLRLH | PTSQPLFAFE |
| | VQDIHPTVPN | PYNLLCALPP | QRSWYTVLDL | KDAFFCLRLH | PTSQPLFAFE |
| | VQDIHPTVPN | PYNLLCALPP | QRSWYTVLDL | KDAFFCLRLH | PTSQPLFAFE |
| | VQDIHPTVPN | PYNLLCALPP | QRSWYTVLDL | KDAFFCLRLH | PTSQPLFAFE |
| | VQDIHPTVPN | PYNLLCALPP | QRSWYTVLDL | KDAFFCLRLH | PTSQPLFAFE |
| | VQDIHPTVPN | PYNLLCALPP | QRSWYTVLDL | KDAFFCLRLH | PTSQPLFAFE |
| | VQDIHPTVPN | PYNLLCALPP | QRSWYTVLDL | KDAFFCLRLH | PTSQPLFAFE |
| | VQDIHPTVPN | PYNLLCALPP | QRSWYTVLDL | KDAFFCLRLH | PTSQPLFAFE |
| | VQDIHPTVPN | PYNLLCALPP | QRSWYTVLDL | KDAFFCLRLH | PTSQPLFAFE |
| | VQDIHPTVPN | PYNLLCALPP | QRSWYTVLDL | KDAFFCLRLH | PTSQPLFAFE |
| | VQDIHPTVPN | PYNLLCALPP | QRSWYTVLDL | KDAFFCLRLH | PTSQPLFAFE |
| | VQDIHPTVPN | PYNLLCALPP | QRSWYTVLDL | KDAFFCLRLH | PTSQPLFAFE |

VQDIHPTVPN PYNLLCALPP QWSWYTVLDL KDVFFCLRLH PTSQPLFAFE
VQDIHPTVPN PYNLLCALPP QWSWYTVLDL KDVFFCLRLH PTSQPLFAFE
VQDIHPTVPN PYNLLCALPP QWSWYTVLDL KDVFFCLRLH PTSQPLFAFE
VQDIHPTVPN PYNLLCALPP QRSWYTVLDL KDAFFCLRLH PTSQPLFAFE
VQDIHPTVPN PYNLLCALPP QRSWYTVLDL KDAFFCLRLH PTSQPLFAFE
VQDIHPTVPN PYNLLCALPP QRSWYTVLDL KDAFFCLRLH PTSQPLFAFE
VQDIHPTVPN PYNLLCALPP QRSWYTVLDL KDAFFCLRLH PTSQPLFAFD
VQDIHPTVPN PYNLLCALPP QRSWYTVLDL KDAFFCLRLH PTSQPLFAFE
VQDIHPTVPN PYNLLCALPP QRSWYTVLDL KDAFFCLRLH PTSQPLFAFD
VQDIHPTVPN PYNLLCALPP QRSWYTVLDL KDAFFCLRLH PTSQPLFAFE
VQDIHPTVPN PYNLLSALPP ERNWYTVLDL KDAFFCLRLH PTSQPLFAFE
VQDIHPTVPN PYNLLSALPP ERNWYTVLDL KDAFFCLRLH PTSQPLFAFE
VQDIHPTVPN PYNLLSALPP ERNWYTVLDL KDAFFCLRLH PTSQPLFAFE
VQDIHPTVPN PYNLLSALPP ERNWYTVLDL KDAFFCLRLH PTSQPLFAFE
VQDIHPTVPN PYNLLSALPP ERNWYTVLDL KDAFFCLRLH PTSQPLFAFE
VQDIHPTVPN PYNLLSALPP ERNWYTVLDL KDAFFCLRLH PTSQPLFAFE
VQDIHPTVPN PYNLLSALPP ERNWYTVLDL KDAFFCLRLH PTSQPLFAFE
VQDIHPTVPN PYNLLSALPP ERNWYTVLDL KDAFFCLRLH PTSQPLFAFE
VQDIHPTVPN PYNLLSALPP ERNWYTVLDL KDAFFCLRLH PTSQPLFAFK
VQDIHPTVPN PYNLLSALPP ERNWYTVLDL KDAFFCLRLH PTSQPLFAFE
VQDIHPTVPN PYNLLSALPP KRNWYTVLDL KDAFFCLRLH PTSQPLFAFE
VQDIHPTVPN PYNLLSALPP ERNWYTVLDL KDAFFCLRLH PTSQPLFAFE
VQDIHPTVPN PYNLLSALPP ERNWYTVLDL KDAFFCLRLH PTSQPLFAFE
VQDIHPTVPN PYNLLSALPP ERNWYTVLDL KDAFFCLRLH PTSQPLFAFE
VQDIHPTVPN PYNLLCALPP QRSWYTVLDL KDAFFCLRLH PTSQPLFAFE
VQDIHPTVPN PYNLLSALPP ERNWYTVLDL KDAFFCLRLH PTSQPLFAFK
VQDIHPTVPN PYNLLCALPP QRSWYTVLDL KDAFFCLRLH PTSQPLFAFE
VQDIHPTVPN PYNLLCALPP QRSWYTVLDL KDAFFCLRLH PTSQPLFAFE
VQDIHPTVPN PYNLLCALPP QRSWYKVLDL KDAFFCLRLH PTSQPLFAFE
VQDIHPTVPN PYNLLSALPP ERNWYTVLDL KDAFFCLRLH PTSQPLFAFE
VQDIHPTVPN PYNLLSALLP ERNWYTVLDL KDAFFCLRLH PTSQPLFAFE
VQDIHPTVPN PYNLLSALPP ERNWYTVLDL KDAFFCLRLH PTSQPLFAFE
VQDIHPTVPN PYNLLSALPP ERNWYTVLDL KDAFFCLRLH PTSQPLFAFE
VQDIHPTVPN PYNLLSALPP ERNWYTVLDL KDAFFCLRLH PTSQPLFAFE
VQDIYPTVPN PYNLLSALPP ERNWYTVLDL KDAFFCLRLH PSSQPLFAFE
V-DIYPTVPN PYNLLSALPP E-NWYTVLDL KDAFFCLRLH PTSQPLFAFE
VQDIHPMVPN PYNLLSALPP KRNWYTVLDL KNAFFCLRLH PTSQPLFAFE
VQDIHPTVPN PYNLLCALPP QRSWYTVLDL KDAFFCLRLH PTSQPLFAFE
VQDIHPTVPN PYNLLCALPP QRSWYTVLDL KDAFFCLRLH PTSQPLFAFD
VQDIHPTVPN PYNLLCALPP QRSWYTVLDL KDAFFCLRLH PTSQPLFAFK
VQDIHPTVPN PYNLLSALPP ERNWYTVLDL KDAFFCLRLH PTSQPLFAFE
VQDIHPTVPN SYNLLSALPP ERNWYTVLDL KDAFFCLRLH PTSQPLFAFE
VQDIHPTVPN SYNLLSALPP ERNWYTVLDL KDAFFCLRLH PTSQPLFAFE
VQDIHPTVPN PYNLLCALPP QWSWYTVLDL KDVFFCLRLH PTSQPLFAFE

VQDIHPTVPN PYNLLSALPP E--WYTVLDL KD-FFCLRLH PTSQPLFAFE
VQDIHPTVPN PCNLLSALPP ERIWYTVLDL KDAFFCLRLH LDSQPLFAFE
VQDIHPTVPN PYNLLSTLPP GRTWYTVLDL KDAFFCLRLH PNSQPLFAFE
VQDIHPTVPN PYNLLSTLPP GQTWYTVLDL KDAFFCLRLH PNSQPLFAFE
VQDIHPTVPN PYNLLSTLPP GRTWYTVLDL KDAFFCLRLH PNSQPLFAFE
VQDIHPTVPN PYNLLSTLPP GRT-YTVLDL KDAFFCLRLH PNSQPLFAFE
VLDIHPTVPN PYNLLSSLPP ERTWYTVLDL KDAFFCLRLH PKSQLLFAFE
VLDIHPTVPN PYNLLSSLPP ERTWYTVLDL KDAFFCLHLH PKSQLLFAFE
VLDIHPTVPN PYNLLSSLPP ERTWYTVLDL KDAFFCLRLH PKSQLLFAFE
VLDIHPTVPN PYNLLSSLPP ERTWYTVLDL KDAFFCLRLH PKSQLLFAFE
VLDIHPTVSN PYNLLSSLPP ERTWYTVLDL KDAFFCLRLH TKSQLLFAFE
VLDIHPTVPN PYNLLSSLPP ERTWYTVLDL KDAFFYLRLH PKSQLLFAFE
VLDIHPTVPN PYNLLSSLPP ERTWYTVLDL KDAFFCLRLH PKSQLLFAFE
VQDIHPTVPN PYNLLSSLPP ERTWYTVLDL KDAFFCLRLH PNSQPLFAFE
VQDIHPTVPN PYNLLSSLPP SHTWYSVLDL KDAFFCLKPH PNSQPLFAFE
VQDIHPTVPN PYNLLSSLPP SYTWYSVLDL KDAFFCLRLH PNSQPLFAFE
VQDIHPTVPN PYNLLSSLPP ERKWYTVLDL KDAFFCLKLH PFSQPIFAFE
VQDIHPTVPN PYNLLSSLPP ERKWYTVLDL KDAFFCLKLH PSSQPIFAFE
VQDIHPTVPN PYNLLSSLPP ERKWYTVLDL KDAFFCLKLH PSSQPIFAFE
VQDIHPTVPN PYNLLSSLPP ERKWYTVLDL KDAFFCLKLH PSSQPIFAFE
VQDIHPTVPN PYNLLSSLPP ERKWYTVLDL KDAFFCLKLH PSSQPIFAFE
VQDIHPMVPN PYNLLISLPP EK-WYTVLDL KDAFFCLKLH PSSQPIFAFE
VQDIHPTVPN PYNLLSALPP KRS-YTVLDL KDAFFCLRLH LDSQSLFAFE
VEDVHPIVPN PYTLLSTLPP ERIWYTVLDL KDAFFSLRLH PNSQHLFAFE
VRDIHPTVPN PYNLLSSLHP DRQWYTVLDP KDAFFCLKLH PMSQVIFAFE
MQDIHPTVPN P-NLLSSLPP EQQWYTVLDL KDAFFRLKLH SRSQPIFAFE
VQDIHPTIPN PYTLLSSIPS ERTWYTVLDL RDAFFCLRLH PSSQPIFTFE
VLDIHPTVPN PYNLLSSLPP SHVWYTVLDF KDAFFCLRLH PTSQPMFAFE
VQDIHPTVPN PYNLLSSLPP SHVWYTVLDL KDAFFCLRLH PTSQPMFAFE
VQDIHPTVPN PYNLLSSLPP SHVWYTVLDL KDAFFCLRLH PTSQPMFAFE
VQDIHPTVPN PYNLLSTLPP ERTWYTVLDL KDAFFCLRLH PNSQPLFAFE
IQDIHPTLPN PYNLLSSLPK E---YTVLDL KDAFFCLRLH PSSQPIFTFE
VEDIHPTVPN PYNLLSGLPP SHQWYTVLDL KDAFFCLRLH PTSQPLFAFE
VEDIHPTVPN PYNLLSGLPP SHQWYTVLDL KDAFFCLRLH PTSQPLFTFE
VEDIHPTVPN PYNLLSGLPP SHQWYTVLDL KDAFFCLRLH PTSQPLFAFE
VEDIHPTVPN PYNLLSGLPP SHQWYTVLDL KDAFFCLRLH PTSQPLFAFE
VEDIHPTVPN PYNLLSGLPP SHQWYTVLDL KDAFFCLRLH PTSQPLFAFE
VEDIHPTVPN PYNLLSGLPP SHQWYTVLDL KDAFFCLRLH PTSQPLFAFE
VEDIHPTVPN PYNLLSGLPP SHQWYTVLDL KDAFFCLRLH PTSQPLFAFE
VEDIHPTVPN PYNLLSGLPP SHQWYTVLDL KDAFFCLRLH PTSQSLFAFE
VEDIHPTVPN PYNLLSGLPP SHQWYTVLDL KDAFFCLRLH PTSQSLFAFE
VEDIHPTVPN PYNLLSGLPP SHQWYTVLDL KDAFFCLRLH PTSQPLFAFE
VEDIHPTVPN PYNLFSTLPP THTWYTVLDL KDAFFCLWLS PKSQPLFAFE
VEDIHPTVPN PYNLFSTLPP THTWYTVLDL KDAFFCLWLS PKSQPLFAFE

```
VEDIHPTVPN PYNLLSTLPP SHPWYTVLDL KDAFFCLRLH SESQLLFAFE
TMDIHPTVPN PYNLLSTLSP DRTWYTVLDL KDAFFCLPLA PQSQELFAFE
TVDIHPTVPN PYNLLSTLKP DYSWYTVLDL KDAFFCLPLA PQSQELFAFE
VADLHPTVPN PYNLLSTLPP QHTWYTVLDL KDAFFCLRLS PLSQPYFAFE
MQDMHPTVPN PHNLLSLLPP EQ-WYTVFDL KDAFFCPKLN SRS-AIFAFE
VQDIHPT-PN -YNLLSALLP ERNWYTV-DL KDAFFCLRLP PTQPLLFE--
VQDIHPTVPN PYNLLSLLPT ERKWYIVLDL KDAFFCLKLH SSSQLIFAFD
V-DLYPTVPK PYNLLSSLPP DRKWYSVLDL KDAFF-LRLH HSSQSIFAFE
VEDIHPTVPN PYTLLSHLPP SHVWYTTLDL KDAFFSIALA PSSQHIFAFE
VQDIXPTXPN PXN------- EQKWYTVFDL KDVSFFLKLH PSSQPIFAFE
VETIHPTVPN PYTLLSLLPP DRIWYSVLDL KDAFFCIPLA PESQLIFAFE
VQDFYPAFPN LYNLLSSLPP KQK-YTVLDL KNAFFCLKLH PSSQPIFAFE
VQDFYPAVPN LYNLLSSLPP KQK-YTVLDL KKCFLVTSLQ PAHIRLV---
VQDIHPTVPN TYNLLTSLPN DRQWYMVMDL NYSFFFLKLH PMSQVIFAFK
V-NIHPAVPN SYNLLSSLHP DRHCYIVLHL KDTFFCLKLH PMSQVIFAFK
L-DIYPTVPN SYNLLSSLYP DRHCYIVLHL KDTFFFLKLH PMSQVIFAFK

WRDPGTGRTG QLTWTRLPQG FKNSPTIFDE ALHRDLANFR IQHPQVTLLQ
WRDPGTGRTG QLTWTRLPQG FKNSPTIFDE ALHRDLANFR IQHPQVTLLQ
WRDPGTGRTG QLTWTRLPQG FKNSPTIFDE ALHRDLANFR IQHPQVTLLQ
WRDPDTGRTG QLTWTRLPQG FKNSPTIFDE ALHRDLANFR IQHPQVTLLQ
WRDPGTGRTG QLTWTRLPQG FKNSPTIFDE ALHRDLANFR IQHPQVTLLQ
WRDPGAGRTG QLTWTRLPQG FKNSPTIFDE ALHRDLANFR IQHPQVTLLQ
WRDPGAGRTG QLTWTRLPQG FKNSPTIFDE ALHRDLANFR IQHPQVTLLQ
WRDPGAGRTG QLTWTRLPQG FKNSPTIFDE ALHRDLANFR IQHPQVTLLQ
WRDPGAGRTG QLTWTRLPQG FKNSPTIFDE ALHRDLANFR IQHPQVTLLQ
WRDPGAGRTG QLTWTRLPQG FKNSPTIFDE ALHRDLANFR IQHPQVTLLQ
WRDPGAGRTG QLTWTRLPQG FKNSPTIFDE ALHRDLANFR IQHPQVTLLQ
WRDPGAGRTG QLTWTRLPQG FKNSPTIFDE ALHRDLANFR IQHPQVTLLQ
WRDPGAGRTG QLTWTRLPQG FKNSPTIFDE ALHRDLANFR IQHPQVTLLQ
WRDPGAGRTG QLTWTRLPQG FKNSPTIFDE ALHRDLANFR IQHPQVTLLQ
WRDPGTGRTG QLTWTRLPQG FKNSPTIFDE ALHRDLANFR IQHPQVTLLQ
WRDPGAGRTG QLTWTRLPQG FKNSPTIFDE ALHRDLANFR IQHPQVTLLQ
WRDPGAGRTG QLTWTRLPQG FKNSPTIFDE ALHRDLANFR IQHPQVTLLQ
WRDPGAGRTG QLTWTRLPQG FKNSPTIFDE ALHRDLANFR IQHPQVTLLQ
WKDPGTGRTG QLTWTRLPQR FKNAPTIFDE ALHRDLANFR IQHPQVTLLQ
WRDPGAGRTG QLTWTRLPQG FKNSPTIFDE ALHRDLANFR IQHPQVTLLQ
WKDPGTGRTG QLTWTRLPQR FKNAPTIFDE ALHRDLANFR IQHPQVTLLQ
WRDPGTGRTG QLTWTRLPQG FKNSPTIFDE ALHRDLANFR IQHPQVTLLQ
WRDPGTGRTG QLTWTRLPQG FKNSPTIFDE ALHRDLANFR IQHPQVTLLQ
WRDPGTGRTG QLTWTRLPQG FKNSPTIFDE ALHRDLANFR IQHPQVTLLQ
WRDPGTGRTG QLTWTRLPQG FKNSPTIFDE ALHRDLANFR IQHPQVTLLQ
WRDPGTGRTG QLTWTRLPQG FKNSPTIFDE ALHRDLANFR IQHPQVTLLQ
WRDPGTGRTG QLTWTRLPQG FKNSPTIFDE ALHRDLANFR IQYPQVTLLQ
```

WRDPGTGRTG QLTWTRLPQG FKNSPTIFDE ALHRDLANFR IQHPQVTLLQ
WRDPGTGRTG QLTWTRLPQG FKNSPTIFDE ALHRDLANFR IQHPQVTLLQ
WRDPGTGRTG QLTWTRLPQG FKNSPTIFDE ALHRDLANFR IQHPQVTLLQ
WRDPGTGRTG QLTWTRLPQG FKNSPTIFDE ALHRDLANFR IQHPQVTLLQ
WRDPGTGRTG QLTWTRLPQG FKNSPTIFDE ALHRDLANFR IQHPQVTLLQ
WRDPGTGRTG QLTWTRLPQG FKNSPTIFDE ALHRDLANFR IQHPQVTLLQ
WRDPGTGRTG QLTWTRLPQG FKNSPTIFDE ALHRDLANFR IQHPQVTLLQ
WRDPGTGRTG QLTWTRLPQG FKNSPTIFDE ALHRDLANFR IQHPQVTLLQ
WRDPGTGRTG QLTWTRLPQG FKNSPTIFDE ALHRDLANFR IQHPQVTLLQ
WRDPGAGRTG QLTWTRLPQG FKNSPTIFDE ALHRDLANFR IQHPQVTLLQ
WRDPGTGRTG QLTWTRLPQG FKNSPTIFDE ALHRDLANFR IQHPQVTLLQ
WRDPGAGRTG QLTWTRLPQG FKNSPTIFDE ALHRDLANFR IQHPQVTLLQ
WRDPGAGRTR QLTWTRLPQG FKNSPTIFDE ALHRDLANFR IQHPQVTLLQ
WRDPGAGRTG QLTWTRLPQG FKNSPTIFDE ALHRDLANFR IQHAQVTLLQ
WRDPGTGRTR QLTWTRLPQG FKNSPTIFDE ALHRDLANFR IQHPQVTLLQ
WRDPGTGRTG QLTWTRLPQG FKNSPTIFDE ALHRDLANFR IQHPQVTLLQ
WRDPGTGRTR QLTWTRLPQG FKNSPTIFDE ALHRDLANFR IQHPQVTLLQ
WRDPGTGRTG QLTWTRLPQG FKNSPTIFDE ALHRDLANFR IQHPQVTLLQ
WRDPGTGRTG QLTWTRLPQG FKNSPTIFDE ALHRDLANFR IQHPQVTLLQ
WRDPGTGRTG QLTWTRLPQG FKNSPTIFDE ALHRDLANFR IQHPQVTLLQ
WRDPGTGRTG QLTWTRLPQG FKNSPTIFDE ALHRDLANFR IQHPQVTLLQ
WRDPGTGRTG QLTWTRLPQG FKNSPTIFDE ALHRDLANFR IQHPQVTLLQ
WRDPGAGRTG QLTWTRLPQG FKNSPTIFDE ALHRDLANFR IQHPQVTLLQ
WKDPGTGRTG QLTWTRLPQR FKNAPTIFDE ALHRDLANFR IQHPQVTLLQ
WRDPGAGRTG QLTWTRLPQG FKNSPTIFDE ALHRDLANFR IQHPQVTLLQ
WRDPGTGRTG QLTWTRLPQG FKNSPTIFDE ALHRDLANFR IQHPQVTLLQ
WRDPGTGRTG QLTWTRLPQG FKNSPTIFDE ALHRDLANFR IQHPQVTLLQ
WRDPGTGRTG QLTWTRLPQG FKNSPTIFDE ALHRDLANFR IQHPQVTLLQ
WRDPGTGRTG QLTWTRLPQG FKNSPTIFDE ALHRDLANFR IQHPQVTLLQ
WRDPGTGRTG QLTWT-LPQG FKNSPTIFDK ALHKDLANFR IQHPQVTLLQ
WRDPNTERTG QLTWTRLPQG FKNSPTIFDE ALHRDLANFR VQHPQVTLLQ
WRDSESGQAG QLTWTRLPQG FKNSPTLFDE ALHRDLALFR ANNPQVTLLQ
WRDSESGQAG QLTWTRLPQG FKNSPTLFDE ALHRDLALFR ANNPQVTLLQ
WRDSESGQAG QLTWTRLPQG FKNSPTLFDE ALHRDLALFR ANNPQVTLLR
WRDSESGQAG QLTWTRLPQG FKNSPTLFDE ALHRDLALFR ANNPQVTLLQ
WRDPEGGQTG QLTWTRLPQG FKNSPTLFDE ALHRDLAPFR ARNPQLTLLQ
WRDPEGGQTG QLTWTRLPQG FKNSPTLFDE ALHRDLAPFR ARNPQLTLLQ
WRDPEGGQTG QLTWTRLPQG FKNSPTLFDE ALHRDLVPFR AQNPQLTLLQ
WRDPEGGQTG QLTWTRLPQG FKNSPTPFDE ALHRDLAPFR ARNPQLTLLQ
WRDPEGRQTG QLTWTRLPQG FKNSPTLFDE ALHRDLAPFH AQNPQLTLLQ
WRDPEGGHTG QLTWTRLPQV FKNSPTLFDE ALHRDLAPFH ARNPQLTLLQ
WRDPEGGQTG QLTWTRLPQG FKNSPTLFDE ALHRDLAPFR AQNPQLTLLQ
WRDPEGGHTG QLTWTRLPQG FKNSPTLFDE ALHRDLAPFR AQNPQISLLQ
WRDPEKGNTG QLTWTRLPQG FKNSPTLFDE ALHRDLASFR ALNPQVVMLQ

```
WKDPEKGNTG QLTWTRLPQG FKNSPTLFDE ALHRDLAPFR ALNPQVVLLQ
WRDPDTGQTG QLTWTRLPQG FKNSPTLFDE ALHRDLASFR AENSQVTLLQ
WRDPDTGQMG QLNWTRLPQE FKNSPTLFDE ALHRDLASFR AENSQVTLLQ
WRDPDTGQMG QLTWTRLPQE FKNSPTLFDE ALHRDLASFL AENSQVTLLQ
WRDPDTGQMG QLTWTRLPE- FKNSPTLFDE ALHRDLASFR AENSQVTLLQ
WRDPDTGQMG QLTWTRLPE- FKNSPTLFDE ALHRDLASFR AENSQVTLLQ
WRDPDTGQTG QLTWTRLPRG FKNSPTLFDE ALHGDLASFR AKKPQLILLQ
WRDPDTGRTG -VNWTHLP-G FKNSPTIFDK ALHRDLANFR VQHPHLTLLQ
WKGPETGHSG QLTWT-LPQG FKNSPTLFDE ALHRDLTIFR TNNPHLTLLQ
WRDPDSGQAG QLTWTRLPQG FKNSPTLFYE ALHWDLVLFR SENPQISLLQ
WKDPDTGQTG QLTWTRLPQG FKNSPTLFDE ALHWDLATFR TAHPQGTLLQ
-RDPNTGQVG QLTWTRLPQG FKNSPTLFDE ALHRDLAAFR PTTPQVTLLQ
W-DPQDGITG QLTWTRLPQG FKNSPTIFDE ALHQDLMPFR VSHPQVTLLK
WKDPQDGTTG QLTWTRLPQG FKNSPTIFDE ALHQDLTPFH ASHPQVTLLQ
WKDPQDGTTG QLMWTRLPQG FKNSPTIFDE ALHQDLTPFR ASHPQVTLLQ
WRDPESEKTG QLTWTRLPQG FKN-PTLFDE ALHRDLSSFQ ANDPQVTLVQ
W-DPESGVKG KLTWARLPQG FKNSPTIFDE ALHRDLASFR ATNPQVTLLQ
WRDPEMGISG QLTWTRLPQG FKNSPTLFDE ALHRDLADFR IQHPDLILLQ
WRDPEMRISG QLTWTRLPQG FKNSPTLFDE ALHRDLADFR IQHPDLILLQ
WRDPEMGISG QLTWTRLPQG FKNSPTLFDE ALHRDLAGFR IQHPDLILLQ
WRDPEMGISG QLTWTRLPQG FKNSPTLFDE ALHRDLADFR IQHPDLILLQ
WRDPEMGISG QLTWTRLPQG FKNSPTLFDE ALHRDLAGFR IQHPDLILLQ
WRDPEMGISG QLTWTRLPQG FKNSPTLFDE ALHRDLAGFR IQHPDLILLQ
WRDPEMGISG QLTWTRLPQG FKNSPTLFDE ALHRDLADFR IQHPDLILLQ
WRDPEMGISG QLTWTRLPQG FKNSPTLFDE ALHRDLADFR IQHPDLILLQ
WRDPEMGISG QLTWTRLPQG FKNSPTLFDE ALHRDLADFR IQHPDLILLQ
WRDPEMGISG QLTWTRLPQG FKNSPTLFDE ALHRDLADFR IQHPDLILLQ
WKDPEIGLSG QLTWTRLPQG FKNSPTLFDE ALHRDLADFR VQHPTLILLQ
WKDPEIGLSG QLTWTRLPQG FKNSPTLFDE ALHRDLADFR VQHPTLILLQ
WRDPEIGLSG QLTWTRLPQG FKNSPTLFDE ALHSDLADFR VRYPALVLLQ
WRDPERGISG QLTWTRLPQG FKNSPTLFDE ALHRDLTDFR TQHPEVTLLQ
WKDPERGISG QLTWTRLPQG FKNSPTLFDE ALHRDLTDFR TQHPEVTLLQ
WKDPASGISG QLTWTCLPQE FKNSPTIFDE ALHQDLALYR ESNPQVTLLQ
WKDPDTGQTG QLTWTRLPQG FKYSPTLFNE ALHRDLATFR TAQPQ-TLLQ
-RDPGT-EEG QLTWTRLPQG -KNSPTIFDE ALHRDANFIQ HPTP-----Q
WRDPDSGMLG QLTWTRLLQG FKNSPTLFHE ALHRDLATFR TENPQVTLLQ
WRDPDTGLVG LLTWTRFPQG FKNSPTLFDE ALHWDLATFR AENPQVTLFQ
WNNGNTGTPG QLTWTRLPQG FKNSPTLFNE ALNQDLDSFR QSHNSVTLLQ
WRDLNTGQTG QLT-TQLPQR FKNTPILFYE ALH-DLVSF- AKNSQITLFQ
WADAEEGESG QLTWTRLPQG FKNSPTLFDV ALNRDLQGFR LDHPSVSLLQ
WTDLDMGQMG QLTLKRLPQG FKNSPTLFHK SLHRDLASF- AENCQITLLQ
--ERSHGTNG TIDLEAVTPG VQKLPHPFFI SLYIEIPPSE ARLCFSVLM-
WRNPDSGQAI QLTWTLLPTR FKNSTTLFYE ALLWDMAPFR SENLQISLLQ
WRDPGSGHIY LKIYLKIY-- LKNSTTLFDE AFHPDMALFR SEIPQVFLLH
```

WRDPGAGQTN FETFTK---- FKNSPTLFYE AFHSDLALFR AEIPQVFLLH

YVDDLLLAGA TKQDCLEGTK ALLLELSDLG YRASAKKAQI CRREVTYLGY
YVDDLLLAGA TKQDCLEGTK ALLLELSDLG YRASAKKAQI CRREVTYLGY
YVDDLLLAGA TKQDCLEGTK ALLLELSDLG YRASAKKAQI CRREVTYLGY
YVDDLLLAGA TKQDCLEGTK ALLLELSDLG YRASAKKAQI CRREVTYLGY
YVDDLLLAGA TKQDCLEGTK ALLLELSDLG YRASAKKAQI CRREVTYLGY
YVDDLLLAGA TKQDCLEGTK ALLLELSDLG YRASAKKAQI CRREVTYLGY
YVDDLLLAGA TKQDCLEGTK ALLLELSDLG YRASAKKAQI CRREVTYLGY
YVDDLLLAGA TKQDCLEGTK ALLLELSDLG YRASAKKAQI CRREVTYLGY
YVDDLLLAGA TKQDCLEGTK ALLLELSDLG YRASAKKAQI CRREVTYLGY
YVDDLLLAGA TKQDCLEGTK ALLLELSDLG YRASAKKAQI CRREVTYLGY
YVDDLLLAGA TKQDCLEGTK ALLLELSDLG YRASAKKAQI CRREVTYLGY
YVDDLLLAGA TKQDCLEGTK ALLLELSDLG YRASAKKAQI CRREVTYLGY
YVDDLLLAGA TKQDCLEGTK ALLLELSDLG YRASAKKAQI CRREVTYLGY
YVDDLLLAGA TKQDCLEGTK ALLLELSDLG YRASAKKAQI CRREVTYLGY
YVDDLLLAGA TKQDCLEGTK ALLLELSDLG YRASAKKAQI CRREVTYLGY
YVDDLLLAGA TKQDCLEGTK ALLLELSDLG YRASAKKAQI CRREVTYLGY
YVDDLLLAGA TKQDCLEGTK ALLLELSDLG YRASAKKAQI CRREVTYLGY
YVDDLLLAGA TKQDCLEGTK ALLLELSDLG YRASAKKAQI CRREVTYLGY
YVDDLLLAGA TKQDCLEGTK TLLLELSDIG YRASTKKAQI CRREVTYLGY
YVDDLLLAGA TKQDCLEGTK ALLLELSDLG YRASAKKAQI CRREVTYLGY
YVDDLLLAGA TKQDCLEGTK ALLLELSDLG YRASAKKAQI CRREVTYLGY
YVDDLLLAGA TKQDCLEGTK ALLLELSDLG YRASAKKAQI CRREVTYLGY
YVDDLLLAGA TKQDCLEGTK ALLLELSDLG YRASAKKAQI CRREVTYLGY
YVDDLLLAGA TKQDCLEGTK ALLLELSDLG YRASAKKAQI CRREVTYLGY
YVDDLLLAGA TKQDCLEGTK ALLLELSDLG YRASAKKAQI CRREVTYLGY
YVDDLLLAGA TKQDCLEGTK ALLLELSDLG YRASAKKAQI CRREVTYLGY
YVDDLLLAGA TKQDCLEGTK ALLLELSDLG YRASAKKAQI CRREVTYLGY
YVDDLLLAGA TKQDCLEGTK ALLLELSDLG YRASAKKAQI CRREVTYLGY
YVDDLLLAGA TKQDCLEGTK ALLLELSDLG YRASAKKAQI CRREVTYLGY
YVDDLLLAGA TKQDCLEGTK ALLLKLSDLG YRASAKKAQI CRREVTYLGY
YVDDLLLAGA TKQDCLEGTK ALLLELSDLG YRASAKKAQI CRREVTYLGY
YVDDLLLAGA TKQDCLEGTK ALLLELSDLG YRASAKKAQI CRREVTYLGY
YVDDLLLAGA TKQDCLEGTK ALLLELSDLG YRASAKKAQI CRREVTYLGY
YVDDLLLAGA TKQDCLEGTK ALLLELSDLG YRASAKKAQI CRREVTYLGY
YVDDLLLAGA TKQDCLEGTK ALLLELSDLG YRASAKKAQI CRREVTYLGY
YVDDLLLAGA TKQDCLEGTK ALLLELSDLG YRASAKKAQI CRREVTYLGY
YVDDLLLAGA TKQDCLEGTK ALLLELSDLG YRASAKKAQI CRREVTYLGY
YVDDLLLAGA TKQDCLEGTK ALLLELSDLG YRASAKKAQI CRREVTYLGY
YVDDLLLAGA TKQDCLEGTK ALLLELSDLG YRASAKKAQI CRREVTYLGY
YVDDLLLVGA TKQDCLEGTK -LLLELSDLG YRASAKKAQI CRREVTYLGY
YVDDLLLAGA TKQDCLEGTK ALLLELSDLG YRASAKKAQI CRKEVTYLGY

YVDDLLLAGA TKQDCLEGTK ALLLELSDLG YRASAKKAQI CRREVTYLGY
YVDDLLLAGA TKQDCLEGTK ALLLELSDLG YRASAKKAQI CRKEVTYLGY
YVDDLLLAGA TKQDCLEGTK ALLLELSDLG YRASAKKAQI CRREVTYLGY
YVDDLLLAGA TKQDCLEGTK ALLLELSDLG YRASAKKAQI CRREVTYLGY
YVDDLLLAGA TKQDCLEGTK ALLLKLSDLG YRASAKKAQI CRREVTYLGY
YVDDLLLAGA TKQDCLEGTK ALLLKLSDLG YRASAKKAQI CRREVTYLGY
YVDDLLLAGA TKQDCLEGTK ALLLELSDLG YRASAKKAQI CRREVTYLGY
YVDDLLLAGD TKQDCLKGTK ALLLELSDLG YRASAKKAQI CRREVTYLGY
YVDDLLLAGA TKQDCLEGTK TLLLELSDIG YRASTKKAQI CRREVTYLGY
YVDDLLLAGA TKQDCLEGTK ALLLELSDLG YRASAKKAQI CRREVTYLGY
YVDDLLLAGA TKQDCLEGMK ALLLELSDLG YRASAKKAQI CRREVTYLGY
YVDDLLLAGA TKQDCLEGTK ALLLELSYLG YRASAKKAQI CRREVTYLG-
YVDDLLLAGA TKQDCLEGTK ALLLELSYLG YRASAKKAQI CRREVTYLG-
YVDDLLLAGA TKQDCLECTK ALLLELSDLG YRASAKKAQI CRREVTYLGY
FEDDLLLAGA TKQDCLEGTK ALLLELSDLG YRASAKKAQI CRREVTYLGY
YVDDLLLAGA TQQDCLKGTK ALLLELTDLG YRASAKKAQI CKREVTYLGY
YVDDLLLAAE TREDCEIGTQ NLLGELGKLG YRASAKKAQL CQIEVTYLGY
YVDDLLLAAE TREDCEIGTQ NLLGELGKLG YRASAKKAQL CQIEVTYLGY
YVDDLLLAAE TREDCEIGTQ NLLGELGKLG YRASAKKAQL CQIEVTYLGY
YVDDLLLAAE TREDCEIGTQ NLLGELGKLG YRASAKKAQL CQIEVTYLGY
YVDDLLVAAA SKELCHQGTE RLLAELSDLG YRVSAKKAQI CQTEVTYLGY
YVDDLLVAAA SKELCHQGTE RLLAELSDLG YRVSAKKARI CQTEVTYLGY
YVDDLLVAAA SKELCHQGTE RLLAELSDLG YRVSAKKAQI CQTEVTYLGY
YVDDLLVAAA SKELCHQGTE RLLAELSDLG YRVSAKKAQI CQTEVTYLGY
YVDDLLVAAA SKELCHQGTE RLLAELSDLG YRVSAKKVQI CQTEVTYLGY
YVDDLLVAAA SKELCHQGTE RLLTELSDLG YRVSAKKAQI CQTEVTYLGY
YVDDLLIAAA SKELCQQGTE RLLTELGNLG YRVSAKKAQI CQTEVIYLGY
YVDDLLLAAS TQELCREGTK RLLNELGELG YRVSAKKAQL CRTEVTYLGY
YVDDLLVAAP TYRDCKEGTR RLLQELSKLG YRVSAKKAQL CREEVTYLGY
YVDDLLVAAP TYEDCKKGTQ KLLQELSKLG YRVSAKKAQL CQREVTYLGY
YVDDLLLAAT TERECWQGTK KLLVELGELG YRASAKKAQL CQMEVVYLGY
YVDDLLLAAT TERECWQGTK RLLVELGELG YRASAKKAQL CQMEVVYLGY
YVDDLLLAAT TERECWQGTK RLLVELGELG YRASAKKAQL CQMEVVYLGY
YVDDLLLAAT IERECWQGTK RLLVELGELG YRASAKKAQL CQMEVVYLGY
YVDDLLLAAT IERECWQGTK RLLVELGELG YRASAKKAQL CQMEVVYLGY
YADDLLLGAT SEQECRQGTE KLLTELGELG YRASAKKAQL CQMEVVYLGY
YVDDLLLAGA TQQNCLKGMK AQLLELTDLG Y-ASAKKAQI CKREVTYLGY
YVDDLLLGAE TQGECQKGTE QLLAELGCLG YRASAKKAQL CRTEVTYLGY
YADDLLLAAT TKQECRLGTE KLLAELSKLG YKASAKKA-I CQTEVTYLGY
YVDDLLVAAS TKEECLRGTK DLLTELGELG YRASTKKAQL CQTEVIYLGY
YVDDLLLSVA TEDECWRGTE DLLQELGELG YRTLAKKAQL CWKEVTYLGY
YVDDLLIAGA TETECREATM ALLTELSRLG Y-ASAKKALL CQREVTYLGY
YVDDLLIARA SEAECQEATM ALLTELSPLG YRASAKKAQL CRQEVTYLGY
YVDDLLIAGA SEAECREATM ALLTELSRLG YRASAKKAQL CRQEVTYLGY

```
YVDDLLIAAG TREECELGTQ NILVELGELG Y-VSAKKGAV IPNRGDLL-W
YVDDLLLAAT TKEECQQGAK NLLAELGELG YWTSAKKAQL CWTEVTYLGY
YVDDLLLAAT SELDCQQGTR ALLQTLGNLG YRASAKKAQI CQKQVKYLGY
YVDDLLLAAT SELDCQQGTR ALLQTLGNLG YRASAKKAQI CQKQVKYLGY
YVDDLLLAAT SELDCQQGTR ALLQTLGDLG YRASAKKAQI CQKQVKYLGY
YVDDLLLAAT SELDCQQGTR ALLQTLENLG YRASAKKAQI CQKQVKYLGY
YVDDLLLAAT SELDCQQGTR ALLQTLGDLG YRASAKKAQI CQKQVKYLGY
YVDDLLLAAT SELDC-QGTR ALLQTLGDLG YQASAKKAQI CQKQVKYLGY
YVDDLLLAAT SELDCQQGTR ALLQTLGDLG YRASAKKAQI CQKQVKYLGY
YVDDLLLAAT SELDCQQGTR ALLQTLGDLG YRASAKKAQI CQKQVKYLGY
YVDDLLLAAT SELDCQQGTR ALLQTLGDLG YRASAKKAQI CQKQVKYLGY
YVDDLLLAAT SELDCQQGTR ALLQTLGNLG YRASAKKAQI CQKQVKYLGY
YVDDLLLVAT SETACHQGTE SLLQTLGRLG YRASAKKGQI CQTQVTYLGY
YVDDLLLVAT SETACHQGTE SLLQTLGRLG YRASAKKGQI CQTQVTYLGY
YVDDLLLAAA TRTECLEGTK ALLETLGNKG YRASAKKAQI CLQEVTYLGY
YVDDLLLAAP TEEACTRGTK HLLRELGDKG YRASAKKAQI CQTKVTYLGY
YVDDLLLAAP TKKACTQGTR HLLQELGEKG YRASAKKAQI CQTKVTYLGY
YVDDILLAAE TQEDCIKGTE KLLTELRTLG YRASAKKAQI CQQQVSYLGY
-VDDLLVAAS TKEECFHGTK ELLTELGELG YRASV-KAQL CQTEVTYLGY
YVE--LLGAT K--DC-EGTK ALLLELSP-S YRA-AKKAQI CRREVTYLGY
YVDDLLLATT TQAECQQGTE KPLTELGELG YRASAKNVRL CQTEVTYLGY
YVDDLLLAAT TELECRKGTE KLLTELGELG YQASAKKAQL CQTEVTYLGY
YVDDLLLAAP SEAECRQATG DLLQELGQLG YRASAKKAQI CRQTVTYLGY
YVDDLLLAAK SERECLQGSK RLLAELGELG YRASTKKAQL CQMEVVYLGY
YVDDLLIAAD TQAACLSATR DLLMTLAELG YRVSGKKAQL CQEEVTYLGF
CADDLHLAAA TEREFWQGTK RLLGELGELG W-ASVKKAQL CHIEVVYLGY
---------- --------TC SLLQENSGTG PRDCPKKAQL CHIEVVYLGY
YVDDLLLAAT TEQEARLGTE KLLAELSELG YKASAKKAQL CQAEVTYLGY
YVYDLLLAAT TEKCCRQGTE KLLAELSELG YKASAKKAQN YQTEVTYLGY
YVYDLLLAAT TEKWCRLGTE KLLAELSELG YKASANKAQN CQTEVTYLGY

SLRDGQRWLT EARKKTVVQI PAPTTAKQMR EFLGTAGFCR LWIPGFATLA
SLRDGQRWLT EARKKTVVQI PAPTTAKQMR EFLGTAGFCR LWIPGFATLA
SLRDGQRWLT EARKKTVVQI PAPTTAKQMR EFLGTAGFCR LWIPGFATLA
SLRDGQRWLT EARKKTVVQI PAPTTAKQMR EFLGTAGFCR LWIPGFATLA
SLRDGQRWLT EARKKTVVQI PAPTTAKQMR EFLGTAGFCR LWIPGFATLA
SLRGGQRWLT EARKRTVVQI PAPTTAKQVR EFLGTAGFCR LWIPGFATLA
SLRGGQRWLT EARKRTVVQI PAPTTAKQVR EFLGTAGFCR LWIPGFATLA
SLRGGQRWLT EARKRTVVQI PAPTTAKQVR EFLGTAGFCR LWIPGFATLA
SLRGGQRWLT EARKRTVVQI PAPTTAKQVR EFLGTAGFCR LWIPGFATLA
SLRGGQRWLT EARKRTVVQI PAPTTAKQVR EFLGTAGFCR LWIPGFATLA
SLRGGQRWLT EARKRTVVQI PAPTTAKQVR EFLGTAGFCR LWIPGFATLA
SLRGGQRWLT EARKRTVVQI PAPTTAKQVR EFLGTAGFCR LWIPGFATLA
SLRGGQRWLT EARKRTVVQI PAPTTAKQVR EFLGTAGFCR LWIPGFATLA
```

```
SLRGGQRWLT EARKRTVVQI PAPTTAKQVR EFLGTAGFCR LWIPGFATLA
SLRGGQRWLT EARKKTVVQI PAPTTAKQVR EFLGTAGFCR LWIPGFATLA
SLRGGQRWLT EARKRTVVQI PAPTTAKQVR EFLGTAGFCR LWIPGFATLA
SLRGGQRWLT EARKRTVVQI PAPTTAKQVR EFLGTAGFCR LWIPGFATLA
SLRGGQRWLT EARKRTVVQI PAPTTAKQVR EFLGTAGFCR LWIPGFATLA
SLRDGQRWLT EARKKTVVQI PAPTTAKQMR EFLGTAGFCR LWIPGFATLA
SLRGGQRWLT EARKRTVVQI PAPTTAKQVR EFLGTAGFCR LWIPGFATLA
SLRGGQRWLT EARKRTVVQI PAPTTAKQVR EFLGTAGFCR LWIPGFATLA
SLRDGQRWLT EARKKTVVQI PAPTTAKQMR EFLGTAGFCR LWIPGFATLA
SLRGGQRWLT EARKKTVVQI PAPTTAKQVR EFLGTAGFCR LWIPGFATLA
SLRGGQRWLT EARKKTVVQI PAPTTAKQVR EFLGTAGFCR LWIPGFATLA
SLRGGQRWLT EARKKTVVQI PAPTTAKQVR EFLGTAGFCR LWIPGFATLA
SLRGGQRWLT EARKKTVVQI PAPTTAKQVR EFLGTAGFCR LWIPGFATLA
SLRGGQRWLT EARKKTVVQI PAPTTAKQVR EFLGTAGFCR LWIPGFATLA
SLRGGQRWLT EARKKTVVQI PAPTTAKQVR EFLGTAGFCR LWIPGFATLA
SLRGGQRWLT EARKKTVVQI PAPTTAKQVR EFLGTAGFCR LWIPGFATLA
SLRGGQRWLT EARKKTVVQI PAPTTAKQVR EFLGTAGFCR LWIPGFATLA
SLRGGQRWLT EARKKTVVQI PAPTTAKQVR EFLGTAGFCR LWIPGFATLA
SLRGGQRWLT EARKKTVVQI PAPTTAKQVR EFLGTAGFCR LWIPGFATLA
SLRGGQRWLT EARKKTVVQI PAPTTAKQVR EFLGTVGFCR LWIPGFATLA
SLRGGQRWLT EAWKKTVVQI PAPTTAKQVR EFLGTAGFCR LWIPGFATLA
SLRDGQRWLT EARKKTVVQI PAPTTAKQVR EFLGTAGFCR LWIPGFATLA
SLRGGQRWLT EARKKTVVQI PAPTTAKQVR EFLGTAGFCR LWIPGFATLA
SLRGGQRWLT EARKRTVVQI PAPTTAKQVR EFLGTAGFCR LWIPGFATLA
SLRGGQRWLT EARKKTVVQI PAPTTAKQVR EFLGTAGFCR LWIPGFATLA
SLRGGQRWLT EARKKTVVQI PAPTTAKQVR EFLGTAGFCR LWIPGFATLA
SLRGGQRWLT EARKRTVVQI PAPTTAKQVR EFLGTAGFCR LWIPGFATLA
SLQGGQRWLT EARKKTVVQI PAPTTAKQVR EFLGTAGFCR LWIPGFATLA
SLRDGQRWLT EARKKTVVQI PAPTTAKQVR EFLGTAGFCR LWIPGFATLA
SLRDGQRWLT EARKKTVVQI PAPTTAKQVR EFLGTAGFCR LWIPGFATLA
SLRDRQRWLT EARKKTVVQI LAPTTAKQVR EFLGTAGFCR LWIPGFATLE
SLRGGQRWLT EARKRTVVQI PAPTTAKQVR EFLGTAGFCR LWIPGFATLA
SLRGGQRWLT EARKKTVVQI PAPTTAKQVR EFLGTAGFCR LWIPGFATLA
SLRDGQRWLT EARKRTVVQI LAPTTAKQVR EFLGTAGFCR LWIPGFTTLA
SLRDEQRWLT EARKRTVVQI LAPTTAKQVR EFLGTAGFCR LWIPGFTTLA
SLRDGQRWLT EARKRTVVQI LAPTTAKQVR EFLGTAGFCR LWIPGFATLA
SLQDGQRWLT EARKRTVAQI PTPTTAKQVR EFLGTA---- ----GFATLA
SLRGGQRWLT EARKKTVVQI PAPTTAKQVR EFLGTAGFCR LWILGFATLT
SLRGGQRWLT EARKRTVVQI PAPTTAKQVR EFLGTAGFCR LWIPGFATLT
SLRGGQRWLT EARKKTVVQI PAPTTAKQVR EFLGTAGFCR LWIPGFATLA
SLRDGQRWLT EARKRTVVQI LVPTTAKQVR EFLGTAGFC- LWIPGFATLA
SLRDGQRWLT EARKRTVVQI LVPTTAKQVR EFLGTAGFC- LWIPGFATLA
SLWGGQRWLT GAWKKTVVQI PAPTTAKQVR E-LGTAGFCR LWIPRFATLA
SLRD-QRWLT EARKRTVVQI PAPTTAKQVR EFLGTAGFCR LWIPGFSTLA
```

SLRDGQRWLT EARKRTVVQI LVPTMSRQFR EFLGTAGFCR LWILGFATLA
VLRDGQRWLT EARKQAVMQI PAPTTARQVR EFLGTAGFCR LWIPGFATLA
VLRDGQRWLT EARKQAVMQI PAPTTARQVR EFLGTAGFCR LWIPGFATLA
VLRDGQRWLT EARKQAVMQI PAPTTARQVR EFLGTAGFCR LWIPGFATLA
VLKDEQRWLT EAKKQAVMQI PTPTTARQVR EFLGTAGFCR LWIPGFATLA
TLRGGKRWLT EARKKTVMMI PSPTTPRQVR EFLGTAGFCR LWIPGFATLA
TLRGGKRWLT EARKKTVMMI PSPTTPRQVR EFLGTAGFCR LWIPGFATLA
TLRGGKRWLT EARKKTVMMI PSPTTPRQVR EFLGTAGFCR LWIPGFATLA
TLRGGKRWLT EARKKTVMMI PSPTTPRQVR EFLGTAGFCR LWIPGFATLA
TLRGGKRWLT EARKKTFMMI PSPTTPRQVR EFLGTAGFCR LWIPGFATLA
TLRGGKRWLT EARKKTVMMI PSPTTPRQVR EFLGTAGFCR LWIPGFATLA
TLRGGKRWLT EARKKTVMMI PPPTTPRQVR EFLGTAGFCR LWIPGFATLA
TLREGKRWLT EARKKTVMQI PTPTTPRQVR EFLGTAGFCR LWIPGFATLA
LLKGGKRWLT PARKATVMKI PTPTTPRQVR EFLGTAGFCR LWIPGFASLA
LLKEGKRWLT PARKATVMKI PVPTTPRQVR EFLGTAGFCR LWIPGFASLA
ILRDGKRWLT EARKRTVTQI PTPATPRQVR EFLGTAGFCR LWIPGFATLA
ILRDGKRWLT EARKRTVTQI PTLATPRQVR EFLGTAGFCR LWIPGFATLA
IL----RWLT EARKRTVTQI PTLATPRQVR EFLGTAGFCR LWIPGFATLA
ILWDGKRWLT EARKRTVTQI PTLATPRQVR EFLGTAGFCR LWIPGFATLA
ILWDGKRWLT EARKRTVTQI PTLATPRQVR EFLGTAGFCR LWIPGFATLA
TL-DGKQWLM EARKKTVTQI PVPATPRQVR EFLGTAGFCR LWIPGFATLA
SLWDGKRWLA EAQKRTVVQI LVPTMARQVR EFLGTAGFCR LWILGFATLA
ILRDGKRWLT EARKKTVLQI STPQMPLQVR EFLGTAGFCR LWIPGFAALA
TLRGGKRWLT EAQKKTVTQI PIPTTSKQVR EFLGIAGFCR LWIPSFATLA
SLKEGKRWLT EARKKTVTQI PVPTTARQVR EFLGTAGFCR LWIPGFAPMA
LLKEGQQWLT EARKKTVTSI PVPRNPRQVH EFLGTAGYCR IWTTGFATLA
ILKDGKRWLT EARKKTVTQI PVPSTHRQVR EFLGTAGFCR LWIPRYATLA
ILKDGKRWLT KALKKTVTQI PVPSTHRQVR EFLSTAGFCQ LWIPRYASLV
ILKDGKRWLT EARKKTVTQI PVPSTHRQVR EFLGTAGFCR LWIPRYASL-
VLRNGQ-WLT DTRKQTVIRI PTPTTPRQVR EFLGTAGFCR LWIPGFATLA
ALRDGKRWLM EAQKKTVTQI PTPTTPLQVR EFLGKVGFCR LWIPGFTTL-
LLKEGQRWLT EARKETVMGQ PTPKTPRQLR EFLGTAGFCR LWIPGFAEMA
LLKEGQRWLT EARKETVMGQ PTPKTPRQLR EFLGTAGFCR LWIPGFAEMA
LLKEGQRWLT EARKETVMGQ PTPKTPRQLR EFLGTAGFCR LWIPGFAEMA
LLKEGQRWLT EARKETVMGQ PTPKTPRQLR EFLGTAGFCR LWIPGFAEMA
LLKEGQRWLT EARKETVMGQ PIPKTPRQLR EFLGTAGFCR LWIPGFAEMA
LLKEGQRWLT EARKETVMGQ PIPKTPRQLR EFLGTAGFCR LWIPGFAEMA
LLREGQRWLT EARKETVMGQ PVPKTPRQLR EFLGTAGFCR LWIPGFAEMA
LLKEGQRWLT EARKETVMGQ PTPKTPRQLR EFLGTAGFCR LWIPGFAEMA
LLKEGQRWLT EARKETVMGQ PTPKTPRQLR EFLGTAGFCR LWIPGFAEMA
LLKEGQRWLT EARKETVMGQ PTPKTPRQLR EFLGTAGFCR LWIPGFAEMA
QLRDGQRWLT PARKQTVTGI PAPKNGRQLR -FLGKAGFCR LWIPGFAEMA
QLRDGQRWLT PARKQTVTGI PAPKNGRQLR -FLGKAGFCR LWIPGFAEMA
SLKDGQRWLT KARKEAILSI PVPKNSRQVR EFLGTAGYCR LWIPGFAELA

ILSEGKRWLT PGRIETVARI PPPQSPREVR EFLGTAGFCR LWIPGFAELA
ILSEGKRWLT PGRIETVARI PPPRNPREVR EFLGTAGFCR LWIPGFAELA
LLKGGQRWLT ESRKDTVAQI PAPKNARQVR EFWGMAGFCR LWIPGFAKLT
SLKEGKWWLT EVRKKTVTQI PVPTTP-QVK EFLGKAGFCR LWIPGFATMA
SLRDGQRWLT EARKKTVVQI PAPTTAKQVE RVLGTAGFCR LWIPGFATL-
TLHDGKRWLT EARKQTVTQI LAPTTPCQVR EFLGTAGFCR LWIPGYSLFL
TL-DGKRWLM EARKKIVTQM P-----PQLH EFLGTAGFCR LWIPGFVTSA
KLKEGTRWLT EAMKETILRL PVPTSAREVR EFLGTTGYCR LWILGYAEIA
TL-DGKQWLA EARKRTVTQI LALATPRQVR EFLGTTGFCR LWIPVFATLA
KIHKGSRSLS NSRTQAILQI PVPKTKRQVR EFLGTIGYCR LWIPGFAELA
TLRDGKQWLT EARKRTVTQI PTLATPRQVR EFLGTSGFCR LWKPGLATPV
TLRDLKQWLT EARKRTVTQI PTLATPRQVR EFLGTAGFCR LWKPGFATLA
TLR-GRKWLP EAEKDCDSN- ---PNPNNLE EGVGISG--H GWVPKLCQPG
SVRVRKWWVT EACKKTVTQI PLPTSLTQVR EFVGMAGFCR L-IPSYATLA
SLRKRKWWLT EAWKKTVTQI HIPTTLKQVR EFVGMAGFCR L-IPSFASLA

APLYPLTKEK GEFSWAPEHQ KAFDAIKKAL LSAPALALPD VTKPFTLYVD
APLYPLTKEK GEFSWAPEHQ KAFDAIKKAL LSAPALALPD VTKPFTLYVD
APLYPLTKEK GEFSWAPEHQ KAFDAIKKAL LSAPALALPD VTKPFTLYVD
APLYPLTKEK GEFSWAPEHQ KAFDAIKKAL LSAPALALPD VTKPFTLYVD
APLYPLTKEK GEFSWAPEHQ KAFDAIKKAL LSAPALALPD VTKPFTLYVD
APLYPLTKEK GEFSWAPEHQ KAFDAIKKAL LSAPALALPD VTKPFTLYVD
APLYPLTKEK GEFSWAPEHQ KAFDAIKKAL LSAPALALPD VTKPFTLYVD
APLYPLTKEK GEFSWAPEHQ KAFDAIKKAL LSAPALALPD VTKPFTLYVD
APLYPLTKEK GEFSWAPEHQ KAFDAIKKAL LSAPALALPD VTKPFTLYVD
APLYPLTKEK GEFSWAPEHQ KAFDAIKKAL LSAPALALPD VTKPFTLYVD
APLYPLTKEK GEFSWAPEHQ KAFDAIKKAL LSAPALALPD VTKPFTLYVD
APLYPLTKEK GEFSWAPEHQ KAFDAIKKAL LSAPALALPD VTKPFTLYVD
APLYPLTKEK GEFSWAPEHQ KAFDAIKKAL LSAPALALPD VTKPFTLYVD
APLYPLTKEK GEFSWAPEHQ KAFDAIKKAL LSAPALALPD VTKPFTLYVD
APLYPLTKEK GEFSWAPEHQ KAFDAIKKAL LSAPALALPD VTKPFTLYVD
APLYPLTKEK GEFSWAPEHQ KAFDAIKKAL LSAPALALPD VTKPFTLYVD
APLYPLTKKK GEFSWAPEHQ KAFDAIKKAL LSAPALALPD VTKPFTLYVD
APLYPLTKEK G-FSWAPEHQ KAFDAIKKAL LSAPALALPD VTKHFTLYVD
APLYPLTKEK GEFSWAPEHQ KAFDAIKKAL LSAPALALPD VTKPFTLYVD
APLYPLTKEK GEFSWAPEHQ KAFDAIKKAL LSAPALALPD VTKPFTLYVD
APLYPLTKEK GEFSWAPEHQ KAFDAIKKAL LSAPALALPD VTKPFTLYVD
APLYPLTKEK GEFSWAPEHQ KAFDAIKKAL LSAPALALPD VTKPFTLYVD
APLYPLTKEK GEFSWAPEHQ KAFDAIKKAL LSAPALALPD VTKPFTLYVD
APLYPLTKEK GEFSWAPEHQ KAFDAIKKAL LSAPALALPD VTKPFTLYVD
APLYPLTKEK GEFSWAPEHQ KAFDAIKKAL LSAPALALPD VTKPFTLYVD
APLYPLTKEK GEFSWAPEHQ KAFDAIKKAL LSAPALALPD VTKPFTLYVD
APLYPLTKEK GEFSWAPEHQ KAFDAIKKAL LSAPALALPD VTKPFTLYVD
APLYPLTKEK GEFSWAPEHQ KAFDAIKKAL LSAPALALPD VTKPFTLYVD

APLYPLTKEK GGFSWAPEHQ KAFDAIKKAL LSAPALALPD VTKPFTLYVD
APLYPLTKEK GEFSWAPEHQ KAFDAIKKAL LSAPALALPD VTKPFTLYVD
APLYPLTKEK GEFSWAPEHQ KAFDAIKKAL LSAPALALPD VTKPFTLYVD
APLYPLTKEK GEFSWAPEHQ KAFDAIKKAL LSAPALALPD VTKPFTLYVD
APLYPLTKEK GEFSWAPEHQ KAFDAIKKAL LSAPALALPD VTKPFTLYVD
APLYPLTKEK GEFSWAPEHQ KAFDAIKKAL LSAPALALPD VTKPFTLYVD
APLYPLTKEK GEFSWAPEHQ KAFDAIKKAL LSAPALALPD VTKPFTLYVD
APLYPLTKEK GEFSWAPEHQ KAFDAIKKAL LSAPALALPD VTKPFTLYVD
APLYPLTKEK GEFSWAPEHQ KTFDAIKKAL LSTPALALPD VTKPFTLYVN
APLYQLTKEK GEFSWAPEHQ KAFDAIKKAL LSAPALALPD VTKPFTLYVD
APLYPLTKEK GEFSWAPEHQ KAFDAIKKAL LSAPALALPD VTKPFTLYVE
APLYPLTKEK REFSWAPEHQ KAFDAIKKAL LSAPALALPD VTKPFTLYVD
APLYPLTKEK GEFSWAPEHQ KAFDAIKKAL LSAPALALPD VTKPFTLYVD
APLYPLTKEK GEFSWAPEHQ NAFDAIKKAL LSAPALALSD VTKPFTLYVD
APLYPLTKEK GEFSWAPEHQ KAFDAIKKAL LSAPALALPD VTKPFTLYVD
APLYPLSKEK GEFSWAPEHQ KAFDAIKKAL LSTPALALPD ITKPFTLYVD
APLYPLTKEK GEFSWAPEHQ KAFDAIKKAL LSTPALALPD VTKPFTLYVD
APLYPLTKEK GEFSWAPEHQ KAFDAIKKAL LSAPALALPD VTKPFTLYVD
APLYPLTKEK GEFSWAPEHQ KAFDAIKKAL LSTPALALPD VTKPFTLYVD
APLYPLTKEK GEFSWAPEHQ KAFDAIKKAL LSTPALALPD VTKPFTLYVN
APLYPLTKEK GEFSWAPEHQ KAFDAIKKAL LSAPALALPD VTKPFTLYVD
APLYPLTKEK GEFSWAPEHQ KAFDAIKKAL LSAPALALPD VTKPFTLYVD
APLYPLTKEK GEFSWAPEHQ KAFDASKKAL LSAPALALPD VTKPFTLYVD
APLYPLTKEK GKFSWAPEHQ KTFDAIKKAL LSAPALALPD VTKPFTLYVD
APLYPLTKEK GEFSWAPEHQ KAFDAIKKAL LSAPALALPD VTKPFTLYVD
APLYPLTKEK GEFSCAPKHQ KAFDAIKKAL LSAPALALPD VTKPFTLYVD
APLYPLTKEK GEFSCAPEHQ KAFDAIKKAL LSAPALALPD VTKPFTLYVD
APLYPLTKEK GEFSWAPEHQ KAFDAIKKAL LSAPALALPD VTKPFTLYVD
APLYPLTKEK GEFSWAPEHQ KAFDAIKKAL LSAPALALPD VTKPFTLYVD
APFYSLTKEK GKFTWTSEHQ GAFDAVKKAL LSVPALALLD MTKPFTLYVD
APLYPLTKEK GEFTWTREHQ LAFETLKKAL LQAPALALPD LNKPFTLYID
APLYPLTKEK GEFTWTREHQ LAFETLKKAL LQAPALALPD LNKPFTLYID
APLYPLTKEK GEFTWTREHQ LAFETLKKAL LQTPALALPD LNKPFTLYID
ALLYPLTKEK GEFTWTREHQ LAFETLKKAL LQAPALALPD LNKPFTLYID
APLYPLTKEG VPFEWKEEHQ RAFEAIKSSL MTAPALALPD LTKPFVLYVD
APLYPLTKEG VPFEWKEEHQ RAFEAIKSSL MTAPALALPD LTKPFVLYVD
APLYPLTKKG VPFEWKEEHQ RAFEAIKSSL MTAPALALPD LTKPFVLYVD
APLYPLTKEG VPFEWKEEHQ RAFEAIKSSL MTAPAIALPD LTKPFVLYVD
APLYPLTKEG VPFEWKEEHQ RAFEAIKSSL MTAPALALPD LTKPFVLYVD
APLYPLTKEG VPFEWKEEHQ RAFEAIKSSL MTAPALALPD LTKPFVLYVD
APLYPLTREG IPFEWKEEHQ RAFEAIKSSL MTAPALALPD LTKSFVLYVD
APLYPLTKEK VPFTWTEEHQ RAFEDIKAAL LAAPALALPD LTKPFTLYVD
APLYPLTREK VPFTWTEAHQ EAFGRIKEAL LSAPALALPD LTKPFALYVD
APLYPLTKES IPFIWTEEHQ QAFDHIKKAL LSAPALALPD LTKPFTLYID

```
APLYPLTKES GEFRWTSEHQ KAFENIKKAL LTAPALALPD LTKPFILYVD
APLYPLTKES GEFRWTSEHQ KAFENIKKAL LTASALALPD LTKPFILYVD
APLYPLTKES GEFRWTSEHQ KAFENIKKAL LTAPALALPD LTKPFILYVD
APLYPLTKES GEFRWTSEHQ KAFKNIKKAL LTAPALALPD LTKPFILYVD
APLYPLTKES GEFRWTSEHQ KAFKNIKKAL LTAPALALPD LTKPFILYVD
APLYPLTKES GEFK-TPEHQ RAYN-IKEAL LTVPALAMPD LTKPFTLYVD
APLYSLTKEK EGFTWTSKHQ RVFDTIKKAL LSAPALALSD VTKPFTLYVD
APLHSLTRSE TEFQWKPEHQ AAFDAIKQAL LTTPALVLPD LTKPFTLFVD
APLYPLTKET GTFSW---HQ EAFKEIKKAL LSALPLALPD LTRPVTLYVD
APLYPLTKEG KTFSWTPNHQ KAFNEIKKAL LSAPALALTY LTKPFTLYVD
APLYSLT-EG VDFNWQSEHQ KAFEQLKKAL LSAPALALPN LTKPFYLYVN
APLYPLTKNS TPFIWGPGQQ EAFDAIKQAL LTAPALALPD VTKPFTLFID
APLYPLTKIS TPFTWGPEQQ EAFDTIKQAL LTALALALPD V----TLFVD
APLYPLTKNS TPFARGPEQQ EAFDTIKQA- LTAPALALPD VTKPFTLFVD
APLYSITKER VEFIWTQEHQ SAFETLQKGL LQAPALALPD VNKPFTLYID
--LYPLTKEG GEFEWTETQQ EAFDKIKQ-- ---------D LTEPFTLYVD
APLYPLTKTG TLFNWGPDQQ KAYQEIKQAL LTAPALGLPD LTKPFELFVD
APLYPLTKTG TLFNWGPDQQ KAYQEIKQAL LTAPALGLPD LTKPFELFVD
APLYPLTKTG TLFNWGPDQQ KAYQEIKQAL LTAPALGLPD LTKPFELFVD
APLYPLTKTG TLFNWGPDQQ KAYQEIKQAL LTAPALGLPD LTKPFELFVD
APLYPLTKTG TLFNWGPDQQ KAYQEIKQAL LTAPALGLPD LTKPFELFVD
APLYPLTKTG TLFNWGPDQQ KAYQEIKQAL LTAPALGLPD LTKPFELFVD
APLYPLTKTG TLFNWGPDQQ KAYQEIKQAL LTAPALGLPD LTKPFELFVD
APLYPLTKTG TLFNWGPDQQ KAYQEIKQAL LTAPALGLPD LTKPFELFVD
APLYPLTKTG TLFEWGPDQQ KAYQEIKQAL LTAPALGLPD LTKPFELFVD
APLYPLTKTG TLFNWGPDQQ KAYQEIKQAL LTAPALGLPD LTKPFELFVD
APLYPLTKLG VLFQWGPEQQ KAFNNIK-AL LSSPALGLPD ITKPFELFID
APLYPLTKLG VLFQWGPEQQ KAFNNIK-AL LSSPALGLPD ITKPFELFID
APLYPLTRPG TLFQWGTEQQ LAFEDIKKAL LSSPALGLPD ITKPFELFID
APLYALTKES APFTWQEKHQ SAFEALKEAL LSAPALGLPD TSKPFTLFID
APLYALTKES TPFTWQTEHQ LAFEALKKAL LSAPALGLPD TSKPFTLFLD
APLYPLTKNS TPFVWGDKEQ RAFDQIKRAL LSAPALGLPD VTKPFHLYVA
APLNPLTKEL KTIS-TPDHQ KAFDEIKKAL LSSPALALPD LTKPFTLFVD
APLYPLTKEK GEFS-APEHQ KA-DAIKKAL LSAPALALPD VT-PFTLY-D
WLLSTLTKEG GKFHWTSEFQ EAFDKIKQAL LPAPALALPD LTKLFTLYVD
APLYPLTKEG GVCV-DPRPP ESLGN-QEGP TDGPSLS-PA LNQALYFYVD
KPLYEATKDK VPWAWGSDQQ KAYDELKVAL LRAPALALPD PLKPFTLFVD
APLNPLNKES GEFRWISEYQ RDFENIKEAL LTAPALALTD ITKPFILYVD
QPLYAATRGG NPLVWGEKEE EAFQSLKLAL TQPPALALPS LDKPFQLFVE
----PLDQEK ----WDIRTS ESLKYGGPAD CASLGIARSN QT---LLPLC
TLLYPLTKKS GEFRWTSEHQ KAFENIMEAL LTVPALALPD LTKPCILYVD
SPTLPPYRD- ----WCPIHQ EAFKEIKKAL LSIHGLGLPD LTKPFTLYVD
ATLYPFTKET GTFS-TPIHQ YAFKEIKKLS LS-------- ------IYIE
ATLYPFTKET GIFSWTPIHQ -ACKEIKKAL LSAPALSLQD LTKPVTLY--
```

ERKGVARGVL TQTLGPWRRP VAYLSKKLDP VASGWPICLK AIAAVAILVK
ERKGVARGVL TQTLGPWRRP VAYLSKKLDP VASGWPICLK AIAAVAILVK
ERKGVARGVL TQTLGPWRRP VAYLSKKLDP VASGWPICLK AIAAVAILVK
ERKGVARGVL TQTLGPWRRP VAYLSKKLDP VASGWPVCLK AIAAVAILVK
ERKGVARGVL TQTLGPWRRP VAYLSKKLDP VASGWPICLK AIAAVAILVK
ERKGVARGVL TQTLGPWRRP VAYLSKKLDP VASGWPVCLK AIAAVAILVK
ERKGVARGVL TQTLGPWRRP VAYLSKKLDP VASGWPVCLK AIAAVAILVK
ERKGVARGVL TQTLGPWRRP VAYLSKKLDP VASGWPVCLK AIAAVAILVK
ERKGVARGVL TQTLGPWRRP VAYLSKKLDP VASGWPVCLK AIAAVAILVK
ERKGVARGVL TQTLGPWRRP VAYLSKKLDP VASGWPVCLK AIAAVAILVK
EHKGVARGVL TQTLGPWRRP VAYLSKKLDP VASGWPVCLK AIAAVAILVK
ERKGVARGVL TQTLGPWRRP VAYLSKKLDP VASGWPICLK AIAAVAILVK
ERKGVARGVL TQTLGPWRRP VAYLSKKLDP VASGWPVCLK AIAAVAILVK
ERKGVARGVL TQTLGPWRRP VAYLSKKLDP VASGWPVCLK AIAAVAILVK
ERKGVARGVL TQTLGPWRRP VAYLSKKLDP VASGWPVCLK AIAAVAILVK
ERKGVARGVL TQTLGPWRRP VAYLSKKLDP VASGWPICLK AIAAVAILVK
ERKGVARGIL TQTLGPWRRP VAYLSKKLDP VASGWPICLK AIAAVAILVK
ERKGVARGVL TQTLGPWRRP VAYLSKKLDP VASGWPVCLK AIAAVAILVK
ERKGVARGVL TQTLGPWRRP VAYLSKKLDP VASGWPICLK AIAAVAILVK
ERKGVARGVL TQTLGPWRRP VAYLSKKLDP VASGWPVCLK AIAAVAILVK
ERKGVARGVL TQTLGPWRRP VAYLSKKLDP VASGWPVCLK AIAAVAILVK
ERKGVARGVL TQTLGPWRRP VAYLSKKLDP VASGWPICLK AIAAVAILVK
ERKGVARGVL TQTLGPWRRP VAYLSKKLDP VASGWPVCLK AIAAVAILVK
ERKGVARGVL TQTLGPWRRP VAYLSKKLDP VASGWPVCLK AIAAVAILVK
ERKGVARGVL TQTLGPWRRP VAYLSKKLDP VASGWPVCLK AIAAVAILVK
ERKGVARGVL TQTLGPWRRP VAYLSKKLDP VASGWPVCLK AIAAVAILVK
ERKGVARGVL TQTLGPWRRP VAYLSKKLDP VASGWPVCLK AIAAVAILVK
ERKGVARGVL TQTLGPWRRP VAYLSKKLDP VASGWPVCLK AIAAVAILVK
ERKGVARGVL TQTLGPWRRP VAYLSKKLDP VASGWPVCLK AIAAVAILVK
ERKGVARGVL TQTLGPWRRP VAYLSKKLDP VASGWPVCLK AIAAVAILVK
ERKGVARGVL TQTLGPWRRP VAYLSKKLDP VASGWPVCLK AIAAVAILVK
ERKGVARGVL TQTLGPWRRP VAYLSKKLDP VASGWPVCLK AIAAVAILVK
ERKGVAQGVL TQTLGPWRRP VAYLSKKLDP VASGWPVCLK AIAAVAILVK
ERKGVARGVL TQTLGPWRRP VAYLSKKLDP VASGWPVCLK AIAAVAILVK
ERKGVARGVL TQTLGPWRRP VAYLSKKLDP IASGWPVCLK AIAAVAILVK
ERKGVARGVL TQTLGPWRRP VAYLSKKLDP VASGWPICLK AIAAVAILVK
ERKGVARGVL TQTLGPWRRP VAYLSKKLDP VASGWPVCLK AIAAVAILVK
ERKGVARGVL TQTLGPWRRP VAYLSKKLDP VASGWPVCLK AIAAVAILVK
ERKGVARGVL TQTLGPWRRP VAYLSKKLDP VASGWPVCLK AIAAVAILVK
ERKGVARGVL TQTLGPWRRP VAYLSKKLDP VASGWPVCLK AIAAVAILVK
ERKGVARGVL TQTLGPWRRP VAYLSKKLDP VASGWPVCLK AIAAVAILVK
EHKGVARGVL TQTLGPWRRP VAYLSKKLDP VASGWPVCLK AIAAVAILVK
ERKGVARGVL T-TLGPWRRP VAYLSKKLDP IASGWPVCLK AIAAVAILVK

ECNGVARGVL TQTLGPWRRP VAYLSKKLDP VASGWPVCLK AIAAVAILVK
ERKGVARGVL TQTLGPWRRP VAYLSKKLDP VASGWPICLK AIAAVAILVK
ERKGVARGVL TQTLGPWRRP VAYLSKKLDP VASGWPVCLK AIAAVAILVK
ERKGVARGVL TQSLGPWRRP VAYLSKKLDP VASGWPICLK AIAAVAILVK
ERKGVARGVL TQSLGPWRRP VAYLSKKLDP VASGWPICLK AIAAVAILVK
EHKGVARGVL TQSLGPWRRP VAYLSKKLDP VASGWPVCLK AIAAVAILVK
ERKGIARGVL TQTLGPWRRP VAYLSKKLDP VASGWPVCLK AIAAVAILVK
ERKGVARGVL TQTLGPWRRP VAYLSKKLDP VASGWPICLK AIAALAILVK
EHKEIARGVL TQTLGPWRRP VAYLSKKLDP VASGWPVCLK AIAAVAILVK
ERKGVAQGVL TQSLGPWRRP VDYLSKKLDP VASGWPVCLK AIAAMAILVK
ERKGVAQGVL TQSLGPWRRP VDYLSKKLDP VASGWPVCLK AIAAMAILVK
ERKGVAQGVL TQSLGPWRRP VDYLSKKLDP VASGWPVCLK AIAAMAILVK
ERKGVARGVL TQTLGPWRRP VAYLSKKLDP VASGWPVCLK AIAAVAILVK
ECKGVAQGVL TQSLGPWRRP VAYLSKKLDP VASGWPVCLK AIAAVAILVK
ERKGIARGVL TQSLGPWRRP IAYLSKKLDP VASGWPVCLK AIVAVSTLIK
ERNGVARGVL TQVLGPWKRP VAYLSKKLDA VASGWPSCLR AIAATAVLVK
ERNGVARGVL TQVLGPWKRP VAYLSKKLDA VASGWLSCLR AIAATAVLVK
EKNGVARGVL TQVLGPWKRP VAYLSKKLDA VASGWPSCLR AIAATAVLVK
KKNGVARGVL TQVLGP-KRP VAYLSKKLDA VASGWPSYLR AIAATAVLVR
ERAGVARGVL TQALGPWKRP VAYLSKKLDP VASGWPTCLK AIAAVALLIK
ERAGVARGVL TQALGPWKRP VAYLSKKLDP VASGWPTCLK AIAAVALLIK
ERAGVARGVL TQALGPWKRP VAYLSKKLDP VASGWPTCLK AIAAVALLIK
ERAGVARGVL TQALGPWKRP VAYLSKKLDP VASGWPTCLK AIAAVALLIK
ERAGVARGVL TQALGPWKRP VAYLSKKLDP VASGWPTCLK AIAAVALLIK
ERAGVARGVL TQALGPWKRP VAYLSKKLGP VASGWPTCLK AIAAVALLIK
ERAGIARGVL TQALGPWKRP VAYLSKKLDP VASGWPTCLK AIAAVALLIK
ERAGVARGVL TQALGPWKRP VAYLSKKLDP VASRWPSCLK AIAAVALLVK
EKEGVARGVL TQTLGPWRRP VAYLSKKLDP VASGWPTCLK AIAAVALLLK
ERAGVARGVL TQTLGPWRRP VAYLSKKLDP VASGWPTCLK AVAAVALLLK
ERAGVARGVL TQALGPWKRP VAYLSKKLDP VASGWPTCLK AVAAVALLVK
ERAGVARGVL TQALGPWKRP VAYLSKKLDP VASGWPTCLK AVAAVALLVK
ERAGVARGVL TQALGPWKRP VAYLSKKLDP VGSGWPTCLK AVAAVALLVK
ERAGVARGVL TQALGPWKRP VAYLSKKLDP VGSGWPTCLK AVAAVALLVK
ERAGVARGVL TQALGPWKRP VAYLSKKLDP VGSGWPTCLK AVAAVALLVK
ERAGEARGVL TQTLGPWKRP MAYLSKKLDP VASEWPTCLK AVATIALLVK
EHKGIAQGVL TQSLGPWRRP VAYLSKKLDS VASGWPVCLK SIVAMPTLIK
EKAGVAKGVL TQTLGPWKRP VAYLSKKLDP VASGWPSCLK IIAAVAILVK
ERAGIARGVL TQALGP-KRL VAYLSKKLDP VASGWPSCLK AITAVALLLK
ERAGVARGVL MQTLGPWKRP VAYLSKKLDP VVSG-PSHLR VIAAVALLVK
ERAGEA-GVL TQPLGPWKRP VAYLSKKLDP VASG-PFCLK AIAAVTVLVK
ERKGVARGVL TQSLGPWKRP VAYLSKKLDP VASGWPTCLQ AIAAVASLVR
ERKGVARGVL TQPLGPWKRP VAYLSKKLDL VASGWPTCLR AIAAVASLVK
ERKGVARGVL TQLLGPWKRP VAYLSKKLDP VASGWPTCLR AIAAVASLVK
ERNGVARGVL TQTIGPLKHP VAYLSKKLDS VASGWPSCL- AIAATVVLVK

```
Q-AGIPRGVL TQTLGPWKRP MAYLSKKLDP VTSGWPSCLK AIAAMALLVK
EKQGYAKGVL TQKLGPWRRP VAYLSKKLDP VAAGWPPCLR MVAAIAVLTK
EKQGYAKGVL TQKLGPWRRP VAYLSKKLDP VAAGWPPCLR MVAAIAVLTK
EKQGYAKGVL TQKLGPWRRP VAYLSKKLDP VAAGWPPCLR MVAAIAVLTK
EKQGYAKGVL TQKLGPWRRP VAYLSKKLDP VAAGWPPCLR MIAAIAVLTK
EKQGYAKGVL TQKLGPWRRP VAYLSKKLDP VAAGWPPCLR MVAAIAVLTK
EKQGYAKGVL TQKLGPWRRP VAYLSKKLDP VAAGWPPCLR MVAAIAVLTK
EKQGYAKGVL TQKLGPWRRP VAYLSKKLDP VAAGWPPCLR MVAAIAVLTK
EKQGYAKGVL TQKLGPWRRP VAYLSKKLDP VAAGWPPCLR MVAAIAVLTK
EKQGYAKGVL TQKLGPWRRP VAYLSKKLDP VAAGWPPCLR MVAAIAVLTK
EKQGYAKGVL TQKLGPWRRP VAYLSKKLDP VAAGWPPCLR MVAAIAVLTK
KKQGYAKGVL TQRLGPWKRP IAYLSKKLDR VASGWPPCLR MVAAIAVLTK
KKQGYAKGVL TQRLGPWKRP IAYLSKKLDR VASGWPPCLR MVAAIAVLTK
ENSGFAKGVL VQKLGPWKRP VAYLSKKLDT VASGWPPCLR MVAAIAILVK
EKQGIAKGVL TQKLGPWKRP VAYLSKKLDP VAAGWPPCLR IMAATAMLVK
ERQGIAKGVL TQKLGPWKRP VAYLSKKLDP VAAGWPPCLR IMAATAMLVK
ENKGIAKGVL TQKLGPWNRP VAYLSKKMDP VVSGWPTCLK IIAAVAVLVK
ERAGVDSGVL TQTLAPWKRP GAYLFKKLDP VSSGWPSCLR ATVAVALLVK
ERKGVARGVL T-TLGPWRRP VAYLSKKLDP IASGWPVCLK AIAAVAILVK
EQAGVARGVI TQSLGPWKRP VAYLSKKLDP VASGWPSCLK AIAAVALLVK
ERAGVARGVL TQTLGPWERP VAYLSKKLDP VASGWVYCLK AIAAVALLVK
ERRGIAKGVL MQRLGPWKRP VAYLSKKLDP VAAGWPPCLR IIAAVALMVK
ERLGVARGVL TQARGPWKRP KAYLSKKLDP VASR-PKCLK VVAAVALLIK
ETSGAAKGVL TQALGPWKRP VAYLSKRLDP VAAGWPRCLR AIAAAALLTR
EGVG-AKGVL TQALGPWNRL VEYSSKKLDP VARRWPTCLK AVAAVALLIK
ERAWGARGVL TQALGPWKRL VAYLSKELDP VARRWPTFLK SVAAVALLI-
ERTGIARGVL TQALGP-KTP VAYLSKNLEP VASGWPSCLK AITAVGLLLK
E-AGIESGDL T-ALGP-KQP VEYWSKKLDP VSSGWPFC-K AIDALALLLK
--RGMSRGVL I-ALGP-KQP VKYWSKKLDP VSSGWPFCLK AINALALLLK

DADKLTLGQN ITVTAPHALE NTVRQPPDRW MTNARMTHYQ SLLLTERVTF
DADKLTLGQN ITVIAPHALE NIVRQPPDRW MTNARMTHYQ SLLLTERVTF
DADKLTLGQN ITVIAPHALE NIVRQPPDRW MTNARMTHYQ SLLLTERVTF
DADKLTLGQN ITVIAPHALE NIVRQPPDRW MTNARMTHYQ SLLLTERVTF
DADKLTLGQN ITVIAPHALE NIVRQPPDRW MTNARMTHYQ SLLLTERVTF
DADKLTLGQN ITVIAPHALE NIVRQPPDRW MTNARMTHYQ SLLLTERVTF
DADKLTLGQN ITVIAPHALE NIVRQPPDRW MTNARMTHYQ SLLLTERVTF
DADKLTLGQN ITVIAPHALE NIVRQPPDRW MTNARMTHYQ SLLLTERVTF
DADKLTLGQN ITVIAPHALE NIVRQPPDRW MTNARMTHYQ SLLLTERVTF
DADKLTLGQN ITVIAPHALE NIVRQPPDRW MTNARMTHYQ SLLLTERVTF
DADKLTLGQN ITVIAPHALE NIVRQPPDRW MTNARMTHYQ SLLLTERVTF
DADKLTLGQN ITVIAPHALE NIVRQPPDRW MTNARMTHYQ SLLLTERVTF
DADKLTLGQN ITVIAPHALE NIVRQPPDRW MTNARMTHYQ SLLLTERVTF
DADKLTLGQN ITVIAPHALE NIVRQPPDRW MTNARMTHYQ SLLLTERVTF
```

DADKLTLGQN ITVIAPHALE NIVRQPPDRW MTNARMTHYQ SLLLTERVTF
DADKLTLGQN ITVI-PHALE NIVRQPPDRW MTNARMTHYQ SLLLTERVTF
DADKLTLGQN ITVIAPHTLE NIVRQPPDRW MTNARMTHYQ SLLLTERVTF
DADKLTLGQN ITVIAPHALE NIVRQPPDRW MTNARMTHYQ SLLLTERVTF
DADKLTLGQN ITVIAPHALE NIVRQPPDRW MTNARMTHYQ SLLLTERVTF
DADKLTLGQN ITVIAPHALE NIVRQPPDRW MTNARMTHYQ SLLLTERVTF
DADKLTLGQN ITVIAPHALE NIVRQPPDRW MTNARMTHYQ SLLLTERVTF
DADKLTLGQN ITVIAPHALE NIVRQPPDRW MTNARMTHYQ SLLLTERVTF
DADKLTLGQN ITVIAPHALE NIVRQPPDRW MTNARMTHYQ SLLLTERVTF
DADKLTLGQN ITVIAPHALE NIVRQPPDRW MTNARMTHYQ SLLLTERVTF
DADKLTLGQN ITVIAPHALE NIVRQPPDRW MTNARMTHYQ SLLLTERVTF
DADKLTLGQN ITVIAPHALE NIIRQPPDRW MTNARMTHYQ SLLLTERVTF
DADKLTLGQN ITVIAPHALE NIVRQPPDRW MTNARMTHYQ SLLLTERVTF
DADKLTLGQN ITVIAPHALE NIVRQPPDRW MTNARMTHYQ SLLLTERVTF
DADKLTLGQN ITVIAPHALE NIVRQPPDRW MTNARMTHYQ SLLLTERVTF
DADKLTLGQN ITVIAPHALE NIVRQPPDRW MTNARMTHYQ SLLLTERVTF
DADKLTLGQN ITVIAPHALE NIVRQPPDRW MTNARMTHYQ SLLLTERVTF
DADKLTLRQN ITVIAPHALE NIVRQPPDRW MTNARMTHYQ SLLLTERVTF
DADKLTLGQN ITVIAPHALE NIVRQSPDRW MTNARMTHYQ SLLLTERVTF
DADKLTLGQN ITVIAPHALE NIVRQPPDRW MTNARMTHYQ SLLLTERVTF
DADKLTLGQN ITIIAPHALE NIVRQPPDRW MTNARMTHYQ SLLLTERIMF
DADKLTLGQN ITVIAPHALE NIVRQPPDRW MTNARMTHYQ SLLLTERVTF
DADKLTLGQN ITVIAPHALE NIVRQPPDRW MTNARMTHYQ SLLLTERVTF
DADKLTLGQN ITVIAPHALE NIVRQPPDRW MTNARMTHYQ SLLLTERVTF
DADKLTLGQN ITVIAPHALE NIVRQPPDRW MTNA-MTHYQ SLLLTERVTF
DADKLTLGQN ITVIAPHALE NIVRQPPDRW MTNARMTHYQ SLLLTERVTF
DADKLTLGQN ITVIAPHALE NIVRQPPDRW MTNARMTHYQ SLLLTERVTF
DADKLTLGQN ITIIAPHALE NIIRQPPDRW MTNACMSHYQ SLLLTERITF
DADKLTLGQN ITIIAP-ALE NIVRQPPDRW MTNARMTHYQ SLLLTERITF
DADKLTLGQN ITIIAPHALE NIVQQPPDRW MTNARMSHYQ SLLLTERITF
DADKLTLGQN ITVIAPHALE NIVRQPPDRW MTNARMTHYQ SLLLTERITF
DADKLTLGQN ITVIAPHALE NIVRQPPDRW MTNARMTHYQ SLLLTERITF
DADKLTLGQN ITVIAPHALE NIVRQPPDRW MTNARMTHYQ SLLLTERITF
DADKLTLGQN ITVIAPHALE NIVRQPPDRW MTNARMTHYQ SLLLTERITF
DADKLTLGQN ITVIAPHALE NIVRQPPDRW MTNARMTHYQ SLLLTERITF
DADKLTLGQN ITVIAPHALE NIVRQPPDRW MTNARMTHYQ SLLLTERVTF
DADKLTLGQN ITVIAPHALE NIIRQPPD-W MTNARMTHYQ SLLLTERVTF
DADKLTLGQN ITVIAPHTLE NIVRQPPDRW MTNAHMTHYQ SLLLTERVTF
DADKLTLGQN ITIIAPHALE NIVRQPPDRW MTNARMTHYQ SLLLTERITF
DADKLTLGQN ITIIAPHALE NIVRQPPDRW MTNACMTHYQ SLLLTERITF
DADKLTLGQN ITIIAPHALE NIVRQPPDRW MTNACMTHYQ SLLLTERITF
DADKLTLGQN ITVIAPHALE NIVRQPPDRW MTNVCMTRYQ SLLLTERVTF
DTDKLTLGQN ITIIAPHALE NIVRQPPDRW MTNARMTHYQ SLLLTERITF
DADKLTLEDN ---LPPHALE SIVRQPPDRW ITNAHMTHYQ SLLLTNRITF

DADKLTMGQN VTIVAPHSLE SIIRQPPDRW MTNARMTHYQ SLLLTERVSF
DADKLTMGQN VTIVAPHSLE SIIRQPPDRW MTNARMTHYQ SLLLTERVSF
DADKLTMGQN VTIVAPHSLE SIIRQPPDRW MTNARMTHYQ SLLLTERVSF
DADKLTMGQN VAIVAPHSLE SIIRQPPDHW MTNARMTHYQ SLLLTERVSF
DADKLTMGQQ VTVVAPHALE SIVRQPPDRW MTNARMTHYQ SLLLNERVTF
DADKLTMGQQ VTVVAPHALE SIVRQPPDRW MTNARMTHYQ SLLLNERVTF
DADKLTMGQQ VTVVAPHALE SIVRQPPDRW MTNARMTHYQ SLLLNERVTF
DADKLTMGQQ VTVVAPHALE SIVRQPPDRW MTNARMTHYQ SLLLNERVTF
DADKLTMGQQ VTVVAPHALE SIVRQPPDRW MTNARMTHYQ SLLLNERVTF
VADKLTMGQQ VTVVAPHALE SIVRQPPDRW MTNARMTHYQ SLLLNERVTF
DADKLTMGQQ VTVVAPHALE SIVRQPPDRW MTNARMTHYQ SLLLNDRVTF
DADKLTLGQH VTVIAPHALE SIVRQPPDRW MTNARMTHYQ SLLLNERVTF
DADKLTLGQN VLVIAPHNLE SIVRQPPDRW MTNARMTHYQ SLLLNERVSF
DADKLTLGQN VTVIASHSLE SIVRQPPDRW MTNARMTHYQ SLLLNERVSF
DADKLTLGQQ ITVVAPHSLE SIIRQPPDRW MTNARMTHYQ SLLLTERVVF
DADKLTLGQQ ITVVAPHSLE SIIRQPPDRW MTNARMTHYQ SLLLTERVVF
DADKLTLGQQ ITVVAPHSLE SIIRQPPDRW MTNARMTHYQ SLLLTERVVF
DADKLTLGQQ ITVVAPHSLE SIIRQPPDRW MTNARMTHYQ SLLLTERVVF
DADKLTLGQQ ITVVAPHSLE SIIRQPPDRW MTNARMTHYQ SLLLTERVVF
DADKLTLGQ- ITVVAPHSLE SIICQPPDRW MTNARMTDYQ SLLLTERIVF
DADKLTLGQK ITVIAPHAL- STVQQPPDQW MTNAQMAHYQ SLLLTDRVMF
DSDKLTLGQP VTIIAPQALE SFIRQPPDRW ITNSRLTHYQ SLLLDDRVLF
DADKLTLGQQ TIVVAPHALE SIVRQPPDRW MTNARMTHYQ SLLLTERVKF
DADKLTLGQQ MTIIAPHSLE SIV-QPPDRW MMNARMTHYQ SLLLTGRITF
DFDKLTLGQR VTVVAPHTLE SIIRQPPDRW LTNARMTHYQ SLLLTDRVTF
DADKLTLGQR APCIRPHALE SIIRQPPERW MSNARMTHYQ SLLLDDRIGF
DADRLTLGQK LTVIAPHTLE SIIRQPPERW MSNAQMTHYQ SLLLNDRIEF
DADKLTLGQK LTVIAPHTLE SIIRQPPERW MSNARMTHYQ SLLLNDRIEF
DADKLTMGQN VTVVAPHSLE SIIRQPPDRW MTNAQMTHYQ SLLLTDRVTF
DADKLTLGQK VTIVAHHTLE SIIRQPPE-W MTNAHMTHYQ ALLLTNHVTF
DAGKLTMGQP LVILAPHAVE ALVKQPPDRW LSNARMTHYQ AMLLDDRVQF
DAGKLTMGQP LVILAPHAVE ALVKQPPDRW LSNARMTHYQ AMLLDDRVQF
DAGKLTMGQP LVILAPHAVE ALVKQPPDRW LSNARMTHYQ AMLLDDRVQF
DAGKLTMGQP LVILAPHAVE ALVKQPPDRW LSNARMTHYQ AMLLDDRVQF
DAGKLTMGQP LVILAPHAVK ALVKQPPDRW LSNARMTHYQ AMLLDDRVQF
DAGKLTMGQP LVILAPHAVE ALVEQPPDRW LSNARMTHYQ AMLLDDRVQF
DAGKLTMGQP LVILAPHAVE ALVKQPPDRW LSNARMTHYQ AMLLDDRVQF
DAGKLTMGQP LVILAPHAVE ALVKQPPDRW LSNARMTHYQ ALLLDDRVQF
DAGKLTMGQP LVILAPHAVE ALVKQPPDRW LSNARMTHYQ ALLLDDRVQF
DAGKLTMGQP LVILAPHAVE ALVKQPPDRW LSNARMTHYQ ALLLDDRVQF
DAGKLTLGQP LTILAPHVVE ALIKQPPDRW LSNSRMTHYQ ALLLDERVQF
DAGKLTLGQP LTILAPHVVE ALIKQPPDRW LSNSRMTHYQ ALLLDERVQF
DAGKLTLGQP LTILTSHPVE ALVRQPPNKW LSNARMTHYQ AMLLDERVHF
DSAKLTLGQP LTVITPHALE AIVRQPPDRW ITNARLTHYQ ALLLDDRVQF

```
DSAKLTLGQP LTVITPHTLE AIVRQPPDRW ITNARLTHYQ ALLLDDRVQF
DADKLTLGQN LTITAPHALE NVIRQPPDRW LTNARMTHYQ TLLLNDRIKF
DADKLTLGQQ MTIIAPHSFE SIVGQSPDQW MSNARMTHYQ SLLLTDRINF
DADKLTLGQN -TIIAPHAL- NIVRQPPDRW MTNARMTHYQ SLLLTERVTF
DADKLTLGQQ IIIVAPHAPE SVIRQPPDRW MISTHMTHYQ SLLLTEHVVF
DADKLTLGQQ ITIVAPHALE SIIWQPPDRW MTNTR----- ----------
DADKLTFGQH LKVVTPHAIE GVLKYPPGRW MTNARLTHYQ GLLLDPRIIF
DADKLTLRQQ ITMVAPHPLE SIIRQPSD-- MANA----MT SLLLTEWVMF
EASKLTFGQD IEITSSHNLE SLLRSPPDKW LTNARITQYQ VLLLDPRVRF
DADKLTLGQQ ITVEAPHSLE STICQPPDLW IANAWMTHYQ SLLLTEQIMF
DADKLTLGQQ ITVEAPHSLE STICQPPDLW IANAWMTHYQ SLLLTEGVMF
DAKKLTLGQR TIVVVPHGLE SILNHPPDRW MTNAHITHYQ SLLLTERLKF
DADKLTLGQR MLVLDPHVLD NIIRPPPDCW MTNAHMTHYQ KLLLTEIVQF
DADKLTLGQR TIVVDPHVLE NIIRPPPDRW MTNANMTHYQ KLLLTERVQF

APPAALNPAT LLPEETDEPV THDCHQLLIE ETGVRKDLTD IPLTGEVLTW
APPAALNPAT LLPEETDEPV THDCHQLLIE ETGVRKDLTD IPLTGEVLTW
APPAALNPAT LLPEETDEPV THDCHQLLIE ETGVRKDLTD IPLTGEVLTW
APPAALNPAT LLPEETDEPV THDCHQLLIE ETGVRKDLTD IPLTGEVLTW
APPAALNPAT LLPEETDEPV THDCHQLLIE ETGVRKDLTD IPLTGEVLTW
APPAALNPAT LLPEETDEPV THDCHQLLIE ETGVRKDLTD IPLTGEVLTW
APPAALNPAT LLPEETDEPV THDCHQLLIE ETGVRKDLTD IPLTGEVLTW
APPAALNPAT LLPEETDEPV THDCHQLLIE ETGVRKDLTD IPLTGEVLTW
APPAALNPAT LLPEETDEPV THDCHQLLIE ETGVRKDLTD IPLTGEVLTW
APPAALNPAT LLPEETDEPV THDCHQLLIE ETGVRKDLTD IPLTGEVLTW
APPAALNPAT LLPEETDEPV THDCHQLLIE ETGVRKDLTD IPLTGEVLTW
APPAALNPAT LLPEETDEPV THDCHQLLIE ETGVRKDLTD IPLTGEMLTW
APPAALNPAT LLPEETDEPV THDCHQLLIE ETGVRKDLTD IPLTGEVLTW
APPAALNPAT LLPEETDEPV THDCHQLLIE ETGVRKDLTD IPLTGEVLTW
APPAALNPAT LLPEETDEPV THDCHQLLIE ETGVRKDLTD IPLTGEVLTW
APPAALNPAT LLPEETDEPV THDCHQLLIE ETGVRKDLTD IPLTGEMLTW
APPAALNPAT LLPEETDKPV THDCHQLLIE ETGVRKDLTD IPLTGEVLTW
APPAALNPAT LLPEETDEPV THDCHQLLIE ETGVRKDLTD IPLTGEVLTW
APPAALNPAT LLPEETDEPV THDCHQLLIE ETGVRKDLTD IPLTGEVLTW
APPAALNPAT LLPEETDEPV THDCHQLLIE ETGVRKDLTD IPLTGEVLTW
APPAALNPAT LLPEETDEPV THDCHQLLIE ETGVRKDLTD IPLTGEVLTW
APPAALNPAT LLPEETDEPV THDCHQLLIE ETGVRKDLTD IPLTGEVLTW
APPAALNPAT LLPEETDEPV THDCHQLLIE ETGVRKDLTD IPLTGEVLTW
APPAALNPAT LLPEETDEPV THDCHQLLIE ETGVRKDLTD IPLTGEVLTW
APPAALNPAT LLPEETDEPV THDCHQLLIE ETGVRKDLTD IPLTGEVLTW
APPAALNPAT LLPEETDEPV THDCHQLLIE ETGVRKDLTD IPLTGEVLTW
APPAALNPAT LLPEETDEPV THDCHQLLIE ETGVRKDLTD IPLTGEVLTW
APPAALNPAT LLPEETDEPV THDCHQLLIE ETGVRKDLTD IPLTGEVLTW
APPAALNPAT LLPEETDEPV THDCHQLLIE ETGVRKDLTD IPLTGEVLTW
```

APPAALNPAT LLPEETDEPV THDCHQLLIE ETGVRKDLTD IPLTGEVLTW
APPAALNPAT LLPEETDEPV THDCHQLLIE ETGVRKDLTD IPLTGEVLTW
APPAALNPAT LLPEETDEPV THDCHQLLIE ETGVRKDLTD IPLTGEVLTW
APPAALNPAT LLPEETDEPV THDCHQLLIE ETGVRKDLTD IPLTGEVLTW
APPAALNPAT LLPEETDEPV THDCHQLLIE ETGVRKDLTD IPLTGEVLTW
APPAALNPAT LLPEETDEPV THDCHQLLIE ETGVRKDLID IPLTGEVLTW
APPAALNPAT LLPEETDEPV THDCHQLLIE ETGVRKDLTD IPLTGEVLTW
APPAALNPAT LLPEETDEPV THDCHQLLIE ETGVRKDLTD IPLTGKVLTW
APPATLNPAT LLPEETDEPV THDCHQLLIE ETGVHKDLTD IPLTGEVLTW
APPAALNPAT LLPEEADEPV THDCHQLLIE ETGVRKDLTD IPLTGEVLTW
APPAALNPAT LLPEETDEPV THDCHQLLIE ETGVRKDLTD IPLTGEVLTW
APPAALNPAT LLPEETDESV THDCHQLLIE ETGVRKDLTD IPLTGEVLTW
APPAALNPAT LLPEETDEPV THDCHQLLIK ETGVRKDLTD IPLTGEVLTW
APPAALNPAT LLPEETDEPV THDCHQLLIE ETGVRKDLID IPLTGEVLTW
APPATLNPAT LLPEETDEPV THDCHQLLIE ETGVRKDLTD IPLTGEVLTW
TLPAALNPAT LLPEETDEPL THDCHQLLIE E--VRKDLTD IPLTGEVLTW
TLPAALNPAT LLPEETDEPL THDCHQLLIE E--VRKDLTD IPLTGEVLTW
TLPAALNPAT LLPEETDEPL THDCHQLLIE E--VRKDLTD IPLTGEVLTW
TLPAALNPAI LLPEETDEPL THDCHQLLIE E--VRKDLTD IPLTGEVLTW
APPAALNPAT LLPEETDEPV THDCHQLLIE ETGVRKDLTD IPLTGEVLTW
APPAALNPAT LLPEETDEPV THDCHQLLIE ETGVRKDLTD IPLTGEVLTW
APPATLNPAT LLPEETDEPV THDCHQLLIE ETGVRKDLTD IPLTGEVLTW
APPAALNPAT LLPEETDEPV THDCHQLLIE ETGVRKDLTD IPLTGEMLTW
APPAALNPAT LLPEETDEPV THDCQQLLIE ETGVHKDLTD IPLTGEVLTW
TPPAALNPAT LLPEETDEPV THDCQQLLIE ETGVHKDLTD IPLTGEVLTW
TPPAALNPAT LLPEETDEPV THDCQQLLIE ETGVHKDLTD IPLTGEVLTW
APRAALNPAT LLPEETDEPV THDCHQLLIE ETGVRKDLTD IPLTGEVLTW
APPAALNPAT LLPEETDEP- THDCHQLLIE EIGVHKDLPD IPLTGEVLTW
APPAILNPAT LLPEETDEPA THDCHQMLVE EIGI-KDLTD VSLTGGTLTW
APPAILNPAS LLPEA-DEAP AHKCEEILAE ETGIRPDLTD QPWPG-AMTW
APPAILNPAS LLPEA-DEAP AHKCEEILAE ETGIRPDLTD QPWPG-AMTW
APPAILNPAS LLPEA-DEAP VHKCEEILAE ETGIRPDLTD QPWPG-AMTW
APPAILNPAS LLPEA-DEAP AHKCEEILAE ETGIRPDLTD QPWPG-AMTW
APPAILNPAT LLPLTNDSAP VHQCMDILAE ETGTRSDLTD QPWPG-APSW
APPAILNPAT LLPLTNDSVP VHQCMDILAE ETGTRSDLTD QPWPG-APSW
APPAILNPAT LLPLTNDSVP VHQCMDILAE ETGTRSDLTD QPWPG-APSW
APPAILNPAT LLPLTNDSVP VHQCMDILAE ETGTRRDLTD QPWPG-APSW
APPAILNPAT LLPLTNDSVP VHQCMDILAE ETGARSDLTD QPWPG-APSW
APPAILNPAT LLPLTNDSVP VHQCTDILAE ETGTRRDLTD QPWPG-ALSW
APPAILNPAT LLPLTNDSVP VHRCADILAE EIGTRKDLTD QPWPG-APSW
APPAILNPAT LLPEIHNSTP IHQCVDILAE ETGTRKDLTD RPWPG-VPAW
APPAILNPAT LLPVESDDTP IHICSEILAE ETGTRPDLRD QPLPG-VPAW
APPAVLNPAT LLPVESEATP VHRCSEILAE ETGTRRDLED QPLPG-VPTW
APPAILNPAT FLPETDDSSP VHCCADILAE EIGIRSDLCD QPWPG-VPNW

```
APPAILNPAT FLPETDDSSP VHCCADILAE EIGIRSDLRD QPWPG-VPNW
APPAILNPAT FLPETDDSSP VHCCADILAE EIGIRSDLRD QPWPG-VPNW
APPAILNPAT FLPETDDSSP VHCCADILAE EIGIRSDLRD QPWPG-VPNW
APPAILNPAT FLPETDDSSP VHCCADILAE EIGIRSDLRD QPWPG-VPNW
APSVILNPAT PLPEMDDFSP IHCCADILVE ETGIRSDLRD QPWPS-IPSW
TPPAILNPAT LLPEENDEPV THDFHQLLVE ETGIRKDLTD VPLTRGTLTW
GAPVILNPAT LLPEATEQKP IHVCADILAE AFGVREDLTD VPLLG-CPSW
---------- ---ETDDSLP VHRCMDILAE ETRTKKDLTD QPWPD-CPNW
---------- ---ETEDSTP AHKCEEILAE ETGIRKDLRD QTWPG-GSTW
APLDILNPAT LLPEAEERAP IHDCQDILAE ETSTQKDLTD QLWPG-CPNW
GTPATLNPAT LLPETSVN-V THSCQEILAE ETGTWKDLKD QPLKESLLTW
GTPVTLNPAT LLPDISAE-V THSCQEILAE EAGTRRDLRD QPLKGSLLTW
VTPVTLNPAT LLPDISTE-V THSCQEILAE EAGTRQDLRD QPLKGNLLTW
ASPAVLNPDT LLPETNET-P VHHCGEILAE ETGTRPDLSD QPWPG-AATW
---------- ---EIGDSTV IHQCADILAE ETRIRRDLKD QPWVG-SPSW
GPVVALNPAT LLPLPGKE-P HHDCLEILAE THGTRPDLTD QPLPDADHTW
GPVVALNPAT LLPLPGKE-P HHDCLEILAE THGTRPDLTD QPLPDADHTW
GPVVALNPAT LLPLPGKE-P HHDCLEILAE THGTRPDLTD QPLPDADHTW
GPVVALNPAT LLPLPGKE-P HHDCLEILAE THGTRPDLTD QPLPDADHTW
GLVVALNPAT LLPLPGKE-T PHDCLEILAE THGTRPDLTD QPLPNADHTW
GPVVALNPAT LLPLPGKE-T PHDCLEILAE THGTRPDLTD QPLPNADHTW
GPVVALNPAT LLPLPEKG-A PHDCLEILAE THGTRPDLTD QPIPDADHTW
GPIVTLNPAT LLPLPEEG-L QHDCLDILAE AHGTRPDLTD QPLPDADHTW
GPIVALNPAT LLPLPEEG-L QHDCLDILAE AHGTRPDLTD QPLPDADHTW
GPVVALNPAT LLPLPEEG-L QHNCLDILAE AHGTRPDLTD QPLPDADHTW
GPIVALNPVT LFPLPEEA-E QHDCLQILAE VHGTRQDLSD QPLQNADHTW
GPIVALNPVT LFPLPEEA-E QHDCLQILAE VHGTRQDLSD QPLQNADHTW
GPTVSLNPAT LLPLPSGG-N HHDCLQILAE THGTRPDLTD QPLPDADLTW
GPPVTLNPAT LLPAPKDQQS AHDCRQVLAE THGTREDLKD QELPDADHSW
GPPVTLNPAT LLPVPENQPS PHDCRQVLAE THGTREDLKD QELPDADHTW
APATGLNPAT LLPDPDLEGS THDCQEVLAA AHGSRPDLTD LPLPDADFTW
APPAVLNHAT LPLDTEDSTP AHQCEYILAE ETGMRKDLRG QPWPE-GSTW
APPAASNPAT LLPEETDEPV THDCHQVIEE T--VRKDLID IPLTGEVLTW
ALPAILNPTT LLPEAEELCP IHHCADILAE EVGTCRNLRD HPWAG-SPNW
---------- ---ETDDFSP IHHCTDILAE ETGIRHDRKD QPWPG-SPSW
AEPTALNPAT LLPTPDLRAP LHDCQEIMAE VTQVRPDLQD TALPNSELVW
CPPANVKPSL WKKNWDPM-- ---CSTQL-- ---------D QPWPG-VPNW
KQTAALNPAT LLPETDDTLP IHHCLDTLDS LTSTRPDLTD QPLAQAEATL
APPAILNPAT FLPETDNSSP HHCCANILAE EIEIQSNLRD QPWPG-VPNW
APLAILNPAT FLPETDNSSP GHCCANILAE EIGIQSDLRD QPWPG-VP--
APPTILNPAI LLLESHDSLP VYQCMDILEE ETRTRKDFTD QPWPG-CPNW
LPPAILNSAT LL-SADKSLP EHPCIGILAE ETGTKEELSD HPWPD----W
---------- ---------- -HQCMGILAE ETGTKKELTD QPWPD-CPNW
```

FTDGSSYVVE GKRMAGPAVV AGTRPIWASS LPEGTSAQKA ELMALTQALR
FTDGSSYVVE GKRMAGPAVV AGTRPIWASS LPEGTSAQKA ELMALTQALR
FTDGSSYVVE GKRMAGPAVV AGTRPIWASS LPEGTSAQKA ELMALTQALR
FTDGSSYVVE GKRMAGAAVV DGTRTIWASS LPEGTSAQKA ELMALTQALR
FTDGSSYVVE GKRMAGAAVV DGTRTIWASS LPEGTSAQKA ELMALTQALR
FTDGSSYVVE GKRMAGAAVV DGTRTIWASS LPEGTSAQKA ELMALTQALR
FTDGSSYVVE GKRMAGAAVV DGTRTIWASS LPEGTSAQKA ELMALTQALR
FTDGSSYVVE GKRMAGAAVV DGTRTIWASS LPEGTSAQKA ELMALTQALR
FTDGSSYVVE GKRMAGAAVV DGTRTIWASS LPEGTSAQKA ELMALTQALR
FTDGSSYVVE GKRMAGAAVV DGTRTIWASS LPEGTSAQKA ELMALTQALR
FTDGSSYVVE GKRMAGAAVV DGTRTIWASS LPEGTSAQKA ELMALTQALR
FTDGSSYMVE GKRMAGAAVV DGTRTIWASS LPEGTSAQKA ELMALTQALR
FTDGSSYVVE GKRMAGAAVV DGTRTIWASS LPEGTSAQKA ELMALTQALR
FTDGSSYVVE GKRMAGAAVV DGTRTIWASS LPEGTSAQKA ELMALTQALR
FTDGSSYVVE GKRMAGAAVV DGTRTIWASS LPEGTSAQKA ELMALTQALR
FTDGSSYVVE GKRMAGAAVV DGTRTIWASS LPEGTSAQKA ELMALTQALR
FTDGSSYVVE GKRMAGAAMV DGTRTIWASS LPEGTSAQKA ELMALTQALR
FTDGSSYVVE GKRMAGAAVV DGTRTIWASS LPEGTSAQKA ELMALTQALR
FTDGSSYVVE GKRMAGAAVV DGTRTIWASS LPEGTSAQKA ELMALTQALR
FTDGNNYVVE DKRMAGAVVV DGTRKIWASS LPEGTSAQKA ELMALTQALR
FTDGNNYVVE DKRMAGAVVV DGTRKIWASS LPEGTSAQKA ELMALTQALR
FTDGSSYVVE GKRMAGAAVV DGTRMILASS LPEGTSAQKA ELVALTQ---
FTDGSSYVVE GKRMAGAAVV DGTRTIWASS LPEGTSAQKA ELMALTQALR
FTDGSSYVVE GKRMAGAAVV DGTRTIWASS LPEGTSAQKA ELMALTQALR
FTDGSSYVVE GKRMAGAAVV DGTRTIWASS LPEGTSAQKA ELMALTQALR
FTDGSSYVVE GKRMAGAAVV DGTRTIWASS LPEGTSAQKA ELMALTQALR
FTDGSSYVVE GKRMAGAAVV DGTRTIWASS LPEGTSAQKA ELMALTQALR
FTDGSSYVVE GKRMAGAAVV DGTHTIWASS LPEGTSAQKA ELMALTQALR
FTDGSSYVVE GKRMAGAAVV DGTRTIWASS LPEGTSAQKA ELMALTQALR
FTDGSSYVVE GKRMAGTAVV DGTHTIWASS LPEGTSAQKA ELMALTQALR
FTDGSSYVVE GKRMAGAAVV DGTRTIWASS LPEGTSAQKA ELMALTQALR
FTDGSSYVVE GKRMAGAAVV DGTRTIWASS LPEGTSAQKA ELMALTQALR
FTDGSSYVVE GKRMAGAAVV DGTRTIWASS LLEGTSAQKA ELMALTQALR
FTDGSSYVVE GKRMAGAAVV DGTRTIWASS LPEGTSAQKA -LMALTQALR
FTDGSSYVVE GKRMAGAAVV DGTRTIWASS LPEGTSAQKA ELMALTQALR
FTDGSSYVV- GKRMAGAAVV DGTRTI-ASS LPEGTSAQKA ELMALTQALR
FTDGNSYVVE GKRMAGAAVV DGTRTIWASS LPEGTSAQKA ELMALTQALR
FTDGSSYVVE CKRMAGAAVV DGTRTIWASS LPEGTSAQKA ELMALTQALR
FTDRSSYVVE GKRMAGAAVV DGTHTIWASS LPEGTSAQKA ELMALTQALR
FTDGSSYVVE GKRMAGAAVV DGTHTIWASS LPEGTSAQKA ELMALTQALW
FTDGSSYVVE GKRMAGVAVV DGTRTIWASS LPEGTSAQKA ELMALTQALG
FTDGSSYVVE GKRMAGAAVV DGTRTIWASS LPEGTSVQKA ELMALTQALR
FTDGSSYVVE GKRMAGAAVV DGTRTIWASS LPEGTSAQKA ELMALTQALR
FTDGSSYVVE GKRMAGAAVV DGTRTIWATS LPEGTSVQKA ELMALTQALR

```
FTDGSSYVVE GKRVAGAAVV DGTRTIWASS LPEGTSAQKA ELVALTQALR
FTDGSSYVVE GKRVAGAAVV DGTRTIWASS LPEGTSAQKA ELVALTQALR
FTDGSSYVVE GKRVAGAAVV DGTRTIWASS LPEGTSAQKA ELVALTQALR
FTDGSSYVVE GKRVAGAAVV DGTRMIWASS LPEGTSAQKA ELVALTQALR
FTDGSSYVVE GKRMARAAVV DGTRTIWASS LSEGTSAQKA ELVALTQALR
FTDGSSYVVE GKRMAGAAVV DGTRTIWASS LPEGTSAQKA ELMALTQALR
FTDGNNYVVE DKRMAGAVVV DGTRKIWASS LPEGTSAQKA ELMALTQALR
FTDGSSYVVE GKRMAGAAMV DGTRTIWASS LPEGTSAQKT ELMALTQALR
FTDGSSYVVE GKRMAGGAVV GGTRTIWARS LPEGISAQKA ELVALMQALR
FTDGSSYVVE GKRMAGGAVV GGTRTIWARS LPEGISAQKA ELVALMQALR
FTDGSSYVVE GKRMAGGAVV GGTRTIWARS LPEGISAQKA ELVALMQALR
FTDGSSYVVE GKRMAGAAVV DGTHTIWASS LLEGTSAQKA ELMALTQALQ
FTDGSSYVVE GKRMAGAAVV DRTRMIWASS LPEGTSAQKA ELVALMQALR
FTDGSSYIVE GKRMAGVAIV DGM-TVWASS LPEGTLAQKA ELMALTQALQ
FTDGSSFVVE GKRKAGAAVV DGKAVIWASS LPEGTSAQKA ELIALIQALR
FTVGSSFVVE GKRKAGAAVV DGKAVIWASS LPEGTSAQKA ELIALIQALR
FTDGSSFVVK GKRKAGAAVV DGKAVIWASS LPEGTSAQKA ELIALIQALR
FTDGSSFVVE GKRRAGAAVV DKKAVIWASS LPEGTSAQKA ELIALIQALR
YTDGSSFLIE GKRKAGAAVV DGKKVIWASA LPEGTSAQKA ELIALIQALR
YTDGSSFLIE GKRKAGAAVV DGKKVIWASA LPEGTSAQKA ELIALIQALR
YTDGSSFLIK GKRKAGAAVV DGKKVIWASA LPEGTSAQKA ELIALIQALR
YTDGSSFLIE GKRKAGAAVV DGKKVIWASA LPEGTSAQKA ELIALIQALR
YTDGSSFLIE GKRKAGAAVV DGKKVIWASA LPEGTSAQKA ELIALIQALR
YTDGSSFLIE GKRKAGAAVV DRKKVIWASA LPEGTSAQKA ELIALIQAPR
YTDGSSFLIE GKRRAGAAVV DGKKVIWASA LPEGTSAQKA ELIALTQALR
YTDGSSFVVE GKRRAGAAVV DGKQVIWASS LPEGTSAQKA ELVALTQALR
YTDGSSFIMD GRRQAGAAIV DNTRTVRASN LPEGTSAQKA ELIALTQALR
YTDGSSFITE GKRRAGAPIV DGKRTVWASS LPEGTSAQKA ELVALTQALR
YTDGSSFLVE GKRMAGAAVV DGKRVVWAGS LPEGTSAQKA ELIALTQALW
YTDGSSFLVE GKRMAGAAVV DGKRVVWAGS LPEGTSAQKA ELIALTQALR
YTDGSSFLVE GKRMAGAAVV DGKRVVWAGS LPEGTSAQKA ELIALTQALR
YTDGSSFLVE GKRMAGAAVV DGKRVVWAGS LPEGTSAQKA ELIALTQALR
YTDGSSFLVE GKRMAGAAVV DGKRVVWAGS LPEGTSAQKA ELIALTQALR
YTDRSSFLVE GKRMAGVAVV DGQRVVWASS LPEGTSAQKA ELIAMTQALQ
FTDGSSYMVE GKRMAGAAIV DG--TVWASS LPEGTAAQKA ELVALTQAL-
YTDGSSFLVE GKRRAGAAVV DGKKTIWFSG LPEGTSAQKA ELIALTQALR
YMDGSSFLVE GKRKVGAAVV NGKQTIWASS LPEGTTAQRA ELIALTQALR
YMDGSSFVVE GKQVAGAAVV DRTDVIWASR LPEGTSAQKA ELIALIQALQ
FMDGSSFVVE GKRMAGAAVV DGKRTIWASN LPEGTSAQRA ELIALTQALR
YSDGSSYMTE GKRVAGAAVV DDSTTIWASS LPGGISAQWA ELIALTQALK
YTDGSSCIAE GKQVAGAAVV DDSTIILASR LPEGTSAQWA ELIALTQALK
YTDGRSYIVE GKRVAGAAVV DDSTIIWASR LPEGTSAQRA ELIALTQALK
FTDRSSFVVE GKR-AGAAVV DGKSVI-SSS LPEGTSAQKA ELIALTQALR
YTDGSSFVVE GKR------- ---------- --------KA ELIVLIQALC
```

YTDGSSFLQE GQRRAGAAVT TETEVIWAKA LPAGTSAQRA ELIALTQALK
YTDGSSFLQE GQRRAGAAVT TETEVIWAKA LPAGTSAQRA ELIALTQALK
YTDGSSFLQE GQRRAGAAVT TETEVIWAKA LPAGTSAQRA ELIALTQALK
YTDRSSFLQE GQRRAGAAVT TETEVIWAKA LPAGTSAQPA ELIALTQALK
YTDGSSFLQE GQRKAGAAVT TETEVIWAKA LPAGTSAQRA ELIALTQALK
YTDGSSFLQE GQRKAGAAVT TETEVIWAKA LPAGTSAQRA ELIALTQALK
YTDGSSFLQE GQRKAGAAVT TETEVIWARA LPAGTSAQRA ELIALTQALK
YTDGSSFLQE GQRKAGAAVT TETEVIWAKA LPAGTSAQRA ELIALTQALK
YTDGSSFLQE GQRKAGAAVT TETEVVWAKA LPAGTSAQRA ELIALTQALK
YTDGSSLLQE GQRKAGAAVT TETEVIWAKA LPAGTSAQRA ELIALTQALK
YTDGSSFLAD GERKAGAAVT TEDKVIWART LAAGTSAQRA ELIALTQALK
YTDGSSFLAD GERKAGAAVT TEDKVIWART LAAGTSAQRA ELIALTQALK
YTDGSSFIRN GEREAGAAVT TESEVIWAAP LPPGTSAQRA ELIALTQALK
YTDGSSYIDS GTRRAGAAVV DGHHIIWAQS LPPGTSAQKA ELIALTKALE
YTDGSSYLDS GTRRAGAAVV DGHNTIWAQS LPPGTSAQKA ELIALTKALE
FTDGSSFLEG GKRRAGAAVV DGKQVIWAAA LPQGTSAQRA ELIAMTRALE
CMDGSSFMVE GKRMAGAAVV DGTDVIWGSH LSEGTSAQKA ELIALI-ALQ
FTDGSSYVVE GKRMAGAA-V DGTRTIWASS LPEGTSAQKA ELMALTQALR
YTDGSSFVVE GKQKAGAAVV DEKWVIWASS LPEGTSAQKA ELIVFIQALR
YTDGSSFVVE GKWKAGAVVV DRNQVIRASS LPEGTSAQKA ELVALIQSLR
YTDGSSFVID GVRRAGAAVV DGGNIIWSAS LSPGTSAQKA ELIALAEALE
HTDGSSFLVE GKRMAGVVVV DDK-IVWASS LPEGT-VQKA ELIALTQALW
FTDGSSYVRD GKRYAGAAVV TLDSVIWAEP LPIGTSAQKA ELIALTKALE
YTDRSSFLLE GKIMAGAAVV DGKRDVWAGS LQEGKSAQKA ELIALTQAL-
FQDRSSFLV- SKRMAGAALV DGKRDVWAGS LQEGKSAQKA ELIALTQAL-
DTDGNSFLVR GKRKVGKVVV NGKQTILASS LPEGKTAQRA ELIILNQACS
CTDDSRFLVK GKQKAEAALG NGKQTICIST LPERTITQRA ELIALTQTLQ
CTDDSRFLFK GKQKAGAALG NGKQTICTST LPEGTITQRA ELIALTQALQ

LAEGKSINIY TDSRYAFATA HVHGAIYKQR GLLTSAGREI KNKEEILSLL
LAEGKSINIY TDSRYAFATA HVHGAIYKQR GLLTSAGREI KNKEEILSLL
LAEGKSINIY TDSRYAFATA HVHGAIYKQR GLLTSAGREI KNKEEILSLL
LAEGKSINIY TDSRYAFATA HVHGAIYKQR GLLTSAGREI KNKEEILSLL
LAEGKSINIY TDSRYAFATA HVHGAIYKQR GLLTSAGREI KNKEEILSLL
LAEGKSINIY TDSRYAFATA HVHGAIYKQR GLLTSAGREI KNKEEILSLL
LAEGKSINIY TDSRYAFATA HVHGAIYKQR GLLTSAGREI KNKEEILSLL
LAEGKSINIY TDSRYAFATA HVHGAIYKQR GLLTSAGREI KNKEEILSLL
LAEGKSINIY TDSRYAFATA HVHGAIYKQR GLLTSAGREI KNKEEILSLL
LAEGKSINIY TDSRYAFATA HVHGAIYKQR GLLTSAGREI KNKEEILSLL
LAEGKSINIY TDSRYAFATA HVHGAIYKQR GLLTSAGREI KNKEEILSLL
LAEGKSINIY TDSRYAFATA HVHGAIYKQR GLLTSAGREI KNKEEILSLL
LAEGKSINIY TDSRYAFATA HVHGAIYKQR GLLTSAGREI KNKEEILSLL
LAEGKSINIY TDSRYAFATA HVHGAIYKQR GLLTSAGREI KNKEEILSLL
LAEGKSINIY TDSRYAFATA HVHGAIYKQR GLLTSAGREI KNKEEILSLL

LAEGKSINIY TDSRYAFATA HVHGAIYKQR GLLTSAGREI KNKEEILSLL
LAEGKSINIY TDNRYAFATA HVHGAIYKQR GLLTSAGREI KNKEEILSLL
LAEGKSINIY TDSRYAFATA HVHGAIYKQR GLLTSAGREI KNKEEILSLL
LAEGKSINIY TDSRYAFATA HVHGAIYKQR GLLTSAGREI KNKEEILSLL
LAKGKSINIY TDSRYAFATA HVHGAIYKQR GLLTSAGREI KNKEEILSLL
LAKGKSINIY TDSRYAFATA HVHGAIYKQR GLLTSAGREI KNKEEILSLL
LAEGKSINIY TDSRYAFATV HVHGAIYKQR GLLTSAGREI KNKEEILSLL
LAEGKSINIY TDSRYAFATA HVHGAIYKQR GLLTSAGREI KNKEEILSLL
LAEGKSINIY TDSRYAFATA HVHGAIYKQR GLLTSAGREI KNKEEILSLL
LAEGKSINIY TDSRYAFATA HVHGAIYKQR GLLTSAGREI KNKEEILSLL
LAEGKSINIY TDSRYAFATA HVHGAIYKQR GLLTSAGREI KNKEEILSLL
LAEGKSINIY TDSRYAFATA HVHGAIYKQR GLLTSAGREI KNKEEILSLL
LAEGKSINIY TDSRYAFATA HVHGAIYKQR GLLTSAGREI KNKEEILSLL
LAEGKSINIY TDSRYAFATA HVHGAIYKQR GLLTSAGREI KNKEEILSLL
LAEGKSINIY TDSRYAFATA HVHGAIYKQR GLLTSAGREI KNKEEILSLL
LAEGKSINIY TDSRYAFATA HVHGAIYKQR GLLTSAGREI KNKEEILSLL
LAEGKSINIY TDSRYAFATA HVHGAIYKQR GLLTSAGREI KNKEEILSLL
LAEGKSINIY TDSRYAFATA HVHGAIYKQR GLLTSAGREI KNKEEILSLL
LAEGKSINIY TDSRYAFATA HVHGAIYKQR GLLTSAGREI KNKEEILSLL
LAEGKSINIS TDSRYAFATA HVHGAIYKQR GLLTSAGREI KNKEEILSLL
LAEGKSINIY TDSRYAFATA HVHGAIYKQR GLLTSAGREI KNKDEILSLL
LAEGKSINIY TDSRYAFATA HVHGAIYKQR GLLTSAGREI KNKEKILSLL
LAEGKSINIY TDSRYAFATA HVHGAIYKQR GLFTSAGREI KNKEEILSLL
LAEGKSINIY TDSRYAFATA HVHRAIYKQR GLLTSAGREI KNKEEILSLL
LAEGKSINIY TDSRYAFATA HVHGAIYKQR VLLTSAGREI KNKEEILSLL
LAEGKSINIY TDSRYAFATV HIHGAIYKQR GLLTSAGREI KNKEEILSLL
LAEGKSINIY TDSRYAFATA HVHGAIYKQR GLLTSAGREI KNKEEILSLL
LAEGKSINIY TDSRYAFATA HVHGAIYKQR GLLTSAGREI KNKEEILSLL
LAEGKSINIY TDSRYAFATA HVHGAIYKQK GLLTSAGREI KNKEEILSLL
LAEGKSINIY TYSRYALATA HVHGAIYKQR GLLTSAGREF KNKEEILSLL
LAEGKSINIY TYSRYALATA HVHGAIYKQR GLLTSAGREF KNKEEILSLL
LAEGKSINIY TYSRYAFATA HVHGAIYKQR GLLTSAGREV KNKEEILSLL
LAEGKSINIY TYSRYAFATA HVHGAIYKQR GLLTSAGREV KNKEEILSLL
LAEGKSINIY TDSRYAFATA HVHGAIYKQR GLLTSAGREV KNKEKILSLL
LAEGKSINIY TDSRYAFATA HIHGAIYKQR GLLTSAGREI KNKEEILSLL
LAKGKSINIY TDSRYAFATA HVHGAIYKQR GLLTSAGREI KNKEEILSLL
LAEGKSINIY TDSRYAFATA HVHGAIYKQR GLLTSTGRKI KNKEEILSLL
LAEGKSINIY TDSRYAFVTA QVHGAIYKQR GLLTSTGREV KNKEEILSLL
LAEGKSINIY TDSRYAFVTA QVHGAIYKQR GLLTSTGREV KNKEEILSLL
LAEGKSINIY TDSRYAFVTA QVHGAIYKQR GLLTSTGREV KNKEEILSLL
LAEGKSIDIY TDSRYAFATA HVHGAIYKQR GLLTSAGREI KNKEEILSLL
LAKGKSINIY TDSRYAFATA HVHGAIYKQR GLLTSAGREV KNKEEILSLL
LAEGKSVNIY TDSR-AFATA HVHGAIYKQR GLLTSAGKEI KNKEEFLSLL
LAEGRALNVY TDSRYAFATA HVHGAIYRHR GLLTSAGKDI KNKEEILSLL

```
LAEGRALNVY TDSRYAFATA HVHGAIYRHR GLLTSAGKNI KNKEEILSLL
LAEGRALNVY TDS--AFATA HVHGAIYRHR GLLTSAGKDI KNKEEILSLL
LAEERALNVY TDSQYAFATA HVHGAIYRHH GLLTSAGKDI KNKEEILSLL
EAKGKIVNIY TDSRYAFATA HIHGAIYRQR GLLTSAGKDI KNKEEILALL
EAKGKIVNIY TDSRYAFATA HIHGAIYRQR GLLTSAGKDI KNKEEILALL
EAKGKIINIY TDSRYAFATA HIHGAIYRQR GLLTSAGKDI KNKEKILALL
EAKGKIVNIY TDSRYAFATA HIHGAIYRQR GLLTSAGKDI KNKEEILALL
EAKGKIVNIY TDSRYAFATA HIHGAIYRQR GLLTSAGKDI KNKEEILALL
EAKGKIVNIY TDSRYAFATA HIHAAIYRQR GLLTSAGKDI KNKEEILALL
EAEGKIINIY TDSRYAFATA HIHGAIYRQR GLLTSAGKDI KNKEEILALL
LAEGKAINIY TDSRYAFATA HIHGAIYKQR GLLTSAGRDI KNKEEILALL
LAEGKSINIY TDSRYAFATA HVHGAIYKQR GLLTSAGKDI KNKEEILALL
LAEGKNINIY TDSRYAFATA HIHGAIYKQR GLLTSAGKDI KNKEEILALL
MAEGQSINIY TDSRYAFATA HVHGAIYRQR GLLTSAGKDI KNKEEILSLL
MAEGQSINIY TDSRYAFATA HVHGAIYRQR GLLTSAGKDI KNKEEILSLL
MAEGQSINIY TDSRYAFATA HVHGAIYRQR GLLTSAGKDI KNKEEILSLL
MAEGQSINIY TDSRYAFATA HVHGAIYRQR GLLTSAGKDI KNKEEILSLL
MAEGQSINIY TDSRYAFATA HVHGAIYRQR GLLTSAGKDI KNKEEILSLL
MAEGRPINIY TDSRYAFATA HIHGAFYRQR GLL--SGKDI KNKEEILSLL
LAEGKSVNIY MDSRYAFATA HVHGAIYQQR GLLTSAGKEV KNKEEILCLL
LAEGKNINIY TDSRYAFATA HVHGAIYQQR GLLTSTGKEV KNKDEILSLL
LAKEKNINIY TDSRYAFATA HIRGTIYRQR GLLTSAGKDV KSKEEILSLL
MAEGKTVNIY TDSRYAFATV HVHGAIYRQR VLLTSAGKDI KNKEEILSLL
LASGNCVNIY MDSSYAFATA HIHGAIYKQR ELLTSGGEEI KNKEEILSLL
LAEGKRLNLY TDSCYAFATA HVHGAIYQWR GLLTSAGKDI KNRQEILELL
LAEKKRLNLY TDSRYAFATA HVHGAIYRQR GLLTSAGKDI KNKQEILELL
LAEKKRLNVY TDSRYAFATA HVHRAIYRQR GLLTSAGKDI KSKQDIIGLL
LAEGKAINIY TDC-YAFATA HVHGAIYRQR GLLTSAGKDI KNKEEILGLL
LAEGKSISIY TDSRYTFATA HVHGAIYQQR GLLTSAGKEI KNKEEILSLL
MAEGKKLNVY TDSRYAFATA HVHGEIYRRR GLLTSEGKEI KNKGEILALL
MAEGKKLNVY TDSRYAFATA HVHGEIYRRR GLLTSEGKEI KNKGEILALL
MAEGKKLNVY TDSRYAFATA HVHEKIYRRR GLLTSEGKEI KNKGEILALL
MAEGKKLNVY TDSRYAFATA HVHGKIYRRR GLLTSEGKEI KNKGEILALL
MAEGKKLNVY TDSRYAFATA HVHGEIYRRR GLLTSEGKEI KNKGEILALL
MAEGKKLNVY TDSRYAFATA HVHGEIYRRR GLLTSEGKEI KNKGEILALL
MAEGKKLNVY TDSRYAFATA HVHGEIYRRR GLLTSEGKEI KNKSEILALL
MAEGKKLNVY TDSRYAFATA HIHGEIYRRR GLLTSEGKEI KNKDEILALL
MAEGKKLNVY TDSRYAFATA HIHGEIYRRR GLLTSEGKEI KNKDEILALL
MAEGKKLNVY TDSRYAFATA HIHGEIYRRR GLLTSEGKEI KNKDEILALL
MAKGKRLNVY TDSRYAFATA HIHGEIYRRQ GLLTSEGKDI KNKAEMLALL
MAKGKRLNVY TDSRYAFATA HIHGEIYRRQ GLLTSEGKDI KNKAEMLALL
MAEGKKLTVY TDSRYAFATT HVHGEIYRRR GLLTSEGKEI KNKNEILALL
LSEGKKANIY TDSRYAFATA HTHGSIYERR GLLTSEGKEI KNKAEIIALL
LSKGKKANIY TDSRYAFATA HTHGSIYERR GLLTSEGKEI KNKAEIIALL
```

```
MAENKKVNIY TDSRYAFATA HIHGTIYQQR GLLTSSGREI KNKDEIVALL
MAEGKNVNIY TDSRHSFATA HVHGAI-RQR ELLTSAGKDI KNKEEILSLL
LARRESINIY TAGRYAFATA HVHGAIYKQR GLLTSAGREI KNKEEILSLL
LAQGKNINIY MDSRYAFATA HIHGAIYRQQ GLLTSAGKEI KNKEEILSLL
MAEGKPINIY TDSRYAFATA HIHGAIYSQR GLLTSAGKDI --KEEILSLL
RAEGRRVTVY TDSRYAFGTV HVHGAIYRER GFVTAEGKAL RNLPEVRRLL
MTEGRPINIY TNSRYAFATA HIHGAIYRQR GLLTSVGKDI KNK-EILNFL
WSKDKSVNIY TDSRYAFATL HVHGMIYRER GLLTAGGKAI KNAPEILALL
IAEGQSINIY TDSRYVFATA HGQVAMYRQR GLLMSAGKDI KNKEEILSLL
IAQGQSINIY TDSRYVFATA HVHVAMYRQR GLLTSAGKDI KNKEEILSLL
LKRRTSINIY TDRRYAFATA HIHWAIYRQR GLLTSAGEDI KNKEEILSLL
LAKEKNNNIY TDSR-VFTTT YTCGTIYRQL GLLTSARKDI NNNKKILSRL
LAKEKNNNIY TDSR-VFTTT HICGTIYSQR GLLTSAGKDL ----------

EAVHLPKRLA IIHCPGH
EAVHLPKRLA IIHCPGH
EAVHLPKRLA IIHCPGH
EALHLPKRLA IIHCPGH
EAVHLPKRLA IIHCPGH
EALHLPKRLA IIHCPGH
EALHLPKRLA IIHCPGH
EALHLPKRLA IIHCPGH
EALHLPKRLA IIHCPGH
EALHLPKRLA IIHCPGH
EALHLPKRLA IIHCPGH
EALHLPKRLA IIHCPGH
EALHLPKRLA IIHCPGH
EALHLPKRLA IIHCPGH
EALHLPKRLA IIHCPGH
EALHLPKRLA IIHCPGH
EALHLPKRLA IIHCPGH
EALHLPKRLA IIHCPGH
EAVHLPKRLA IIHCPGH
EALHLPKRLA IIHCPGH
EALHLPKRLA IIHCPGH
EAVHLPKRLA IIHCPGH
EALHLPKRLA IIHCPGH
EALHLPKRLA IIHCPGH
EALHLPKRLA IIHCPGH
EALHLPKRLA IIHCPGH
EALHLPKRLA IIHCPGH
EALHLPKRLA IIHCPGH
EALHLPKRLA IIHCPGH
EALHLPKRLA IIHCPGH
```

EALHLPKRLA IIHCPGH

EALHLPKRLA IIHCPGH

EALHLPKRLA IIHCPGH

EALHLPKRLA IIHCPGH

EALHLPKRLA IIHCPGH

EALHLPKRLA IIHCPGH

KALHLPKRLA IIHCPGH

EALHLPERLA IIHCPGH

EALHLPKRLA IIHCPGH

EALHLPKRLA IIHCPGH

EALHLPKRLA IIHCPGH

EALHLPKRLA IIHCPGH

EALHLPKRLA IIHCPGH

EALHLPKRLA IIHCPGH

EALHLPKRPA IIHCLGH

EALHLPKRLA IIHCLGH

EALHLPKRLA IIHCLGH

EALHLPKRLA IIHCLGH

EALHLPKRLA IIHCPGH

EALHLPKRLA IIHCPGH

EALHLPKRLA IIHCPGH

EALHLPKRLA IIHCPGH

EALHLPKRLA IIHCLGH

EALHLPKRLA IIHCLGH

EALHLPKRLA IIHCLGH

EALHLPKRLA IIHCPGH

EALHLPKRLA IIHCLGH

EALYLPKRLA IMHCPGH

EAVHLPHRVA IIHCPGH

EAVHLPRRVA IIHCPGH

EAVHLPRRVA IIHCPGH

EAVHLPRRVA IIHCPGH

EAIHAPKKVA IIHCPGH

EAIHAPKKVA IIHCPGH

EAIHAPKKVA IIHCPGH

EAIHAPKKVA IIHCPGH

EAIHAPKKVA IIHCPGH

EAIHAPKKVA IIHCPGH

EAIHAPKKVA IIHCPGH

EAVHLPKKVA IIHCPGH

EAIHLPKRVA IIHCPGH

EAIHLPRRVA IIHCPGH

EAIHLPAKVA IIHCPGH

EAIHLPAKVA IIQCPGH

```
EAIHLPAKVA IIHCPGH
EAIHLPAKVA IIHCPGH
EAIHLPAKVA IIHCPGH
EAIHLPTKVA IMHCPGH
EALHLPKRLA IMHCPGH
EAVHLPRKVA IIHCPGH
EAIHLPKKLA IIHCPGH
EAIHLPKKVA IIHCPGH
KAIHLPKKVA IIHCPGH
TALHLPAELA IIHCPGH
TAIHLPAELA IIHCPGH
TAIHLPAELA IIHCPGH
EAVHLPLKVA IIHCPGH
EAIHLPKRVA IMHCPGH
KALFLPKRLS IIHCPGH
KALFLPKRLS IIHCPGH
KALFLPKRLS IIHCPGH
KALFLPKRLS IIHCPGH
KALFLPKRLS IIHCPGH
KALFLPKRLS IIHCPGH
KALFLPKRLS IIHCPGH
KALFLPKRLS IIHCPGH
KALFLPKRLS IIHCPGH
KALFLPKRLS IIHCPGH
EALFLPKRLS IMHCPGH
EALFLPKRLS IMHCPGH
EALFLPKRLS IIHCPGH
KALFLPRRVA IIHCPGH
KALFLPQEVA IIHCPGH
VALMLPTKVS IIHCPGH
EAIYLSKKVA IIHCIGH
EALHLPKRLV RPHSPLH
EAIHLPAKVA IIHFPGH
EAIHLP-NVA IIHCPGH
MAVQMPRAVA VVHIPGH
EAIHLPAKVA IIHCPGH
TAVWLPKRVA VMHCKGH
EGIHLPAKVA IIHFPGH
EAIHLPAKVA IIHFPGH
EAIHLRKNLV IIHYPCH
EAL------A TIHSPGH
---------A TIYCPGH
```

Env

148 334

| | | | | | |
|---|---|---|---|---|---|
| AEMK02000133.1 | VNGMSWGIVY | YGGSGRKGSI | LTIRLRIETE | PPVAIGPNKC | LAEQGPPIQE |
| CM000812.5_c | VNGMSWGIVY | YGGSGKKGSV | LTIRLRIETE | PPVAIGPNKG | LAEQGPPIQE |
| LT634572.1_d | VNGMSWGIVY | YGGSGKKGSV | LTIRLRIETE | PPVAIGPNKG | LAEQGPPIQE |
| CM000818.5_b | VNGMSWGIVY | YGGSGKKGSV | LTIRLRIETE | PPVAIGPNKG | LAEQGPPIQE |
| CM000812.5_b | VNGMSWGIVY | YGGSGKKGSV | LTIRLRIETE | PPVAIGPNKG | LAEQGPPIQE |
| CM000817.5 | VNGMSWGIVY | YGGSGKKGSV | LTIRLRIKTE | PPVAIGPNKG | LAEQGPPIQE |
| LUXX01006697.1 | VNGMSWGIVY | YGGSGKKGSV | LTIRLRIETE | PPVAIGPNKG | LAEQGPPIQE |
| LUXS01031219.1 | VNGMSWGIVY | YGGSGKKGSV | LTIRLRIETE | PPVAIGPNKG | LAEQGPPIQE |
| LUXX01090830.1 | VNGMSWGIVY | YGGSGKKGSV | LTIRLRIETE | PPVAIGPNKG | LAEQGPPIQE |
| LUXU01038095.1 | VNGMSWGIVY | YGGSGKKGSV | LTIRLRIETE | PPVAIGPNKG | LAEQGPPIQE |
| PERV-A | VNGISWGIVY | YGGSGKKGSV | LTIRLRIETE | PPVAIGPNKG | LAEQGPPIQE |
| CM000819.5_b | VNGMSWGIVY | YGGSGKKGSV | LTIRLRIETE | PPVAIGPNKG | LAEQGPPIQE |
| CM000823.5_b | VNGMSWGIVY | YGGSGKKGSV | LTIRLRIETE | PPVAIGPNKG | LAEQGPPIQE |
| CM000830.5_c | VNGMSWGIVY | YGGSGKKGSV | LTIRLRIETE | PLVAIGPNKG | LAEQGPPIQE |
| CM000824.5_b | VNGMSWGIVY | YGGSGKKGSV | LTIRLRIETE | PPIAIGPNKG | LAEQGPPIQE |
| CM000816.5_b | VNGMSWGIVY | YGGSGKKGSV | LTIRLRIETE | PPVAIGPNKG | LAEQGPPIQE |
| CM000826.5_a | VNGMSWGIVY | YGGSGKKGSV | LTIRLRIETE | PPVAIGPNKG | LAEQGPPIQE |
| LUXV01020898.1 | VNGMSWGIMY | YGGTGRKGSV | LTIRLRIETE | PPVAIGPNKG | LAEQGPPIQE |
| CM000828.5_a | VNGMSWGIMY | YGGTGRKGSV | LTIRLRIETE | PPVAIGPNKG | LAEQGPPIQE |
| CM000814.5_a | VNGMSWGIVY | YGGSGKKGSV | LTIRLRIETE | PPVAIRPNKG | LAEQGPPIQE |
| CM000828.5_c | VNSMSWGIVY | YGGSGKKRSV | LTIRLRIKTE | PPVTIGPNKG | LAEQGPPIQE |
| LT634572.1_a | VNGMSWGIMY | YGGSGRKGFV | LTIRLRIETE | PPVAIGPNKG | LAEQGPLIQE |
| LT634572.1_b | VNGMSWGIMY | YGGSGRKGSV | LTIRLRIETE | PPVAIGPNKG | LAEQGPPIQE |
| AJKK01232762.1 | VNGMS-KIMY | YGGSGRKGSI | LTIRLRIQTE | PPVAIGPNKG | LAEQGPPIQE |
| LUXW01074277.1 | VNGMS-KIMY | YGGSGRKGSI | LTIRLRIETE | PPVAIGPNKG | LAEQGPPIQE |
| AOCR01193540.1 | INGMSWGIVF | YRGSGEERSI | LTIHLRIETE | PPVAVGPNKV | LTEQGPPIQE |
| CM000814.5_b | INGMSWGIVF | YRGSGEERSI | LTIHLRIETE | PPVAVGPNKV | LTEQGPPIQE |
| LUXT01066854.1 | INGMSWGIVF | YRGSGEERSI | LTIHLRIETE | PPVAVGPNKV | LTEQGPPIQE |
| LUXX01079974.1 | INGMSWGIVF | YRGSGEERSI | LTIHLRIETE | PPVAVGPNKV | LTEQGPPIQE |
| KQ003728.1 | INGMSWGIVF | YRGSGEERSI | LTIHLRIETE | PPVAVGPNKV | LTEQGPPIQE |
| LUXY01106894.1 | INGMSWGIVF | YRGSGEERSI | LTIHLRIETE | PPVAVGPNKV | LTEQGPPIQE |
| LUXV01069975.1 | INGMSWGIVF | YRGSGEERSI | LTIHLRIETE | PPVAVGPNKV | LTEKGPPIQE |
| LUXS01071149.1 | INGMSWGIVF | YRGSGEERSI | LTIHLRIETE | PPVAVGPNKV | LTEQGPPIQE |
| LUXW01019135.1 | INGMSWGIVF | YRGSGEERSI | LTIHLRIETE | PPVAVGPNKV | LTEQGPPIQE |
| LUXR01116336.1 | INGMSWGIVF | YRGSGEERSI | LTIHLRIETE | PPVAVGPNKV | LTEQGPPIQE |
| LUXQ01024825.1 | VNGMSWGIVF | YRGSGEERSI | LTIHLRIETE | PPVAVEPNKV | LTEQGPPIQE |
| LIDP01000003.1_a | INGMSWGIVF | YRGSGEERSI | LTIHLRIETE | PPVAVGPNKV | LTEQGPPIQX |
| AOCR01017097.1 | VNGMSWGMVY | YGGSGQPGSI | LTIRLKI-NE | PPMAIGPNTV | LTGQRPPTQG |
| AOCR01011636.1 | VNGMSWGMVY | YGGSGQPGSI | LTIRLKI-NE | PPMAIGPNTV | LTGQRPPTQG |
| AOCR01098665.1 | VNGMSWGMVY | YGGSGQPGSI | LTIRLKI-NE | PPMAIGPNTV | LTGQRPPTQG |
| AOCR01116418.1 | VNGMSWGMVY | YGGSGQPGSI | LTIRLKI-NE | PPMAIGPNTV | LTGQRPPTQG |
| AOCR01214488.1 | VNGMSWGMVY | YGGSGQPGSI | LTIRLKI-NE | PPMAIGPNTV | LTGQRPPTQG |

| | | | | | |
|---|---|---|---|---|---|
| AOCR01214680.1 | VNGMSWGMVY | YGGSGQPGSI | LTIRLKI-NE | PPMAIGPNTV | LTGQRPPTQG |
| PERV-C | VNGMSWGMVY | YGGSGQPGSI | LTIRLKI-NE | PPMAIGPNTV | LTGQRPPTQG |
| KQ004162.1 | VNGMSWGIIY | YAGWGQPGSI | LTIRLKI-SE | PPMAVGPNTV | LTGQRTPT-- |
| LUXQ01128968.1 | VNGMSWGIIY | YAGWGQPGSI | LTIRLKI-SE | PPMAVGPNTV | LTGQRTPT-- |
| LUXR01088996.1 | VNGMSWGIIY | YAGWGQPGSI | LTIRLKI-SE | PPMAVGPNTV | LTGQRTPT-- |
| CM000824.5_a | VNGISWEMVY | YGGSGQPGSI | LTIRLKI-NE | PPMAIGPNTV | LTGQRSPAQG |
| LUXU01026052.1 | VNGISWEMVY | YGGSGQPGSI | LTIRLKI-NE | PPMAIGPNTV | LTGQRSPAQG |
| KQ003791.1 | VNGISWEMVY | YGGSGQPGSI | LTIRLKI-NE | PPMAIGPNTV | LTGQRSPAQG |
| LUXT01031517.1 | VNGISWEMVY | YGGSGQPGSI | LTIRLKI-NE | PPMAIGPNTV | LTGQRSPAQG |
| LUXQ01071671.1 | VNGISWRMVY | YGG-GQPGSI | LTIRLKI-NE | PPMAIGPNTV | LTGQRPPAQG |
| LUXV01065321.1 | VNGISWRMVY | YGG-GQPGSI | LTIRLKI-NE | PPMAIGPNTV | LTGQRPPAQG |
| LUXT01067056.1 | INGMSWGIIY | YTGWDQPGSI | LTIRLKI-SE | PPMAIGPNTV | LTGQRTPT-- |
| LUXU01073709.1 | INGMSWGIIY | YTGWDQPGSI | LTIRLKI-SE | PPMAIGPNTV | LTGQRTPT-- |
| AEMK02000197.1 | INGMSWGIVF | YKYGGGAGST | LTIRLRIETE | PPVAVGPDKV | LAEQGPPALE |
| CM000812.5_a | INGMSWGIVF | YKYGGGAGST | LTIRLRIETE | PPVAVGPDKV | LAEQGPPALE |
| CM000819.5_a | INGMSWGIVF | YKYGGGAGST | LTIRLRIETE | PPVAVGPDKV | LAEQGPPALE |
| LUXR01004647.1 | INGMSWGIVF | YKYGGGAGST | LTIRLRIETE | PPVAVGPDKV | LAEQGPPALE |
| CM000822.5_b | INGMSWGIVF | YKYGGGAGST | LTIRLRIETE | PPVAVGPDKV | LAEQGPPALE |
| PERV-B | INGMSWGIVF | YKYGGGAGST | LTIRLRIETE | PPVAVGPDKV | LAEQGPPALE |
| CM000820.5 | INGMSWGIVF | YKYGGGAGST | LTIRLRIETE | PPVAVGPDKV | LAEQGPPGLE |
| CM000827.5 | INGMSWGIVF | YKYGGGAGST | LTIRLRIETE | PPVAVGPDKV | LAEQGPPALE |
| CM000830.5_d | INGMSWGIVF | YKYGGGAGST | LTIRLRIETE | PPVAVGPDKI | LAEQGPPALE |
| AEMK02000137.1 | INGMSWGIVF | YKYGGGAGST | LTIRLRIETE | PPVAVGPDKV | LAEQGPPALE |
| CM000815.5 | INGMSWGIVF | YKYGGGAGST | LTIRLRIETE | PPVAVGPDKV | LAEQGPPALE |
| CM000814.5_c | INGMSWGIVF | YKYGGGAGST | LTIRLRIETE | PPVAVGPDKV | LAEQGPPALE |
| CM000825.5_b | INGMSWGIVF | YKYGGGAGST | LTIRLRIETE | PPLAVGPDKV | LAEQGPPALE |
| LUXQ01076544.1 | INGMSWGIVF | YKYGGGAGST | LTIRLRIETE | PPVAVGPDKV | LAEQGPPALE |
| LUXY01101100.1 | INGMSWGIVF | YKYGGGAGST | LTIRLRIETE | PPVAVGPDKV | LAEQGPPALE |
| AEMK02000476.1 | INGMSWGIVF | YKYGGGAGST | LTIRLRIETE | PPVAVGPDKV | LAEQGPPALE |
| AOCR01107158.1 | INGMSWGIVF | YKYGGGAGST | LTIRLRIETE | PPVAVGPDKV | LAEQGPPALE |
| CM000826.5_b | INGMSWGIVF | YKYGGGAGST | LTIRLRIETE | PPVAVGPDKV | LAEQGPPALE |
| LUXW01069599.1 | INGMSWGIVF | YKYGGGAGST | LTICLRIETE | PPVAVGSNKV | LAEQGPPALE |
| LUXR01068772.1 | INGMSWGIVF | YKYGGGAGST | LTICLRIETE | PPVAVGSNKV | LAEQGPPALE |
| LUXS01081675.1 | INGMSWGIVF | YKYGGGAGST | LTICLRIETE | PPVAVGSNKV | LAEQGPPALE |
| LUXY01013808.1 | INGMSWGIVF | YKYGGGAGST | LTICLRIETE | PPVAVGSNKV | LAEQGPPALE |
| LUXX01033707.1 | INGMSWGIVF | YKYGGGAGST | LTICLRIETE | PPVAVGSNKV | LAEQGPPALE |
| LUXV01015952.1 | INGMSWGIVF | YKYGGGAGST | LTICLRIETE | PPVAVGSNKV | LAEQGPPALE |
| LUXU01085042.1 | INGMSWGIVF | YKYGGGAGST | LTICLRIETE | PPVAVGSNKV | LAEQGPPALE |
| CM000818.5_a | INGMSWGIVF | YKYGGGAGST | LTIRLRVETE | PPVAVGPNKV | LAEQGPPALE |
| LUXX01080744.1 | INGMSWGIVF | YKYGGGAGST | LTIRLRVETE | PPVAVGPNKV | LAEQGPPALE |
| CM000828.5_d | INGMSWGIVF | YKYGGGAGST | LTIRLRIETE | PPVAVGPNIV | LAEQGPLALE |
| LUXR01022139.1 | INGMSWGIVF | YKYGGGAGST | LTIRLRIETE | PPVAVGPDKV | LAEQGPPALE |
| LUXX01045907.1 | INGMSWGIVF | YKYGGGAGST | LTIRLRIETE | PPVAVGPDKV | LAEQGPPALE |
| LUXW01068183.1 | INGMSWGIVF | YKYGGGAGST | LTIRLRIETE | PPVAVGPNIV | LAEQGPPALE |

| | | | | | |
|---|---|---|---|---|---|
| KQ003125.1 | INGMSWGIVF | WKWGGGAGST | LTIRLRVEPE | PPVAVGPNKV | LAEQGPPALE |
| LUXR01063446.1 | INGMSWGIVF | WKWGGGAGST | LTIRLRVEPE | PPVAVGPNKV | LAEQGPPALE |
| AORO02006834.1 | INGMSWGIVF | WKWGGGAGST | LTIRLRVEPE | PPVAVGPNKV | LAEQGPPALE |
| LUXV01061131.1 | INGMSWGIVF | WKWGGGAGST | LTIRLRVEPE | PPVAVGPNKV | LAEQGPPALE |
| CM000823.5_a | INGMSWGIVF | WKWGGGAGST | LTIRLRVEPE | PPVAVGPNKV | LAEQGPPALE |
| LUXS01013084.1 | INGMSWGIVF | WKWGGGAGST | LTIRLRVEPE | PPVAVGPNKV | LAEQGPPALE |
| LUXW01070394.1 | INGMSWGIVF | WKWGGGAGST | LTIRLRVEPE | PPVAVGPNKV | LAEQGPPALE |
| LUXX01073178.1 | INGMSWGIVF | WKWGGGAGST | LTIRLRVEPE | PPVAVGPNKV | LAEQGPPALE |
| LUXU01008622.1 | INGMSWGIVF | WKWGGGAGST | LTIRLRVEPE | PPVAVGPNKV | LAEQGPPALE |
| LUXT01085568.1 | INGMSWGIVF | WKWGGGAGST | LTIRLRVEPE | PPVAVGPNKV | LAEQGPPALE |
| LUXQ01001340.1 | INGMSWGIVF | WKWGGGAGST | LTIRLRVEPE | PPVAVGPNKV | LAEQGPPALE |
| LUXY01039945.1 | INGMSWGIVF | WKWGGGAGST | LTIRLRVEPE | PPVAVGPNKV | LAEQGPPALE |
| LIDP01000012.1 | INGMSWGIVF | WKWGGGAGST | LTIRLRVEPE | PPVAVGPNKV | LAEQGPPALE |
| AOCR01201355.1 | INGMSWGIVF | WKWGGGAGST | LTIRLRVEPE | PPVAGGPNKV | LAEQGPPA-- |
| CM000830.5_a | INGMNWGIVF | YKYGGGAGST | LTIRLRIETE | PPVAVGPDKV | LTEQGPPALE |
| LUXW01049000.1 | INGMNWGIVF | YKYGGGAGST | LTIRLRIETE | PPVAVGPDKV | LTEQGPPALE |
| eJJRV_NW_004504375.1 | IKGKTWGLRF | YMTGWDKGLT | FTIRLKIEPP | TPRPVGPNKV | LVDQGPPCGI |
| eJJRV_NW_004504378.1 | IKGKTWGFRF | YMTGWDKGLP | FTIRLKIE-- | -PRPVGPNKV | LVDQGPPN-- |
| eJJRV_NW_004504334.1 | IKGKTWGLRF | YMTGWAKGLT | FTI-LKIEPP | TPSSIGPNNV | LADQGLPN-- |
| KoRV | LQGRTWGLRF | YVT-GHPGVQ | LTIRLVITSP | PPVVVGPDPV | LAEQGPPRKI |
| GALV | ITGKTWGLRF | YVS-GHPGVQ | FTIRLKITNM | PAVAVGPDLV | LVEQGPPRTS |
| MMERV_31 | IGGMSWGVVF | YNYGSRPGSL | LHVRLNIESP | PALPVGPNKV | LPEPNRPFLP |
| MMERV_29 | IGGMTWGVVF | YNYGSRPGSL | LHVRLNIESP | PALPVGPNKV | LPEPNRPFLP |
| MMERV_37 | IGGMTWGVVF | YNYGSRPGSL | LHVRLNIESP | PALPVGPNKV | LPEPNRPLLP |
| MCERV_7 | LRGMTWGIVF | YNYGSKPGSL | LHVRLHIESP | LALPVGPNKG | LPEQNQLPLP |
| MMERV_17 | LRGMTWGIVF | YNYGSKPGSL | LHVRLHTERP | LALPVGPNKI | LPEQN-LSLP |
| MPERV_4 | LEGMTWGIVF | YKHGGG-GSR | LQVHLKVEHR | PTQLVGPNQV | LDDQSPPPLI |
| MSERV_11 | DLGQTWG--- | ---------- | -WARIATSGP | PKDRTSPHSA | TPGTGP--RP |
| MMERV_24 | --GMTWGIVF | YRHGGGQGLR | LQVRLKIEHL | PARPVGPNQV | LADQGP--LI |
| MCERV_12 | DLGQTWG--- | ---------- | -RARIATSGP | PKDRTSPDSA | TPGTGP--RP |
| MCERV_16 | HKGMTWDIVF | YRHEGEVGSC | LLIRLKIENP | PAQPVGLNKV | LPDQGLPT-- |
| MSERV_14 | HKGMTWGIVF | YH-GGEVGSR | LLICLKIENP | PAQPVGPNKV | LPDQGPPT-- |
| MMERV_19 | LKGNRWGWRV | YIPLRDPGFI | FTIRLTVRDP | AVTLVGPNKV | LIEQGPPVVP |
| MMERV_5 | LKGNRWGWRV | YIPLRDPGFI | FTIRLTVRDP | AVTLVGPNKV | LIEQGPPVVP |
| MMERV_11 | LKGNRWGWRV | YIPLRDPGFI | FTIRLTVRDP | AVTLVGPNKV | LIEQGPPVVL |
| MMERV_1 | LKGNRWGWRV | YIPLRDPGFI | FTIRLTVRDP | AVTLVGPNKV | LIEQGPPVVL |
| MDEV | LKGNRWGWRV | YIPIRDPGFI | FTIRLTVRDL | AVTSIGPNKV | LTEQAPPVAP |
| MCERV_11 | LKGNCWGWRI | YIGGRDPGSI | FTIRLKIESP | PTLAVGPNSA | L--DGAPAPI |
| OOEV | INGHSWGIRF | YKNGYDSGIL | MTLKLKIETP | APVSIGPNPV | L---GPPPLP |
| ratNor4542 | WRGRTWGLFF | PDRKRDLSAT | FTIQLKIESL | PTQPVCPNQV | LPEQGTPSPD |
| MMERV_21 | TKGYTWGLRI | YKERYDEGLL | FTIRLKIET- | PYNPLGPPTK | FTPQPTPVIA |
| MMERV_10 | TKGYTWGLRI | YKERYDERLL | FTIRLKIET- | PYNPLGPPTK | FTPQPTPVIA |
| MMERV_3 | TKGYTWGLRI | YKERYDEGLL | FTIRLKIET- | PYNPLGPPTK | FTPQPTPVIA |
| MMERV_6 | DGPKVWGLRL | YRSGTDPVTR | FSLTRQVLNG | PRVPIGPNPV | ITDQLPPSRP |

```
MMERV_26            DGPKVWGLRL YRSGTDPVTR FSLTRQVLNG PRVPIGPNPV ITDQLPPSRP
M-CRV               DAPKVWGLRL YRSGADPVTR FSLTRQVLNG PRVPIGPNPV ITEQLPPSQP
MMERV_12            DGPKVWGLRL YRSGTDPVTR FSLTRQVLNG PRIPIGPNPV ITDQLPPSRP
R-MuLV              ITGHYWGLRL YVSGQDPGLT FGIRLKYQNG PRVPIGPNPV LADQLSFPLP
F-MuLV              TTGHYWGLRL YVSGQDPGLT FGIRLSYQNG PRIPIGPNPV LADQLSFPLP
M-MuLV              TTGHYWGLRL YVSGQDPGLT FGIRLRYQNG PRVPIGPNPV LADQQPLSKP
FeLV                DGPKIWGLRL YRTGYDPIAL FTVSRQVSAT PPQAMGPNLV LPDQKPPSRQ
ratNor2288          LEGRRWGLRL YVAGQDPGIL LDIRLKVEPV SPKAMGPHPV LKDQRLLARA
ratNor293           ARGFEWGIRF YVTNRDPGLT FKIKLDIKNT SSSPVGPNKV LPDQG-----
RfRV                LSGLTWGFQL WAWGPHPGGL LTIRLSVET- ISTQVGPNKV LAPLVPTKNP
REV                 CTGIIWDPAA YVSGG----- ---------- -----GPTDM IREESVRERL
ePCRV_KN678005.1    VDGLTWGIVF WKYSRGEGSG LHVRLNIKRP AAQSIGPNKV LSGPRPLPEP
ePCRV_KN680906.1    VDGLTWGIVF WKYGGGEGSG LHVRLNIKRL PAQSIGPNKV LSGPRPLPEP
ePCRV_KN676491.1    INGLTWGVVF WKYGVGVGSS LHIRLKIESP PAQSVGPNKV LSGQKPLPAL
ePCRV_KN676182.1    INGLTWGVVF WKYGGRVGSS LHIRLKIESP PAQSVGPNKV LSGQGPLPAL
ePCRV_KN677924.1    INGLTWGVVF WKYG-GVGSS LHIRLKIESS PAQSVGPNKV LSGQKPLPAP
ePCRV_KN678690.1    INGLTWGVVF WKYGGGVGSS LHIRLKIESP PAQSVGPNKV LSGRGPLPAL
ePCRV_KN676905.1    INGLTWGVVF WKYGGCVWSS LHICLKIESP PAQLVGLNKV LSGQKPLPTL

                    QRPSLNPSDY NTASGSVPTE PNITIKTGAK LFSLIQGAFQ ALNSTTPEAT
                    QRPSPNPSDY NTTSGSVPTE PNITIKTGAK LFNLIQGAFQ ALNSTTPEAT
                    QRPSPNPSDY NTTSGSVPTE PNITIKTGAK LFNLIQGAFQ ALNSTTPEAT
                    QRPSPNPSDY NTTSGSVPTE PNITIKTGAK LFNLIQGAFQ ALNSTTPEAT
                    QRPSPNPSDY NTTSGSVPTE PNITIKTGAK LFNLIQGAFQ ALNSTTPEAT
                    QRPSPNPSDY NTTSGSVPTE PNITIKTGAK LFNLIQGAFQ ALNSTTPEAT
                    QRPSPNPSDY NTTSGSVPTE PNITIKTGAK LFSLIQGAFQ ALNSTTPEAT
                    QRPSPNPSDY NTTSGSVPTE PNITIKTGAK LFSLIQGAFQ ALNSTTPEAT
                    QRPSPNPSDY NTTSGSVPTE PNITIKTGAK LFSLIQGAFQ ALNSTTPEAT
                    QRPSPNPSDY NTTSGSVPTE PNITIKTGAK LFSLIQGAFQ ALNSTTPEAT
                    QRPSPNPSDY NTTSGSVPTE PNITIKTGAK LFSLIQGAFQ ALNSTTPEAT
                    QRPSPNPSDY NTTSGSVPTE PNITIKTGAK LFSLIQGAFQ ALNSTTPEAT
                    QRPSPNPSDY NTTSGSVPTE PNITIKTGAK LFSLIQGAFQ ALNSTTPEAT
                    QRPSPNPSDY NTTSGSVPTE PNITIKTGAK LFSLIQGAFQ ALNSTTPEAT
                    QRPSPNPSDY NTTSGSVPTE PNITIKTGAK LFSLIQGAFQ ALNSKTPEAT
                    QRPSPNPSDY NTTSGSVPTE PNITIKTGAK LFSLIQGAFQ ALNSTTPEAT
                    QRPSPNPSDY NTTSGSVPTE PNITIKTGTK LFSLIQGAFQ ALNSTTPEAT
                    QRPSPNPSVY NTTSGLVPPE PNFTIKTGAK LFSLIQGAFQ ALNSTTPEAT
                    QRPSPNPSVY NTTSGLVPPE PNFTIKTGAK LFSLIQGAFQ ALNSTTPEAT
                    QRPSPNPSDY NTTSGSVPTE PNITIKTGAK LFNLIQGAFQ ALNSTTPEAT
                    QRPSPNPSDY NTTSGSVPTE PNITIKTGAK LFNLIQGAFQ ALNSTTPEAT
                    QRPSPNPSDY NTTSGSVPTE PNITIKTVVK LFSLIQGDFQ ALNSTTPEAT
                    QRPSPNPSDY NTTSGSVPTE PNITIKTAAK LFSLIQGDFQ ALNSTTPEAT
                    QRPSPNPSDY NTTSGSVPTE PNITIKTGAK LFSLIQGAFQ ALNSTTPEAT
                    QRPSPNPSDY NTTSGSVPTE PNITIKTGAK LFSLIQGAFQ ALNSTTPEAT
```

QRPSPNPSDF NTTSGSVPTE PNITIKTGAK LFSLIQGAFQ ALNSKAPEAT
QRPSPNPSDF NTTSGSVPTE PNVPVKTGQR LFSLIQGAFQ ALNSTTPEAT
QRPSPNPSDF NTTSGSVPTE PNVPVKTGQR LFSLIQGAFQ ALNSTTPEAT
QRPSPNPSDF NTTSGSVPTE PNVPVKTGQR LFSLIQGAFQ ALNSTTPEAT
QRPSPNPSDF NTTSGSVPTE PNVPVKTGQR LFSLIQGAFQ ALNSTTPEAT
QRPSPNPSDF NTTSGSVPTE PNVPVKTGQR LFSLIQGAFQ ALNSTTPEAT
QRPSPNPSDF NTTSGSVPTE PNVPVKTGQR LFSLIQGAFQ ALNSTTPEAT
QRPSPNPSDF NTTSGSVPTE PNVPVKTGQR LFSLIQGAFQ ALNSTTPEAT
QRPSPNPSDF NTTSGSVPTE PNVPVKTGQR LFSLIQGAFQ ALNSTTPEAT
QRPSPNPSDF NTTSGSVPTE PNVPVKTGQR LFSLIQGAFQ ALNSTTPEAT
QRPSPNPSDF NTTSGSVPTE PNVPVKTGQR LFSLIQGAFQ ALNSTTPEAT
QRPSPNPSDF NTTSGXVPTE PNVPVKTGQR LFSLIQGAFQ ALNSTTPEAT
PGPS-----S NITSGSDPTE SNSTTKMGAK LFSLIQGAFQ ALNSTTPEAT
PGPS-----S NITSGSDPTE SNSTTKMGAK LFSLIQGAFQ ALNSTTPEAT
PGPS-----S NITSGSDPTE SNSTTKMGAK LFSLIQGAFQ ALNSTTPEAT
PGPS-----S NITSGSDPTE SNSTTKMGAK LFSLIQGAFQ ALNSTTPEAT
PGPS-----S NITSGSDPTE SNSTTKMGAK LFSLIQGAFQ ALNSTTPEAT
PGPS-----S NITSGSDPTE SNSTTKMGAK LFSLIQGAFQ ALNSTTPEAT
PGPS-----S NITSGSDPTE SNSTTKMGAK LFSLIQGAFQ ALNSTTPEAT
PGPS-----S DITSELDPTE SNSTTKTGAK LFSLIQGAFQ ALKSTSPEAT
PGPS-----S DITSELDPTE SNSTTKTGAK LFSLIQGAFQ ALKSTSPEAT
PGPS-----S DITSELDPTE SNSTTKTGAK LFSLIQGAFQ ALKSTSPEAT
PGPF-----F NITSGSDPTE SNSTTKTGTK LFSLIQGAFQ ALNSTTPEAT
PGPF-----F NITSGSDPTE SNSTTKTGTK LFSLIQGAFQ ALNSTTPEAT
PGPF-----F NITSGSDPTE SNSTTKTGTK LFSLIQGAFQ ALNSTTPEAT
PGPF-----F NITSGSDPTE SNSTTKTGTK LFSLIQGDFQ ALNSTTPEAT
PGPF-----F NITSGSDPTE SNSTTKTGTK LFSLIQGAFQ ALNSTTPEAT
PGPF-----F NITSGSDPTE SNSTTKTGTK LFSLIQGAFQ ALNSTTPEAT
PGPS-----S DITSKLDPTE SNSTTKTGTK LFSLIQGAFQ ALNSTTPEAT
PGPS-----S DITSKLDPTE SNSTTKTGTK LFSLIQGAFQ ALNSTTPEAT
PRPDITQPPS NSTTGLIPTS PGVPVKTGQR LFSLIQGAFQ AINSTDPDAT
PRPDITQPPS NGTTGLIPTS PGVPVKTGQR LFSLIQGAFQ AINSTDPDAT
PRPDITQPPS NGTTGLIPTS PGVPVKTGQR LFSLIQGAFQ AINSTDPDAT
PRPDITQPPS NGTTGLIPTS PGVPVKTGQR LFSLIQGAFQ AINSTDPDAT
PRPDITQPPS NGTTGLIPTS PGVPVKTGQR LFSLIQGAFQ AINSTDPDAT
PRPDITQPPS NGTTGLIPTS PGVPVKTGQR LFSLIQGAFQ AINSTDPDAT
PRPDITQPPS NGTTGLIPTS PGVPVKTGQR LFSLIQGAFQ AINSTDPDAT
PRPDITQPPS NGTTGLIPTS PGVPVKTGQR LFSLIQGAFQ AINSTDPDAT
PRPDITQPPS NGTTGLIPTS PGVPVKTGQR LFSLIQGAFQ AINSTDPDAT
PRPDITQPPS NGTTGLIPTS PGVPVKTGQR LFSLIQGAFQ AINSTDPDAT
PRPDITQPPS NGTTGLIPTS PGVPVKTGQR LFSLIQGAFQ AINSTDPDAT
PRPDITQPPS NGTTGLIPTS PGVPVKTGQR LFSLIQGAFQ AINSTDPDAT
PRPDITQTPS NGTTGLIPTS PGVPVKTGQR LFSLIQGAFQ AINSTDPDAT
PRPDITQPPS NGTTGLIPTS PGVPVKTGQR LFSLIQGAFQ AINSTDPDAT

```
PRPDITQPPS NGTTGLIPTS PGVPVKTGQR LFSLIQGAFQ AINSTDPDAT
PRPDITQPPS NGTTGLIPTS PGVPVKTGQR LFSLIQGAFQ AINSTDPDAT
PRPDITQPPS NGTTGLIPTS PGVPVKTGQR LFSLIQGAFQ AINSTDPDAT
PRPDITQPPS NGTTGLIPTS PGVPVKTGQR LFSLIQGAFQ AINSTDPDAT
PRPDITQLPD NGTTGLIPTS PGVPVKTGQR LFRLIQGAFQ AITSTDPDAT
PRPDITQLPD NGTTGLIPTS PGVPVKTGQR LFRLIQGAFQ AITSTDPDAT
PRPDITQLPD NGTTGLIPTS PGVPVKTGQR LFRLIQGAFQ AITSTDPDAT
PRPDITQLPD NGTTGLIPTS PGVPVKTGQR LFRLIQGAFQ AITSTDPDAT
PRPDITQLPD NGTTGLIPTS PGVPVKTGQR LFRLIQGAFQ AITSTDPDAT
PRPDITQLPD NGTTGLIPTS PGVPVKTGQR LFRLIQGAFQ AITSTDPDAT
PRPDITQLPD NGTTGLIPTS PGVPVKTGQR LFRLIQGAFQ AITSTDPDAT
PRPDITQPPG NGTTGLIPTS PGVPVKTGQR LFSLIQGVFQ AINSTDPDAT
PRPDITQPPG NGTTGLIPTS PGVPVKTGQR LFSLIQGVFQ AINSTNSDAT
PRPDITQPPD NSTTGLIPTS PGVPVKTGQR LFSLIQGAFQ AINSTDPDAT
PRPDITQPPD NSTTGLIPTS PGVPVKTGQR LFSLIQGAFQ AINSTDPDAS
PRPDITQPPD NSTTGLIPTS PGVPVKTGQR LFSLIQGAFQ AINSTDPDAT
PRPDITQPPD NSTTGLIPTS PGVPVKTGQR LFSLIQGAFQ AINSTDPDAS
-RPNTTQPPG NSTAGLIPT- ------TGQR LFSLIQGAFQ VLNSTDPDAT
-RPNTTQPPG NSTAGLIPT- ------TGQR LFSLIQGAFQ VLNSTDPDAT
-RPNTTQPPG NSTAGLIPT- ------TGQR LFSLIQGAFQ VLNSTDPDAT
-RPNTTQPPG NSTAGLIPT- ------TGQR LFSLIQGAFQ VLNSTDPDAT
-RPNTTQPPG NSTAGLIPT- ------TGQR LFSLIQGAFQ VLNSTDPDAT
-RPNTTQPPG NSTAGLIPT- ------TGQR LFSLIQGAFQ VLNSTDPDAT
-RPNTTQPPG NSTAGLIPT- ------TGQR LFSLIQGAFQ VLNSTDPDAT
-RPNTTQPPG NSTAGLIPT- ------TGQR LFSLIQGAFQ VLNSTDPDAT
-RPNTTQPPG NSTAGLIPT- ------TGQR LFSLIQGAFQ VLNSTDPDAT
-RPNTTQPPG NSTAGLIPT- ------TGQR LFSLIQGAFQ VLNSTDPDAT
-RPNTTQPPG NSTAGLIPT- ------TGQR LFSLIQGAFQ VLNSTDPDAT
-RPNTTQPPG NSTAGLIPT- ------TGQR LFSLIQGAFQ VLNSTDPDAT
-RPNTTQPPG NSTAGLIPT- ------TGQR LFSLIQGAFQ VLNSTDPDAT
-RPDITQPPS NGTTGLIPTS PGVPVKTGQR LFSLIQGAFQ AINSTDPDAT
PRPDITQPPG NGTTGLIPTS PSVPVKTGQR LFSLIQGAFQ AINSTDPDAT
PRPDITQPPG NGTTGLIPTS PSVPVKTGQR LFSLIQGAFQ AINSTDPDAT
AAPTHSGSQD IMTARTTSTN PEAPHRTRQR LFNLVQGAFL AINATNSNAT
VALVSAKTRR IMTARTTLTS PEAPHGKGQR LFNLIQGAFL AINATNPNAT
AALVAAGTCR VTTTRITPTS PVAPHKTGQR LLNLIQGAFL TINATNPNAT
PSPPAS---- -----PIPTA PTPSPTTGDR LFGLVQGAFL ALNATNPEAT
LAPPPSLPDS NATSAQTPTT LTPPPTTGDR LFDLVQGAFL TLNATNPGAT
RTHQMSSPQK NSSPVLYSIA PVELSPTRQR LLGLLQGAFS VLNGTNPNAT
RTHQMSSPQK NSSPVLYSIA PVELSPTRQR LLGLLQGAFS VLNGTNPNAT
RTHQMSSPQK NSSPVLYSTA PVE--PTQQR LLGLLQGALS VLNGTNPNAT
----VSSPRG ILSSTLPPIT PVGPPPTRQR LFSLLQGAFS VLNSTNPNAT
----VSSPQG ILSSALPPIT PVGPPPTRQR LFSLLQGAFS VLNSTNPNAT
PQPFPTLPTS -PHSSLNPFS NVAPAKTGQR LIILVQGAFS VIN-TNPDMT
```

```
PTGTATFPDF ---AHLSPFQ PCCPSQDRTD INQSCTGREP LVTEPIPTLS
PQPSPTLPTS -PQSSLTPSS NVAPAKTGQT LINLVQGAFS MINRTNPDMT
PTGTPTLPTS -PHSSLTPFS NVAPANTGQR LINLVQGAFS MINRTNPDMT
--------PI LGTSSHSPST PLALPSIGQR LLNLIQGAFS VLNGSNPNMM
--------PI LGTSTPSPST PLALPSIGQR LLNLIQGAFS VLNGSNPDMM
AVPAPPTPQP NSLGTNTPLK PPPPLGTENR LVSLVQGAFL ALNRTNPNMT
AVPAPPTPQP NSLGTNTPLK PPPPLGTENR LVSLVQGAFL ALNRTNPNMT
AVPAPPTPQP NSLGTNTLLK PPPPLGTEDR LVSLVQGAFL VLNRTNPNMT
AVPAPPTPQP NSLGTNTLLK PPPPLGTEDR LVSLVQGAFL VLNRTNPNMT
AVPAPPTSRP YSLET----- -PPLLDTENR LVSLVQGAFL VLNRTNPNMT
PYIPPPGKTP FSTAPYPPLT PPSPTRTEDR LFNLLDGAFT VLNRTNPSAT
PPLLPPAPKN NGTSGLKET- -IKPGTRQDR MLSLVQAAFT VLNATNPEAT
APESLPSRPS LTSSPLFP-- -----SIGKR LLNLIQGAFT VLNSFSPNMT
DI-------- --TQPPTPQV PTPEIPSRQR MFNLVRGAFY ALNRTDPSAT
DI-------- --TQPPTPQV PTPEIPSRQR MFNLVRGAFY ALNRTDPSAT
DI-------- --TQPPTPQV PTPAIPSRQR MFNLVRGAFY ALNRTDPSAT
VLPRPPQPPP PGAASIVPET ASQQPGTGDR LLNLVDGAYQ ALNLTSPDKT
VLPRPPQPPP PGAASIVPET ASQQPGTGDR LLNLVDGAYQ ALNLTSPDKT
VLPRPPHPPP SGAASMVPGA PSQQPGTGDR LLNLVKGAYQ ALNLTSPDRT
VLPRPPQPSP TGAASI---- ---QPGTGDR LLNLVDGAYQ ALNLTSPDKT
NLPKPAKSPP VSTPTMIPST PPPPAGTGDR LLNLVQGAYQ ALNLTNPDKT
NLPKPAKSPP ASTPTLIPST PPPPAGTGDR LLNLVQGAYQ ALNLTNPDKT
KV----KSPS VTKPPGTPLS PLPPAGTENR LLNLVDGAYQ ALNLTSPDKT
STGSKVATQR LTESAPRSVA PPKRIGTGDR LINLVQGTYL ALNATDPNKT
RPKVTTTLKP VTFTHSEVFV PPHLPSAQER MFKLIQGTFQ ALNHSNPNLT
---------- ---------- ------TGDR LFNLIRGAYQ ALNQSNPNDT
GSRDKDTTGR IGTQPKTSVT PPATQTTEDS LRKLVRTVYE TLNATSPHLT
EI-------- --IRHSYPSL PPRGVDLDPQ TSDILEATHQ VLNATNPRLA
T--------- -LSPSSIPVT PPAPINTEQR LLNLVQGAFS VLNTSDPSIT
T--------- -LSPSSIPVT PSAPINTEQR LLNLVQGAFN VLNTSDPSIT
T--------- -LSPSSIPIT PPAPTNTEQR LLNLVQGAFS VLNTSNPSIT
T--------- -LSPSSIPIT PPAPTNTEQR LLNLVQGAFS VLNTSNPSII
T--------- -LSPSSIPIM PPTSTNTEQR LLNLVQ-AFS VLNTSNSSIT
T--------- ---------- ---PTNTEQR LLNLVQGAFS VLNTSNPSIT
T--------- -LSPSSIPNN PLHQHRAE-- IAEFSTGSFQ VLNTSNPSIT

SSCWLCLASG PPYYEGTARG GKFNVTKEHD QCTWGSQNKL TLTEVSGKGT
SSCWLCLASG PPYYEGMARG GKFNVTKEHD QCTWGSQNKL TLTEVSGKGT
SSCWLCLASG PPYYEGMARG GKFNVTKEHD QCTWGSQNKL TLTEVSGKGT
SSCWLCLASG PPYYEGMARG GKFNVTKEHD QCTWGSQNKL TLTEVSGKGT
SSCWLCLASG PPYYEGMARG GKFNVTKEHD QCTWGSQNKL TLTEVSGKGT
SSCWLCLASG PPYYEGMARG GKFNVTKEHD QCTWGSQNKL TLTEVSGKGT
SSCWLCLASG PPYYEGMARG GKFNVTKEHD QCTWGSQNKL TLTEVSGKGT
SSCWLCLASG PPYYEGMARG GKFNVTKEHD QCTWGSQNKL TLTEVSGKGT
```

SSCWLCLASG PPYYEGMARG GKFNVTKEHD QCTWGSQNKL TLTEVSGKGT
SSCWLCLASG PPYYEGMARG GKFNVTKEHD QCTWGSQNKL TLTEVSGKGT
SSCWLCLASG PPYYEGMARG GKFNVTKEHD QCTWGSQNKL TLTEVSGKGT
SSCWLCLASG PPYYEGMARG GKFNVTKEHD QCTWGSQNKL TLTEVSGKGT
SSCWLCLASG PPYYEGMARG GKFNVTKEHD QCTWGSQNKL TLTEVSGKGT
SSCWLCLASG PPYYEGMARG GKFNVTKEHD QCTWGSQNKL TLTEVSGKGT
SSCWLCLASG PPYYEGMARG GKFNVTKEHD QCTWGSQNKL TLTEVSGKGT
SSCWLCLASG PPYYEGMAKG GKFNVTKEHD QCTWGSQNKL TLTEVSGKGT
SSCWLCLASG PPYYEGMARG GKFNVTKEHD QCTWGSQNKL TLTEVSEKGT
SSCWLCLASG PPYYEGMARG GKFNVTKEHD QCTWGSQNKL TLTEVSGKGT
SSCWLCLALS PPYYEGMARG GKFNVTKEHD QCTWGSQNKL TLTEVSGKGT
SSCWLCLASG PPYYEEMARE GKFNVTKEHD QCTWGSQNKL TLTEVSGKGT
SSCWLCLTSG PPYYEGMARG GKFNVTKEHD QCTWGSQNKL TLTEVSEKDT
SSCWLCLASG PPYYEGMARG GKFNVTKEHD QCTWGSQNKL TLTEVSGKGT
SSCWLCLASG PPYYEGMARG GKFNVTKEHD QCTWGSQNKL TLTEVSGKGT
SSCWLCLASG PPYYEKMARG GKFNVSKKHD QCT-GSQNKL TLTEVSGKDT
SSCWLCLASG PPYYEKMARG GKFNVTKKHD QCT-GSQNKL TLTEVSGKDT
SSCWLCLASG PPYYEGMARG GKFNVTKEHD QCTWGSQNKL TLTEVSGKGT
SSCWLCLASG PPYYGGMARE GKFNVTKEHD QCTWGSQNKL TLTEVSGKGT
SSCWLCLASG PPYYGGMARE GKFNVTKEHD QCTWGSQNKL TLTEVSGKGT
SSCWLCLASG PPYYGGMARE GKFNVTKEHD QCTWGSQNKL TLTEVSGKGT
SSCWLCLASG PPYYGGMARE GKFNVTKEHD QCTWGSQNKL TLTEVSGKGT
SSCWLCLASG PPYYGGMARE GKFNVTKEHD QCTWGSQNKL TLTEVSGKGT
SSCWLCLASG PPYYGGMARE GKFNVTKEHD QCTWGSQNKL TLTEVSGKGT
SSCWLCLASG PPYYGGMARE GKFNVTKEHD QCTWGSQNKL TLTEVSGKGT
SSCWLCLASG PPYYGGMARE GKFNVTKEHD QCTWGSQNKL TLTEVSGKGT
SSCWLCLASG PPYYEGMARE GKFNVTKEHD QCTWGSQNKL TLTEVSGKGT
SSCWLCLASG PPYYEGMARE GKFNVTKEHD QCTWGSQNKL TLTEVSGKGT
SSCWLCLASG PPYYXGMARR GKFNVTKEHD QCTWGSQNKL TLTEVSGKGT
SSCWLCLALG PPYYEGMARR GKFNVTKEHD QCTWGSQNKL TLTEVSGKGT
SSCWLCLALG PPYYEGMARR GKFNVTKEHD QCTWGSQNKL TLTEVSGKGT
SSCWLCLALG PPYYEGMARR GKFNVTKEHD QCTWGSQNKL TLTEVSGKGT
SSSWLCLASG PPYYEGMARR GKFNVTKEHD QCTWGSQNKL TLTEVSGKGT
SSSWLCLASG PPYYEGMARR GKFNVTKEHD QCTWGSQNKL TLTEVSGKGT
SSSWLCLASG PPYYEGMARR GKFNVTKEHD QCTWGSQNKL TLTEVSGKGT
SSCWLCLASG PPYYEGMARR GKFNVTKEHD QCTWGSQNKL TLTEVSGKGT
SSCWLCLASG PPYYEGMAKR GKFNVTKEHD QCTWGSQNKL TLTEVSGKGT
SSCWLCLASG PPYYEGMARG GKFNVTKEHD QCTWGSQNKL TLTEVSGKGT
SSCWLCLALS PPYYEGMARG GKFNVTKEHD QCTWGSQNKL TLTEVSGKGT
SSCWLCLASG PPYYEGMARG GKFNVTKEYD QCTWGSQNKL TLTEVSGKGT
SSCWLCLASG PPYYEGMARG GKFNVTKEYD QCTWGSQNKL TLTEVSGKGT
SSCWLCLASG PPYYEGMARG GKFNVTKEYD QCTWGSQNKL TLTEVSGKGT
SSCWLCLASG PPYYEGMARG GKFNVTKEYD QCTWGSQNKL TLTEVSGKGT
SSCWLCLASG PPYYEGMARG GKFNVTKEHD QCTWGSQNKL TLTEVSGKGT

SSCWLCLASG PPYYGGMARE GKFNVTKEHD QCTWGSQNKL TLTEVSGKGT
SSCWLCLTSG PPYYKKMAKR EKFNVTKKHD QCTWGSQNKL TLTEVSGKDT
SSCWLCLASG PPYYEGMARG GKFNVTKEHD QCTWGSQNKL TLTEVSGKDT
SSCWLCLSSG PPYYEGMAKE GKFNVTKEHN QCTWGSRNKL TLTEVSGKGT
SSCWLCLSSG PPYYEGMAKE GKFNVTKEHN QCTWGSRNKL TLTEVSGKGT
SSCWLCLSSG PPYYEGMAKE GKFNVTKEHN QCTWGSRNKL TLTEVSGKGT
SSCWLCLSSG PPYYEGMAKE GKFNVTKEHN QCTWGSRNKL TLTEVSGKGT
SSCWLCLSSG PPYYEGMAKE GKFNVTKEHN QCTWGSRNKL TLTEVSGKGT
SSCWLCLSSG PPYYEGMAKE GKFNVTKEHN QCTWGSRNKL TLTEVSGKGT
SSCWLCLSSG PPYYEGMAKE GKFNVTKEHN QCTWGSRNKL TLTEVSGKGT
SSCWLCLSSG PPYYEGMAKE GKFNVTKEHN QCTWGSRNKL TLTEVSGKGT
SSCWLCLSSG PPYYEGMAKE GKFNVTKEHN QCTWGSRNKL TLTEVSGKGT
SSCWLCLSSG PPYYEGMAKE GKFNVTKERN QCTWGSRNKL TLTEVSGKGT
SSCWLCLSSG PPYYEGMAKE GKFNVTKEHN QCTWGSRNKL TLTEISGKGT
SSCWLCLSSG PPYYEGMAKE GKFNVTKEHN QCTWGSQNKL TLTEVSGKGT
SSCWLCLSSG PPYYEGMAKE GKFNVTKEHN QCTWGSRNKL TLTEVSGKGT
SSCWLCLSSG PPYYEGMAKE GKFNVTKEHN QCTWGSRNKL TLTEVSGKGT
SSCWLCLSSG PPYYEGMAKE GKFNVTKEHN QCTWGSRNKL TLTEVSGKGT
SSCWLCLSSG PPYYEGMAKE GKFNVTKEHN QCTWGSRNKL TLTEVSGKGT
SSCWLCLSSG PPYYEGMAKE GKFNVTKEHN QCTWGSRNKL TLTEVSGKGT
SSCWLCLSSG PPYYEGMAKE EKFNVTKEHN QCTWGSRNKL TLTEVSEKGT
SSCWLCLSSG PPYYEGMARE GKFDVTKKHN QCTWGSRNKL TLTEVSGKGT
SSCWLCLSSG PPYYEGMARE GKFDVTKKHN QCTWGSRNKL TLTEVSGKGT
SSCWLCLSSG PPYYEGMARE GKFDVTKKHN QCTWGSRNKL TLTEVSGKGT
SSCWLCLSSG PPYYEGMARE GKFDVTKKHN QCTWGSRNKL TLTEVSGKGT
SSCWLCLSSG PPYYEGMARE GKFDVTKKHN QCTWGSRNKL TLTEVSGKGT
SSCWLCLSSG PPYYEGMARE GKFDVTKKHN QCTWGSRNKL TLTEVSGKGT
SSCWLCLSSG PPYYEGMARE GKFDVTKEHN QCTWGSRNKL TLTEVSGKGT
SSCWLCLSSG LLIMRNGRR- -KFNVTKEHN QCTWGSRNKL TFTEVSGKGT
SFCWLCLSSG PPYYEGMARE GKFNVTKEHN QCTWGSRNKL TFTEVSGKGT
SSCWLCLSSG PPYYEGMARE GKFNVTKEHN QCTWGSRNKL TLTEVSGKGT
SSCWLCLSSG PPYYEGMARE GKFNVTKEHN QCTWGSRNKL TLTEVSGKGT
SSCWLCLSSG PPYYEGMARE GKFNVTKEHN QCTWGSRNKL TLTEVSGKGT
SSCWLCLSSG PPYYEGMARE GKFNVTKEHN QCTWGSRNKL TLTEVSGKGT
SSCWLCLSSG PPYYEGMARE GKFNVTKEHN QCTWGSRYKL TLTEVSGKGT
SSCWLCLSSG PPYYEGMAKE GKFNVTKEHN QCTWGSRNKL TLTEVSGKGT
SSCWLCLSSG PPYYEGMARE GKFNVTKEHN QCTRGSRNKL TLTEVSGKGT
SSCWLCLSSG PPYYEGMARE GKFNVTKEHN QCTRGSRNKL TLTEVSGKGT
SSCWLCLSSG PPYYEGMARE GKFNVTKEHN QCTRGSRNKL TLTEVSGKGT
SSCWLCLSSG PPYYEGMARE GKFNVTKEHN QCTRGSRNKL TLTEVSGKGT
SSCWLCLSSG PPYYEGMARE GKFNVTKEHN QCTRGSRNKL TLTEVSGKGT
SSCWLCLSSG PPYYEGMARE GKFNVTKEHN QCTRGSRNKL TLTEVSGKGT
SSCWLCLSSG PPYYEGMARE GKFNVTKEHN QCTRGSRNKL TLTEVSGKGT
SSCWLCLSSG PPYYEGMARE GKFNVTKEHN QCTRGSRNKL TLTEVSGKGT

```
SSCWLCLSSG PPYYEGMAKE GKFNVTKEHN QCTWGSRNKL TLTEVSGKGT
SSCWLCLSSG PPYYEGMAKE GKFNVTKEHN QCTWGSRNKL TLTEVSGKGT
SSCWLCLSSG PPYYEGMARE GKFNVTKEHN QCTRGSRXKL TXTEVSGKGT
SSCWLCLSSG PPYYEGMAKE GKFNVTKEHN QCTRGSRNKL TLTEVSGKGT
SSCWLCLSSG PPYYEGMARE GKFNVTKKHD RCTRVSQNKL TLTEVSGKGT
SSCWLCLSSG PPYYEGMARE GKFNVTKKHD RCTRVSQNKL TLTEVSGKGT
SACWLCLSSG PSYYEGMATM GEFNVTKEHR QCSWGTKNKL TPPEVSGRGT
SACWLCLSLG PPYYEGMATM GEFNVTKEYS RCPWGTKNKL TLPEVSGRGT
SACWLCLSSG PPNYEGVANV GEFNVTKEHR QCSWGIKNKL TFPDISGSGT
ESCWLCLALG PPYYEGIATP GQVTYASTDS QCRWGGKGKL TLTEVSGLGL
ESCWLCLAMG PPYYEAIASS GEVAYSTDLD RCRWGTQGKL TLTEVSGHGL
KSCWLCLTSG PPYYEGMAVM GTYNNTTSHD HCRWGRSHTL TLTEVSGRGT
KSCWLCLTSG PPYYEGMAVI GTYNNTTSHD HCRWGRSHTL TLTEVSGRGT
KSCWLCLTSG PP-YEGMAVM GTYNNTTSHD HCSWGRSHTL TLTEVSGRGT
KSCWLCLASG PPYYEGMAVM GTYNNTTSHD PCSWRRSHTF TLTEVLSTGT
KSCWLCLASG PPCYEGMAVM RT-NNTTSHD HCSWGCSHTL TLTEVSGTGT
ESCWLCLALG PP-YETIAVS GTFNTTSSHE ACVWGTSHRL TLTEVTGIGT
----LVGFAL QAYYKEIAVS GTFTTTSSHE ARVWGTSQRL TLPEVT--GT
ESCWLCLASV PPYYKEIAVS GTFITTSSHE ACVWGTSQRL TLPEVT--GT
ESCWLCLASG LPYYKEIAVS GTFNTTSSHD ACVWGT-NRL TLPEVTGTGT
ESCWLCLASG PPYYKGIAVS EAFNHTTSQD ACVWGTSYRL TLTEITGVGT
ESCWLCLALG PPYYEGIAVS GAFNNTTSQD ACVWGTSFRL TLTEITGVGT
QSCWLCYTSS PPYYEGIAQI RTYNITSDHS QCLWGENRKL TLAAVSGRGL
QSCWLCYTSS PPYYEGIAQI RAYNITSDHS QCLWGENRKL TLAAVSGRGL
QSCWLCYASS PPYYEGIAQI RTYNITSDHS QCLWGENRKL TLAAVSGRGL
QSCWLCYASS PPYYEGIAQI RTYNTTSDHS QCLWGENRKL TLAAVSGRGL
QSCWLCYASN PPYYEGIAQT RTYNITSDHS QCLWGENRKL TLTAVSGNGL
QSCWLCFAAN PPYYEGIAQV RKYNVTSDYS HCPWGSQRKL TLTAVSGAGL
KSCWLCYAAP PPYYDAIGYS SNYTNVSSPD HCRWKQESKL TLSSVFGNGT
ESCWLCLSSG PPIIHNTFRS GTFNNTTSHD NCAWNSKNRL TLTQVSGKGT
EDCWLCLSSG PPYYEGIAFN GDFNRTSSHT SCSWGTGQKL TLTEVSARGL
EDCWLCLSPG PPYYEGIAFN GDFNRTSSHT SCSWGTGQKL ILTKVSARGL
EDCWLCLSSG PPYYEGIDFN GDFNRTSSHT SCSWGTGQKL TLTEVSARGL
QECWLCLVAG PPYYEGVAVL GTYSNHTSAA NCSVASQHKL TLSEVTGQGL
QECWLCLVAG PPYYEGVAVL GTYSNHTSAA NCSVASQHKL TLSEVTGQGL
QGCWLCLVSG PPYYEGVAVL GTYSNHTSAA NCSVASQHKL TLSEVTGQGL
QECWLCLVSG PPYYEGVAVL GTYSNHTSAA NCSVASQHKL TLSEVTGQGL
QECWLCLVSG PPYYEGVAVL GTYSNHTSAT NCSVASQHKL TLSEVTGRGL
QECWLCLVSG PPYYEGVAVL GTYSNHTSAA NCSVASQHKL TLSEVTGRGL
QECWLCLVAG PPYYEGVAVL GTYSNHTSAA NCSVASQHKL TLSEVTGQGL
KDCWLCLVSR PPYYEGIAIL GNYSNQTNPP SCLSTPQHKL TISEVSGQGL
TECWLCLSTS PSYYEGIATY ANFTKETNPE ACLHTNKPRL TLTEVSGAGT
ESCWLCLTAS PPYYEGI--- -------SHG ECAWQDQGKV TITEVFDSGL
TSCWLCYDVK PPFYEAIGLN ATYNASNGKS QCSWGDRIGL TLQLVSGNGT
```

```
ENCWLCMTLG TPIPAAIPAN GEVTL---DG NCSLSLPFRV QPTGSVDVNC
NSCWLCLASR PPYYEGVAFS GNFNNATSHS LCTWGTQNKL TLTEVTGKGT
NSCWLCLASR PPYYEGVAFS ENFNNATSHS LCTWGTQNKL TLTEVTGKGT
KSCWLCLASR PPYYEGVAFS GNFNNTTSHS LCTWGTQNKL TLTEVTGKGT
KSCWLCLASR PPYYEGVAFS GNFNNTTSHS LCTWGTQNKL TLTEVTGKGT
KSCWLCLASR PPYYEGVALS GNFNNTTSHS LCTWGTQNKL TLTEVTGKGT
KSCWLCLASR PPYYEGVAFS GNFSNTTSHS LCTWGTQNKL TLTEVTGKGT
NSCWLYLASR PPYYEGVAFS GNFNNTTSHS LCTWGTQNKL TLTEVTGKGT

CIGRVPPSHQ HLCNHTEAFN RTSESQYLVP GYDRWWACNT GLTPCVSTLV
CIGMVPPSHQ HLCNHTKAFN RTSESQYLVP GYDRWWACNT GLTPCVSTLV
CIGMVPPSHQ HLCNHTEAFN RTSESQYLVP GYDRWWACNT GLTPCVSTLV
CIGMVPPSHQ HLCNHTEAFN RTSESQYLVP GYDRWWACNT GLTPCVSTLV
CIGMVPPSHQ HLCNHTEAFN RTSESQYLVP GYDRWWACNT GLTPCVSTLV
CIGMVPPSHQ HLCNHTEAFN RTSESQYLVP GYDRWWACNT GLTPCVSTLV
CIGMVPPSHQ HLCNHTEAFN RTSESQYLVP GYDRWWACNT GLTPCVSTLV
CIGMVPPSHQ HLCNHTEAFN RTSESQYLVP GYDRWWACNT GLTPCVSTLV
CIGMVPPSHQ HLCNHTEAFN RTSESQYLVP GYDRWWACNT GLTPCVSTLV
CIGMVPPSHQ HLCNHTEAFN RTSESQYLVP GYDRWWACNT GLTPCVSTLV
CIGMVPPSHQ HLCNHTEAFN RTSESQYLVP GYDRWWACNT GLTPCVSTLV
CIGRVPPSHQ HLCNHTEAFN RTSESQYLVP GYDRWWACNT GLTPCVSTLV
CIGRVPPSHQ HLCNHTEAFN RTSESQYLVP GYDRWWACNT GLTPCVSTLV
CIGRVPPSHQ HLCNHTEAFN QTSESQYLVP GYDRWWACNT GLTPCVSTLV
CIGRVPPSHQ HLCNHTEAFN RTSESQYLVP GYDRWWACNT GLTPCVSTLV
CIGRVPPSHQ HLCNHTEAFN RTSESQYLVP GYDRWWACNT GLTPCVSTLV
CIGRVPPSHQ HLCNHTEAFN RTSESQYLVP GYDRWWACNT GLTPCVSTLV
CIGRVPPSHQ HLCNHTEAFN RTSESQYLVP GYDRWWACNT GLTPCVSTLV
CIGRVPPSHQ HLCNHTEAFN RTSESQYLVP GYDRWWACNT GLTPCVSTLV
CIGMVPPSHQ HLCNHTEAFN RTSESQYLVP GYDRWWACNT GLTPCVSTLV
CIGMVPPSHQ HLCNHTEAFN QTSESQYLVP GYDRWWTCNT GLTPCVSTLV
CIGRVPPSHQ HLCNHTEAFN -TSESQYLVP GYDRWWACNT GLTPCVSTLV
CIGRVPPSHQ HLCNHTEAFN RTSESQ--VS GT---WLQVV GMTPCVSTLV
CIGRVPPSHQ HLCNHTEAFN RTSESQ--VT GT---WLQVV GMTPCFSTLV
CIGRVPPSHQ HLCNHTEAFN RTSKSQVSGT RLDRWWTCNT GLTPCVSTLV
CIGRVPPSHQ HLCNHTEAFN RTSENQYLVP GYDRWWACNT GLTPCVSTLV
YIGRVPPSHQ HLCNHTEAFN RTSENQYLVP GYDRWWACNT GLTPCVSTLV
YIGRVPPSHQ HLCNHTEAFN RTSENQYLVP GYDRWWACNT GLTPCVSTLV
YIGRVPPSHQ HLCNHTEAFN RTSENQYLVP GYDRWWACNT GLTPCVSTLV
YIGRVPPSHQ HLCNHTEAFN RTSENQYLVP GYDRWWACNT GLTPCVSTLV
YIGRVPPSHQ HLCNHTEAFN RTSESQYLVP GYDRWWACNT GLTPCVSTLV
YIGRVPPSHQ HLCNHTEAFN RTSESQYLVP GYDRWWACNT GLTPCVSTLV
YIGMVPPSHQ HLCNHTEAFN RTSESQYLVP GYDRWWACNT GLTPCVSTLV
YIGRVPPSHQ HLCNHTEAFN RTSENQYLVP GYDRWWACNT GLTPCVSTLV
CIGKVPPSHQ HLCNHTEAFN QTSESQYLVP GYDRWWACNT GLTPCVSTLV
```

CIGRVPPSHQ HLCNHTEAFN RTSESQYLVP GYDRWWACNT GLTPCVSTLV
XIGRVPPSHQ HLCNHTEAFN RTSENQYLVP GYDRWWACNT GLTPCVSTLV
CIGKVPPSHQ HLCNHTEAFN QTSESQYLVP GYDRWWACNT GLTPCVSTLV
CIGKVPPSHQ HLCNHTEAFN QTSESQYLVP GYDRWWACNT GLTPCVSTLV
CIGKVPPSHQ HLCNHTEAFN QTSESQYLVP GYDRWWACNT GLTPCVSTLV
CIGKVPPSHQ HLCNHTEAFN QTSESQYLVP GYDRWWACNT GLTPCVSTLV
CIGKVPPSHQ HLCNHTEAFN QTSESQYLVP GYDRWWACNT GLTPCVSTLV
CIGKVPPSHQ HLCNHTEAFN QTSESQYLVP GYDRWWACNT GLTPCVSTLV
CIGKVPPSHQ HLCNHTEAFN QTSESQYLVP GYDRWWACNT GLTPCVSTLV
CIGKVPPSHQ HLCNHTEAFN QTSESQYLVP GYDRWWACNT GLTPCVSTLV
CIGRVPPSHQ HLCNHTEAFN RTSESQYLVP GYDRWWACNT GLTPCVSTLV
CIGKVPPSHQ HLCNHTEAFN QTSESQYLVP GYDRWWACNT GLTPCVSTLV
YIGRVPPSHQ HLCNHTEAFN RTSESQYLVP GYDRWWACNT GLTPCVSTLV
YIGRVPPSHQ HLCNHTEAFN RTSESQYLVP GYDRWWACNT GLTPCVSTLV
YVGRVPPSHQ HLCNHTEAFN RTSESQYLVP GYDRWWACNT GLTPCVSTLV
CIGRVPPSHQ HLYNHTEAFN RTSESQYLVP GYDRWWACNT GLTPCVSTLV
CIGRVPPSHQ HLCNHTEAFN RTSESQYLVP GYDRWWACNT RLTPCVSTLV
YIGRVPPSHQ HLCNHTEAFN RTSESQYLVP GYDRWWACNT RLTPCVSTLV
CIKKVPPSHQ HLYNHTEAFN QTSESQYLVP GYDRWWACNT GLTPCVSTLV
CIKKVPPSHQ HLYNHTEAFN QTSESQYLVP GYDRWWACNT GLTPCVSTLV
CIGKAPPSHQ HLCYSTVVYE QASENQYLVP GYNRWWACNT GLTPCVSTSV
CIGKAPPSHQ HLCYSTVVYE QASENQYLVP GYNRWWACNT GLTPCVSTSV
CIGKAPPSHQ HLCYSTVVYE QASENQYLVP GYNRWWACNT GLTPCVSTSV
CIGKAPPSHQ HLCYSTVVYE QASENQYLVP GYNRWWACNT GLTPCVSTSV
CIGKAPPSHQ HLCYSTVVYE QASENQYLVP GYNRWWACNT GLTPCVSTSV
CIGKAPPSHQ HLCYSTVVYE QASENQYLVP GYNRWWACNT GLTPCVSTSV
CIGKAPPSHQ HLCYSTVVYE QASENQYLVP GYNRWWACNT GLTPCVSTSV
CIGKAPPSHQ HLCYSTVVYE QASENQYLVP GYNRWWACNT GLTPCVSTSV
CIGKAPPSHQ HLCYSTVVYE QASENQYLVP GYNRWWACNT GLTPCVSTSV
CIGKAPPSHQ HLCYSTVVYE QASENQYLVP GYNRWWACNT GLTPCVSTSV
CIGKAPPSHQ HLCYSTVVYE QASENQYLVP GCNRWWACNT GLTPCVSTSV
CIGKAPPSHQ HLCYSTVVYE QASENQYLVP GYNRWWACNT GLTPCVSTSV
CIGKAPPSHQ HLCYSTVVYE QASENQYLVP GYNRWWACNT GLTPCVSTSV
CIGKAPPSHQ HLCYSTVVYE QASENQYLVP GYNRWWACNT GLTPCVSTSV
CIGKAPPSHQ HLCYSTVVYE QASENQYLVP GYNRWWACNT GLTPCVSTSV
CIGKAPPSHQ HLCYSTVVYE QASENQYLVP GYNRWWACNT GLTPCVSTSV
CIGKAPPSHQ HLCYSTVVYE QASENQYLVP GYNRWWACNT GLTPCVSTSV
CIGKAPPSHQ HLCYSTVVYE QASENQYLVP GYNRWWAYNT GLTPCVSTSV
CIGKAPPSHQ HLCNSTMVYE QASENQYLVP GYNRWWACNT GLTPCVSTTV
CIGKAPPSHQ HLCNSTMVYE QASENQYLVP GYNRWWACNT GLTPCVSTTV
CIGKAPPSHQ HLCNSTMVYE QASENQYLVP GYNRWWACNT GLTPCVSTTV
CIGKAPPSHQ HLCNSTMVYE QASENQYLVP GYNRWWACNT GLTPCVSTTV
CIGKAPPSHQ HLCNSTMVYE QASENQYLVP GYNRWWACNT GLTPCVSTTV
CIGKAPPSHQ HLCNSTMVYE QASENQYLVP GYNRWWACNT GLTPCVSTTV

CIGKAPPSHQ HLCNSTMVYE QASENQYLVP GYNRWWACNT GLTPCVSTTV

CIGKAPPSHQ HLCNSTMVYE QASENQYLVP GYNRWWACNT GLTPCVSTTV

CIGKAPPSHQ HLCNSTMVYE QASENQYLVP GYNRWWACNT GLTPCVSTTV

CIGKAPPSHQ HLCNSTMVYE QALENQYLVP GYN-WWACNT GLTPCVSTTV

CIGKAPPSHQ HLCNSTMAYE QALENQYLVP GYN-WWACNT GLTPCVSTTV

CIGKAPPSHQ HLCNSTMVYE QALENQYLVP GYN-WWACNT GLTPCVSTTV

CIGKAPPSHQ HLCNSTMVYE QASENQYLVP GYN-WWACNT GLTPCVSTTV

CIGKAPPSHQ HLCNSTMVYK QASENQYLVP GYNRWWACNT GLTPRVSTTV

CIGKAPPSHQ HLCYSTVVYE QASENQYLVP GYNRWWACNT GLTPCVSTSV

CIGKAPPSHQ HLCNSTMVYK QASENQYLVP GYNRWWACNT GLTPCVSTTV

CIGKAPPSHQ HLCNSTMVYE QASENQYLVP GYNRWWACNT GLTPCVSTTV

CIGKAPPSHQ HLCNSTMVYK QASENQYLVP GYNRWWACNT GLTPRVSTTV

CIGKAPPSHQ HLCNSTMVYK QASENQYLVP GYNRWWACNT GLTPRVSTTV

CIGKAPPSHQ HLCNSTMVYK QASENQYLVP GYNRWWACNT GLTPRVSTTV

CIGKAPPSHQ HLCNSTMVYK QASENQYLVP GYNRWWACNT GLTPRVSTTV

CIGKAPPSHQ HLCNSTMVYK QASENQYLVP GYNRWWACNT GLTPRVSTTV

CIGKAPPSHQ HLCNSTMVYK QASENQYLVP GYNRWWACNT GLTPRVSTTV

CIGKAPPSHQ HLCNSTMVYE QASENQYLVP GYNRWWACNT GLTPCVSTSV

CIGKAPPSHQ HLCIVLWFYE QASENQYLVP GYNRWWACNT GLTPCVSTSV

CIGKAPPSHQ HLCNSTMVYK QASENQYLVP GYNRWWACNT GLTPCVSTTV

CIGKAPPSHQ HLCYSTVVYE QASENQYLVP GYNRWWACNT GLTPCVSTTV

YIGRVPPSHQ HLCNHTEAFN RTSENQYLVP GYDRWWACNT GLSPCVSTLV

YIGRVPPSHQ HLCNHTEAFN RTSENQYLVP GYDRWWACNT GLSPCVSTLV

CIEKPPTSHQ HLCNITLTYN QTSDNQYLVP GYNRWWACDT GLTPCVSTSV

CI-K-PPSHQ HLCNVTLTYS QTSDNQYLVP EYNRWWACDT GLTPCVSTSV

CIGSAPPSHQ HLCNNTLIYN QTSDNQYLMP GYNRWWACDT GLTPCVSPSV

CIGKVPPTHQ HLCNLTIPLN ASHTHKYLLP SNHSWWACNS GLTPCLSTSV

CIGKVPFTHQ HLCNQTLSIN SSGDHQYLLP SNHSWWACST GLTPCLSTSV

CIGKVPLSHR HLCNVTYESP RTKLTYYLSP GPNKWWACST GLTPCISTSV

CIGKVPLSHR HLCNVTYESP RTKLTYYLSP GPNKWWACST GLTPCISTSV

CIGKVPLSHR HLCNVTYESP RTKLTYYLSP GPNKWWACST GLTPCVSTSV

CVGKVSPSHH HLCNETHKSP KTELTYYLSP GPNKWWACST VLTPCVSTSV

CIGKVPPSHR HLCNETHKSP KTELTYYLLP GPNKWWACST VLTPCVSTSV

CLGNPPAAY- HLCNTTLESP STSDIEYLFP GLDK-WACST GLL-VYLLQC

WLGNPLAAYR HLCNTTLESP STSDIEYLFP GLDKWWACSI GLTPCVSTLV

WLGNPLAAYR HLCNTTLESP STSDIEYLFP GLDKWWACST GLTPCVSTLV

CLGKPPAAYR HLCNTTLESP STSDIEYLFP GLDKGWACGT GLTPCVSTLV

CLGNPPAAYK CLCNETLKSP KTSDTRYLFP SLDKWWACST GLTPCVSTSV

CLGNPPAAYK CLCNETLRSP KTSDTRYLFP SLDK-WACSA GLTPCVLTSV

CLGQVPQDKG HLCNQTQNIQ SSKSGQYLVP PLDTVWACNT GLTPCVSMSV

CLGQVPQDKG HLCNQTQNIQ SSKSGQYLVP PLDTVWACNT GLTPCVSMSV

CLGQVPQDKG HLCNQTQNIQ SSKSGQYLVP PLDTVWACNT GLTPCVSMSV

CLGRVPQDKG HLCNQTQNIQ SSKSGQYLVP PLDTVWACNT GLTPCVSMSV

CLGQVPQDKW HLCNQTQNIR PNKGGQYLVP PIDTVWACNT GLTPCISMSV

```
CIGRVPAEYQ PLCNKTIDTL QMSQTS-TCP PPDTWWACGT SLTPCLHISI
CVGTPPQSHR HLCHD---FS LPSGSGYLVP PQDGWWACNT GLTPCVSLEV
CLGNPPAAYR HLCNEILRRG GTSTNRFLVP TLDKWWACGS GLNPCISTSV
CIGTPPSTHK HLCGQIQSVS RTEADYYLVP SPVGWWACNT GLTPCVSTKV
CIGTPPSTHK HLCGQIQSVS RTEADYYLVP SPVGWWACNT GLTPCVSTKV
CIGTPPSTHK HLCGQIQSVS RTEANYYLVP SPVGWWACNT GLTPCVSTKV
CVGAVPKTHQ ALCNTTQKTS --DGSYYLAA PAGTIWACNT GLTPCLSTTV
CVGAVPKTHQ ALCNTTQKTS --DGSYYLAA PAGTIWACNT GLTPCLSTTV
CVGAVPKTHQ ALCNTTQKAS --DGSYYLAA PAGTIWACNT GLTPCLSTTV
CVGAVPKTHQ ALCNTTQNTS --DGSYYLAA PAGTIWACNT GLTPCLSTTV
CIGTVPKTHQ ALCNTTLKTN --KGSYYLVA PAGTTWACNT GLTPCLSATV
CIGTVPKTHQ ALCNTTLKAG --KGSYYLVA PTGTMWACNT GLTPCLSATV
CIGAVPKTHQ ALCNTTQTSS --RGSYYLVA PTGTMWACST GLTPCISTTI
CIGTVPKTHQ ALCNETQQGH --TGAHYLAA PNGAYWACNT GLTPCISMAV
CVGKVPSTHS HLCRH----- ---------- ----LWACSS GLTPCLADSV
CLGNVPPSHR YLCNQTSLSL TTT------- ---QWWACNT GLTPCVATRV
CLGKVPQAKQ SLCASIDSSP SKSDTKWLIP RTDGWWICST GLTPCLSTSV
YAGEIPVGYV HFTNCTSIQE VSNETTRLCP PPGHVFVCGN NMA---YTAL
CLGNVPADKK HLCNQTLTSP HTSTTYYLVP PLEGWWACST GLTPCVATSV
CLGNVPADKK QLCNQTLTSP HTSTTYYLVP PLEGWWAYST RLTPCVATSV
CLGNVPTSRK HLCNQTLASP HTSTTYYLVP PLEGWWACST GLTPCVATSV
CLGNVPTSRK HLCNQTLASP HTSTTYYLVP PLEGWWACST GLTPCVATSV
CLGNVPTSRK HLCNQTLASP HTSTTYYLVP PLEGWWACST GLTPCVATSV
CLGNVPTSRE HLCNQTLASP HTSTTYYLVP PLEGWWACST GLTPCVATSV
CLGNVLTSRK HLCNQTLASP HSSTSYYLVP PLEGWWACST GLTPCVATSV

FNQTKDFCVM VQIVPRVYYY PEKAVLDEYD YRYNQPKREP ISLTLAVMLG
FNQTKDFCVM VQIVPRVYYY PEKAVLDEYD YRYNRPKREP ISLTLAVMLG
FNQTKDFCVM VQIVPRVYYY PEKAVLDEYD YRYNRPKREP ISLTLAVMLG
FNQTKDFCVM VQIVPRVYYY PEKAVLDEYD YRYNRPKREP ISLTLAVMLG
FNQTKDFCVM VQIVPRVYYY PEKAVLDEYD YRYNRPKREP ISLTLAVMLG
FNQTKDFCVM VQIVPRVYYY PEKAVLDEYD YRYNRPKREP ISLTLAVMLG
FNQTKDFCVM VQIVPRVYYY PEKAVLDEYD YRYNRPKREP ISLTLAVMLG
FNQTKDFCVM VQIVPRVYYY PEKAVLDEYD YRYNRPKREP ISLTLAVMLG
FNQTKDFCVM VQIVPRVYYY PEKAVLDEYD YRYNRPKREP ISLTLAVMLG
FNQTKDFCVM VQIVPRVYYY PEKAVLDEYD YRYNRPKREP ISLTLAVMLG
FNQTKDFCVM VQIVPRVYYY PEKAVLDEYD YRYNRPKREP ISLTLAVMLG
FNQTKDFCFM VQIVPRVYYY PEKAVLDEYD YRYNRPKREP ISLTLAVMLG
FNQTKDFCVM VQIVPRVYYY PEKAVLDEYD YRYNRPKREP ISLTLAVMLG
FNQTKDFCVM VQIVPRVYYY PEKAVLDEYD YRYNRPKREP ISLTLAVMLG
FNQTKDFCVM VQIVPRVYYY PEKAVLDEYD YRYNWPKREP ISLTLAVMLG
FNQTKDFCVM VQIVPRVYYY PEKAVLDEYN YRYNRPKREP ISLTLAVMLG
FNQTKDFCVM VQIVPRVYYY PEKAVLDEYD YRYNRPKREP ISLTLAVMLG
FNQTKDFCVM VQIVPRVYYY PEKAVLDEYD YRYNRPKREP ISLTLAVMLG
```

FNQTKDFCVM VQIVPRVYYY PEKAVLDEYD YRYNRPKREP ISLTLAVMLG
FNQTKDFCVM VQIVPRVYYY PEKAVLNKYD YRYNRPKREP ISLTLAVMLG
FNQTKDFCVM VQIVPRVYYY PEKTVLDKYD YRYNRPKKEP ISLTLAVMLG
FNQTKDFCFM VQIVPRVYYY PKKAVLDEYD YRYNRPKREP ISLTLAVMLG
FNQTKDFCVM VQIVPRVYYY PKKAVLDEYG YRYNRPKREP ISLTLAVMLG
FNQTKDFCVM VQIVPRVYYY PKKAVLDEYG YRYNRPKREP ISLTLAVMLG
FNQTKDFCVM VQIVPRVYYY PEKTVLDKYD YRYNRPKREP ISLTLAVMLG
FDQTKGFCVM VQIVPRVYYH PEKAALDKYD YKYNRPKRKP ISLTLAAMLG
FDQTKGFCVM VQIVPRVYYH PEKAALDKYD YKYNRPKRKP ISLTLAAMLG
FDQTKGFCVM VQIVPRVYYH PEKAALDKYD YKYNRPKRKP ISLTLAAMLG
FDQTKGFCVM VQIVPRVYYH PEKAALDKYD YKYNRPKRKP ISLTLAAMLG
FDQTKGFCVM VQIVPRVYYH PEKAALDKYD YKYNRPKRKP ISLTLAAMLG
FDQTKGFCVM VQIVPRVYYH PEKAALDKYD YKYNRPKRKP ISLTLAAMLG
FDQTKGFCVM VQIVPRVYYH PEKAALDKYD YKYNRPKRKP ISLTLAAMLG
FDQTKGFCVM VQIVPRVYYH PEKAALDKYD YKYNRPKRKP ISLTLAAMLG
FDQTKGFCVM VQIVPRVYYH PEKAALDKYD YKYNRPKRKP ISLTLAAMLG
FNQTKDFCVM VQIVPRVYYH PEKAALDKYD YKYNRPKRKP ISLTLAAMLG
FDQTKGFCVM VQIVPRVYYH PEKAVLDKYD YKYNRPKRKP ISLTLAAMLG
FDQTKGFCVM VQIVPRVYYH PEKAALDKYD YKYNRPKRKP ISLTLAAMLG
FNQTKDFCIM VQIVPRVYYY PEKAILDEYD YRNHRQKREP ISLTLAVMLG
FNQTKDFCIM VQIVPRVYYY PEKAILDEYD YRNHRQKREP ISLTLAVMLG
FNQTKDFCIM VQIVPRVYYY PEKAILDEYD YRNHRQKREP ISLTLAVMLG
FNQTKDFCIM VQIVPRVYYY PEKAILDEYD YRNHRQKREP ISLTLAVMLG
FNQTKDFCIM VQIVPRVYYY PEKAILDEYD YRNHRQKREP ISLTLAVMLG
FNQTKDFCIM VQIVPRVYYY PEKAILDEYD YRNHRQKREP ISLTLAVMLG
FNQTKDFCIM VQIVPRVYYY PEKAILDEYD YRNHRQKREP ISLTLAVMLG
FNQTKDFCIM VQIVPRVYYY PEKAILDEYD YRNHRQKREP ISLTLAVMLG
FNQTKDFCIM VQIVPRVYYY PEKAILDEYD YRNHRQKREP ISLTLAVMLG
FNQTKDFCIM VQIVPRVYYY PEKAILDEYD YRNHRQKREP ISLTLAVMLG
FNQTKDFCVM VQIVPRVYYY PEKTVLDEYD YKYHRQKREP ISLTLAVMLG
FNQTKDFCVM VQIVPRVYYY PEKTVLDEYD YKYHRQKREP ISLTLAVMLG
FNETKDFCVM VQIVPRVYYY PEKTVLDEYD YKYHRQKREP ISLTLAVMLG
FNKTKDFCVM VQIVPRVYYY PEKTVLDEYD YRYHRQKREP ISLTLTVMLG
FNQTKDFYIV VQIVPRVYYY PEKTILDEYN YRNHRQKRKP ISLTLAVMLG
FNQTKDFYIV VQIVPRVYYY PEKTILDEYN YRNHRQKRKP ISLTLAVMLG
FNQTKDFYIM VQIVPRVYYY PKKTILDEYD YRNHRQKKKP ISLTLAVMLG
FNQTKDFYIM VQIVPRVYYY PKKTILDEYD YRNHRQKKKP ISLTLAVMLG
FNQSKDFCVM VQIVPRVYYH PEEVVLDEYD YRYNRPKREP VSLTLAVMLG
FNQSKDFCVM VQIVPRVYYH PEEVVLDEYD YRYNRPKREP VSLTLAVMLG
FNQSKDFCVM VQIVPRVYYH PEEVVLDEYD YRYNRPKREP VSLTLAVMLG
FNQSKDFCVM VQIVPRVYYH PEEVVLDEYD YRYNRPKREP VSLTLAVMLG
FNQSKDFCVM VQIVPRVYYH PEEVVLDEYD YRYNRPKREP VSLTLAVMLG
FNQSKDFCVM VQIIPRVYYH PEEVVLDEYD YRYNRPKREP VSLTLAVMLG
FNQSKDFCVM VQIVPRVYYH PEEVVLDEYD YRYNRPKREP VSLTLAVMLG

```
FNQSKDFCVM VQIVPRVYYH PEEVVLDEYD YRYNRPKREP VSLTLAVMLG
FNQSKDFCVM VQIVPRVYYH PEEVVLDEYD YRYNRPKREP VSLTLAVMLG
FNQSKDFCVM VQIVPRVYYH PEEVVLDEYD YRYNRPKREP VSLTLAVMLG
FNQSKDFCVM VQIVPRVYYH PEEVVLDEYD YRYNRPKREP VSLTLAVMLG
FNQSKNFCVM VQIVPRVYYH PEKVVLDEYD YRYNRPKREP VSLTLAVMLG
FNQSKDFCVM VQIVPRVYYH PEEVVLDEYD YRYNQPKREP ISLTLAVMLG
FNQSKDFCVM VQIIPRVYYH PEEVVLDEYD YRYNRPKREP VSLTLAVMLG
FNQSKDFCVM VQIVPRVYYH PEEVVLDEYD YRYNRPKREP VSLTLAVMLG
FNQSKDFCVM VQIIPRVYYH PEEVVLDEYD YRYNRPKREP VSLTLAVMLG
FNQSKDFCVM VQIIPRVYYH PEEVVLDEYD YRYNRPKREP VSLTLAVMLG
FNQSKDFCVM VQIVPRVYYH PEKVVLDEYD YRYNRPKREP VSLTLAVMLG
FNQSKDFCVM VQLIPQVYYH PEEVVIDEYD YRPTRSKREP VTLTLAIILG
FNQSKDFCVM VQLIPQVYYH PEEVVIDEYD YRPTRSKREP VTLTLAIILG
FNQSKDFCVM VQLIPQVYYH PEEVVIDEYD YRPTRSKREP VTLTLAIILG
FNQSKDFCVM VQLIPQVYYH PEEVVIDEYD YRPTRSKREP VTLTLAIILG
FNQSKDFCVM VQLIPQVYYH PEEVVIDEYD YRPTRSKREP VTLTLAIILG
FNQSKDFCVM VQLIPQVYYH PEEVVIDEYD YRPTRSKREP VTLTLAIILG
FNQSKDFCVM VQLIPQVYYH PEEVVIDEYD YRPTRSKREP VTLTLAIILG
FNQSKDFCVM VQLVPRVYYH PEEVVIDEYD YRPTRSKREP VTLTLAVILG
FNQSKDFCVM VQLVPRVYYH PEEVVIDEYD YRPTRSKREP VTLTLAVILG
FNQSKDFCVM VQLVPWVHYH PEEVVIDEYD YRPTRSKREP VTL-------
FNQSKDFCVM VQLVPWVHYH PEEVVIDEYD YRPTRSKREP VTL-------
FNQSKDFCVM VQLVPWVHYH PEEVVIDEYD YRPTRSKREP VTL-------
FNQSKDFCVM VQLVPWVHYH PEEVVIDEYD YRPTRSKREP VTL-------
FNQSKDFCVM VQ-------- PEEVVIDKYD HRPTRSKREP MSLTLAVMLG
FNQSKDFCVM VQ-------- PEEVVIDKYD HRPTRSKREP MSLTLAVMLG
FNQSKDFCVM VQLVPRVYYH PEEVVIDEYD HRPTRSKREP VSLTLAVMLG
FNQSKDFCVM VQLVPRVYYH PEEVVIDEYD HRPTRSKREP VSLTLAVMLG
FNQSKDFCVM VQLVPRVYYH PEEVVIDEYD HRPTRSKREP VSLTLAVMLG
FNQSKDFCVM VQLVPRVYYH PEEVVIDEYD HRPTRSKREP VSLTLAVMLG
FNQSKDFCVM VQLVPRVYYH PEEVVIDEYD HRPTRSKREP VSLTLAVMLG
FNQSKDFCVM VQLVPRVYYH PEEVVIDEYD HRPTRSKREP VSLTLAVMLG
FKQSKDFCVM VQLVPRVYYH PEEVVIDEYD HRPTRSKREP VSLTLAVMLG
FNQSKDFFVM VQLVPRLYYH PEEVVIDEYD HRPTRSKREP VSLTLAVMLG
FNQSKDFCVM VQLVPRVYYH PEEVVIDEYD HRPTRSKREP VSLTLAVMLG
FNQSKDFCVM VQLVPRVYYH PEEVVIDEYD HRPTRSKREP VSLTLAVMLG
FNQSKDFCVM VQLVPRVYYH PEEVVIDXYD HRPTRSKREP VSLTLAVMLG
FNQSKDFCVM VQ-------- PEEVVIDKYD HRPTRSKREP MSLTLAVMLG
FDQTKDFCVM VQIVPWVYYH PEKAVLDEYD YRYNRPKREH ISLTLAAMLR
FDQTKDFCVM VQIVPWVYYH PEKAVLDEYD YRYNRPKREH ISLTLAAMLR
FNPSKNFCIM VQLVPRVYYH PEEVVIDEYD YHSPRSKREP ------VMLG
FNPSKNFCIM VQLVPWVYYH PEEVVIDEYD YHPTRSKREP VTLTLAVILG
FKQSKNLYIM VRLVPQVYYH PEEVVVDEYD Y-----KREP VTLTLAVMLG
FNQSNDFCIQ IQLVPRIYYH PDGTLLQAYE SPHPRNKREP VSLTLAVLLG
```

FNQTRDFCIQ VQLIPRIYYY PEEVLLQAYD NSHPRTKREA VSLTLAVLLG
FNDIRDYCIL VQLVPRVIYH PSETFVDEFD RRLTRLRREP VTMTLAVLLG
FNDIRDYCIL VQLVPRVIYH PSETFVDEFD RRLTRLRREP VTMTLAVLLG
FNDIRDYCIL VQLVPRVIYH PSETFVDEFD RRLTRLRRKP VTMTLAVLLG
FNETRDYCIL VQLVPRVIYH PTENFVDEFD KRPTHLHREP VTMTLAMFGG
FNESRDYCIL VQLVPRVIYH PTENFVDEFD KLPTRLPREP VTMTLAMLLG
LKNTKDYCVL IQLVPRVLFH PTNCFVDKFV RRLTCYRGEP IKLTLAVILG
FDNTKDYCGL IQLVPRVFYH PTNSFVDEFV GRLTRYWKEP VTLTLAVILG
FDNTKDYCVL IQLVPRVFYH PTNSFVDEFV GRLTRYWKEP VTLTLAVILG
FDNTKDYCVL IQSVPRVFHY STNSFVDEFD GRLTRYRKEP VTLTLAVILG
IDDTKDYCVL VQLVPRVFYH PANMLEDEFD KHPTRFWREP ISLTLAVILG
FDDTKDYCVL IQLLSSSFYH P-----DEFD KHPTRFQREP ISLTLAVILG
FNSSKDFCIL VQLIPRLLYH DDSSFLDKFE HR-VRWRREP VTLTLAVLLG
FNSSKDFCIL VQLIPRLLYH DDSSFLDKFE HR-VRWRREP VTLTLAVLLG
FNSSKDFCIL VQLIPRLLYH DDSSFLDKFE HR-VRWKREP VTLTLAVLLG
FNSSKDFCIL VQLIPRLLYH DDSSFLDKFE HR-VRWKREP VTLTLAVLLG
FNSSKDFCIL VQLIPRLLYH DDSSFLDKFE HR-VRWKREP ITLTLAVLLG
FDINRDFCIL IQLVLRVIYH DDKSFIDEFD HR-TRYKREP VTLTLAVLLG
LNTSADFCVL VQLVPRLIYH SDPSFLDEYE GR----KKKK ITLTLAMLLG
-------SIL VQLVPRVYYH PAGTLEDEFD KHLRCFRREP ISLTLAVMLG
FNSSHDFCVM IQLLPRVYYH PASSLEESYA GR--RSKREP ITLTLAAFMG
FNSSHDFCVM IQLLPRVYYH PASSLEESYA GR--RSKREP ITLTLAAFMG
FNSSHDFCVM IQLLPRIYYH PASSLEESYA GR--RSKREP ITLTLAAFME
LDLTTDYCVL VELWPKVTYH SPGYVYGQFE RK-TKYKREP VSLTLALLLG
LDLTTDYCVL VELWPKVTYH SPGYVYGQFE RK-TKYKREP VSLTLALLLG
LNLTTDYCVL VELWPKVTYH SPGYVYDQFE RK-TKYKREP VSLTLALLLG
LNLTTDYCVL VELWPKVTYH SPGYVYGQFE RK-TKYKREP VSLTLALLLG
LNRTTDYCVL VELWPRVTYH PPSYVYSQFE KS-YRHKREP VSLTLALLLG
LNRTTDYCVL VELWPRVTYH PPSYVYSQFE KS-HRHKREP VSLTLALLLG
LNLTTDYCVL VELWPRVTYH SPSYVYGLFE RS-NRHKREP VSLTLALLLG
LNWTSDFCVL IELWPRVTYH QPEYVYTHFA KA-VRFRREP ISLTVALMLG
INNISDYCVM VQLVPRIYYH SPGEFLSLYE PT-PRIKREP ISLTVAVLLG
FNNSRNYCIL IQLLPKIIYH DGPGFKDLLD WRVTRYRREP ISLTLAMLLG
FNAANEFCVL VTVLPRILYH PEESMYSHWD SDSTRSKREP ITLTIATLFS
PNKWIGLCIL ASIVPSIISG EEPIPLPSIE YTAGRHKRAV QFIPLPV--G
FNNSKDFCIL IQLVPRIIYH DSKSFEDQFD WRIYRVRREP VSMALAILLG
FNNSKDFCIL IQLVPRIIYH DSKSLEDKFD SRIYRVRREP VSVTLAILLG
FNSSRDFCIL VQLVPRVIYH DSKSFEAQFD SKIHRTRREP VSMTLAVLLG
FNSSKDFCIL VQLVPRVIYH SKINLIPKYD SKIHRTQREP VSMTLAVLLG
FNNSKDFCIL VQLVPRVIYH DSKSFEDQFD SKIHRIQRDP VSMTLAVLLG
FNSSRDFCIL VQLVPRVIYH DSKSFEDQFD PKIHRTRREP VSMTLAVLLG
FNSSKDFCIL VQLVPRVIFH DSKSFEDQFD SKIHRTRREP VSMTLAVLLG

LGVAAGVGTG TAALITGPQQ LEDLHRIVTE DLQALEKSVS NLEESLTSLS

LGVAAGVGTG TAALITGPQQ LENLHRIVTE DLQALEKSVS NLEESLTSLS
LGVAAGVGTG TAALITGPQQ LENLHRIVTE DLQALEKSVS NLEESLTSLS
LGVAAGVGTG TAALITGPQQ LENLHRIVTE DLQALEKSVS NLEESLTSLS
LGMAAGVGTG TAALITGPQQ LENLHRIVTE DLQALEKSVS NLEESLTSLS
LGVAAGVGTG TAALITGPQQ LENLHRIVTE DLQALEKSVS NLEESLTSLS
LGVAAGVGTG TAALITGPQQ LENLHRIVTE DLQALEKSVS NLEESLTSLS
LGVAAGVGTG TAALITGPQQ LENLHRIVTE DLQALEKSVS NLEESLTSLS
LGVAAGVGTG TAALITGPQQ LENLHRIVTE DLQALEKSVS NLEESLTSLS
LGVAAGVGTG TAALITGPQQ LENLHRIVTE DLQALEKSVS NLEESLTSLS
LGVAAGVGTG TAALITGPQQ LENLHRIVTE DLQALEKSVS NLEESLTSLS
LGVAAGVGTG TAALITGPQQ LENLHRIVTE DLQALEKSVS NLEESLTSLS
LGMAAGVGTG TAALITGPQQ LENLHRIVTE DLQALEKSVS NLEESLTSLS
LGVAAGVGTG TAALITGPQQ LEDLHRIVTE DLQALEKSVS NLEESLTSLS
LGVAAGVGTG TAALITGPQQ LENLHRIVTE DLQALEKSVS NLEESLTSLS
LGMATGVGTG TAALITGPQQ LENLHRIVTE DLQALEKSVS NLEESLTSLS
LGVATGVGTG TAALITGPQQ LENLHRIVTE DLQALEKSVS NLEESLTSLS
LGVAAGVGTG TAALITGPQQ LEDLHRIVTE DLQALEKSVS NLEESLTSLS
LGVAAGVGTG TAALITGPQQ LEDLHRIVTE DLQALEKSVS NLEESLTSLS
LGVAAGVGTG TAALITGPQQ LENLHRIVTE DLQALKKSVS NLEESLTSLS
LGVAAGVGTG TAALITGPQQ LENLHRIVTE DLQALEKSVS NLKKSLTSLS
LGVAAGMGTG TAALITGPQQ LEDLHRIVME DLQALEKSVS NLEESLTSLS
LGVAAGVGTG MAALITGPQQ LEDLHRIVME DL-ALEKSVC NQEESLTSLS
LGVAAGVGTG TAALITGPQQ LEDLHRIVTK DLQALKQSVS NLEESLTSLS
LGVAAGVGTG TAALITGPQQ LEDLHRIVTK DLQALKQSVS NLEESLTSLS
LGVAAGVGTG TAALITGPRQ LEELHRIITE DLQALEKSVS NLEEFLTSLS
LGVAAGVGTG TAALITGPRQ LEELHRIITE DLQALEKSVS NLEEFLTSLS
LGVAAGVGTG TAALITGPRQ LEELHRIITE DLQALEKSVS NLEEFLTSLS
LGVAAGVGTG TAALITGPRQ LEELHRIITE DLQALEKSVS NLEEFLTSLS
LGVAAGVGTG TAALITGPRQ LEELHRIITE DLQALEKSVS NLEEFLTSLS
LGVAAGVGTG TAALITGPRQ LEELHRIITE DLQALEKSVS NLEEFLTSLS
LGVAAGVGTG TAALITGPRQ LEELHRIITE DLQALEKSVS NLEEFLTSLS
LGVAAGVGTG TAALITGPRQ LEELHRIITE DLQALEKSVS NLEEFLTSLS
LGVAAGVGTG TAALITGPRQ LEELHRIITE DLQALEKSVS NLEEFLTSLS
LGVAAGVGTG TAALITGPRQ LEELHRIITE DLQALEKSVS NLEEFLTSLS
LGVAAGVGTG TAALITGPRQ LEELHRIITE DLQALEKSVS NLEEFLTSLS
LGVAAGVGTG TAALXTGPRQ LEELHRIITE DLQALEKSVS NLEEFLTSLS
LGVAAGVGTG TAALVTGPQQ LENLHRIVTE DLQALEKSVS NLEESLTSLS
LGVAAGVGTG TAALVTGPQQ LENLHRIVTE DLQALEKSVS NLEESLTSLS
LGVAAGVGTG TAALVTGPQQ LENLHRIVTE DLQALEKSVS NLEESLTSLS
LGVAAGVGTG TAALVTGPQQ LENLHRIVTE DLQALEKSVS NLEESLTSLS
LGVAAGVGTG TAALVTGPQQ LENLHRIVTE DLQALEKSVS NLEESLTSLS
LGVAAGVGTG TAALVTGPQQ LENLHRIVTE DLQALEKSVS NLEESLTSLS
LGVAAGVGTG TAALVTGPQQ LENLHRIVTE DLQALEKSVS NLEESLTSLS
LGMVAGVGTG TAALVTGPQQ LENLHRIVTE DLQALEKSVS NLEESLTSLS

```
LGMVAGVGTG TAALVTGPQQ LENLHRIVTE DLQALEKSVS NLEESLTSLS
LGMVAGVGTG TAALVTGPQQ LENLHRIVTE DLQALEKSVS NLEESLTSLS
LGVAAGVGTG TAALVTGPQQ LENLHRIVTE DLQALEKSVS NLEESLTSLS
LGVAAGVGTG TAALVTGPQQ LENLHRIVTE DLQALEKSVS NLEESLTSLS
LGVAAGVGTG TAALVTGPQQ LENLHRIVTE DLQALEKSVS NLEESLTSLS
LGVAAGVGTG TAALVTGPQQ LENLHRIVTE DLQALEKSVS NLEESLTSLS
LGVAAGVGTG TAALVTGPQQ LENLHQIVTE DLQALEKSVS NLEESLTSLS
LGVAAGVGTG TAALVTGPQQ LENLHQIVTE DLQALEKSVS NLEKSLTSLS
LGVITGVRTG TAALVTGPQQ LENLHQIVTG NLQALKKSVS NLEKSLTSLS
LGVITGVRTG TAALVTGPQQ LENLHQIVTG NLQALKKSVS NLEKSLTSLS
LGTAVGVGTG TAALITGPQQ LEELHAAMTE DLRALEESVS NLEESLTSLS
LGTAVGVGTG TAALITGPQQ LEELHAAMTE DLRALEESVS NLEESLTSLS
LGTAVGVGTG TAALITGPQQ LEELHAAMTE DLRALEESVS NLEESLTSLS
LGTAVGVGTG TAALITGPQQ LEELHAAMTE DLRALEESVS NLEESLTSLS
LGTAVGVGTG TAALITGPQQ LEELHAAMTE DLRALEESVS NLEESLTSLS
LGTAVGVGTG TAALITGPQQ LEELHAAMTE DLRALEESVS NLEESLTSLS
LGTAVGVGTG TAALITGPQQ LEELHAAMTE DLRALEESVS NLEESLTSLS
LGTAVGVGTG TTALITGPQQ LEELHAAMTE DLRALEESVS NLEESLTSLS
LGTAVGVGTG TAALITGPQQ LEELHAAMTE DLRALEEAVS NLEESLTSLS
LGTAVGIGTG TAALITGPQQ LEELHAAMTE DLRALEESVS NLEESLTSLS
LGTAVGVGTG TAALITGPQQ LEELHAAMTE DLRALEESVS NLEESLTSLS
LGTAVGVGTG TAALITGPQQ LEELHAAMTE DLRALEESVS NLEESLTSLS
LGTAVGVGTG TAALITGPQQ LEELHAAMTE DLRALEESVS NLEESLTSLS
LGTAVGVGTG TAALITGPQQ LEELHAAMTE DLRALEESVN NLEESLTSLS
LGTAVGVGTG TAALITGPQQ LEELHAAMTE DLRALEESVN NLEESLTSLS
LGTAVGVGTG TAALITGPQQ LEELHATMTE DL-ALEESVS NLEESLTSLS
LGTAVGVGTG TAALITGPQQ LEELHAAMTE DLRALEESVS NLEETLTSWS
LGTAVGVGTG TAALITGPQQ LEELHAAMTE DLRALEESVS NLEESLTSLS
LGMA-GVGTG TAALITGPQQ LEELHAAMTE DLRALEESVS NLEETLTSWS
LGMA-GVGTG TAALITGPQQ LEELHAAMTE DLRALEESVS NLEETLTSWS
LGMA-GVGTG TAALITGPQQ LEELHAAMTE DLRALEESVS NLEETLTSWS
LGMA-GVGTG TAALITGPQQ LEELHAAMTE DLRALEESVS NLEETLTSWS
LGMA-GVGTG TAALITGPQQ LEELHAAMTE DLRALEESVS NLEETLTSWS
LGMA-GVGTG TAALITGPQQ LEELHAAMTE DLRALEESVS NLEETLTSWS
LGMA-GVGTG TAALITGPQQ LEELHAAMTE DLRALEESVS NLEETLTSWS
LGMA-GVGTG TAALITGPQQ LEELHAAMTE DLRALEESVS NLEESLTSLS
LGMA-GVGTG TAALITEPQQ LEELHAAMTE DLRALEESVS NLEESLTSLS
---------- -------PQQ LEELHAAITE DLRALEESVN NLEESLTSLS
---------- -------PQQ LEELHAAITE DLRALEESVN NLEESLTSLS
---------- -------PQQ LEELHAAITE DLRALEESVN NLEESLISLS
---------- -------PQQ LEELHAAITE DLRALEESVN NLEESLISLS
LGVTAGVGTG AAALITGPQQ LEAPHAAMTE DLRALEESVS NLEESLTPLS
LGVTAGVGTG AAALITGPQQ LEAPHAAMTE DLRALEESVS NLEESLTPLS
LGVTAGVGTG AAALITGPQQ LEAPHAAMTE DLRALEESVS NLEESLTPLS
```

```
LGVTAGVGTG AAALITGPQQ LEAPHAAMTE DLRALEESVS NLEESLTPLS
LGVTAGVGTG AAALITGPQQ LEAPHAAMTE DLRALEESVS NLEESLTPLS
LGVTAGVGTG AAALITGPQQ LEAPHAAMTE DLRALEESVS NLEESLTPLS
LGVTAGVGTG AAALITGPQQ LEAPHAAMTE DLRALEESVS NLEESLTPLS
LGVTAGVGTG AAALITGPQQ LEAPHAAMTE DLRALEESVS NLEESLTPLS
LGVTAGVGTG AAALITGPQQ LEAPHAAMTE DLRALEESVS NLEESLTPLS
LGVTAGVGTG AAALITGPQQ LEAPHAAMTE DLRALEESVS NLEESLTPLS
LGVTAGVGTG AAALITGPQQ LEAPHAAMTE DLRALEESVS NLEESLTPLS
LGVTAGVGTG AAALITGPQQ LEAPHAAMTE DLRALEESVS NLEESLTPLS
LGVTAGVGTG AAALITGPQQ LEAPHAAMTE DLRALEESVS NLEESLTPLS
LGVTAGVGTG AAALITGPQQ LEAPHAAMTE DLRALEESVS NLEESLTPLS
LGVAAGVGTG TAALITGPRQ LEELHRIITE DLQALEKSVN NLEESLTSLS
LGVAAGVGTG TAALITGPRQ LEELHRIITE DLQALEKSVN NLEESLTSLS
LGIAAGVGSG TTALVMGPQQ LEKLHAVITE DLQALEKSIS NLEESLTSLS
LGIAAGVGTG TTALVTEPQQ LEELHAVITE DLQALEKSVS NLEESLTSLS
LGIATGVGTG TTALITGPQQ LEELHAVITE DLQALQKSIS NLEETLTSLS
LGVAAGIGTG STALIKGPID LQSLQIAMDT DLRALQDSVS KLENSLTSLS
LGITAGIGTG STALIKGPID LQSLQIAIDA DLRALQDSVS KLEDSLTSLS
VGIAAGIGTG AVALDSVPRY YNQLRQAMDT DIAALEQSIT KLKESLTSLS
VGIAAGIGTG AVALDSVPRY YNQLRQAMDT DIAALEQSIT KLKESLTSLS
VGIAAGIGTG AVALDSVPCY YNQLRQAMDT DIAALEQSIT KLKESLTSLS
VGIAAGIGTG AAVLDKVPRY YNQLREAMDT DIAALEQSIT KLKESLTSLS
VGIAAGIRIG AAALDKVP-Y YNQLREAMDT DIAALEQSIT KLKESLTALS
IRVAAGVGTG TTALIRQPHY FQELRSAMND NLRALKKSIT KLKESLTSLS
IGVTAGVGTG TTALLGQLHY FQELRSTMSE YLRALEQSIT KLKESLTSLS
IGVTAGVGTG TTALIGQLHY FQELRSTMSE YLRALGQSIT KLKESLTSLS
IGVTAGVGTG TTALVRQPHY FQELRSAMSE DLRALEQSIT KLKESLTSLS
LGVAAGVETG TAALIQAP-Y FNELRIAMDE DLRALEQSNS KPEESLTSLL
LGVAAGVGTG TAALIQAP-Y FNELRIAMYE DLRA-QQSNS KLEESLTSLL
LGVAAGVGTG TAALIKTPQY YEELRAAMDV DLRTIEQSIT KLEESLTSLS
LGVAAGVGTG TAALIKTPQY YEELRAAMDV DLRTIEQSIT KLEESLTSLS
LGVAAGVGTG TAALIKTPQY YEELRAAMDI DLRTIEQSIT KLEESLTSLS
LGVAAGVGTG TAALIKTPQY YEELRAAMDI DLRTIEQSIT KLEESLTSLS
LGVAAGVGTG TAALIQTPRY FEELRTAMDT DLRAIEHSIT KLEESLTSLS
LGVAAGIGTG AAALIQKPYY YNELRAAMDA DLGALEQSIT KLEESLTSLS
LGITAGIGTG TTALIQQPQY YASLRQAVDI DLRALESSIT QLKESLTSLS
LGVATSVGKG TAALIQNPQY FHELKSAIDE DLQAIEQSIS KLEKSLTSLS
IGMAVGVGTG VSALIEGRQG IQSLRDAVNE DLAAIEKSID ALEKSLTSLS
IGMAVGVGTG VSALIEGRQG IQSLRDAVNE DLAAIEKSID TLEKSLTSLS
IGMAVGVGTG VSALIEGRQG IQSLRDAVNE DLATIEKSID ALKKSLTSLS
MGIAAGVGTG TTALVATK-Q FEQLQAAIHT DLGALEKSVS ALEKSLTSLS
MGIAAGVETG TTALVATK-Q FEQLQAAIHT DLGALEKSVS ALEKSLTSLS
MGIAAGVGTG TTALVATK-Q FEQLQAAIHT DLGALEKSVS ALEKSLTSLS
MGIAS--AAS CTALT----- ---------- -------GVS ALEKSLTSLS
```

```
MGIAAGVGTG TTALVATQ-Q FQQLHAAVQD DFKEVEKSIT NLEKSLTSLS
MGIAAGVGTG TTALVATQ-Q FQQLHAAVQD DLKEVEKSIT NLEKSLTSLS
MGIAAGIGTG TTALMATQ-Q FQQLQAAVQD DLREVEKSIS NLEKSLTSLS
VGIAAGVGTG TKALLETA-Q FRQLQMAMHT DIQALEESIS ALEKSLTSLS
LGVAAGVGTG SAALITGPQQ LQQLNSAISE DIEALEKSIS HLEESLTSLS
LGVAAGVGTG AAALVTQHQG FLSLQAAINE DLRDLQNSIE AL-KSLTYLS
LRI-AGAGTG IASLATQQSG ITSLRAAIDE DIERLETSIS HLEKSLTSLS
LGITGATLTG GTGLGVSVHT YHKLSNQLIE DVQALSGTIN DLQDQIDSLA
AGVVARMGTG TTALIQGSQR YEKLRATMDE DLKTIENSIS KLEESLTSLS
AGVVA-MGTG TTALIQGSQR YEKLRAAMDE DLKTIENSIS KLEESLTSLS
LGVAAGVRTG TTALIQGPLH YEKLRAAMDE DLKAIENSIT KLEESLTSLS
LGVAAGVGTR TTALIQGSHH YEKFRAAMDE DLKAIENSIT KLEESLTSLS
LGVAAGVGTG TTALIQGPHH YEKLRAAMDE DLKAIENSIT KLEESLTSLS
SGVAAKVGTG TTALIQGPHH YEKLRAAMDE DLKAIENS-- --EESLTSRS
LGVAARIETG TTALIQGPHH YEKLRAAMYE DLKAIENSIT KLEESLTSLS
```

```
EVVLQNRRGL DLLFLKEGGL CVALKEECCF YVDH
EVVLQNRRGL DLLFLKEGGL CVALKEECCF YVDH
EVVLQNRRGL DLLFLKEGGL CVALKEECCF YVDH
EVVLQNRRGL DLLFLKEGGL CVALKEECCF YVDH
EVVLQNRRGL DLLFLKEGGL CVALKEECCF YVDH
EVVLQNRRGL DLLFLKEGGL CVALKEECCF YVDH
EVVLQNRRGL DLLFLKEGGL CVALKEECCF YVDH
EVVLQNRRGL DLLFLKEGGL CVALKEECCF YVDH
EVVLQNRRGL DLLFLKEGGL CVALKEECCF YVDH
EVVLQNRRGL DLLFLKEGGL CVALKEECCF YVDH
EVVLQNRRGL DLLFLKEGGL CVALKEECCF YVDH
EVVLQNRRGL DLLFLKEGGL CVALKEECCF YVDH
EVVLQNRRGL DLLFLKEGGL YVALKEECCF YVDH
EVVLQNRRGL DLLFLKEGGL CVALKEECCF YVDH
EVVLQNRRGL DLLFLKEGGL CVALKEECCF YVDH
EVVLQNRRGL DLLFLKEEGL CVALKEECCF YVDH
EMVLQNRRGL DLLFLKKGGL CVALKEKCCF YVNH
EVVLQNRRGL DLLFLKEGGL CVALKEECCF YVDH
EVVLQNRRGL DLLFLKEGGL CVALKEECCF YVDH
KVVLQNRRGL DLLFLKKGEL CVALKEECCF YVDH
EVVLQNRRGL DLLFLKEGGL CVALKEKCCF YVDH
EVVLQNRRGL DLLFLKEGGL CVALKEECCF YVDH
EVVLQNRRGL DLLFLKEGGL CVALKEECCF YVDH
EVVLQNRRGL DLLFLKESGL CVALKEECCF YVDH
EVVLQNRRGL DLLFLKESGL CVALKEECCF YVDH
EVVLQNRRGL DLLFLKKGGL CVALKEEC-- WSSH
EVVLQNRRGL DLLFLKKGGL CVALKEEC-- WSSH
EVVLQNRRGL DLLFLKKGGL CVALKEEC-- WSSH
```

EVVLQNRRGL DLLFLKKGGL CVALKEEC-- WSSH
EVVLQNRRGL DLLFLKKGGL CVALKEEC-- WSSH
EVVLQNRRGL DLLFLKKGGL CVALKEEC-- WSSH
EVVLQNRRGL DLLFLKKGGL CVALKEEC-- WSSH
EVVLQNRRGL DLLFLKKGGL CVALKEEC-- WSSH
EVVLQNRRGL DLLFLKKGGL CVALKEECCF YVDH
EVVLQNRRGL DLLFLKKGGL CVALKEEC-- WSSH
EVVLQNRRGL DLLFLKKGGL CVALKEEC-- WSSH
EVVLQNRRGL DLLFLKKGGL CVALKEEC-- WSSH
EVVLQNRRGL DLLFLKEGGL CVALKEECCF YVDH
EVVLQNRRGL DLLFLKEGGL CVALKEECCF YVDH
EVVLQNRRGL DLLFLKEGGL CVALKEECCF YVDH
EVVLQNRRGL DLLFLKEGGL CVALKEECCF YVDH
EVVLQNRRGL DLLFLKEGGL CVALKEECCF YVDH
EVVLQNRRGL DLLFLKEGGL CVALKEECCF YVDH
EVVLQNRRGL DLLFLKEGGL CVALKEECCF YVDH
EVVLQNRRGL DLLFLKEGGL CVALKEECCF YVDH
EVVLQNRRGL DLLFLKEGGL CVALKEECCF YVDH
EVVLQNRRGL DLLFLKEGGL CVALKEECCF YVDH
EVVLQNRRGL DLLFLKERGL CVALKEECCF YVDH
EVVLQNRRGL DLLFLKERGL CVALKEECCF YVDH
EVVLQNRRGL DLLFLKEGGL CVALKEECCF YVDH
EVVLQNRRGL DLLFLKEGGL CVALKEECCF YVDH
EVVLQNRRGL DLLFLKEGGL CVALKEECCF YVDH
EVVLQNRRGL DLLFLKEGRL CVALKEECCF YVDH
EVVLQNKKGL DLLFLKKRRL CVALKEKCCF YVDH
EVVLQNKKGL DLLFLKKRRL CVALKEKCCF YVDH
EVVLQNRRGL DLLFLREGGL CAALKEECCF YVDH
EVVLQNRRGL DLLFLREGGL CAALKEECCF YVDH
EVVLQNRRGL DLLFLREGGL CAALKEECCF YVDH
EVVLQNRRGL DLLFLREGGL CAALKEECCF YVDH
EVVLQNRRGL DLLFLREGGL CAALKEECCF YVDH
EVVLQNRRGL DLLFLREGGL CAALKEECCF YVDH
EVVLQNRRGL DLLFLREGGL CAALKEECCF YVDH
EVVLQNRRGL DLLFLREGGL CAALKEECCF YVDH
EVVLQNRRGL DLLFLREGGL CAALKEECCF YVDH
EVVLQNRRGL DLLFLREGGL CAALKEECCF YVDH
EVVLQNRRGL DLLFLREGGL CAALKEECCF YVDH
EVVLQNRRGL DLLFLREGGL CAALKEECCF YVDH
EVVLQNRRGL DLLFLREGGL CAALKEECCF YVDH
EVVLQNWRGL DLLFLKEGGL CAALREECCF YVDH
EVVLQNWRGL DLLFLKEGGL CAALREECCF YVDH
EVVLQNRRGL DLLFLREGGL CAALKEECCF YVDH
EVVLQNQRGL DLLFLKEGGL CAALKEE-CF YVDH

EVVLQNRRGL DLLFLREGGL CAALKEECYF YVDH
EVVLQNQRGL DLLFLKEGGL CAALKEE-CF YVDH
EVVLQNQRGL DLLFLKEGGL CAALKEE-CF YVDH
EVVLQNQRGL DLLFLKEGGL CAALKEE-CF YVDH
EVVLQNQRGL DLLFLKEGGL CAALKEE-CF YVDH
EVVLQNQRGL DLLFLKEGGL CAALKEE-CF YVDH
EVVLQNQRGL DLLFLKEGGL CAALKEE-CF YVDH
EVVLQNQRGL DLLFLKEGGL CAALKEE-CF YVDH
EVVLQNRTGG DIWFLKEGGL CAALKEECCF YADH
EVVLQNWRGL DLVFLKEGGL CAALKEECCF YADH
EVVLQNWRGL DLLFLKEGGL CAALRKECCF YVDH
EVVLQNWRGL DLLFLKEGGL CAALREECCF YVDH
EVVLQNWRGL DLLFLKEGGL CAALREECCF YVDH
EVVLQNWRGL DLLFLKEGGL CAALREECCF YVDH
EVVLQNRRGL DLLFLKEGGL CAALKEECCF YVDH
EVVLQNRRGL DLLFLREGGL CAALKEECCF YVDH
EVVLQNRRGL DLLFLKEGGL CAALKEECCF YIDH
EVVLQNRRGL DLLFLKEGGL CAALKEECCF YVDH
EVVLQNRRGL DLLFLKEGGL CAALKEECCF YVDH
EVVLQNRRGL DLLFLKEGGL CAALKEECCF YVDH
EVVLQNRRGL DLLFLKEGGL CAALKEECCF YVDH
EVVLQNRRGL DLLFLKEGGL CAALKEECCF YVDH
EVVLQNRRGL DLLFLKEGGL CAALKEECCF YVDH
EVVLQNRRGL DLLFLKEGGL CAALKEECCF YVDH
EVVLQNRRGL DLLFLKEGGL CAALKEECCF YVDH
EVVLQNRRGL DLLFLKEGGL CAALKEECCF YVDH
EVVLQNRRGL DLLFLXEGGL CAALKEECCF YVDH
EVVLQNRRGL DLLFLKEGGL CAALKEECCF YVDH
EVVLQNRRGL DLLFLKKGGL CVALKEECCF YVDH
EVVLQNRRGL DLLFLKKGGL CVALKEECCF YVDH
EVVLQNRRGL DLLFLKEGGL CAALKKECCF YVVH
EVVLQNWRGL DLLFLKEGGL CAALKEECCF YVDH
EVVLQNWREL DLLFLKEGGL CAALKEECCF YVDH
EVVLQNRRGL DLLFLKEGGL CAALKEECCF YVDH
EVVLQNRRGL DLLFLKEGGL CAALKEECCF YIDH
EVVLQNRRGL DLLFLKEGGL CAALKEECCF YVDH
EVVLQNRRGL DLLFLKEGGL CAALKGECCF YVDH
EVVLQNRRGL DLLFLKEGGL CAALKEECCF YVDH
KVVLQNRRGL DLLFLKEGGL CAALKEECCF YVDH
EVVLQNHRGL DLLFLKEGGL CAALKEECCF YVDH
EVVLQNRRGL DLLFLKERGL CAALKEECYF YIDH
EVVLQNRRGL DLLFLKEGGL CVALKKECCF YKDH
ELVLQNRRGL DLLFLKEGGL CVALKKECCF YKDH
EVVLQNRRGL DLRFLKEGGL CVAIKKECCF YTDH

```
EVVLQNRRGS DFL---DRGL CDALREECCF YVDH
EVVSQNRRES DLLFL-DRGL CNALRQECCF YVDH
EVVLQNRRGL DLLFLKEGGL CAALKEECCF YVDH
EVLLQNRRGL DLLFLKEGGL CAALKEECCF YVDH
EVVLQNRRGL DLLFLKEGGL CAALKEECCF YVDH
EVVLQNRRGL DLLFLKEGGL CAALKEECCF YVDH
EVVLQNRRGL DLLFLKEGGL CAALKEECCF YVDH
EVVLQKRRGL DLLFLKEGGL CAALKEECCF YVDH
EVVLQNRRGL DLLFLKEGGL CAALKEECCF YIDH
EVVLQNRRRL DLLFLKEGGL CAAIREECCF YVDH
EVVLQNRRGL DLLFLKEGGL CAALKEECCF YADH
EVVLQNRRGL DLLFLKEGGL CAALKEECCF YADH
EVVLQNRRGL DLLFLKERGL CAALKEECCF YADH
EVVLQNRRGL DLLFLKEGGL CAALKEECCF YADH
EVVLQNRRGL DLLFLKEGGL CAALKEKCCF YADH
EVVLQNRRGL DLLFLKEGGL CAALKEECCF YADH
EVVLQNRRGL DLLFLKKGRL CAALKEECCF YADH
EVVLQNRRGL DLLFLKEGGL CAALKEECCF YADH
EVVLQNRRGL DLLFLKEGGL CAALKEECCF YADH
EVVLQNRRGL DLLFLKEGGL CAALKEECCF YADH
EVVLQNRRGL DILFLQEGGL CAALKEECCF YADH
EVVLQNRKGL DLLFLKEGGL CAALREECCF YIDH
EVVL-NRRGL DLLFLKEGGL CAALKEECCF YADH
EVVLQNRRGL DLLFLQQGGL CVALGEECCF YADH
EVVLQNRRGL DLLTAEQGGI CLALQEKCCF YANK
EVVLQNRRGL DLLFLKDGGL RAAIKEECCF YVDH
EVVLQNRRGL DLLFLKDGGL CAARKEECCF YVDH
EVVLQNRRGL DLLFLKDGGL CAAIKEECCF YVDH
EVVLQNRRGL DLLFLKDRGL CAAIKEECCF YVDH
EVVLQNRRGL DLLFLKDGGL CAAIKEECCF YVDH
EVVLQNRRGL DLSSLKDGGL CAAIKEECCF YVDH
EVVLQNRRGL DLLFLKDGGL CAAIKEECCF YVNH
```
